# Loss of cyclin A2 in murine colonic epithelial cells disrupts colon homeostasis by triggering DNA damage and dysplasia and high cyclin A2 expression is a good-prognosis factor in patients with colorectal cancer

**DOI:** 10.1101/690404

**Authors:** Yuchen Guo, Monica Gabola, Rossano Lattanzio, Conception Paul, Valérie Pinet, Ruizhi Tang, Hulya Tulari, Julie Bremond, Chloé Maurizy, Quentin Da Costa, Pascal Finetti, Florence Boissière-Michot, Céline Lemmers, Séverine Garnier, François Bertucci, Rania Azar, Jean-Marie Blanchard, Piotr Sicinski, Emilie Mamessier, Bénédicte Lemmers, Michael Hahne

## Abstract

To clarify the function of cyclin A2 in colon homeostasis and colorectal cancer (CRC) we generated mice deficient for cyclin A2 in colonic epithelial cells (CEC). Colons of those mice displayed architectural changes in the mucosa, and signs of inflammation as well as an increased proliferation of CEC associated with the appearance of low- and high-grade dysplasia. The main initial events triggering those alterations in cyclin A2 deficient CEC appear to be abnormal mitoses and DNA damage. Cyclin A2 deletion in CEC promoted the development of dysplasia and adenocarcinomas in the murine colitis-associated cancer model. We next explored the status of cyclin A2 expression in clinical CRC samples at the mRNA and protein level and found higher expression in tumors of stage I and II patients compared to those of stage III and IV. A meta-analysis of 11 transcriptome datasets comprising 2,239 primary CRC tumors displayed different *CCNA2* (the mRNA coding for cyclin A2) expression levels among the CRC tumor subtypes with highest in CMS1 and lowest in CMS4. Moreover, high expression of *CCNA2* was found to be a good prognosis factor independently from other prognostic factors for the CMS1, CMS3 and CMS4 subtypes.

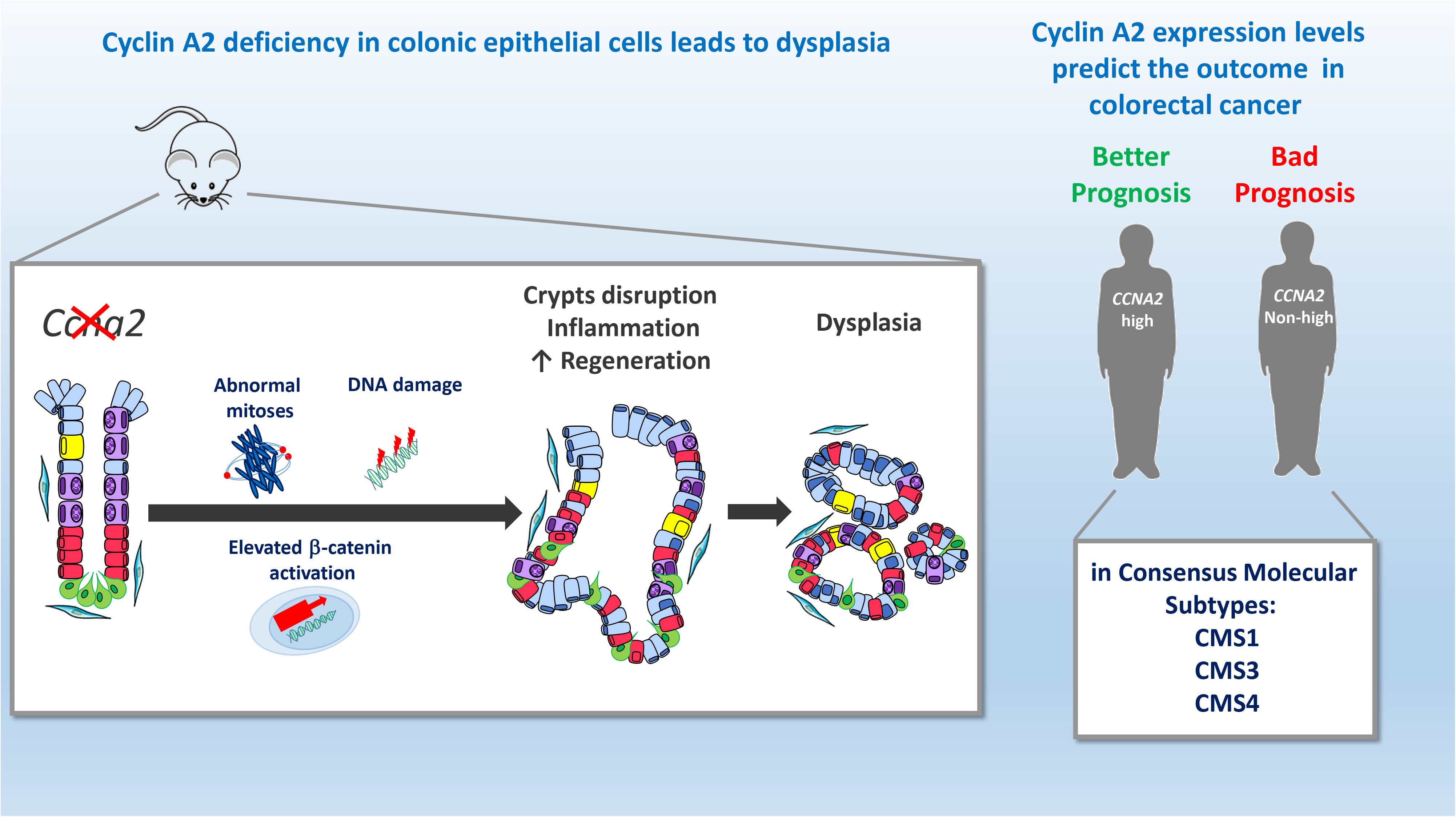

---

Colorectal cancer (CRC) is one of the leading causes of malignancy-related death worldwide with increasing incidence (International Agency for Research on Cancer, WHO; http://gco.iarc.fr/today/fact-sheets-cancers). Disease status of patients can be classified according to histologic tumor grade (differentiation) and anatomic extent of disease (*i.e.*, (TNM, tumors/nodes/metastases, stages I–IV), which describes tumor growth into the wall of rectum or colon (stage II), involvement of regional lymph nodes (stage III) and metastatic spread to other organs (stage IV)(2). Once metastasis has developed, 5-year survival rates following surgery are below 10% (3). Also, adjuvant chemotherapy after surgical resection for stage III CRC and high-risk stage II tumor (*i.e*., tumors with poor differentiation, T4 classification, lymphovascular invasion, perineural invasion) increased the patients survival rates, but many patients will relapse or develop distant metastases (4, 5). Thus, there is an urgent need to better understand the molecular changes leading to the development of CRC to identify novel therapeutic strategies.

An important risk factor for CRC development is inflammation. Early-onset of chronic inflammatory bowel disease and widespread manifestation (i.e., pancolitis) have a higher probability of developing CRC (6). Besides, inherited mutations have been identified for 5% of CRC patients, with diseases traditionally divided into polyposis syndromes, such as the Familial Adenomatous Polyposis (FAP), and non-polyposis syndromes like the Lynch Syndrome, also known as Hereditary Non-Polyposis Colorectal Cancer (HNPCC) (7). Sporadic CRC is a molecularly highly heterogeneous disease. Research teams from different institutions performed recently a meta-analysis on the different reported sub-classifications to define the existence of four distinct gene expression–based subtypes (Consensus Molecular Subtypes or CMS), i.e. immune (CMS1), canonical (CMS2), metabolic (CMS3) and mesenchymal (CMS4) subtype (8). Each CMS has a unique biology and gene expression pattern (8) providing now a frame for designing subtype-specific and personalized treatments. Indeed, CMS display different clinical features including prognosis and response to therapies, with CMS4 having the worst prognosis.

Cyclin A2 is an established regulator of cell proliferation and has been used for molecular diagnostics as a proliferation marker (9). However, several studies suggest that cell cycle regulators can exert additional functions, well beyond cell cycle control (10–12, 12–16). For example, cyclin-CDK inhibitors, p21, p27 and p57, as well as cyclin D1, have been involved in the control of apoptosis, transcription, cytoskeletal dynamics as well as cell migration (12, 13, 13, 16)

We described a role for cyclin A2 in the regulation of cell migration and invasion (17–19). More specifically, we found that deletion of cyclin A2 in a murine mammary gland epithelial cell line (NMuMG) induces epithelial to mesenchymal transition (EMT) and increases invasiveness *in vitro* as well as *in vivo* in the avian embryo model (18, 19). This concurs with a recent report that identified cyclin A2 as a key regulatory component in EMT and metastasis of lung cancer cells by modulating the TGFbeta signaling pathway (20).

Few studies investigated the role of cyclin A2 in tumor development *in vivo* using genetically altered mice. One study described the generation of mutant mice with reduced cyclin A2 expression. These mice displayed spontaneous tumor development, in particular in the lung, and were also more susceptible to chemically-induced skin cancer (21). On the other hand, tissue-specific cyclin A2 inactivation in the liver was shown to impair hepatocellular carcinoma development and recapitulated the phenotype observed in absence of cyclin-dependent kinase 2 (CDK2) (22). These observations point to a tissue-specific role of cyclin A2.

We previously reported lower cyclin A2 expression in hepatic metastases compared to primary tumors derived from the same CRC patients (17). This prompted us to investigate the role of cyclin A2 in colon homeostasis and carcinogenesis using cell type-specific knockout mice.

## Results

### Generation of mice deficient for cyclin A2 in the colonic epithelium

We established two mouse models to study the function of cyclin A2 in colon homeostasis and colorectal cancer development by employing the previously established conditional cyclin A2 (*Ccna2fl/fl*) strain that is based on the Cre/LoxP system (23). Constitutive deletion of cyclin A2 in the intestinal epithelium was obtained by crossing cyclin A2 conditional knock-out (KO) mice with the *villin-Cre* mouse strain (*VilCre*). Expression of Cre under the villin promoter was shown to be active in intestinal epithelial cells (24). In addition, we crossed *Ccna2fl/fl* mice with the inducible *Cre* mouse strain, *villin-CreERT^2^ (VilCreERT2)*, in which *Ccna2* is excised upon intraperitoneal injection of tamoxifen (24).

Efficient inactivation of cyclin A2 in purified colonic epithelial cells (CEC) of *VilCreCcna2fl/fl* mice was confirmed in a RNA sequencing experiment validating the deletion of exons 2 to 7 of the *Ccna2* gene (Supplemental Fig 2A) as well as WB analysis (Fig 1A, left panel). Two tamoxifen injections at day 0 and 2 were sufficient to abrogate cyclin A2 expression in *VilCreERT2Ccna2fl/fl* mice (Fig 1A, right panel).

**Figure 1:**
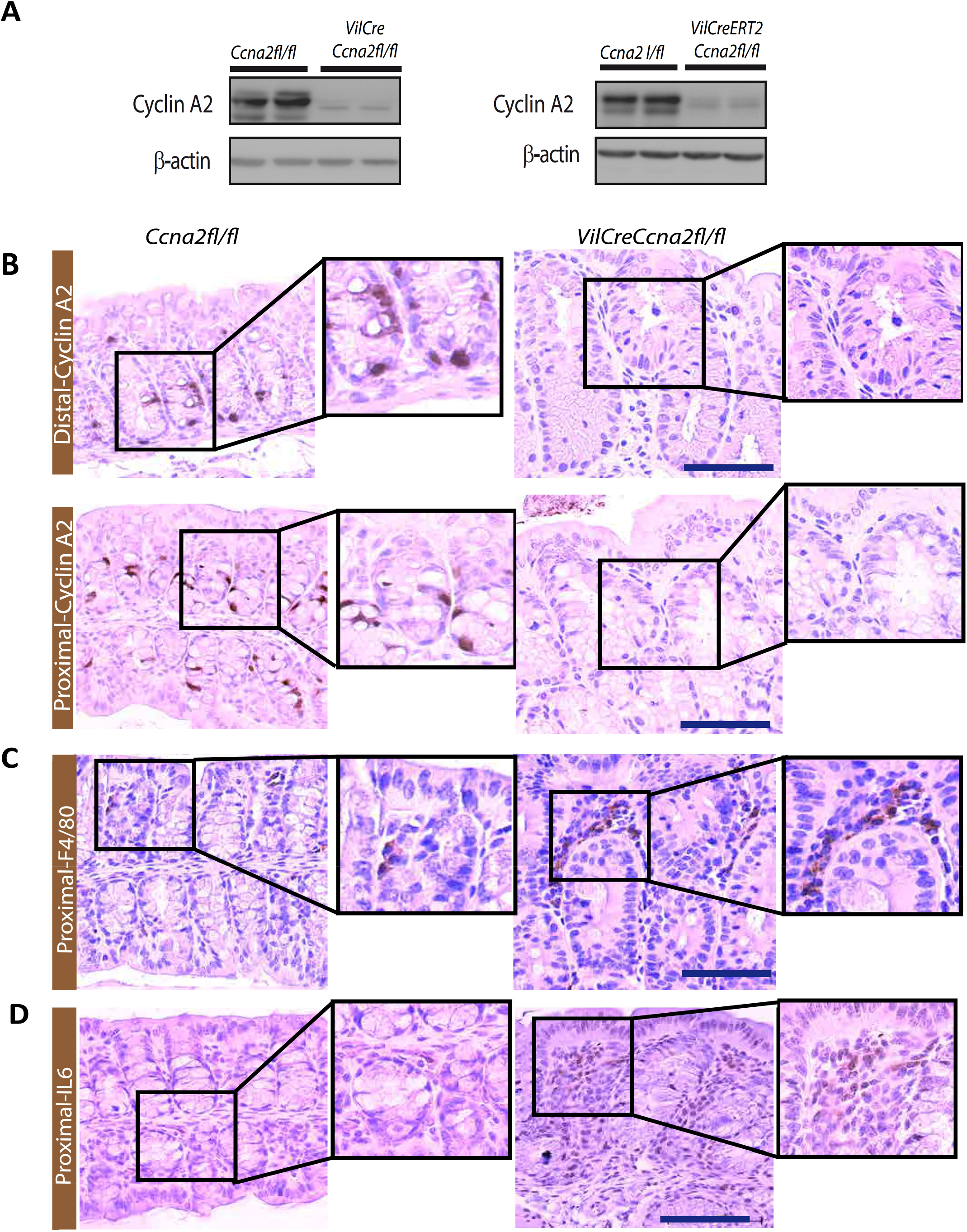
Cyclin A2 deletion leads to epithelial changes in the colon mucosa and inflammation. **(A).** Cyclin A2 depletion at the protein level in colonic epithelial cells (CEC) of constitutive (*VilCreCcna2fl/fl*, left panel) and tamoxifen-induced knockout mice (*VilCreERT2Ccna2fl/fl*, right panel) by comparison to control animals (*Ccna2fl/fl*) displayed by Western Blot analysis. (**B, C, D).** Immunostaining for cyclin A2 (**B**), F4/80 (**C**) and IL-6 (**D**) of the indicated parts of colon derived from control (*Ccna2fl/fl*), and constitutive (*VilCreCcna2fl/fl*) cyclinA2-deficient mice. Blow Up: 2.5X. Scale bars: 100μm.

**Figure 2:**
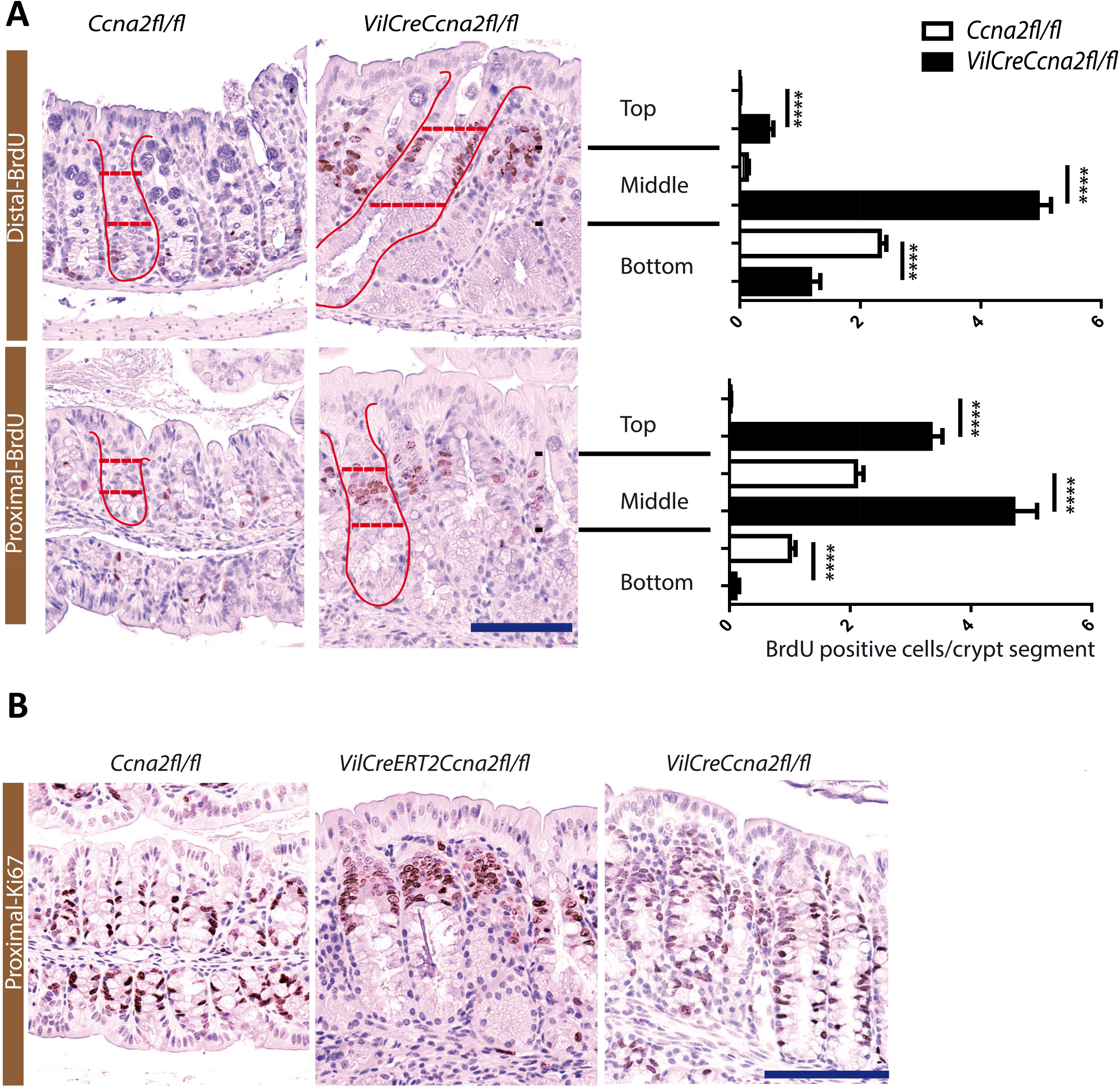
Cyclin A2 depletion in colonic epithelial cells induces cell proliferation. **(A).** Analysis of cell proliferation in colonic crypts of control (*Ccna2fl/fl*) and cyclinA2-deficient (*VilCreCcna2fl/fl*) mice by BrdU incorporation. Left panel: Representative images of immunostainings for BrdU from the distal and proximal parts of the colons taken from control (*Ccna2fl/fl*) and cyclinA2-deficient (*VilCreCcna2fl/fl*) mice 2 hours after intraperitoneal injection of the nucleoside analog. Right panel: Quantification of the numbers of BrdU positive cells per crypt segment. Crypts (depicted in red) were subdivided in 3 parts (dotted lines) for the analysis (bottom, middle and top, n=120 crypts for the distal and n=110 for the proximal part from 3 different mice; mean values ±SEM are provided, **\*\*\*\*** p<0.0001; two-tailed unpaired t-test). **(B).** Representative images for Ki67 immunostaining in the proximal part of the colon from control (*Ccna2fl/fl)* animals, induced knockout mice 8 days after tamoxifen treatment (*VilCreERT2Ccna2fl/fl)* and constitutive knockout mice (*VilCreCcna2fl/fl).* Scale bars: 100μm.

We also confirmed cyclin A2 loss in colonic crypts of *VilCreCcna2fl/fl* mice by immunohistochemistry (Fig 1B). For this, we analyzed the distal region, close to the anus, as well as the proximal region, close to the caecum (Supplemental Fig 1A), since biological and clinical differences for both segments are well established (25). In both regions we detected cyclin A2 expression in epithelial cells of the lower part of the crypts in control mice, whereas cyclin A2 expression was undetectable in colonic crypts of cyclin A2 KO mice (Fig 1B).

### Cyclin A2 depletion in the colon induces epithelial changes in the mucosa and inflammation

During the analysis of cyclin A2 expression we noted epithelial changes in the colonic crypts as well as architectural changes in the mucosa of cyclin A2-deficient mice, but not in control animals (Fig 1B and Supplemental Fig 1C). We also detected immune cell infiltration in cyclin A2-deficient colons. The immune cell infiltrates exhibited morphological characteristics of macrophages and were immunostained using the macrophage marker F4/80. This revealed an increased number of F4/80-positive cells, about 2-fold in constitutive (Fig 1C and Supplemental Fig 1B, left panel) and 1.5-fold in induced (Supplemental Fig 1B, right panel) cyclin A2-deficient colons versus normal ones. Macrophages can be important producers of the pro-inflammatory cytokine IL6 (26) and in fact IL6 was detectable in the stroma of cyclin A2-deficient colons, but not in those of control mice (Fig 1D).

Finally, about 15% males and 10% females of *VilCreCcna2fl/fl* mice developed rectal prolapses within the first three months of age, which can be considered as an indicator of inflammation (27). Taken together, cyclin A2 deficiency in colonic epithelial cells induces epithelial and architectural changes of the colonic crypts and different signs of inflammation.

### RNA sequencing analysis reveals a prominent altered expression of genes involved in cell cycle, mitotic process, chromosome segregation and DNA double-strand break repair

To gain insights into the molecular traits of *Ccna2* mutant mice, RNA sequencing was performed using CEC isolated from the proximal colon of *Ccna2fl/fl* mice (n=4) and *VilCreCcna2fl/fl* mice (n = 4) at four weeks of age. At this age *VilCreCcna2fl/fl* mice already display visible distortion of the crypt architecture that was validated by IHC analysis for the distal region of colons employed for this experiment.

The supervised analysis of the RNAseq data identified 469 upregulated and 640 downregulated genes in *VilCreCcna2fl/fl mice* samples *versus Ccna2fl/fl* mice samples (fold change > 2, adjusted p-value |FDR| >0.05) (Table 1). Pathways analyses of the most altered genes revealed that the most significant biological functions associated with the genes upregulated in *VilCreCcna2fl/fl mice* were related to cell cycle (nuclear division and mitosis, regulation of mitotic cell cycle, p<2,79e^-03^-1,15e^-12^), cellular assembly and organization (chromosome segregation, chromatin segregation, p < 3,01e^-03^-1,80e^-09^), as well as DNA replication, recombination and repair (double-strand break repair and homologous recombination, p<3,01e-^03^-1,80e^-09^) (Table 1, Supplemental Table 4 and Supplemental Fig 2B). In line with this, the analysis highlighted increased levels of *Mki67* (coding for Ki-67), *Brca1, Brca2, Rad18, Rad51, Blm, Exo1* or *Rpa1* (genes involved in the double-strand break repair mechanism) as well as *Cdc45, Bub1, Mad2 and Rad21* (genes related to checkpoint controls of the cell cycle and to abnormal mitosis) (Supplemental Fig 2C).

**Table 1:**
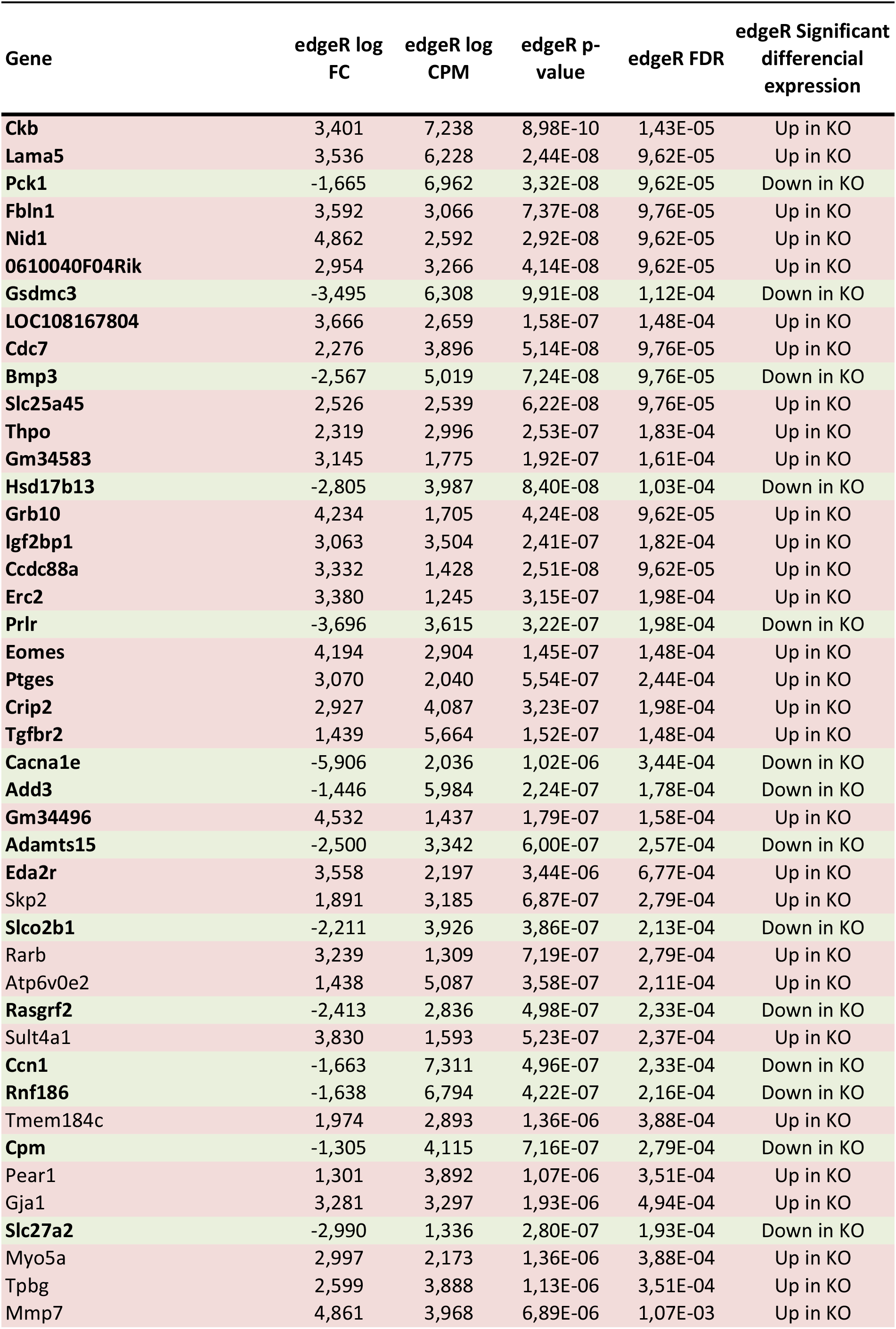
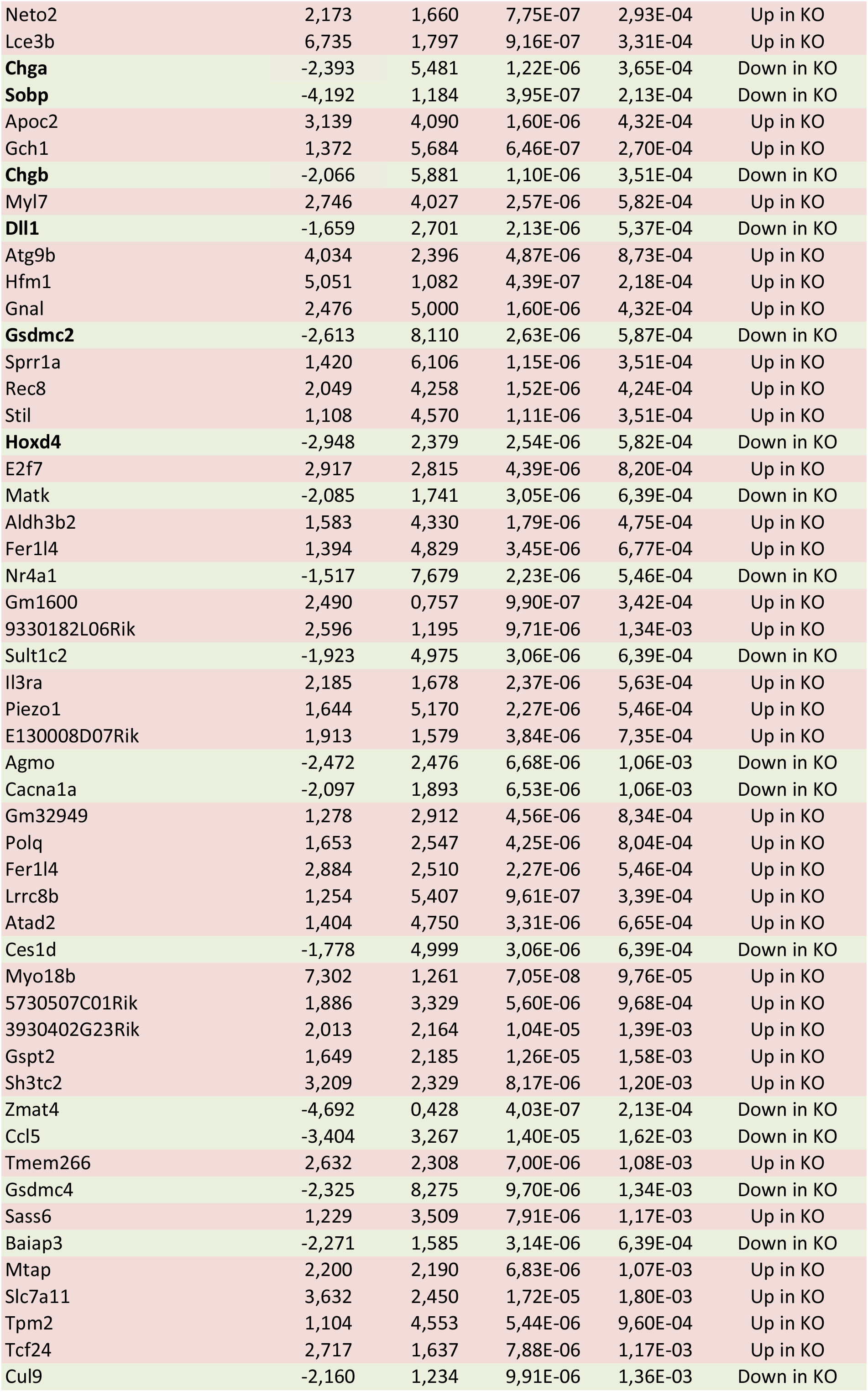
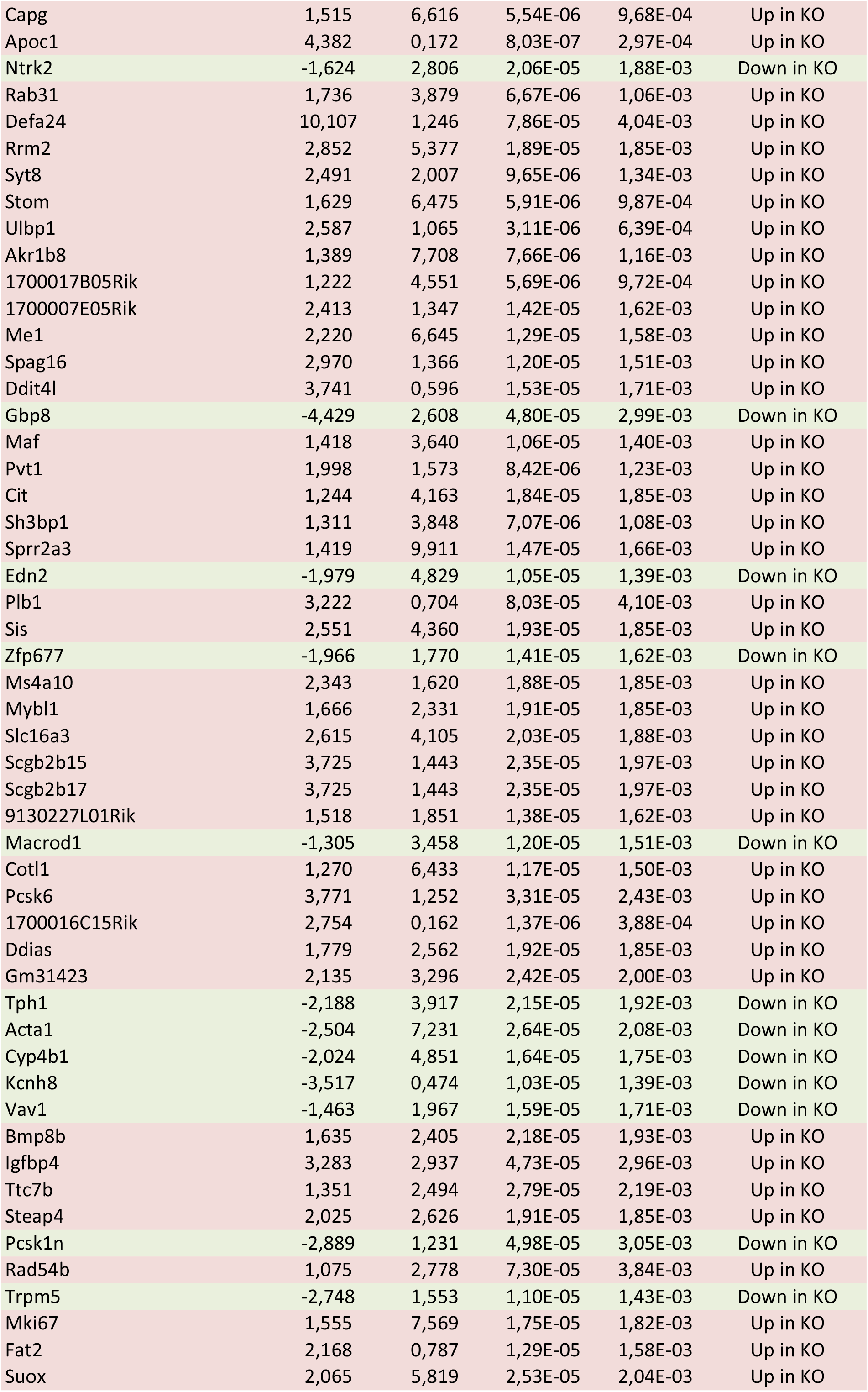
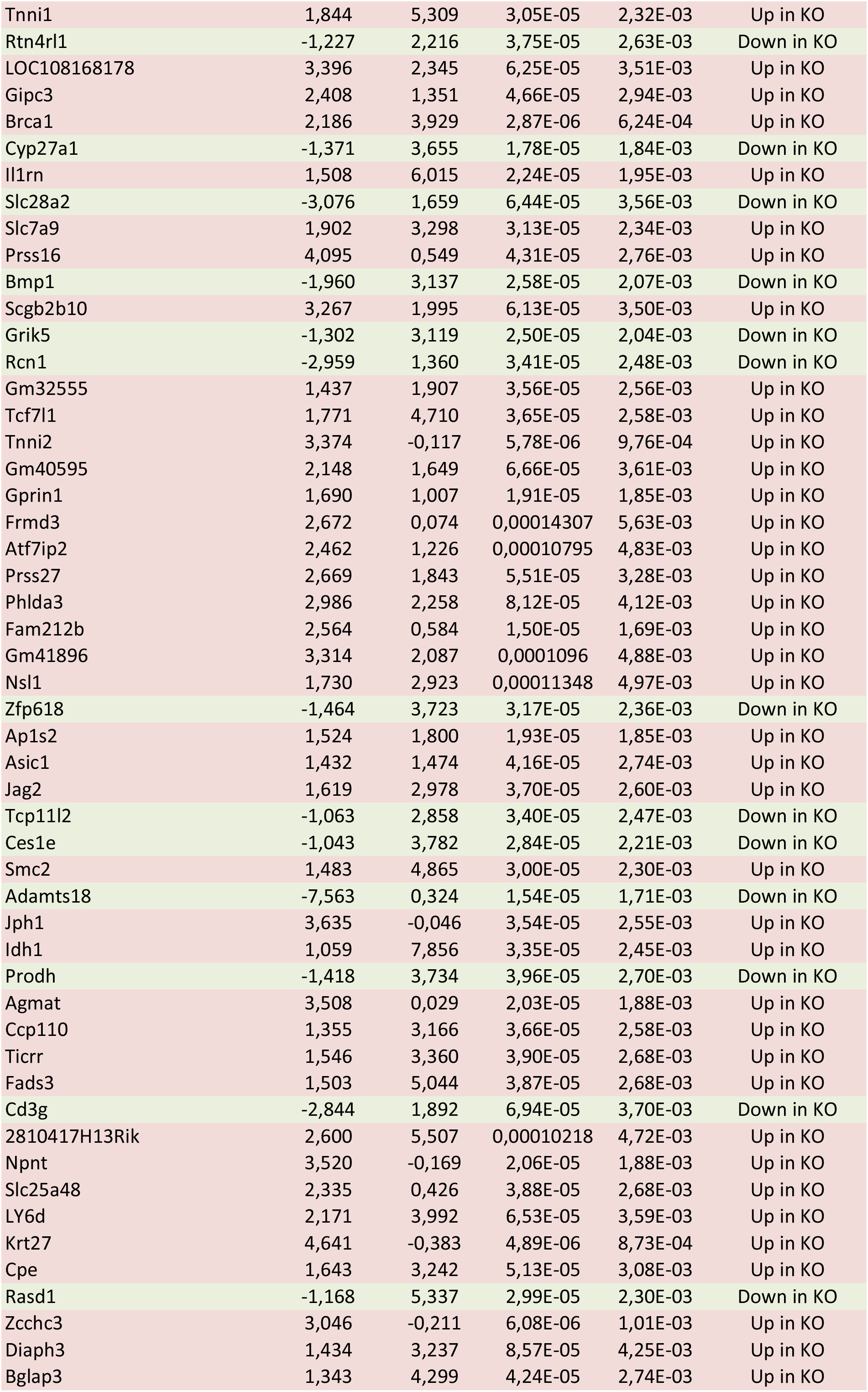
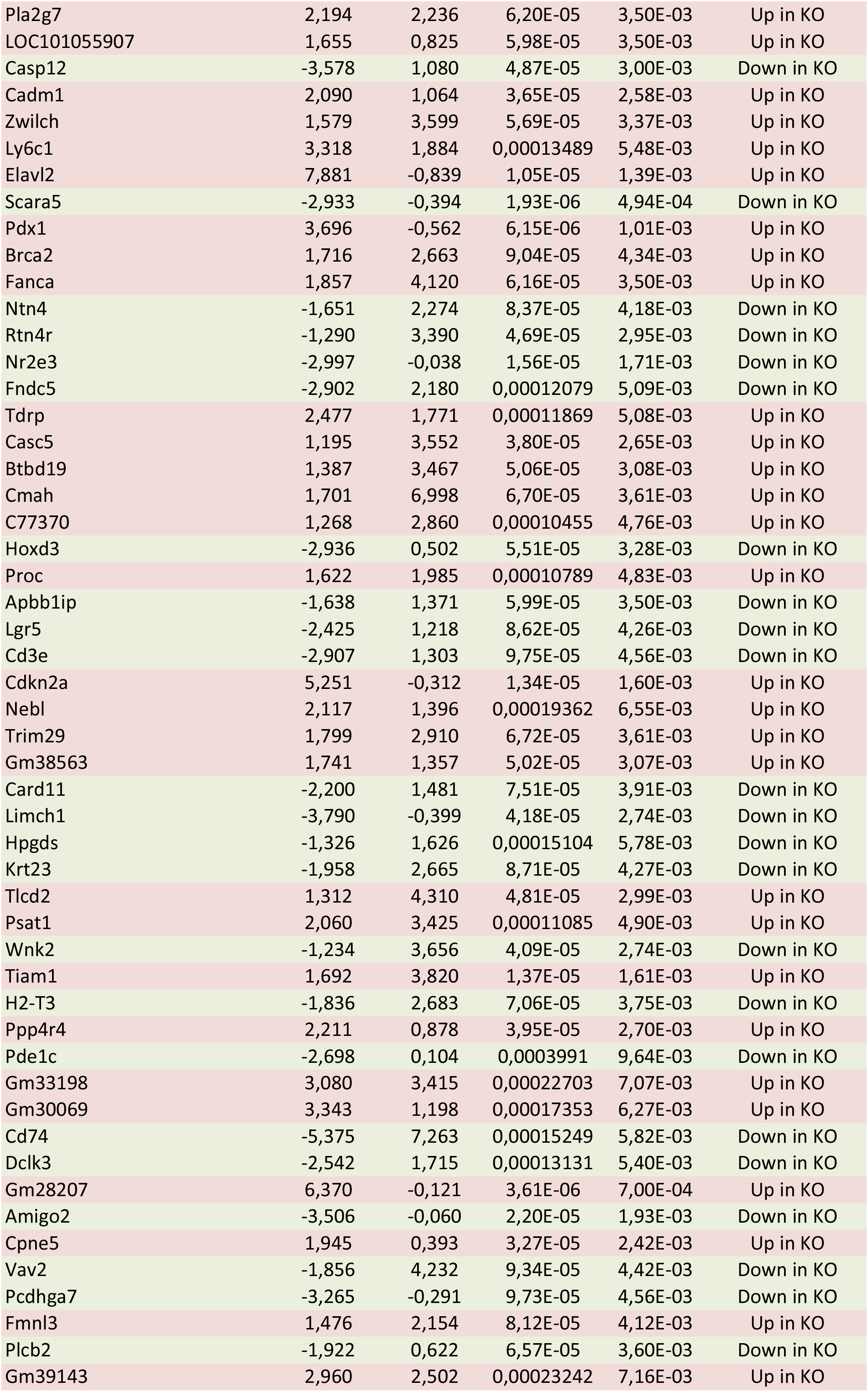
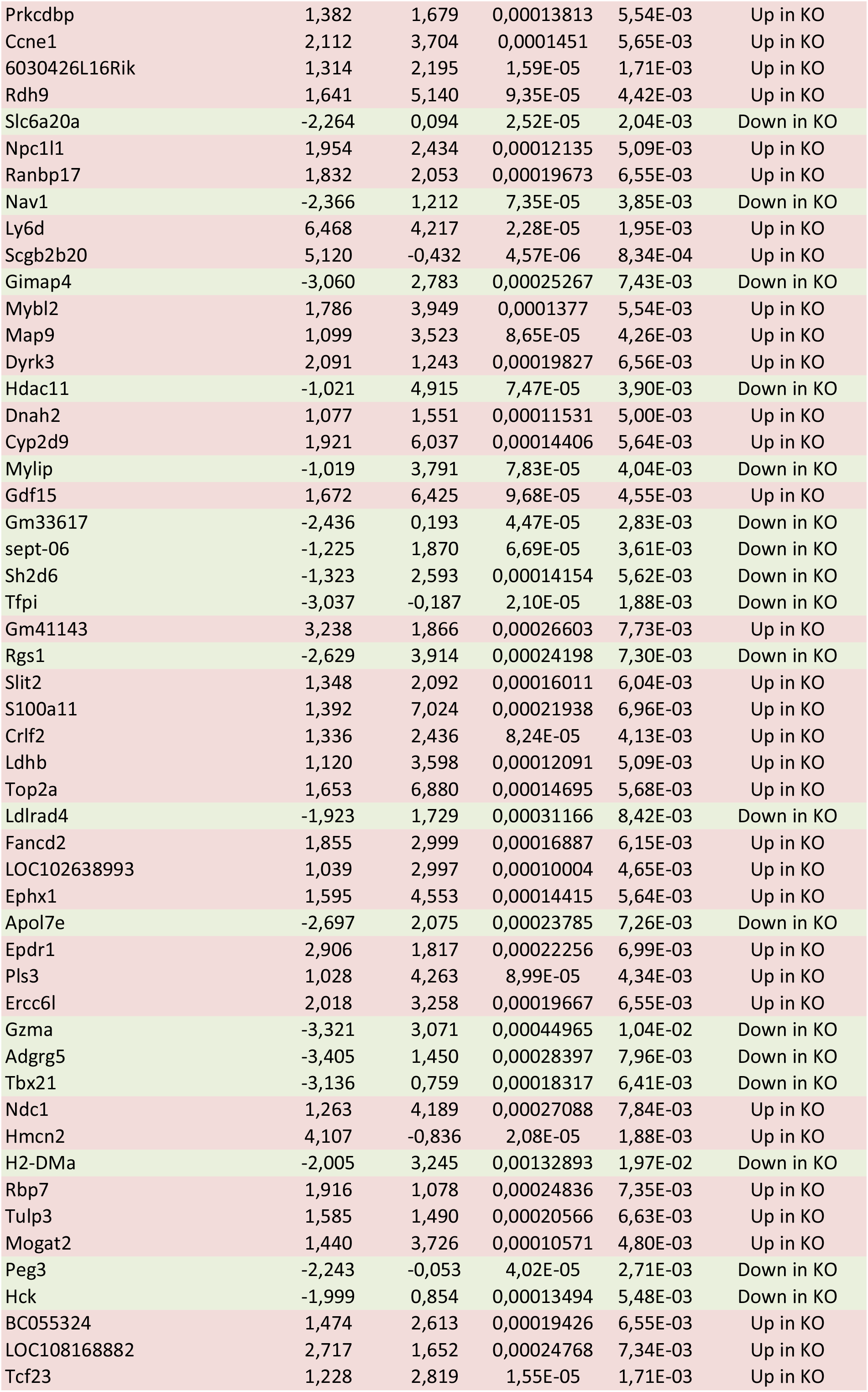
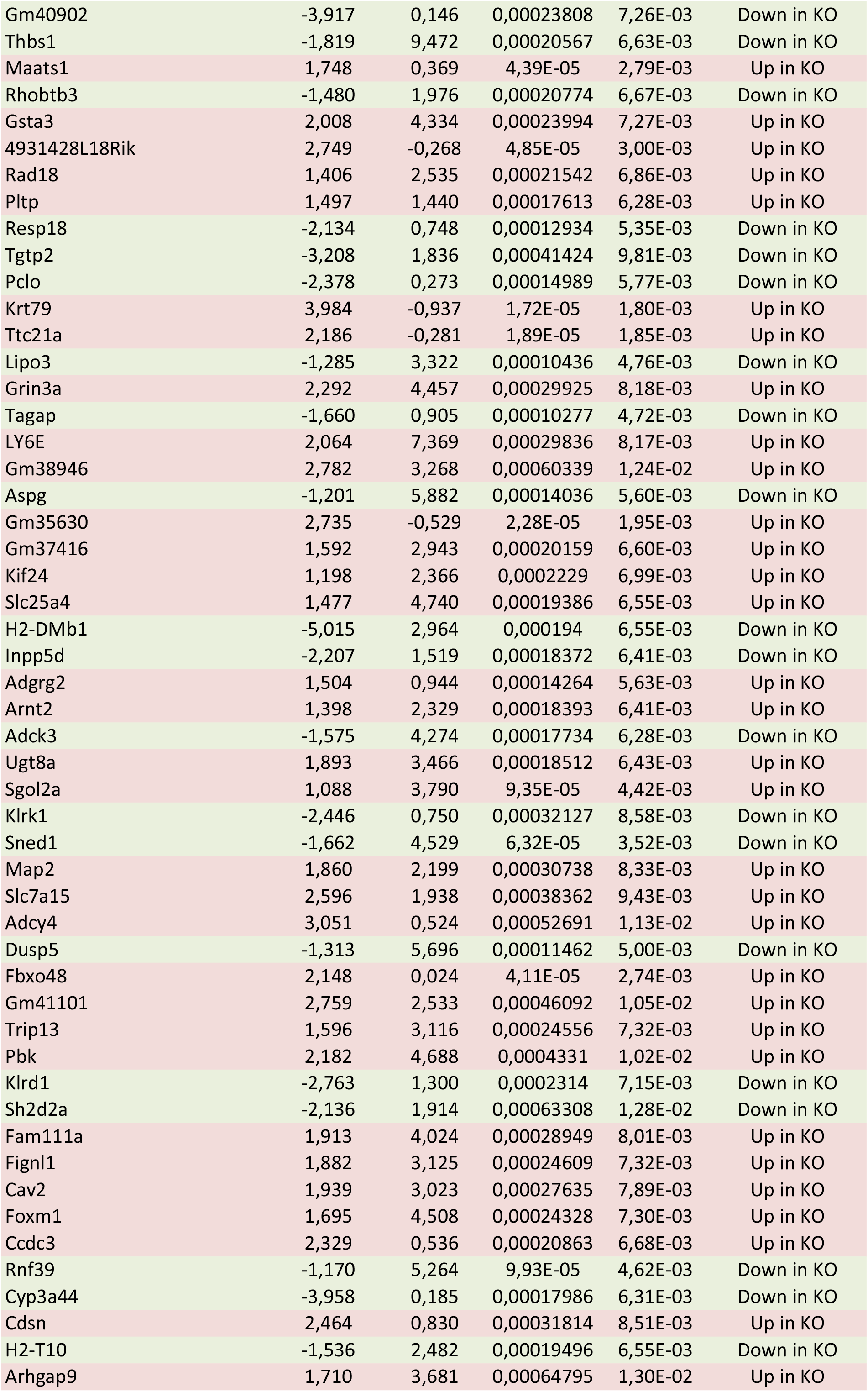
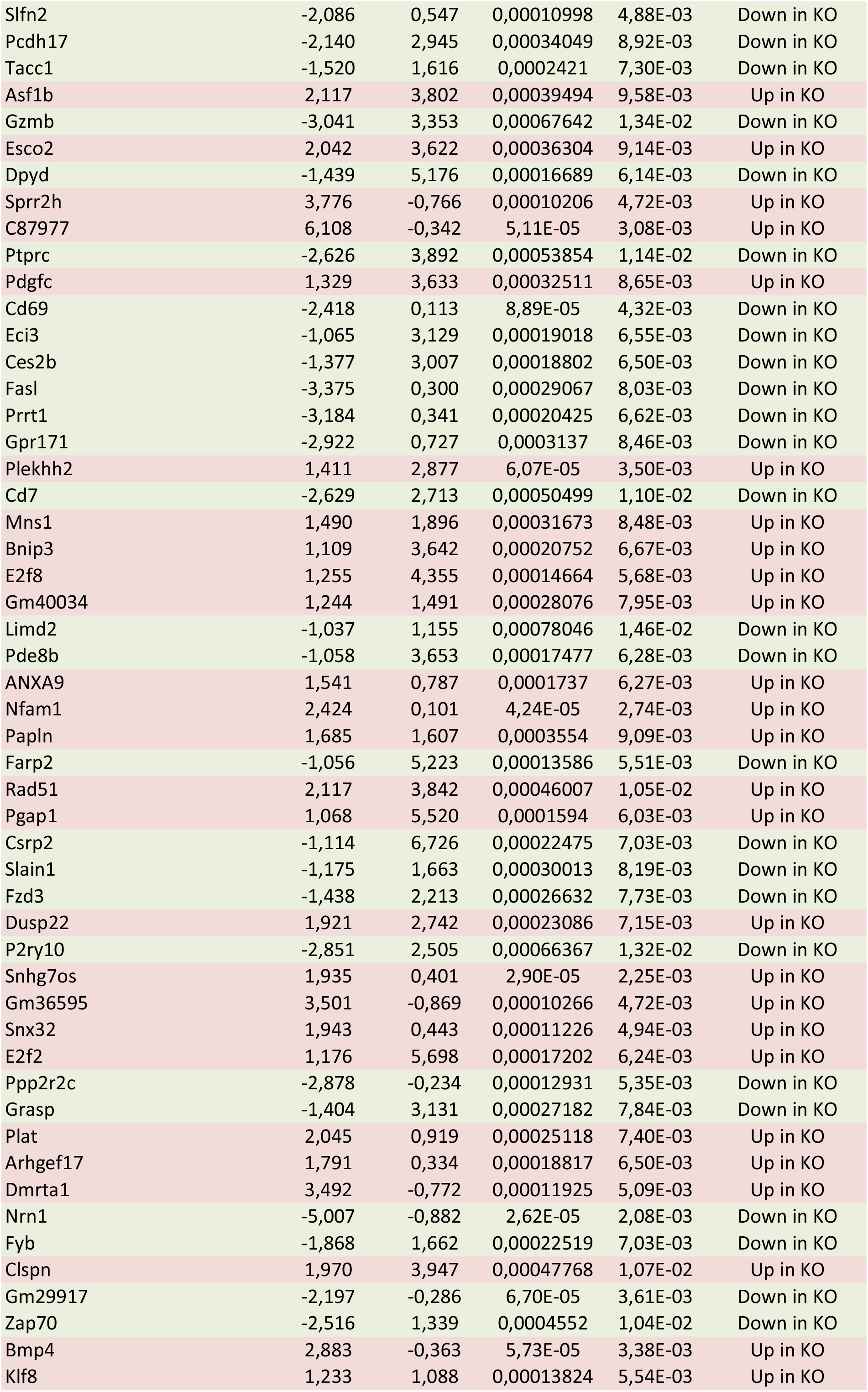
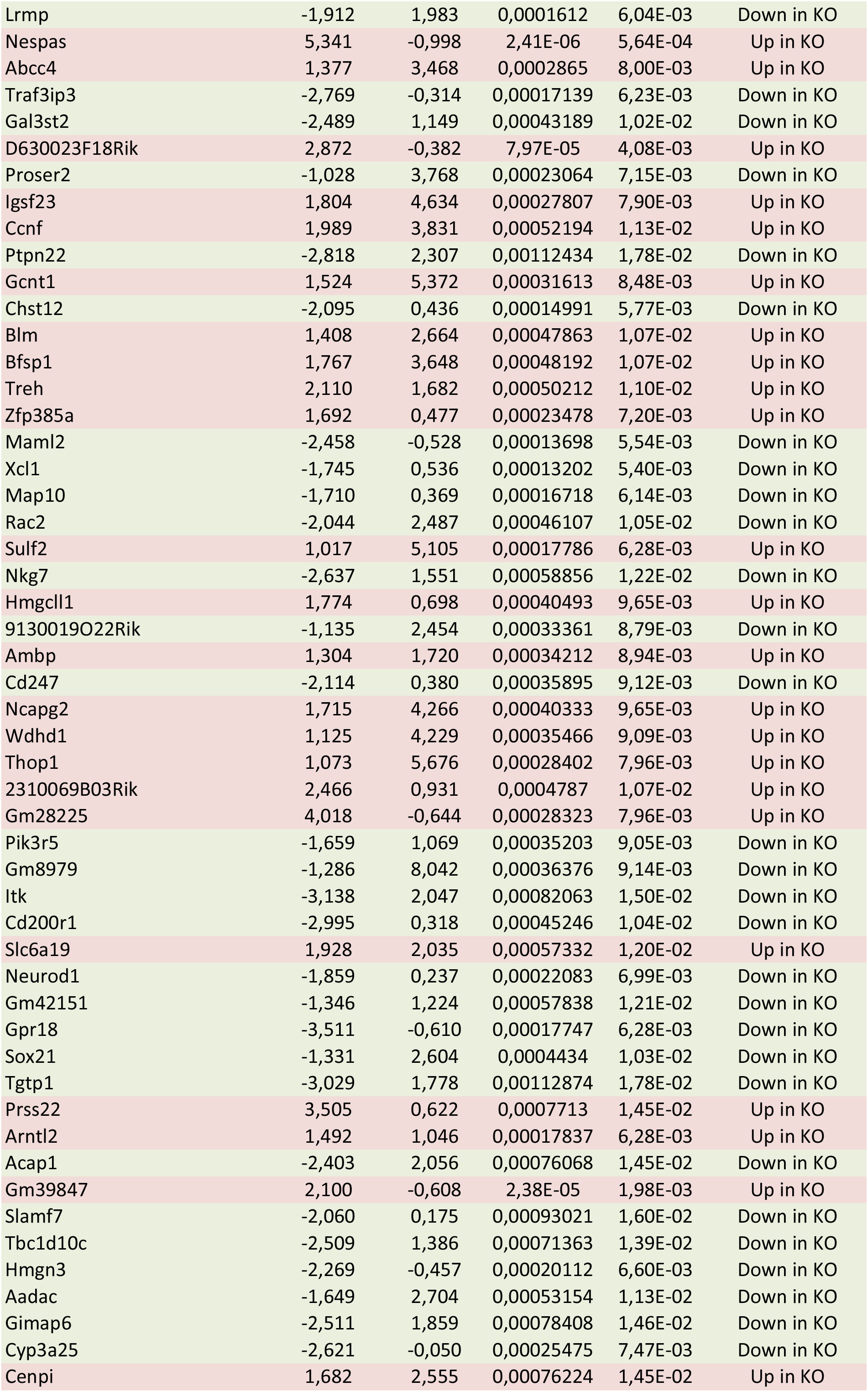
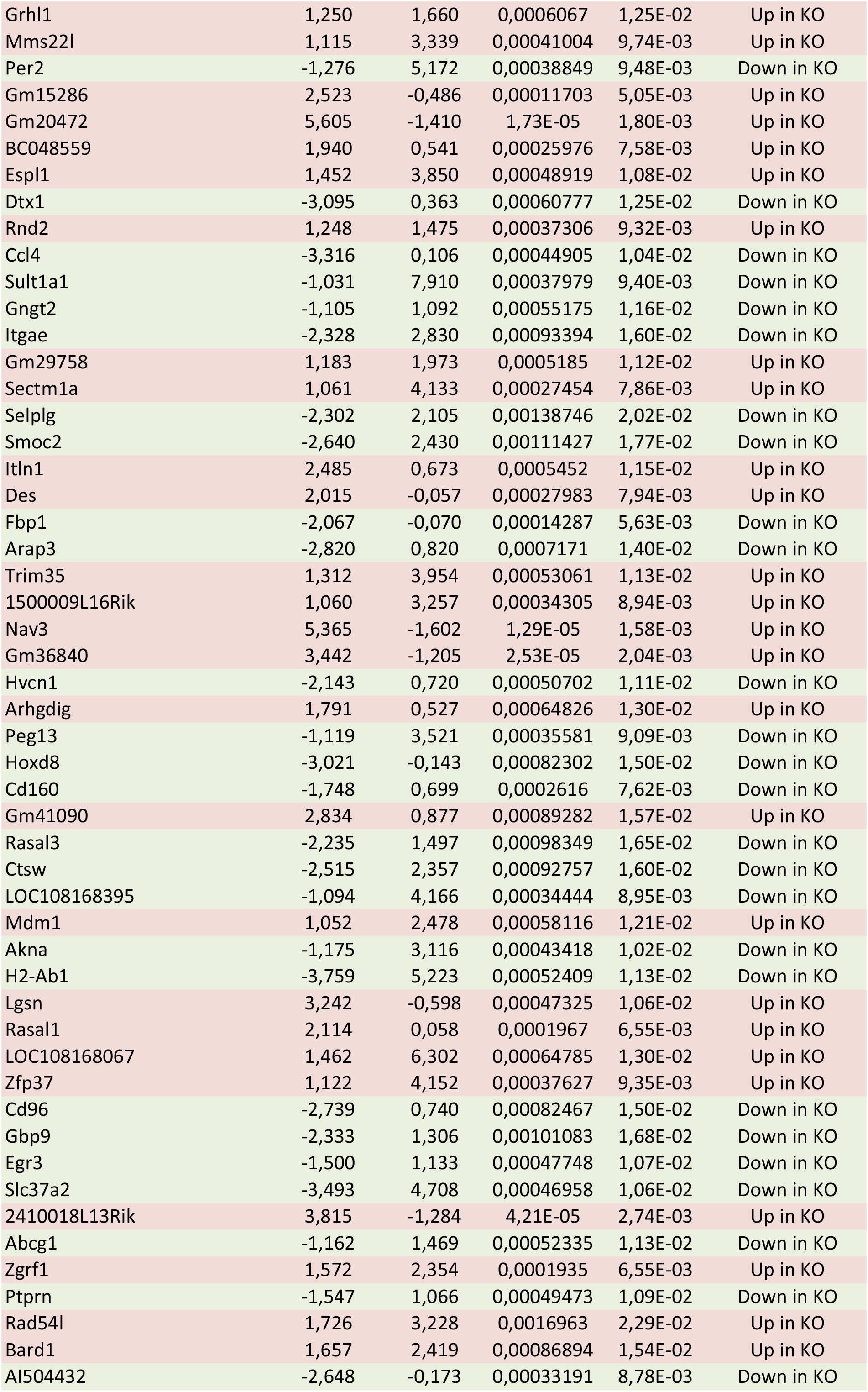
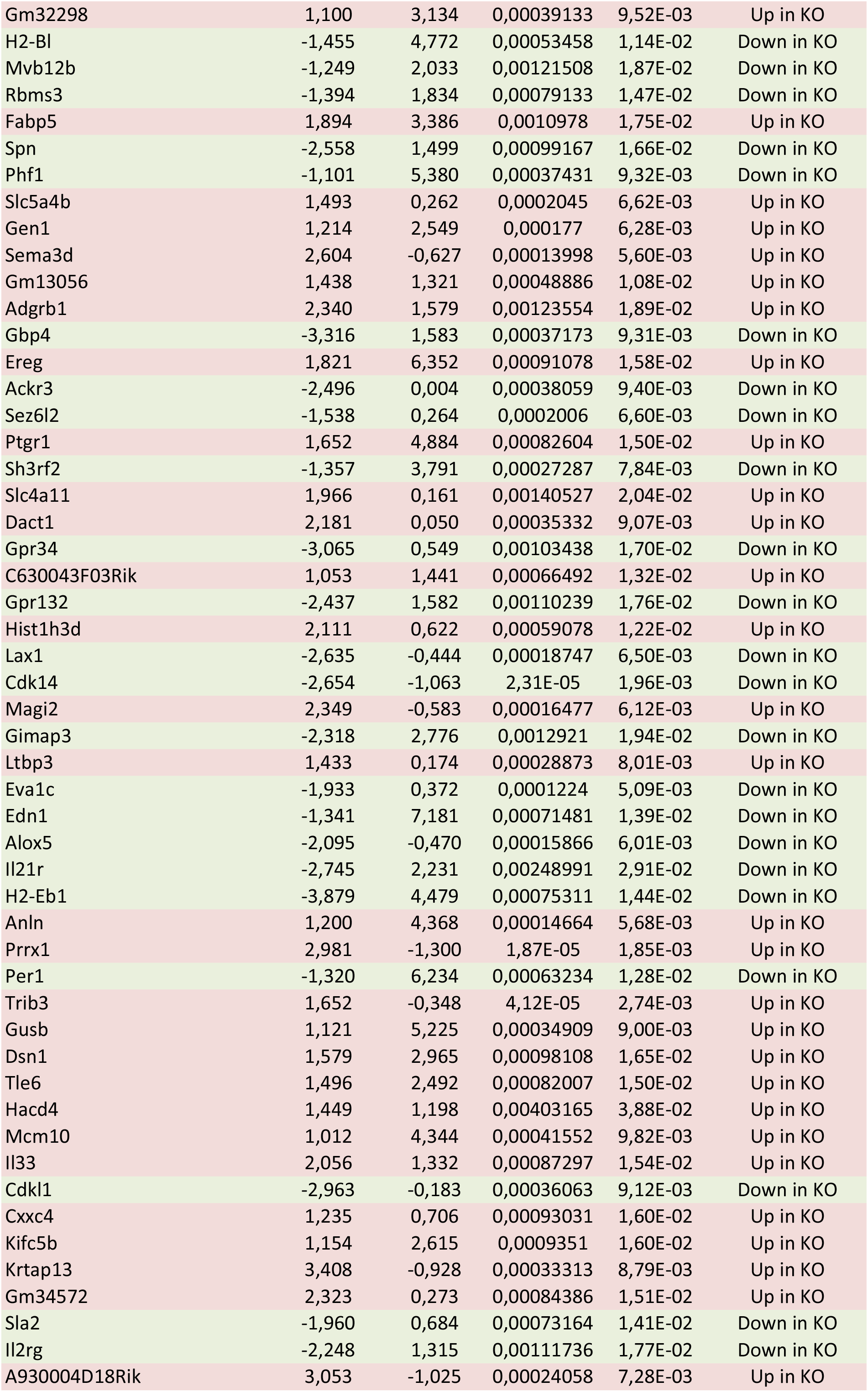
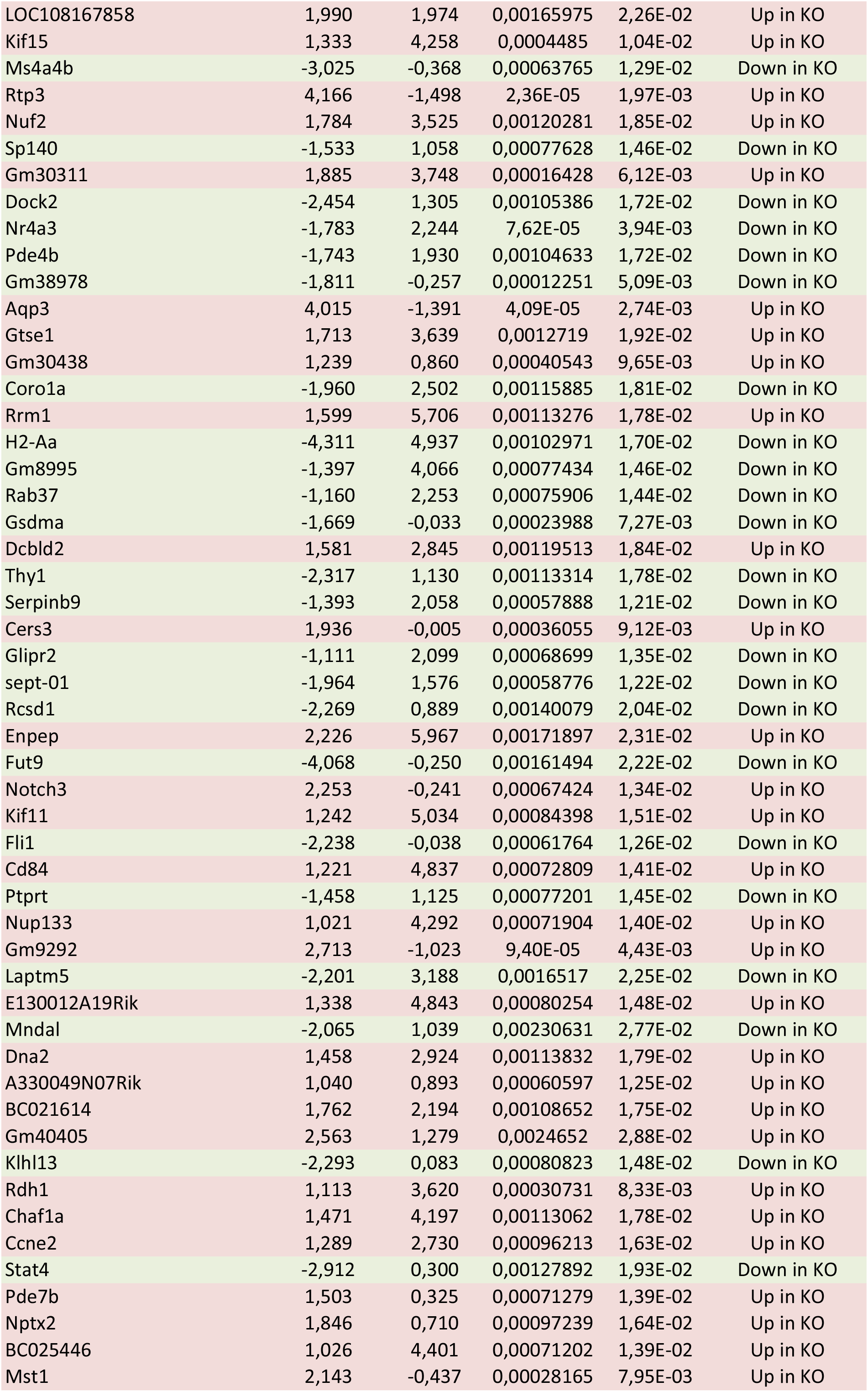
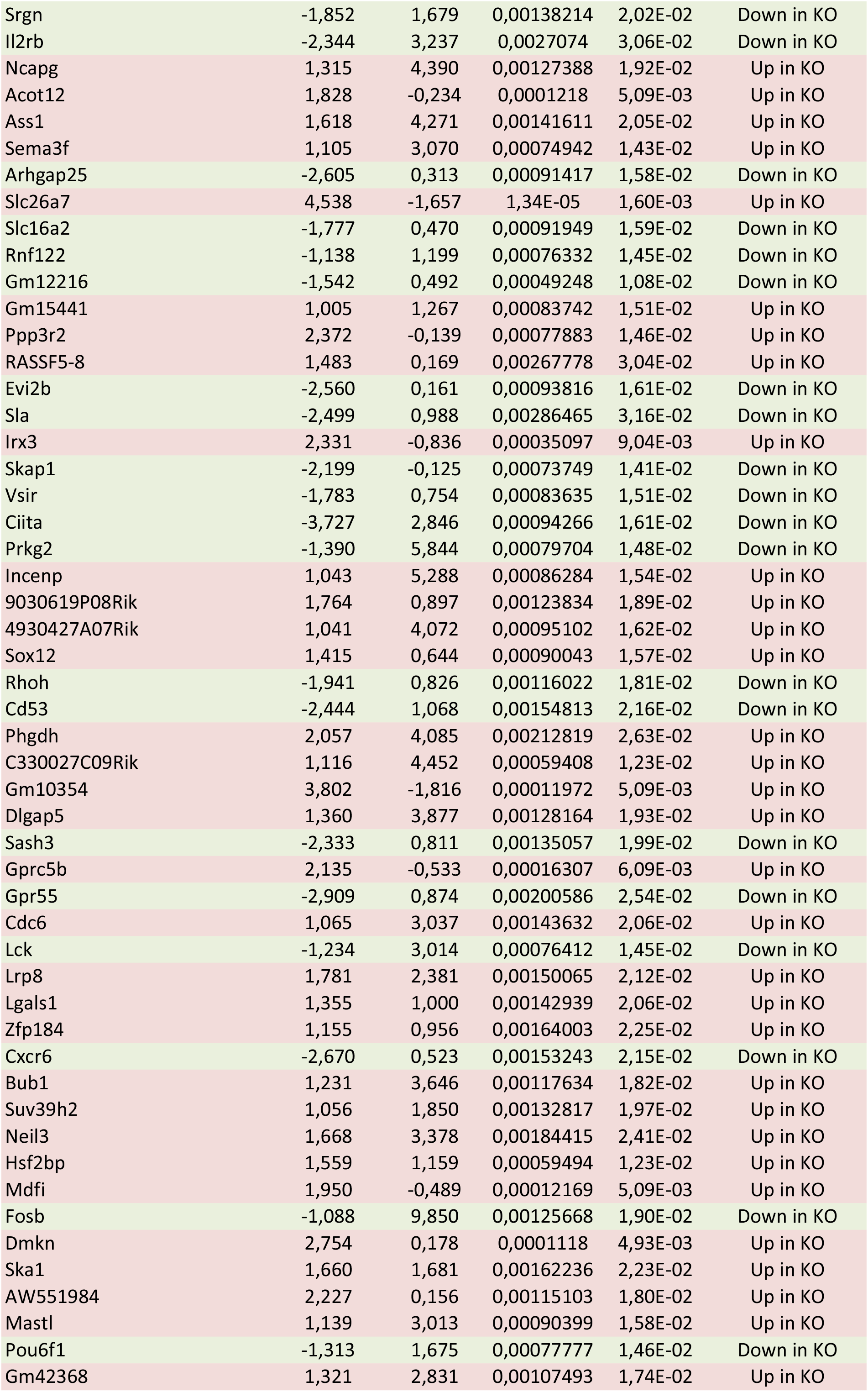
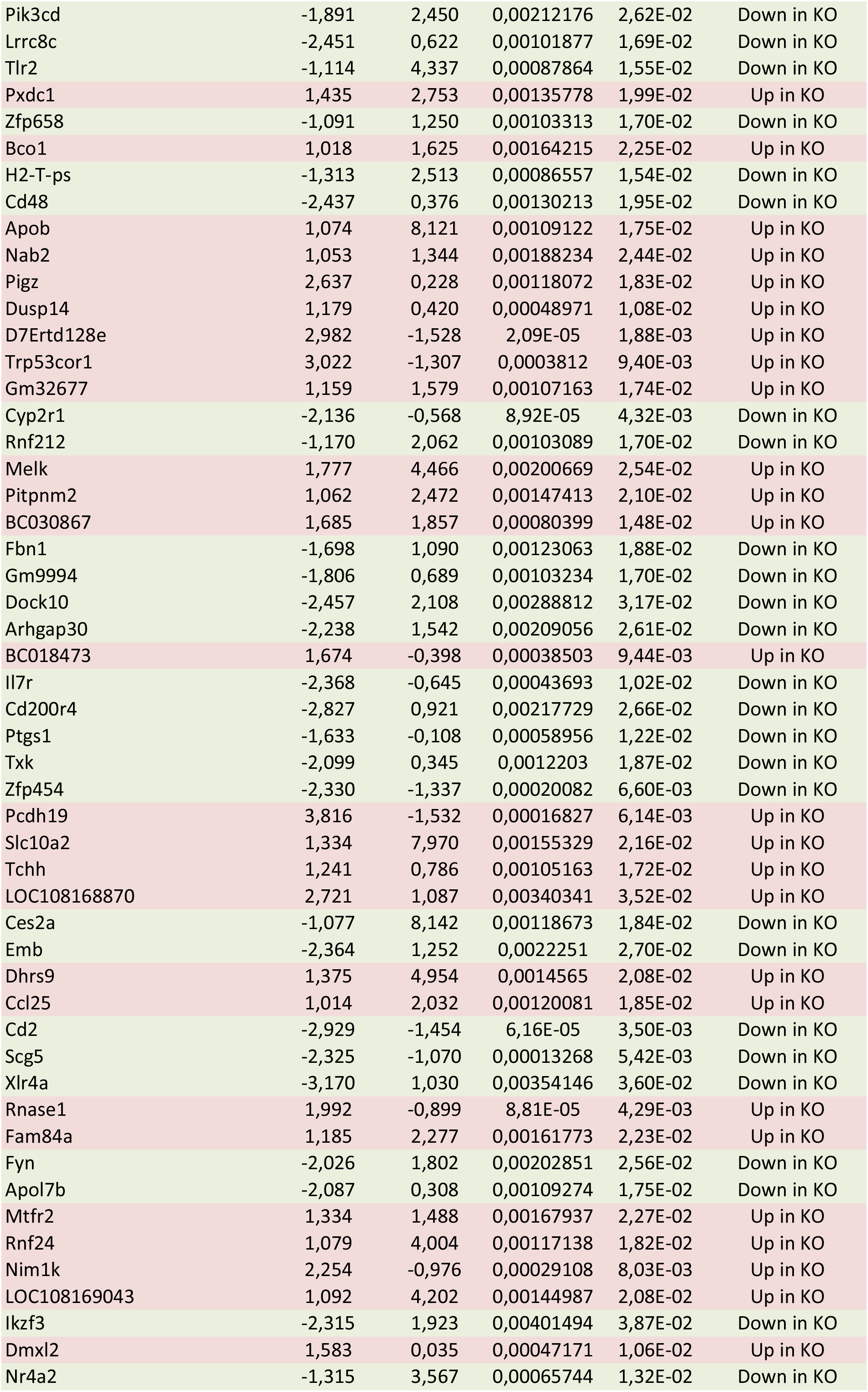
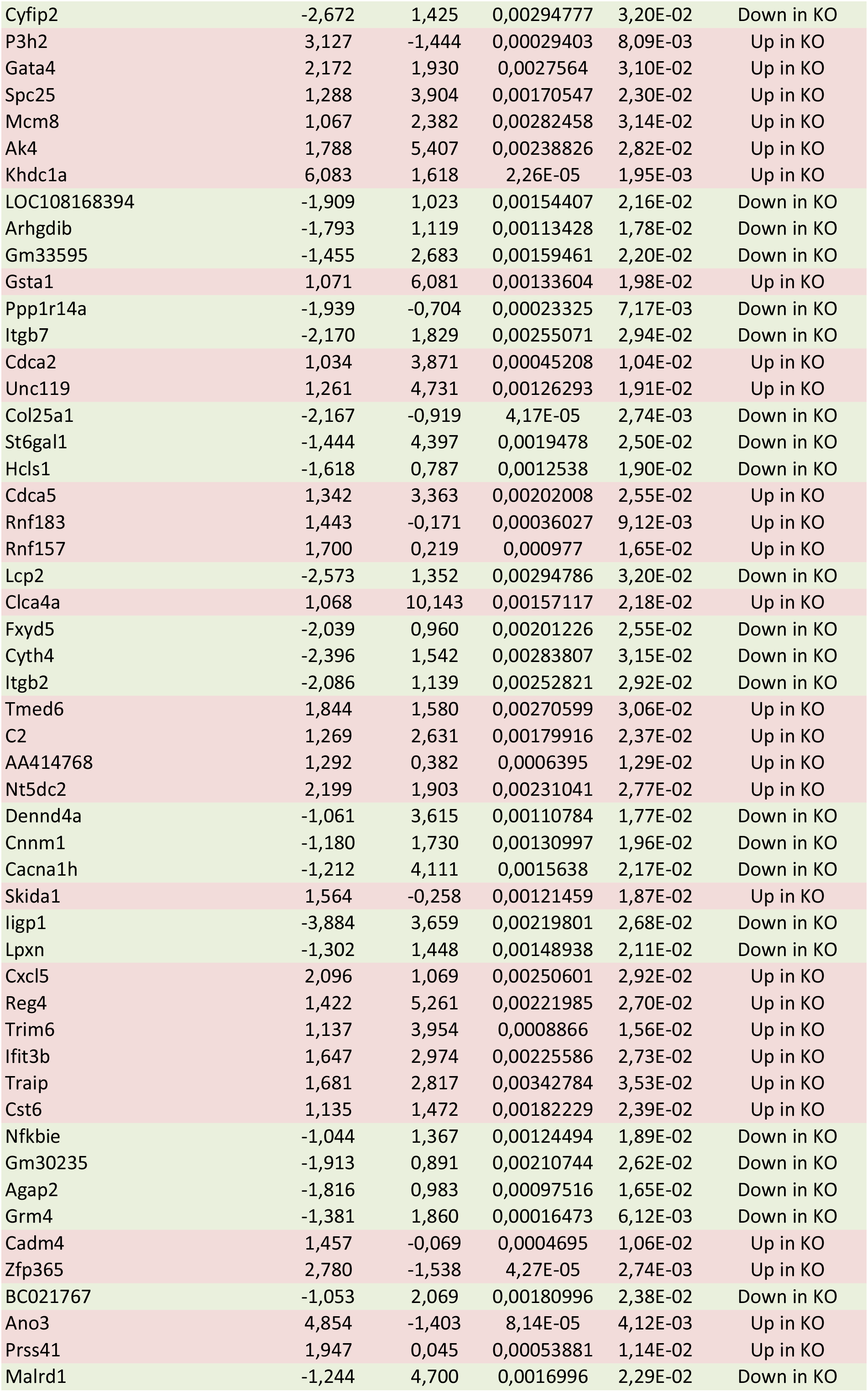
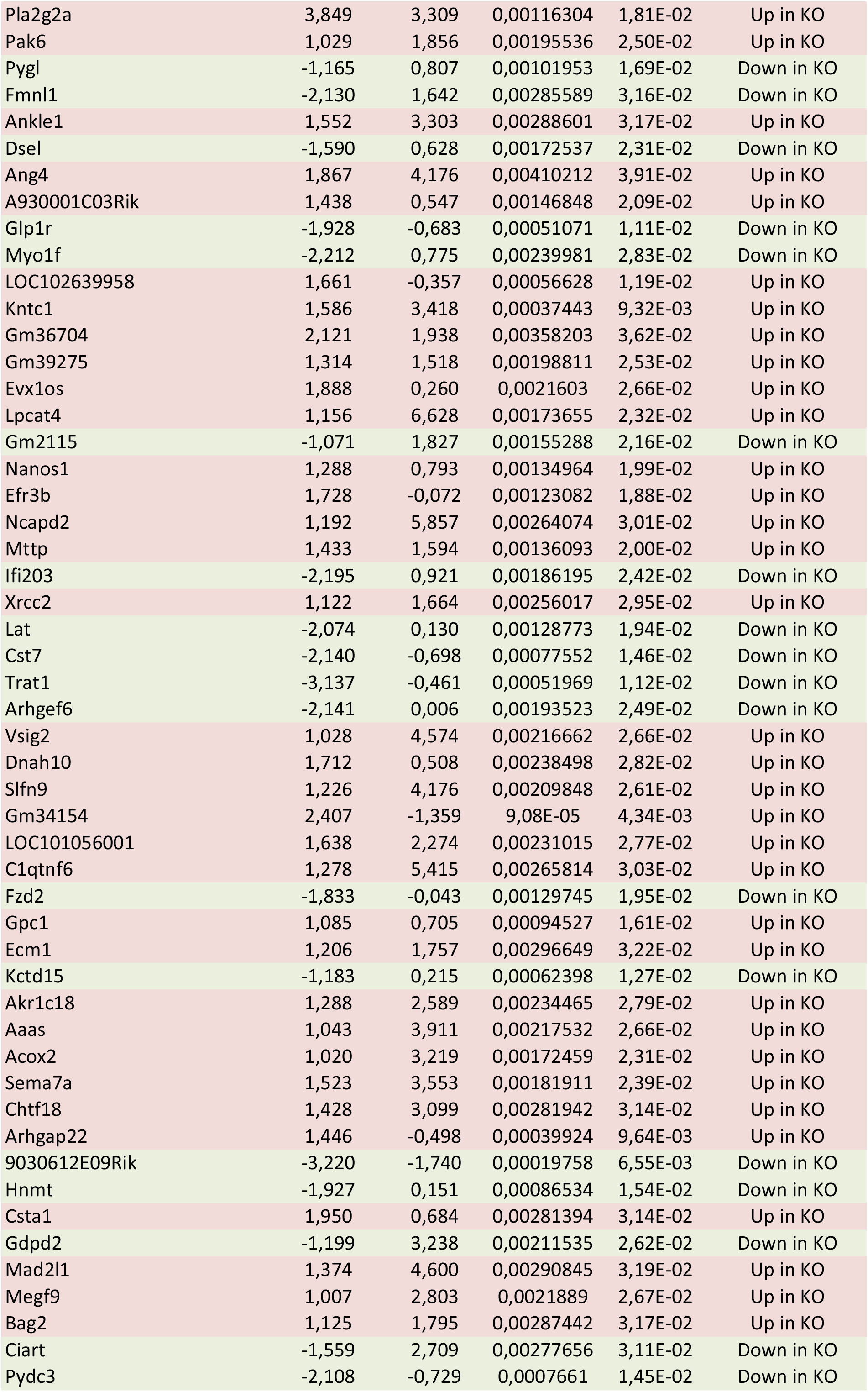
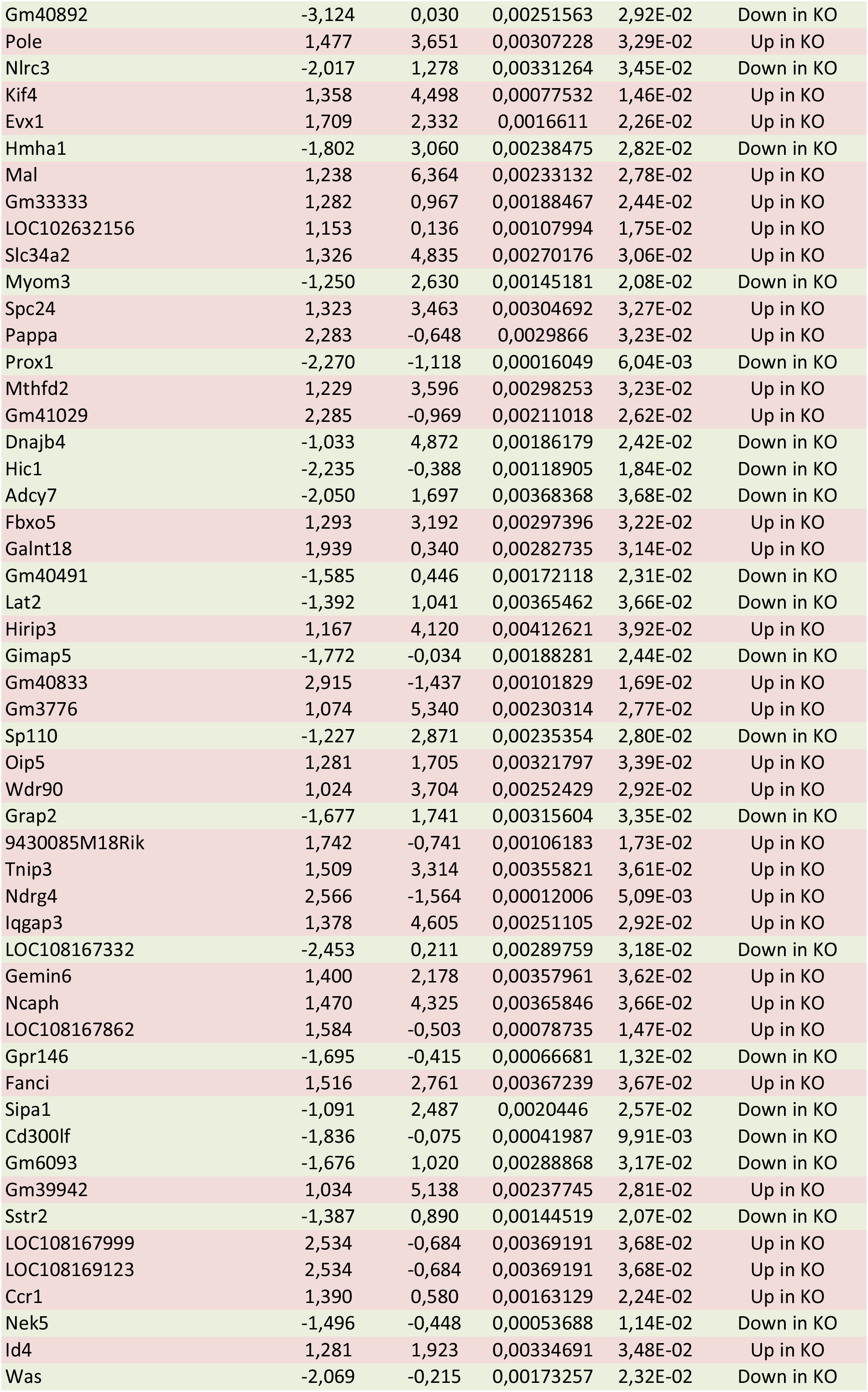
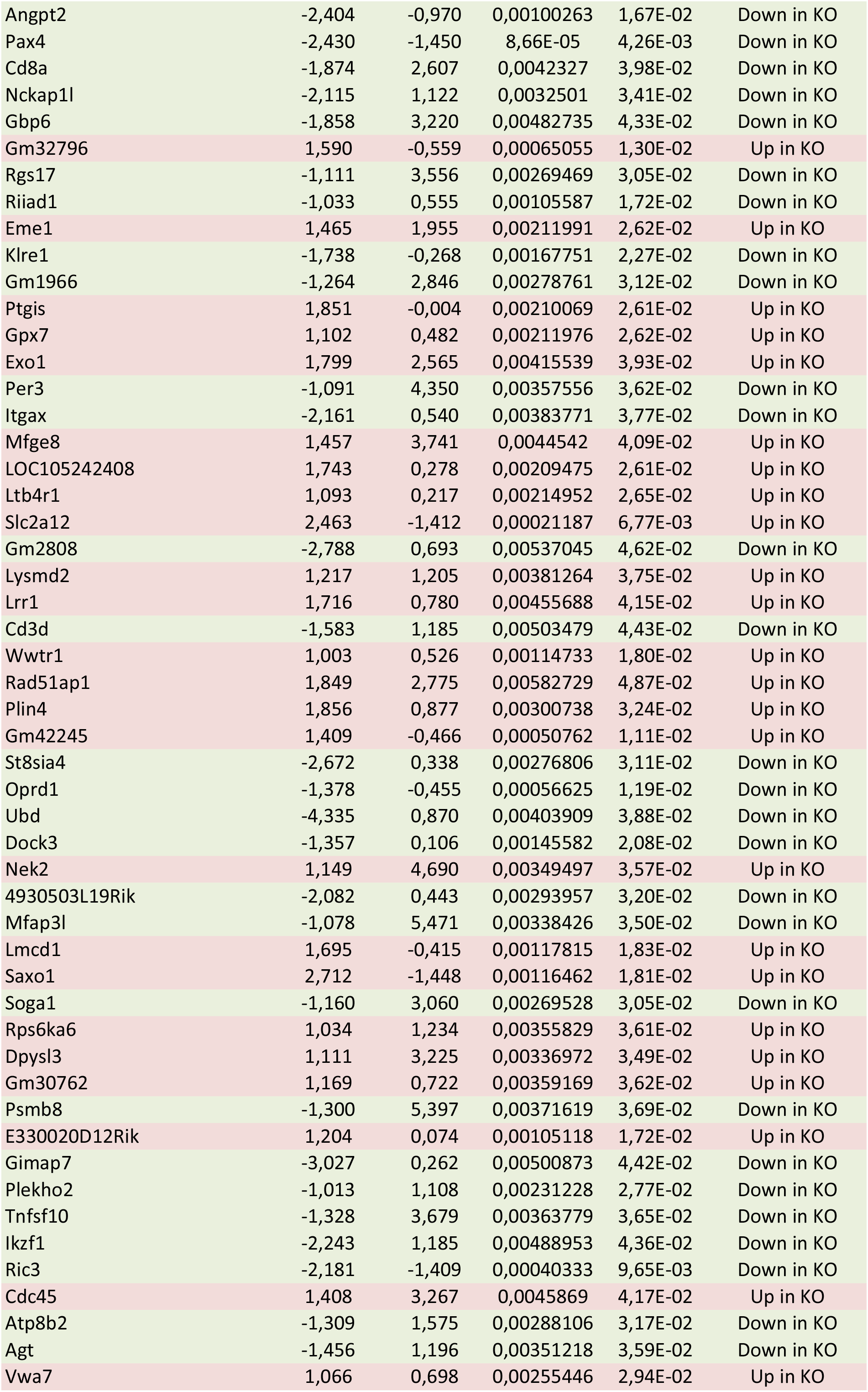
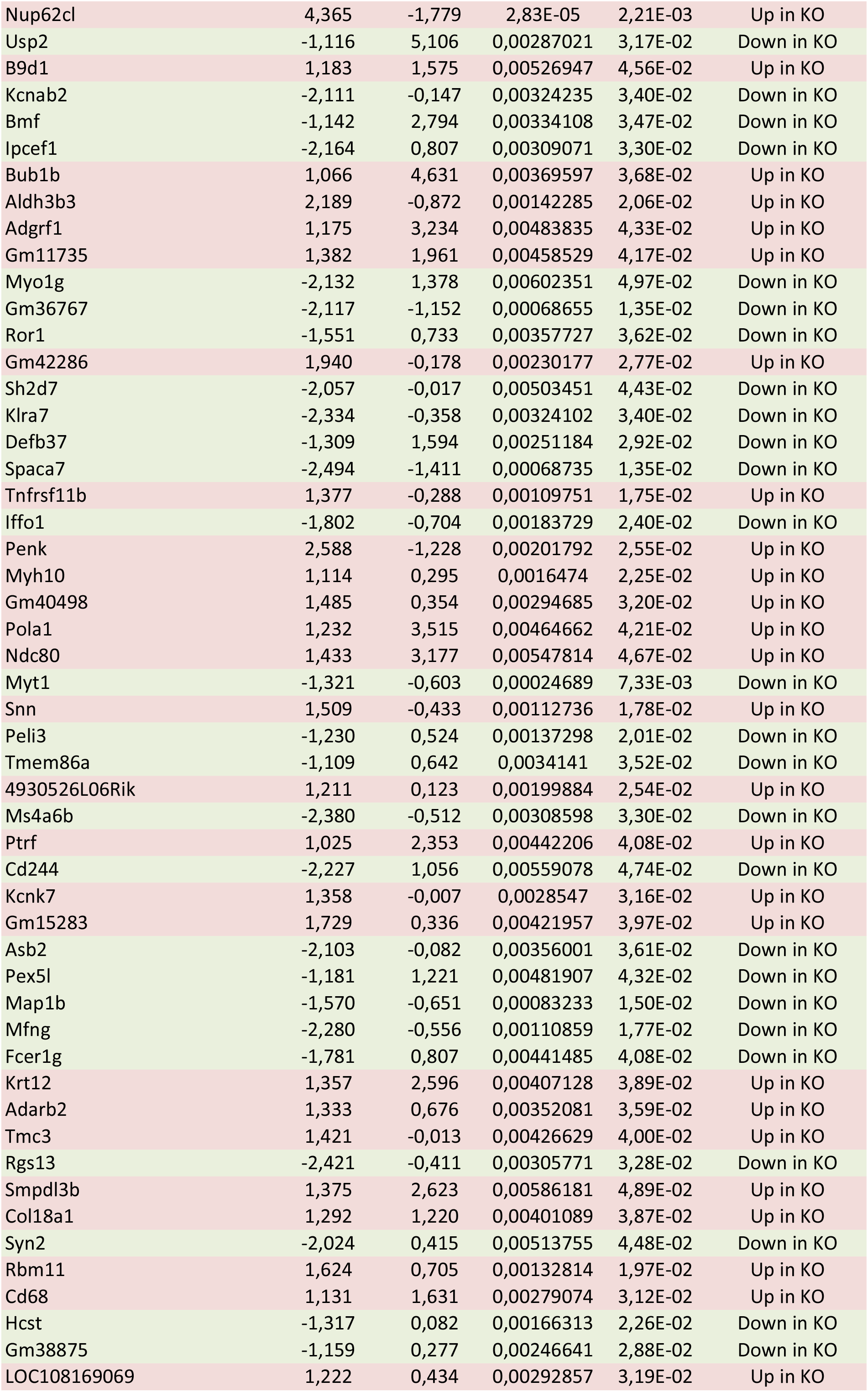
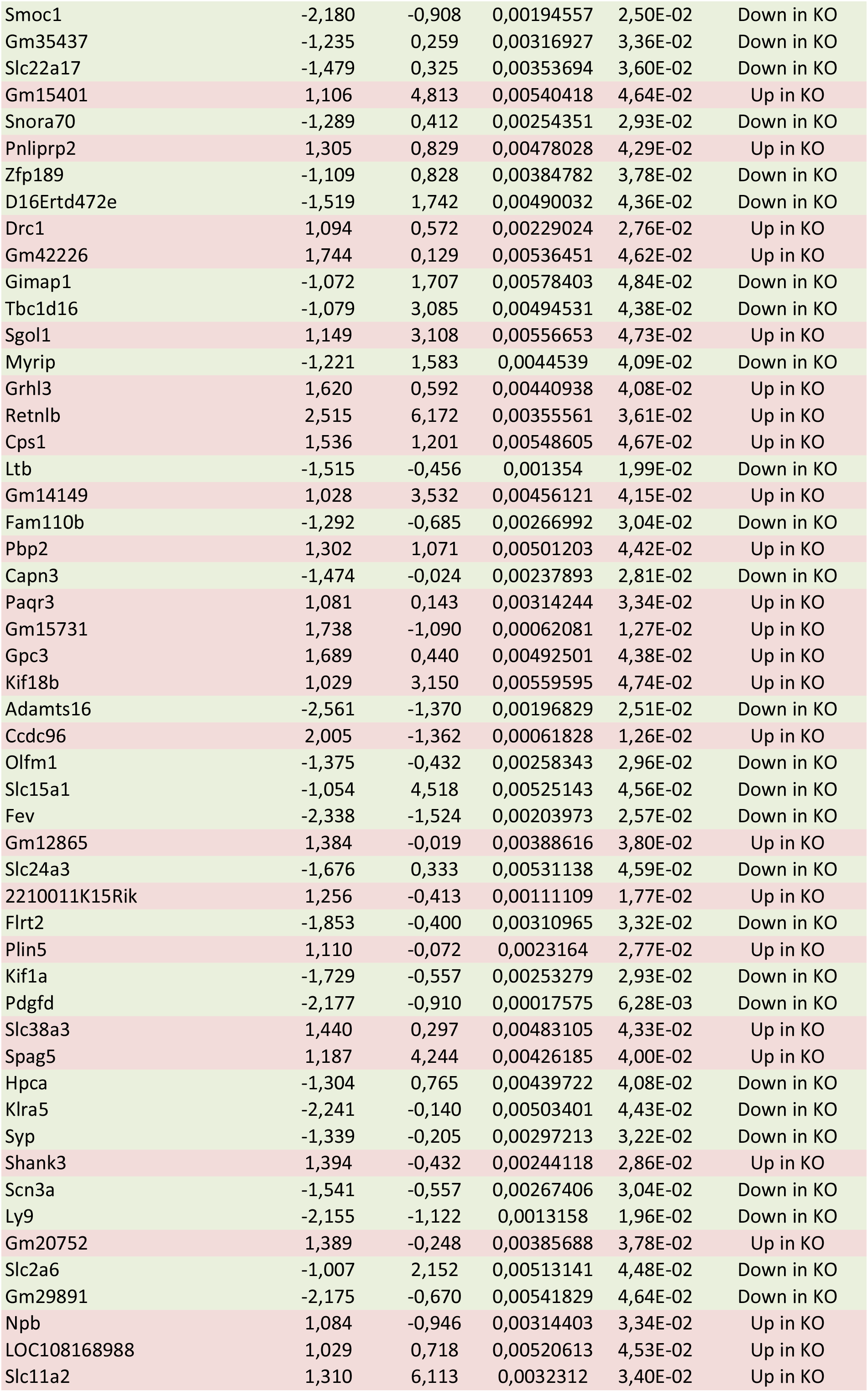
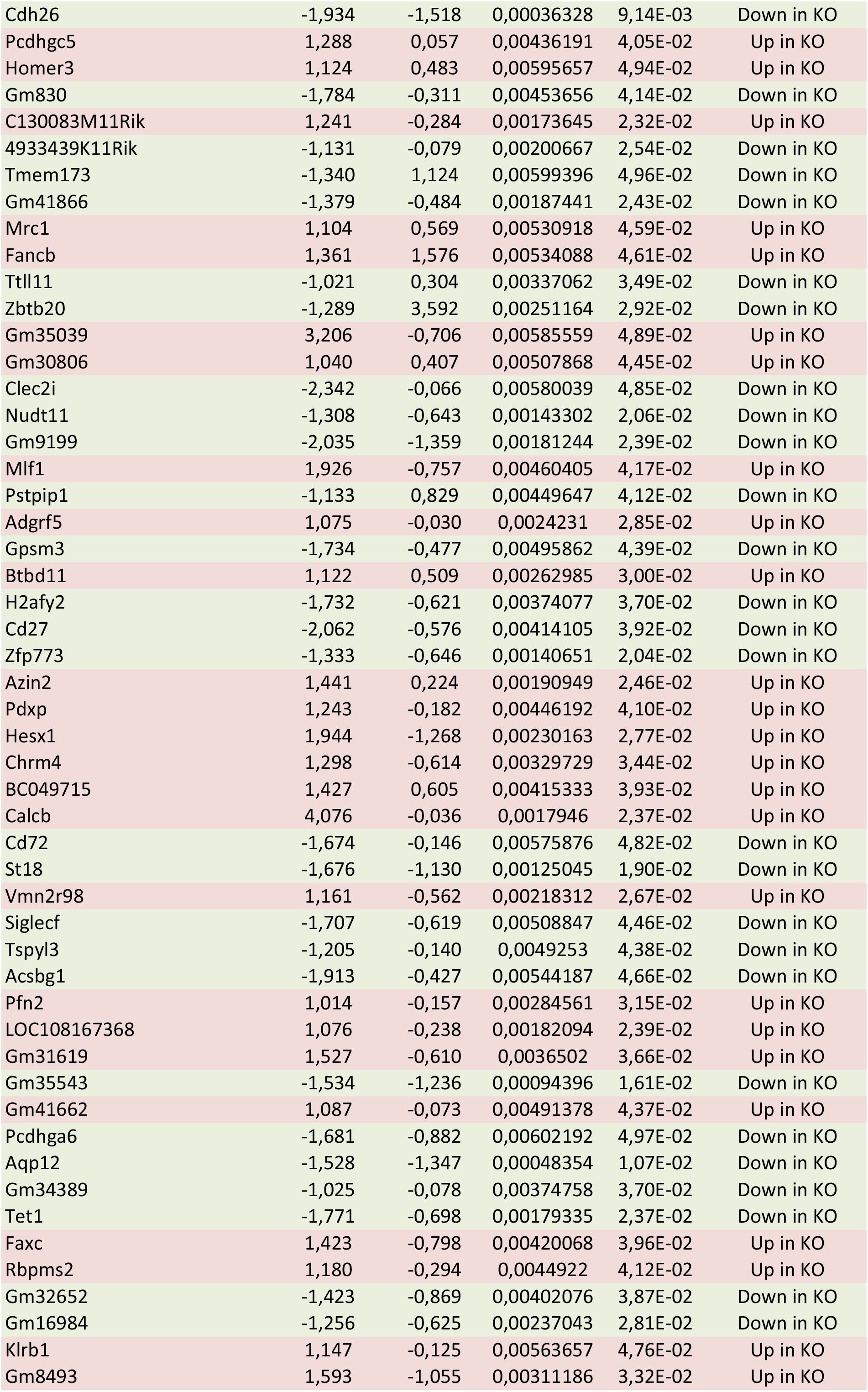
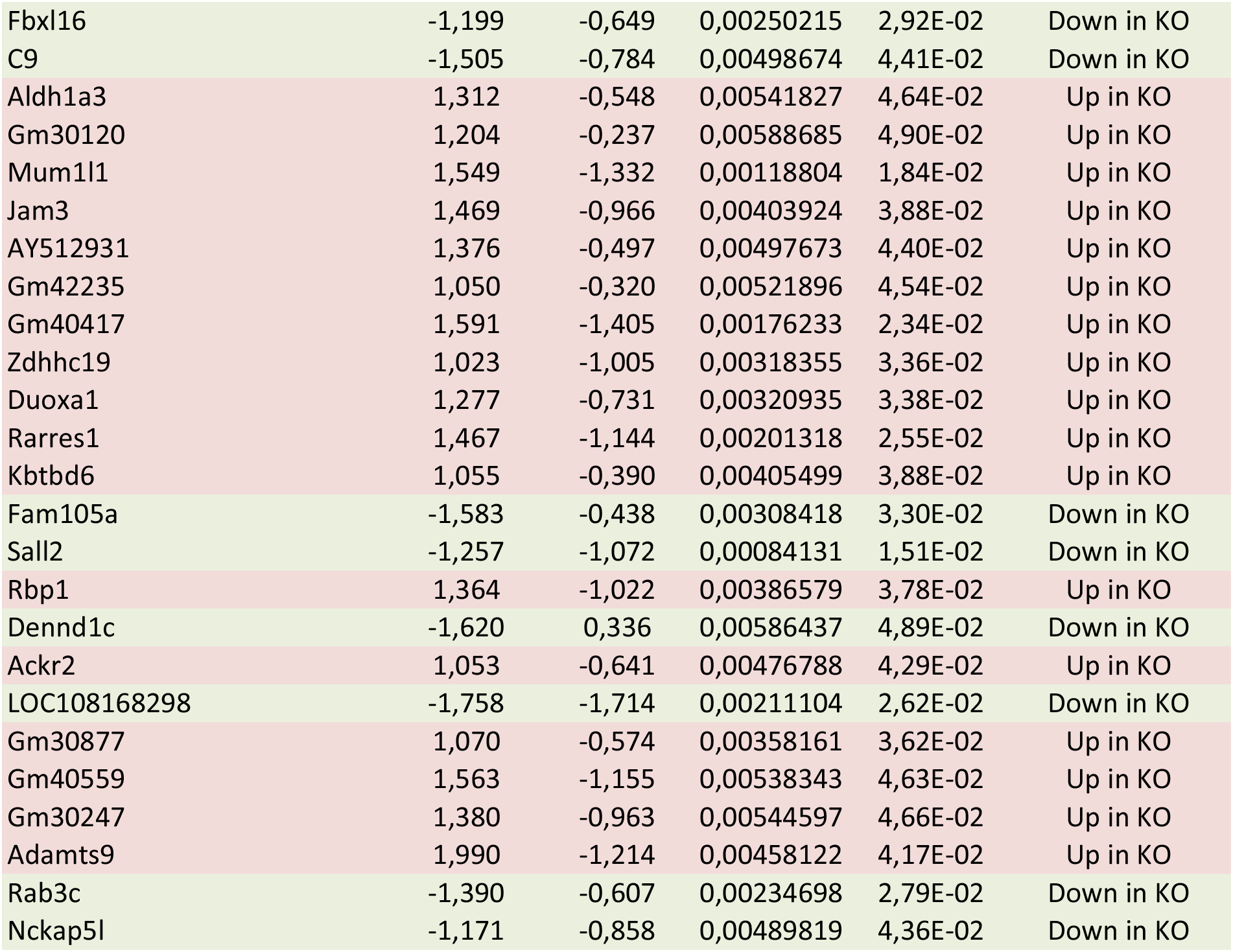
List of the 1109 genes differentially expressed between *VilCreCcna2fl/fl* mice and *Ccna2fl/fl* mice samples (fold change > 2, adjusted p-­-value |FDR| <0.05). FC: Fold change, CPM: Counts-­-per-­-million, FDR: False Discovery Rate

### Cyclin A2 loss in colonic epithelial cells leads to increased proliferation of colonic epithelial cells and onset of dysplasia

To validate the RNAseq results described above, we first tested the proliferation status of cyclin A2-deficient colonic epithelial cells by measuring BrdU-uptake. For this, mice received an intraperitoneal injection of the nucleoside analog BrdU and were sacrificed two hours later to isolate colons. The subsequent IHC analysis was performed by dissecting the different regions of the colon as described above (Supplemental Fig 1A) and by separating the bottom, middle and top parts of the crypts. As shown in Figure 2A, BrdU-positive cells were mainly located in the bottom of the crypts of control mice. In contrast, BrdU incorporation was most prominent in the middle part of the cyclin A2-deficient crypts with an average of more than 4 cells per crypt (n=60, p< 0.0001). Moreover, BrdU-positive cells were detectable in the upper crypt part of *VilCreCcna2fl/fl* but not in control mice. These specific profiles of BrdU incorporation of cyclin A2-deficient colonic epithelial cells were observed equally in distal, transverse as well as the proximal region. To confirm the increased proliferation rate of cyclin A2-deficient CEC, we analyzed Ki-67 expression, which is expressed during all active phases of the cell cycle (i.e., G1, S, G2, and M), but is absent in resting cells (28). Ki-67 expression in the distal region of colons was detectable at the bottom of the crypts in colons from control mice but localized to the middle and upper parts of the colonic crypts from *Ccna2* knockout animals. A similar pattern was observed in the proximal region of colons (Fig 2B, Supplemental Fig 3A and 3B) of constitutive as well as induced knockout mice (Supplemental Fig 3C) therefore confirming the observation made in the BrdU-uptake experiment. Thus, the elevated expression levels of Ki-67 detected by immunostaining in cyclin A2-deficient colons concurs with the upregulation of *Mki67* transcripts detected in the RNAseq analysis.

**Figure 3:**
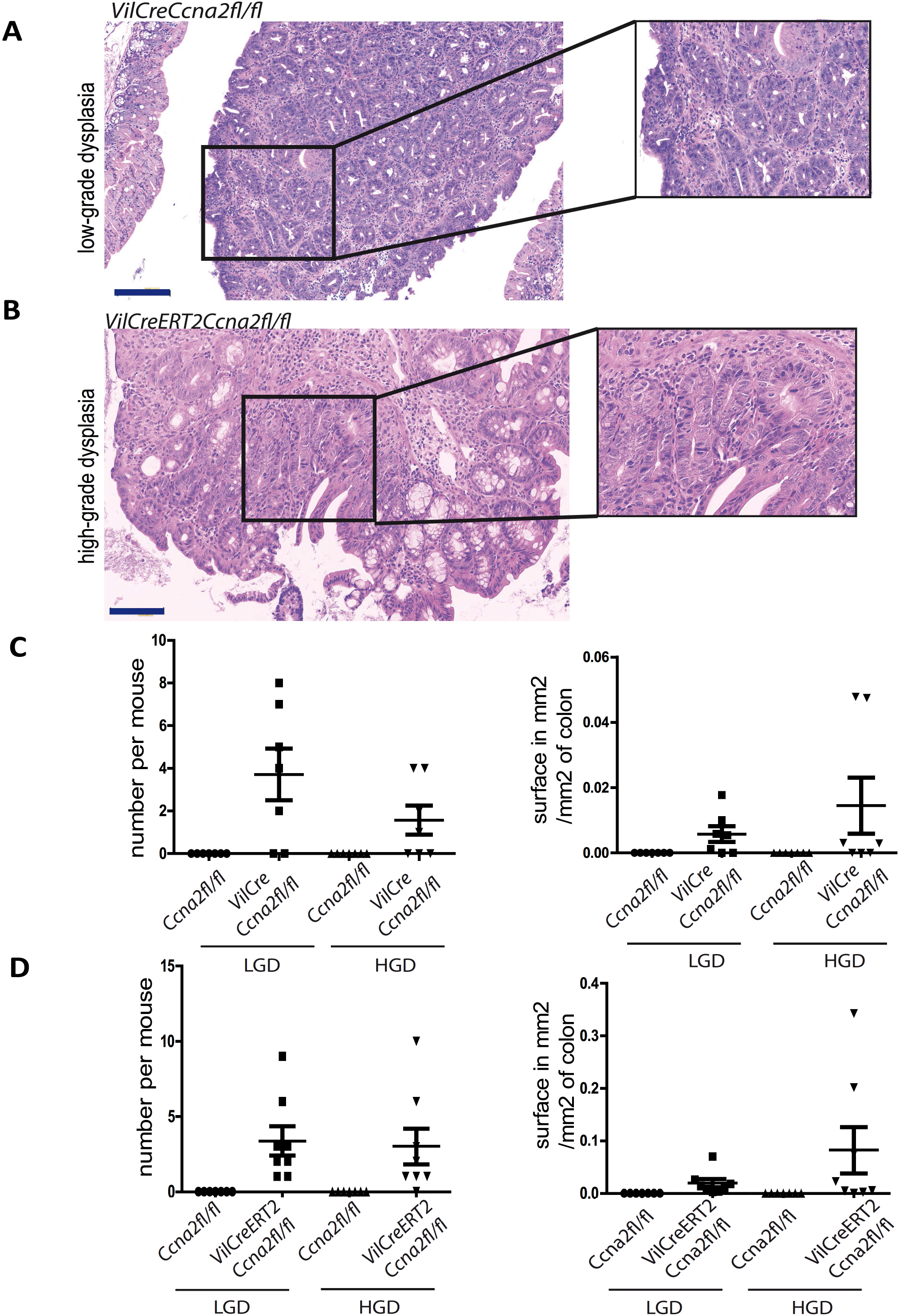
Dysplasia formation in cyclin A2-deficient mice. **(A, B).** Representative images of hematoxylin and eosin staining displaying low- and high-grade dysplasia (LGD and HGD, respectively) in the colons of constitutive (*VilCreCcna2fl/fl)* (**A**), and tamoxifen-induced (*VilCreERT2Ccna2fl/fl)* knockout mice (**B**). **(C, D).** Quantification of the number of low- and high-grade dysplasia and the area of dysplasia (in mm^2^ per mm^2^ of colon) in constitutive (n=7; *VilCreCcna2fl/fl*) and induced knockout mice 8 days after tamoxifen treatment (n=8; *VilCreERT2Ccna2fl/fl*) mice in comparison to control animals (n=7; *Ccna2fl/fl)*. Scale bars: 100μm. Blow-up in **A** and **C** is 2,5x.

Strikingly, colons deficient for cyclin A2, either constitutive or induced, revealed low- and high-grade dysplasia in around 80% of the mice analyzed that was mostly found in the proximal region (Fig 3A and B). Constitutive knockout mice showed an average occurrence of 4 low-grade dysplasia and nearly 2 high-grade dysplasia per colon, while induced knockout mice displayed a mean of about 3 low-grade and high-grade dysplasia per colon, whereas none were detectable in the control mice (Fig 3C and D). Notably, dysplasia has been already detectable eight days after tamoxifen injection in *VilCreERT2Ccna2fl/fl* mice.

### Cyclin A2 depletion in colonic epithelial cells leads to abnormal mitoses and DNA damage

We further explored cell cycle features and noticed an increased incidence of mitoses within the crypts when analyzing colons of constitutive and induced knockout mice (at day 8). Colons with constitutive and induced-*Ccna2* deletion showed an average of 13 and 24 mitotic figures per mm^2^, respectively, whereas in control animals in average only 2 per mm^2^ were found (n=5, p<0.01 for the constitutive as well as induced knockout mouse strain, Fig 4A and B). Besides, CEC deficient for cyclin A2 harbored nuclear pleomorphism characterized by increased nuclear size compared to controls (Figure 4A and Supplemental Fig 4). Finally, abnormal mitosis, detected by combined staining for α-tubulin to visualize mitotic spindles, γ-tubulin to label centrosomes and DAPI to identify chromosomes, were prominent in induced cyclin A2-deficient but not control colons (Supplemental Fig 5 A and B).

**Figure 4:**
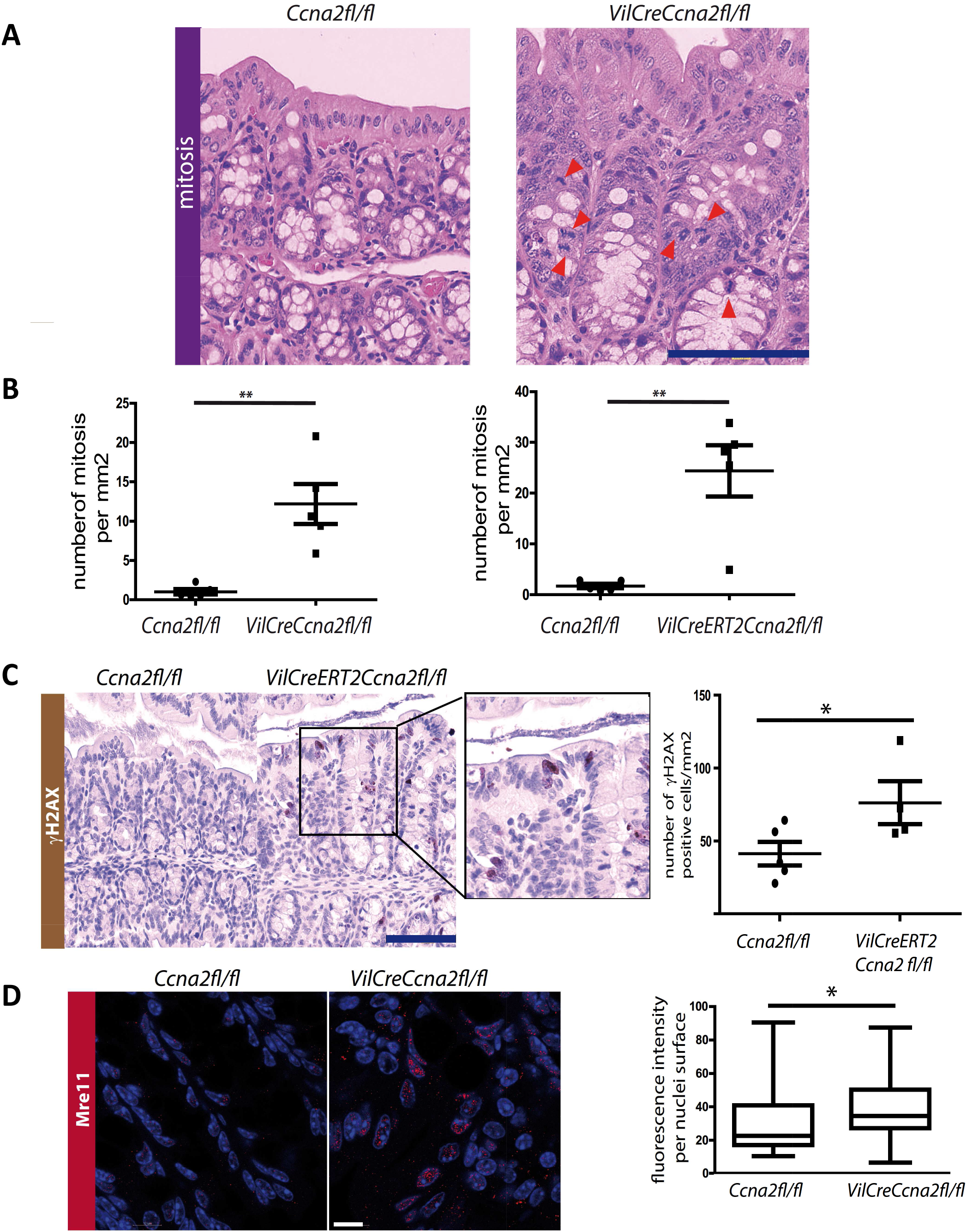
Elevated mitoses, DNA damage and activation of β-catenin following cyclin A2 deletion in the colonic epithelium. **(A).** Representative HE staining showing an increased number of mitoses (red arrowheads) in the proximal region of the colon from cyclin A2-deficient mice compared to a colon of a control animal. Scale bar: 100 μm. **(B).** Quantification of mitotic events in constitutive (left panel; n=5) and induced knockout mice (day 8 following inactivation; right panel; n=5) and control animals (n=5); expressed as mean ±SEM, **\*\*** p<0.01; two-tailed unpaired t-test. **(C).** γH2AX expression analyzed by immunohistochemistry on colon sections of control (*Ccna2fl/fl*; n=5) and tamoxifen-induced knockout (*VilCreERT2Ccna2fl/fl*; n=4) mice at day 8 following cyclin A2 inactivation. Representative γH2AX immunostaining and quantification of the number of γH2AX positive cells per mm^2^ of epithelium from the proximal part of colons are shown (expressed as mean ±SEM**, \***p<0.05; two-tailed unpaired t-test). Scale bar: 100μm. **(D).** Confocal analysis of nuclei foci formation of Mre11 in crypts from the proximal part of colons from control (*Ccna2fl/fl)* and *VilCreCcna2fl/fl* mice. Representative images are shown; Scale bar: 10 μm; (staining of nuclei with DAPI in blue and Mre11 in red) as well as quantification of the Mre11 fluorescence intensity per nuclei surface (right panel; n=83 different nuclear areas for control and n=65 for cyclin A2-deficient mice; mean ±SEM are provided, **\***p<0.05; two-tailed unpaired t-test).

**Figure 5:**
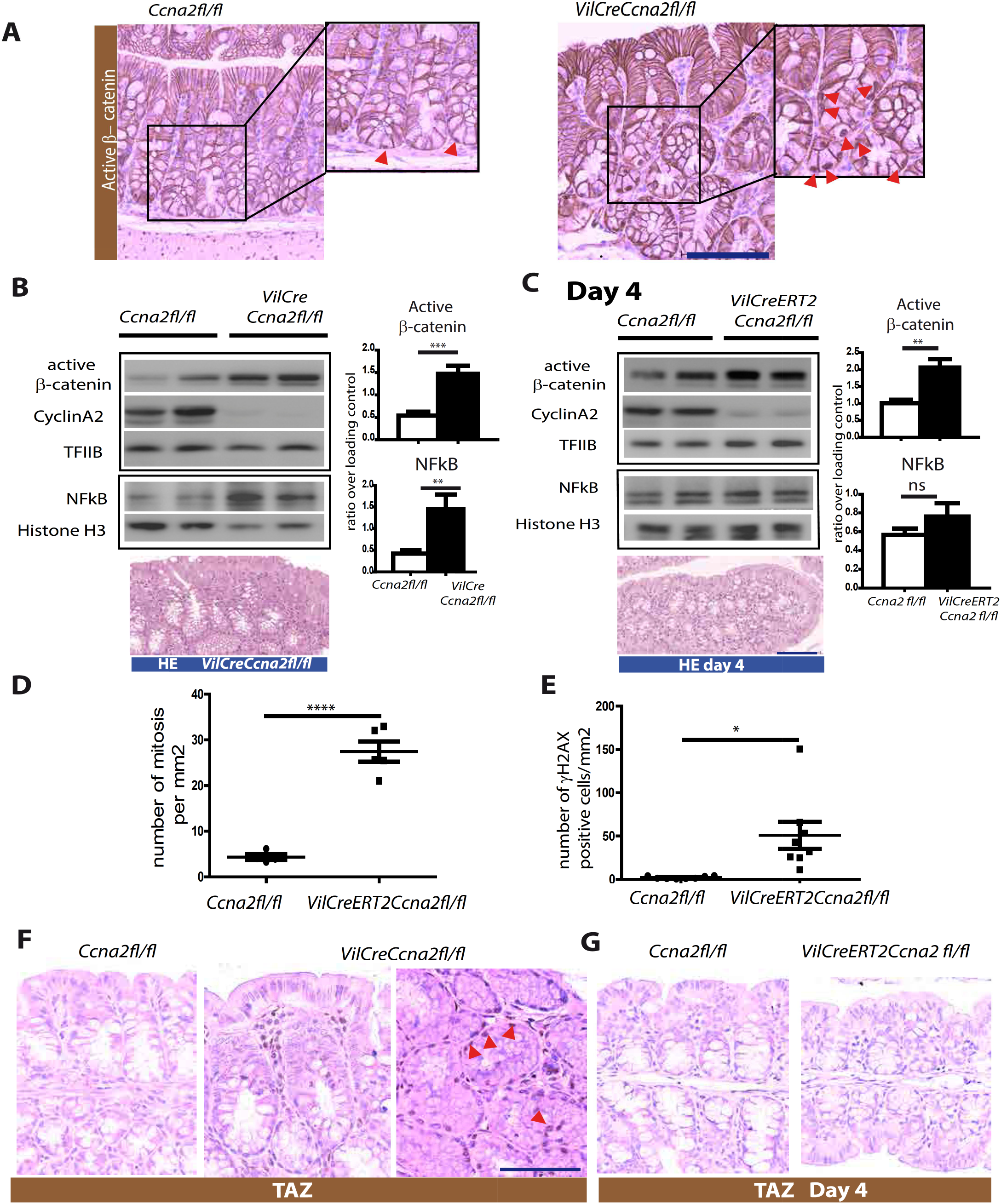
Cyclin A2 deficiency triggers regenerative pathways. **(A).** Representative immunostaining for active β-catenin in control (*Ccna2fl/fl)* and constitutive knockout (*VilCreCcna2fl/fl*) mice. Arrowheads indicate nuclear localization of active β-catenin. Scale bar: 100μm. **(B).** Western blot analysis of active β-catenin and NFκB in CEC extracts derived from constitutive cyclin A2-deficient mice and their control littermates. The right panel shows quantification of active β-catenin and NFκB after normalization with loading controls (TFIIB or Histone H3, n=9). **(C).** Western blot analysis of active β-catenin, and NFκB with their respective quantifications on nuclear extracts derived from CEC of control and tamoxifen-induced knockout mice (*VilCreERT2Ccna2fl/fl)* following deletion of cyclin A2 at day 4 (n=6). Lower panel: Representative HE staining of colons from control and induced knockout mice at day 4 following *Ccna2* inactivation. (**D-E**). Colons of cyclin A2-deficient mice display DNA damage. Quantification of mitoses (**D**) and gH2AX staining (**E**) in colons of control and VilCreERT2Ccna2fl/fl mice at day 4 following tamoxifen-induced Ccna2 depletion (n= 8 mice). (**F, G**). IHC analysis of TAZ expression (indicated by arrow) in constitutive mutant colons (**F**) and in induced knockout mice at day 4 following Ccna2 deletion (**G**), representative images are shown. Scale bar: 100 μm in B, C and D; mean ±SEM are provided, * p<0,05, ** p<0.01 and *** p<0,001; two-tailed unpaired t-test.

Defective mitoses can lead to DNA damage (29), and we thus analyzed colons of tamoxifen-treated *VilCreERT2Ccna2fl/fl* mice by IHC using a marker for double-strand breaks (DSB), *i.e*. an anti-γH2Ax antibody, and found an average of more than 70 γH2AX positive cells/mm^2^ in cyclin A2-deficient CEC compared to 40 in control ones (Fig 4C, n=4, p<0.05). Mre11 is part of the NMR complex involved in the resolution of double-strand DNA (30). We therefore analyzed Mre11 foci formation by confocal microscopy analysis (31) and found that Mre11 foci formation was significantly increased in *VilCreCcna2fl/fl* nuclei (Fig 4D).

Altogether, the alterations found at the protein level correlate with the RNAseq results underpinning the presence of cell cycle perturbations, increased DNA damage and DNA double-strand break repair pathway activation in cyclin A2-deficient CEC.

### Activation of regenerative pathways in cyclin A2-deficient colons

Tissue injury induces inflammation and in turn a regenerative response (32). Several pathways have been described that can promote tissue repair in the intestine including YAP/TAZ, NFκB and WNT/β-catenin signaling (32, 33). We have previously shown that cyclin A2 inactivation in epithelial cells *in vitro* leads to increased β-catenin stability and activity (18, 19). For this reason we first tested β-catenin expression in the colons of constitutive knockout mice by IHC analysis revealing increased nuclear localization of β-catenin in CEC of cyclin A2 deficient colons by IHC (Fig 5A).

In addition, Western blot analysis of nuclear extracts derived from the constitutive knockout mice displayed elevated levels of both active β-catenin as well as the NFκB p65 subunit (Fig 5B). We also detected TAZ expression in crypts as well as surrounding stroma in colons of *VilCreCcna2fl/fl* mice (Fig 5 F).

We next examined the activation sequence of the different pathways modulated in cyclin A2-deficient CEC at the protein level. For this, we took advantage of the inducible Cre model (*VilCreERT2Ccna2fl/fl* mice) and examined the altered pathways four days upon *Ccna2* depletion when colons still not display alterations such as changes of crypt architecture or immune cell infiltration (lower panel in Fig 5B and C). Expression levels of active β-catenin and NFκB were analyzed by Western blot (Fig 5C) and those of γH2AX and TAZ by IHC (Fig 5E-G). Besides, quantification of mitoses was performed on HE stained colons from induced knockout mice (Fig 5D). Elevated levels of active β-catenin, mitoses as well as γH2AX were already detectable at day 4 in cyclin A2-deficient CEC (Fig 5C-E), whereas NFκB and TAZ levels were unaltered at that time point between knockout and control animals (Fig 5C and 5G).

### Cyclin A2 deficiency in CEC promotes colon carcinogenesis in mice

To test whether cyclin A2-deficiency modulates colon carcinogenesis, we subjected *VilCreCcna2fl/fl* and *Ccna2fl/fl* mice to chemically induced colon carcinogenesis, an established mouse model resembling the human pathology (39). This model depends on the intraperitoneal injection of the mutagen azoxymethane (AOM) and the subsequent induction of inflammation by adding dextran sodium sulphate (DSS) in the drinking water and is therefore named colitis-associated carcinogenesis (CAC). DSS is toxic to mucosal epithelial cells specifically in the colon, and the destruction of the mucosal barrier leads to chronic inflammation. In the classical protocol mice receive three DSS treatments for 5 days within 60 days. *VilCreCcna2fl/fl* mice displayed an about twofold more important weight loss at days 10 to 15 following the first DSS administration compared to control mice (Supplemental Fig 6B). We therefore decided to not expose the mice to additional DSS administrations (Supplemental Fig 6A). Mice were sacrificed sixty days after the AOM injection and colons isolated and embedded in paraffin. The grade of colitis was evaluated as described in Supplemental Table 2 (40) and was significantly enhanced in cyclin A2-deficient mice (average score of 4.5 for the mutant against 1.5 for the control animals, n=6, p≤0.05), which notably included submucosa as well as transmural disruption by immune cell infiltration that was not detectable in control mice (Fig 6A-B). The colon architecture was strongly affected in cyclin A2-deficient mice (Fig 6C) and incidence of low- and high-grade dysplasia was significantly higher in *VilCreCcna2fl/fl* mice and rather rare in control mice (Fig 6D). Moreover, the surfaces of both low- and high-grade dysplasia were significantly higher in cyclin A2-deficient *versus* control animals (Fig 6E). In addition, adenocarcinomas were only detectable in cyclin A2 mutant mice, but not in control animals (Fig 6D-E). IHC analysis revealed increased expression levels of activated β-catenin, γH2AX as well as IL-6 in cyclin A2-deficient colons by comparison with controls (Fig 7).

**Figure 6:**
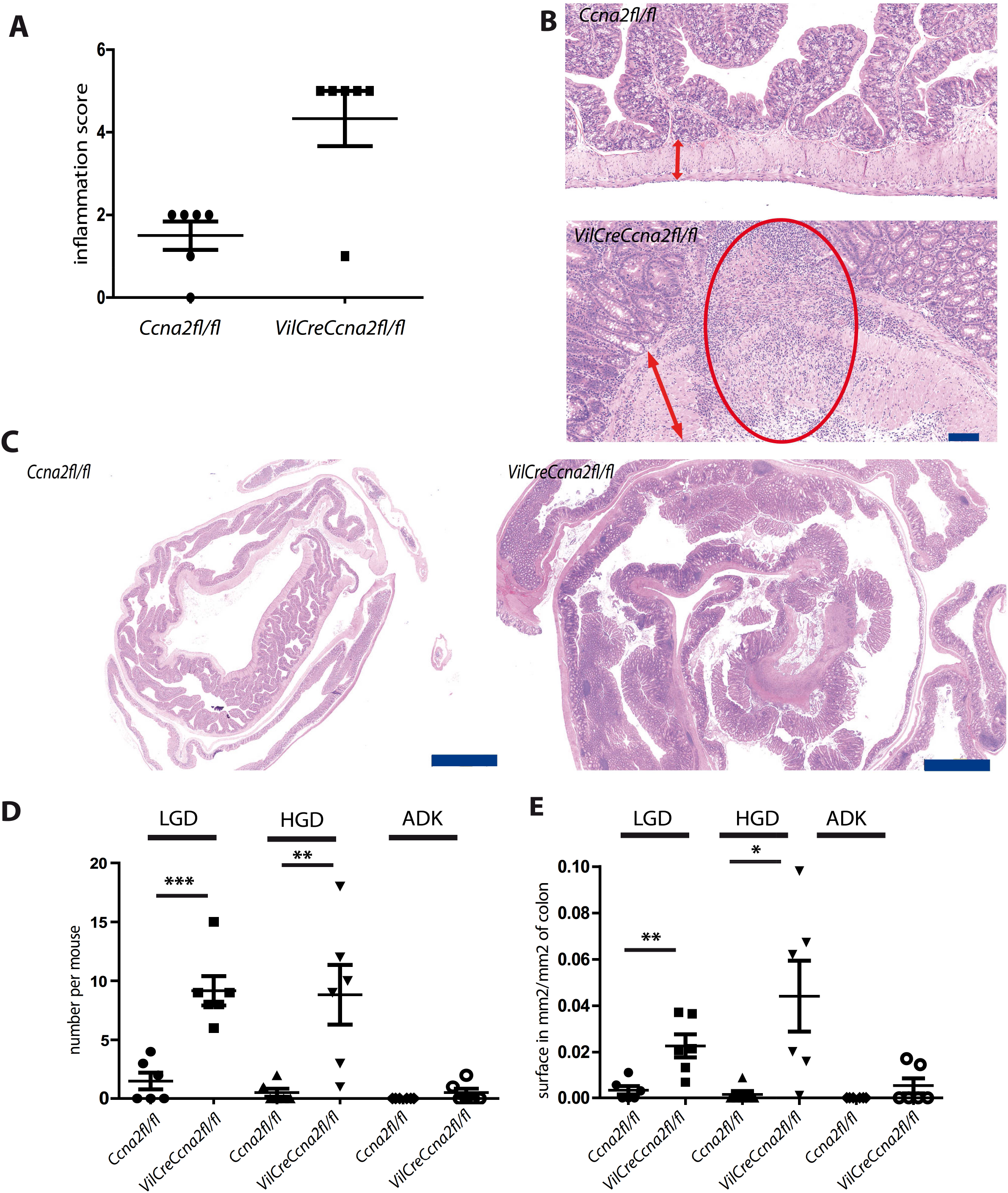
Cyclin A2-deficient mice are more susceptible to colitis-associated colorectal (CAC) cancer. *Ccna2fl/fl* (n=6) and *VilCreCcna2fl/fl* (n=7) mice were exposed to a modified AOM/DSS protocol including only one DSS administration (Supplemental Fig S6A). **(A, B).** Quantification of the inflammatory score of colons from control and *VilCreCcna2fl/fl* mice (**A**) and representative HE staining showing transmural immune cell infiltration (indicated by the red oval) in the knockout mice (**B**) (the red arrows mark the muscularis mucosa). Scale bar: 100 μm **(C)**. Representative images of HE stained colons of control and cyclin A2-deficient mice exposed to the CAC protocol. Scale bar: 2 mm. **(D, E).** Number of dysplasia and adenocarcinoma occurrence per mouse and dysplasia and adenocarcinoma surface expressed in mm^2^ per mm^2^ of colon in control and *VilCreCcna2fl/fl* mice. LGD and HGD: Low- and High-Grade Dysplasia. ADK: Adenocarcinoma. Scale bar: 2 mm. Mean ±SEM are provided**, \*** p<0,05, **\*\*** p<0.01 and **\*\*\*** p<0,001; two-tailed unpaired t-test.

**Figure 7:**
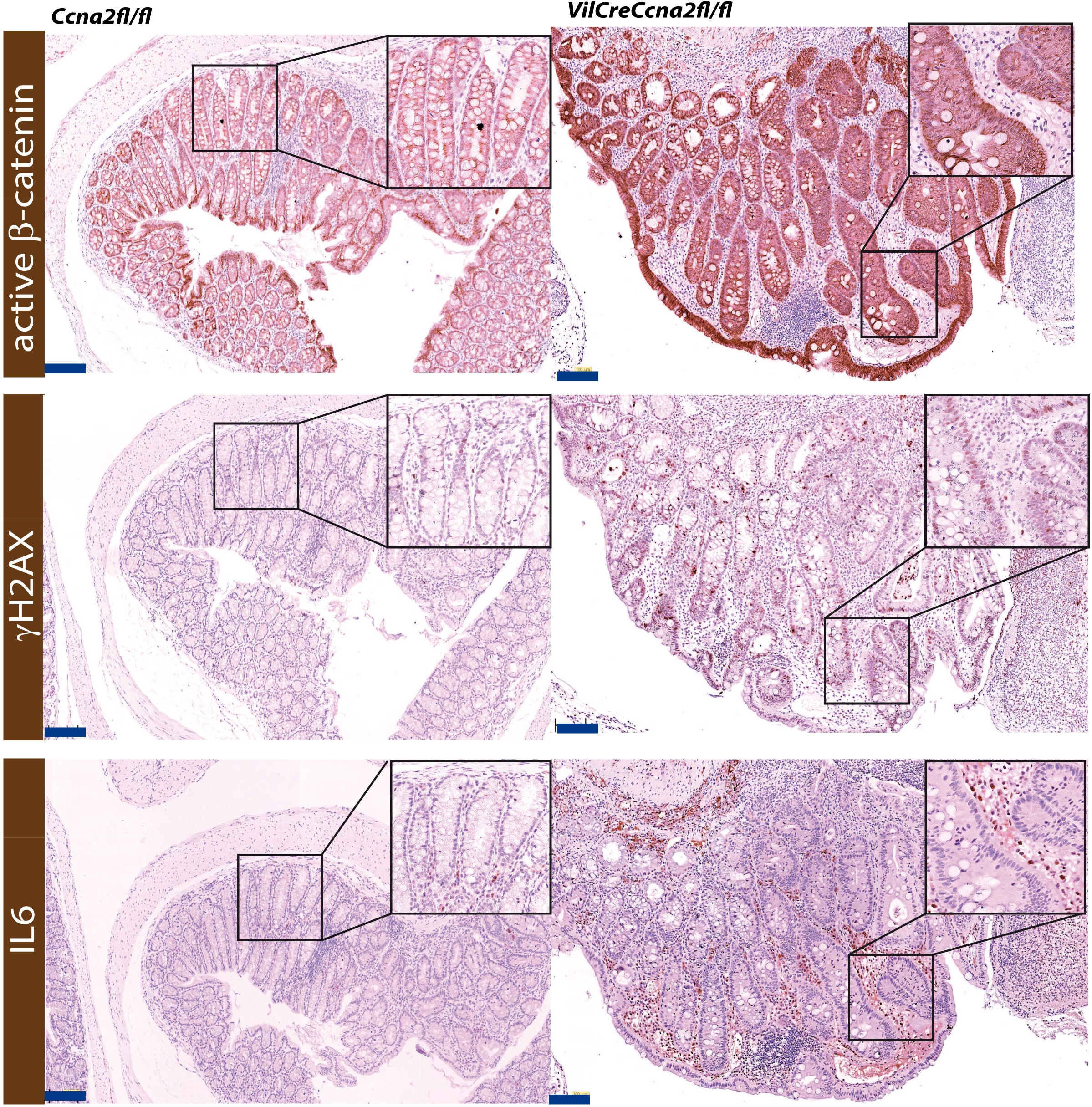
Lesions in colons from cyclin A2-deficient mice exposed to colitis-associated cancer display elevated levels of active β-catenin, DNA damage as well as IL6 expression. Representative immunostainings for active-β-catenin, γH2AX and IL6 in morphologically similar lesions in colons from control and cyclin A2-deficient mice at the end of the AOM/DSS protocol described in Supplemental Fig 6A. Blow Up: 2.5X. Scale bar: 100μm.

Altogether, these observations show that cyclin A2 deficiency in CEC promotes chemically-induced colon carcinogenesis in mice.

### Cyclin A2 expression in CRC patient has a prognostic value

Our finding that cyclin A2 deficiency in mice promotes spontaneous development of dysplasia in the colon as well as induced colon carcinogenesis prompted us to perform a meta-analysis of *CCNA2* expression on available public datasets (Supplemental Table 5). The characteristics of the 2,239 patients with CRC profiled are summarized in Table 2. Briefly, the median patients’ age was 68 years (range, 19 to 97), and the sex-ratio was balanced with 47% of females. The most frequent location was the right colon, followed by the left colon and sigmoid. The pathological stage included more stages III-IV than I-II. The pathological grade was mainly 2, and the mismatch repair (MMR) status was microsatellite stable (MSS) in most of cases. The CMS classification identified 20% of samples as CMS1 (microsatellite instability immune), 32% as CMS2 (canonical), 17% as CMS3 (metabolic), and 31% as CMS4 (mesenchymal).

**Table 2:** Clinicopathological characteristics and correlations with the *CCNA2*-based classification. MMR: Mismatch repair, MSI: Microsatellite instable, MSS: Microsatellite stable, CMS: Consensus Molecular Subtypes, RFS : Relapse Free Survival

| Characteristics |  | All | CCNA2 classes |  | p-value |
| --- | --- | --- | --- | --- | --- |
|  |  |  | Non-high | High |  |
| Patient's age |  |  |  |  | 0,484 |
|  | ≤50 years | 228 (12%) | 105 (11%) | 123 (12%) |  |
|  | >50 years | 1702 (88%) | 830 (89%) | 872 (88%) |  |
| Patients' sex |  |  |  |  | 0,417 |
|  | female | 954 (47%) | 453 (46%) | 501 (48%) |  |
|  | male | 1069 (53%) | 528 (54%) | 541 (52%) |  |
| Tumor colon location |  |  |  |  | 2,04E-02 |
|  | left | 549 (32%) | 253 (31%) | 296 (34%) |  |
|  | right | 777 (46%) | 362 (44%) | 415 (47%) |  |
|  | sigmoid | 291 (17%) | 164 (20%) | 127 (14%) |  |
|  | transversum | 77 (5%) | 35 (4%) | 42 (5%) |  |
| Pathological stage |  |  |  |  | 3,19E-08 |
|  | I-II | 834 (51%) | 347 (44%) | 487 (58%) |  |
|  | III-IV | 785 (49%) | 435 (56%) | 350 (42%) |  |
| Pathological grade |  |  |  |  | 5,99E-03 |
|  | 1 | 26 (6%) | 12 (5%) | 14 (6%) |  |
|  | 2 | 353 (77%) | 190 (83%) | 163 (71%) |  |
|  | 3 | 77 (17%) | 26 (11%) | 51 (22%) |  |
| MMR status |  |  |  |  | 1,48E-05 |
|  | MSI | 128 (19%) | 40 (12%) | 88 (25%) |  |
|  | MSS | 563 (81%) | 298 (88%) | 265 (75%) |  |
| CMS subtypes |  |  |  |  | 1,00E-06 |
|  | CMS1 | 389 (20%) | 80 (8%) | 309 (31%) |  |
|  | CMS2 | 640 (32%) | 298 (30%) | 342 (34%) |  |
|  | CMS3 | 343 (17%) | 162 (17%) | 181 (18%) |  |
|  | CMS4 | 604 (31%) | 439 (45%) | 165 (17%) |  |
| Follow-up median, months (min-max) |  | 42 (1-212) | 40 (1-212) | 42 (1-203) | 5,00E-03 |
| Events, number (%) |  | 496 (26%) | 281 (31%) | 215 (22%) | 2,32E-06 |
| 5-Year RFS [95%CI] |  | 70% [68-73] | 65% [61-69] | 75% [72-79] | 8,86E-06 |

*CCNA2* expression was heterogeneous across the 2,239 primary CRC tumors analyzed (Fig 9A). Compared to normal colon (NC) tissues, *CCNA2* transcripts are up-regulated in most of the primary tumors. In contrast, expression levels in metastases of CRC (ANOVA, p=4.16e^-54^) were significantly lower in comparison to those in primary tumors (Fig 8A). In addition, *CCNA2* transcript levels were found to be lower in the stages III/IV *versus* stages I/II (Fig 8B). In parallel, we evaluated cyclin A2 protein expression by IHC on a TMA of CRC biopsies taken from 65 patients from stage I to IV (described in Supplemental Table 3) employing two different cyclin A2 antibodies (Fig 8C and Supplemental Fig 8A; see material and methods). Cyclin A2 expression was quantified as the number of epithelial positive cells per mm^2^ of tumor area. The IHC analysis confirmed the transcriptome data by revealing cyclin A2 protein levels to be lower in stages III and IV *versus* early stages (Fig 8C and Supplemental Fig 8A). In addition, we observed lower cyclin A2 levels in stage II MSS compared with MSI tumors (Fig 8C and Supplemental Fig 8A). We next evaluated *CCNA2* levels according to the CMS consensus classification (8) and observed that *CCNA2* expression was the highest in the CMS1 class, the lowest in the CMS4 class, and intermediary in CMS2 and CMS3 classes (Fig 8E).

**Figure 8:**
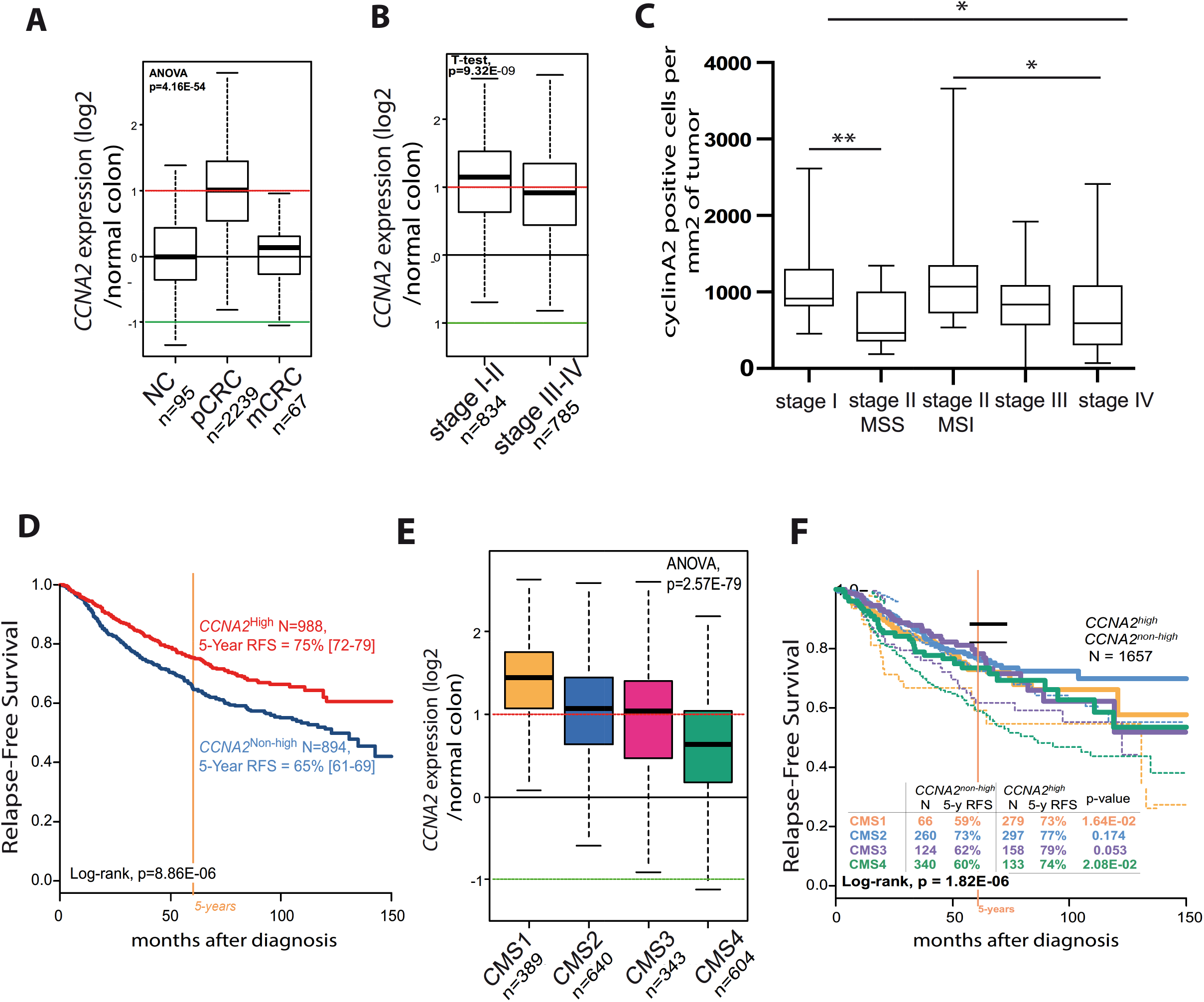
High cyclin A2 expression correlates with better prognosis in CRC patients. **(A).** Bot plots showing *CCNA2* mRNA expression levels (log^2^ over normal colon) in primary CRC tumors (pCRC; n=2239) and metastases (mCRC; n=67). Expression was normalized to normal tissue samples as in (**A**). *CCNA2*^high^ levels were defined as expression above 1 (horizontal red line) and *CCNA2*^non-high^ levels were defined as expression below 1. For each box plot, median and ranges are indicated. *CCNA2* expression was compared between group using the ANOVA (multiple comparisons, p=4.16e^-54^). **(B).** Cyclin A2 mRNA expression levels in stage I-II (n=834) and III-IV (n=785) in primary CRC tumors. Expression was normalized to normal tissue samples as in (**A**). *CCNA2* expression was compared between group using the 2-tailed Student t-test test (expressed as mean ±SD, p=9.32e^-09^). **(C).** Cyclin A2 protein expression determined by immunostaining using the anti-cyclinA2 antibody from Abcam validated on cyclin A2-deficient colon tissue and expressed as number of positive cells per mm^2^ of tumor analyzed from a TMA of CRC tumor samples derived from stage I (n=23), II (n=48 with 24 MSS tumors and 24 MSI tumors =), III (n=30) and IV (n=26) patients. (p<0.01 for the analysis between stage I and II-MSS, p<0.05 for comparison between stage I and stage IV, p=0.01 for stage II-MSI to IV). **(D).** Kaplan-Meier RFS Relapse-Free-Survival (RFS) of patients defined as carrier of *CCNA2*high (n=988) and non-high (n=894) tumors. Survivals were calculated using the Kaplan-Meier method and were compared with the log-rank test (p=8.86^e^-06). **(E).** Analysis of *CCNA2* mRNA levels in the different consensus CRC subtypes, i.e. CMS1 (n=389), CMS2 (n=640), CMS3 (343) and CMS4 (n=604) expressed as log^2^ over normal colon as described in (**A**)(p=2.57e^-79^ by the ANOVA test). **(F).** Kaplan-Meier RFS curves of all patients with pCRC according to both the *CCNA2*-based classification (*CCNA2*^high^ and *CCNA2*^non-high^ classes with plain curve and dashed curve respectively) and the CMS subtypes (p=1.64e^-02^ for the CMS1 subtype, 0.174 for CMS2, 0.053 for CMS3 and 2.08e^-02^ for CMS4).

*CCNA2* expression was then measured as discrete value after comparison with the median expression of the 95 normal colon samples; upregulation, thereafter designated “*CCNA2*^high^” (N=1139), was defined by a CRC/NC ratio ≥2 and no upregulation (“*CCNA2*^non-high^”; N=1100), by a CRC/NC ratio <2. We searched for correlations between the *CCNA2*^high^ or *CCNA2*^non-high^ classification and the clinicopathological and molecular variables of the 2239 primary CRC samples (Table 2) and found no correlation between cyclin A2 mRNA levels and patients’ age or sex. By contrast, when compared with the *CCNA2*^non-high^, the *CCNA2*^high^ class was associated (Fisher’s exact test) with more frequent right and left tumor locations (p=2.04e^-02^), pathological stages I and II (p=3.19e^-08^; Table 2 and Fig 8B), pathological grade 3 (p=5.99e^-03^) as well as MSI status (p=1.48e^-05^) and CMS1 molecular subtype (p=1.00e^-06^).

Relapse-Free-Survival (RFS) data were available for 1,882 out of 2,239 operated pCRC samples. The median follow-up was 42 months (range, 1-212); 496 patients (26%) displayed an event, and the 5-year RFS was 70% (95%CI 68-73; Supplemental Fig 7A). Interestingly, the clinical outcome was different between the two CCNA2-based classes, with 281 events (31%) in the *CCNA2*^non-high^ class (N=894) versus 215 events (22%) in the *CCNA2*^high^ class (N=988; p=2.32e-^06^, Fisher’s exact test; Table 2). The 5-year relapse-free survival (RFS) was lower (65%) for *CCNA2*^non-high^ patients (95%CI 61-69) *versus* 75% (95%CI 72-79) for the *CCNA2*^high^ class, respectively (p=8.86e^-06^; Fig 8D). In univariate analysis for RFS (Table 3), the Hazard Ratio (HR) for the event was 0.67 (95%CI 0.56-0.80) in the *CCNA2*^high^ class when compared to the *CCNA2*^non-high^ class (p=1.01e^-05^, Wald test). Other variables associated with relapse free survival (RFS) included the tumor location (p=1.35e^-06^), the pathological stage (p=2.78e^-15^), the MMR status (p=2.97e^-04^) (with the *CCNA2*^non-high^ class being predominantly associated to the MSS status therefore concurring with the results obtained with the TMA) unlike patients’ age and sex and pathological grade. In multivariate analysis (Table 3), the *CCNA2*-based classification remained associated with RFS (p=2.99e^-03^, Wald test), as well as tumor location (right and sigmoid colon), pathological stage, and MMR status. Of note, it remained significant (p=4.13e^-04^) in multivariate analysis incorporating the CMS molecular classification (Table 3). As shown in Fig 8F, the RFS of the eight classes defined by both the *CCNA2* expression and the CMS was affected (p=1.82e^-06^, log-rank test): *CCNA2* expression correlated with the clinical outcome of three CMS, with a significantly reduced RFS in *CCNA2*^non-high^ tumor patients by comparison to the *CCNA2*^high^ class for the CMS1 (59% *versus* 73, p= 1.64e^-02^), CMS3 (62% *versus* 79%, p= 0.05) and CMS4 consensus subtypes (60% *versus* 74%, p=2.08e^-02^) (Fig 8E).

**Table 3:** Univariate and multivariate prognostic analyses for RFS. MMR: Mismatch repair, MSI: Microsatellite instable, MSS: Microsatellite stable, CMS: Consensus Molecular Subtypes, RFS: Relapse Free Survival, HR: Hazard Ratio

| Variables |  | Univariate |  |  | Multivariate |  |  | Multivariate |  |  |
| --- | --- | --- | --- | --- | --- | --- | --- | --- | --- | --- |
|  |  | N | HR [95CI] | p-value | N | HR [95CI] | p-value | N | HR [95CI] | p-value |
| Patient's age | >50 vs. ≤50 years | 1630 | 1.02 [0.76-1.39] | 0,882 |  |  |  |  |  |  |
| Patients' sex | male vs. female | 1722 | 1.11 [0.92-1.34] | 0,254 |  |  |  |  |  |  |
| Tumor colon location | right vs. left | 1440 | 0.82 [0.67-1.01] | 5,09E-08 | 489 | 0.97 [0.69-1.37] | 0,880 |  |  |  |
|  | sigmoid vs. left |  | 0.33 [0.22-0.50] |  | 489 | 0.37 [0.17-0.81] | 1,28E-02 |  |  |  |
|  | transversum vs. left |  | 0.20 [0.07-0.53] |  | 489 | 0.21 [0.03-1.53] | 0,124 |  |  |  |
| Pathological stage | III-IV vs. I-II | 1374 | 2.65 [2.08-3.38] | 2,78E-15 | 489 | 2.43 [1.72-3.43] | 5,03E-07 |  |  |  |
| Pathological grade | 2 vs. 1 | 408 | 1.40 [0.51-3.84] | 0,463 |  |  |  |  |  |  |
|  | 3 vs. 1 |  | 1.87 [0.61-5.71] |  |  |  |  |  |  |  |
| MMR status | MSS vs. MSI | 606 | 2.59 [1.55-4.32] | 2,97E-04 | 489 | 2.65 [1.47-4.76] | 1,18E-03 |  |  |  |
| CMS subtypes | CMS2 vs. CMS1 | 1657 | 0.81 [0.61-1.07] | 1,44E-04 |  |  |  | 1700 | 0.73 [0.56-0.97] | 2,90E-02 |
|  | CMS3 vs. CMS1 |  | 0.99 [0.73-1.36] |  |  |  |  | 1700 | 0.95 [0.70-1.29] | 0,743 |
|  | CMS4 vs. CMS1 |  | 1.38 [1.06-1.81] |  |  |  |  | 1700 | 1.23 [0.93-1.62] | 0,151 |
| CCNA2 classes | high vs. non-high | 1882 | 0.67 [0.56-0.8] | 1,01E-05 | 489 | 0.61 [0.44-0.85] | 2,99E-03 | 1700 | 0.70 [0.58-0.85] | 4,13E-04 |

To further explore the biological alterations associated with *CCNA2* expression, we compared the whole-genome expression profiles of *CCNA2*^non-high^ (N=222) and *CCNA2*^high^ (N=237) TCGA samples (learning set) (Supplemental Fig 7B and C). We identified 92 genes differentially expressed, including 20 genes overexpressed and 72 genes underexpressed in the *CCNA2*^high^ class (Supplemental Fig 7C; Table 4). A 92-gene GES based on metagene was used to classify samples. As expected, this GES was efficient to class the 459 samples of the learning set (p=1.08e^-52^, t-test). More importantly, its robustness was validated in a completely independent validation set including the 1780 remaining samples (p=9.22e^-180^, t-test; Supplemental Fig 7B). A detailed analysis of the 92 genes revealed that the 20 most significant altered genes were upregulated involved in cell proliferation and DNA repair (mitotic cell cycle regulation, genomic instability and replicative immortality pathways) (Table 4).

**Table 4:**
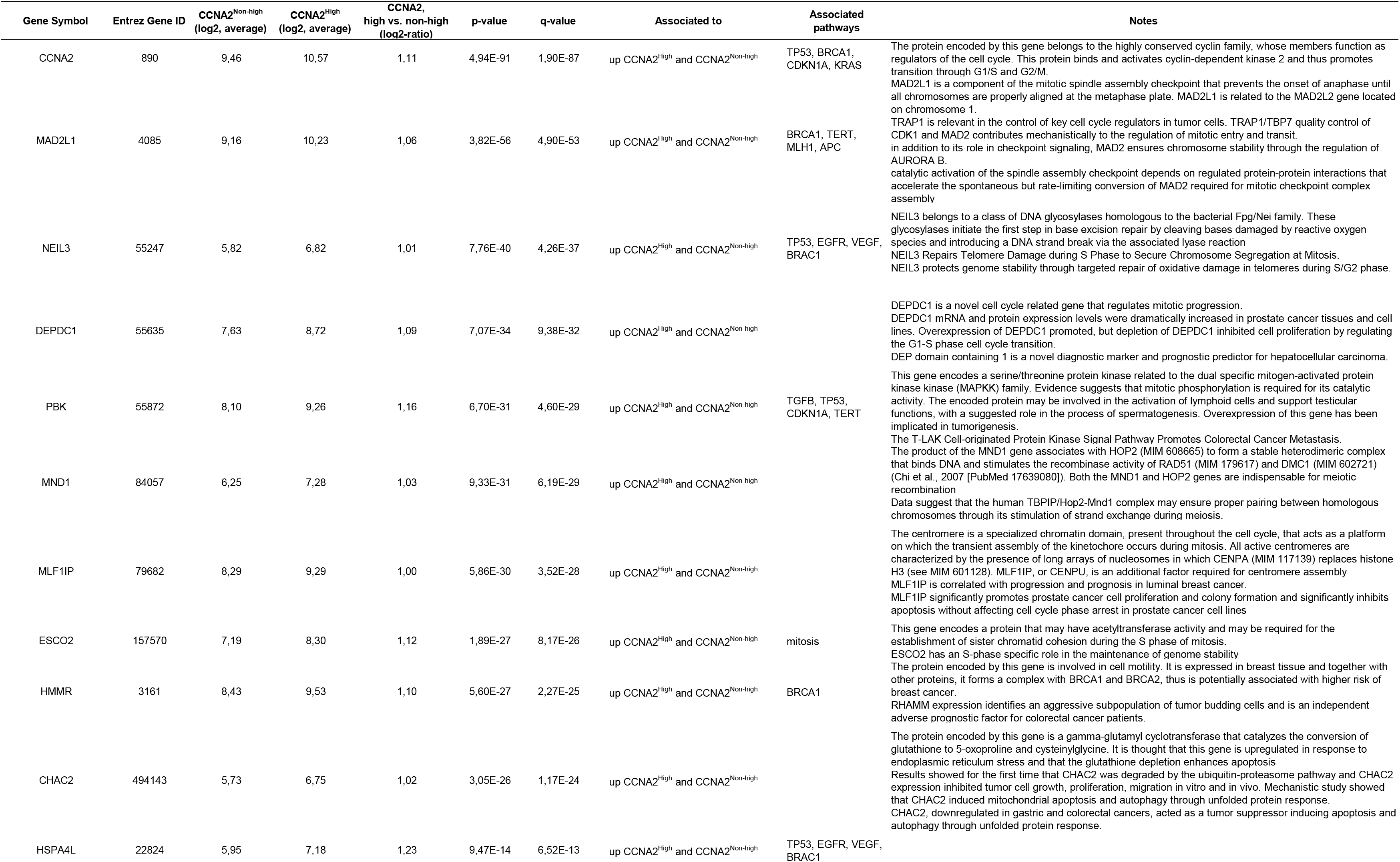
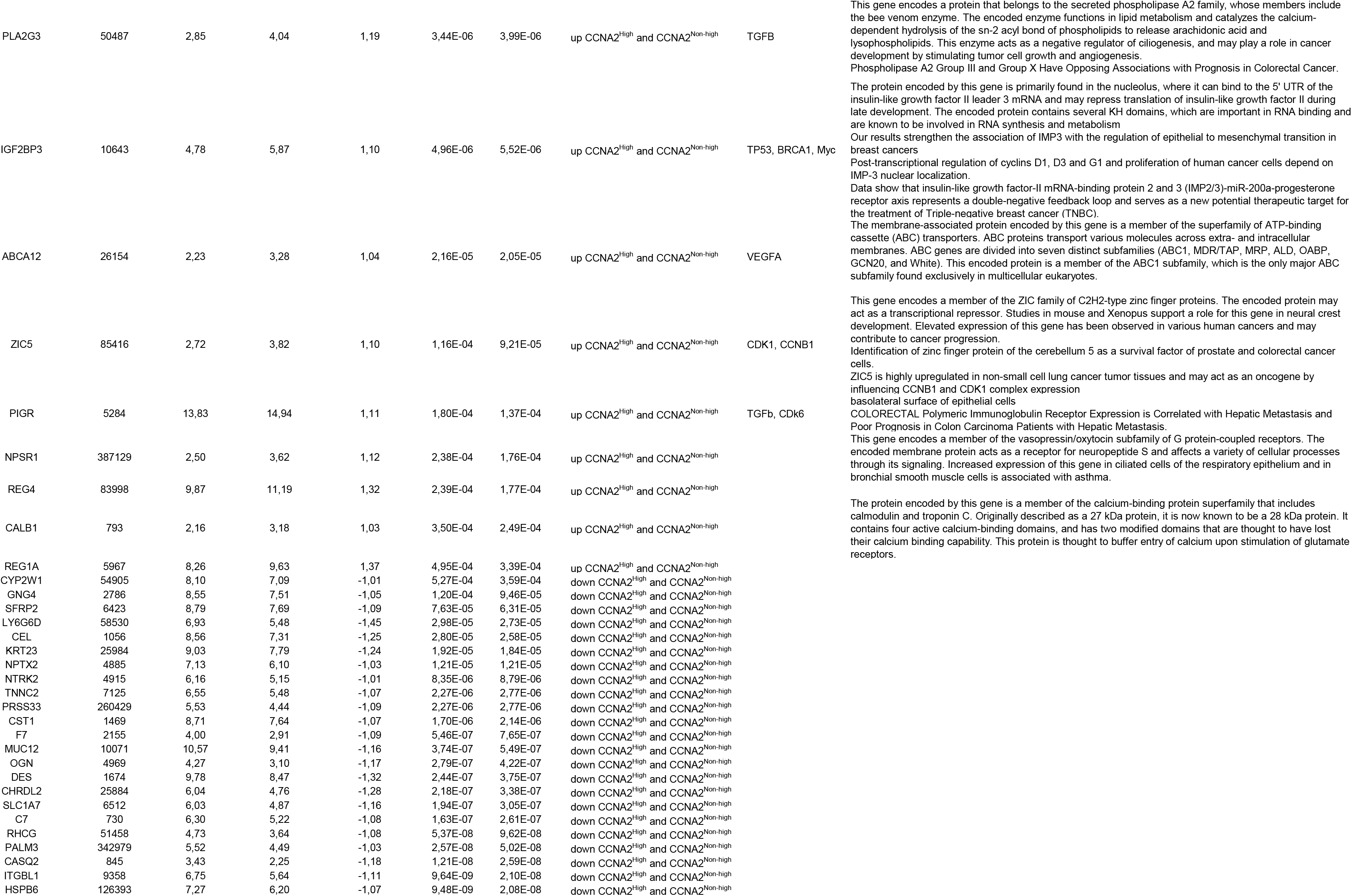
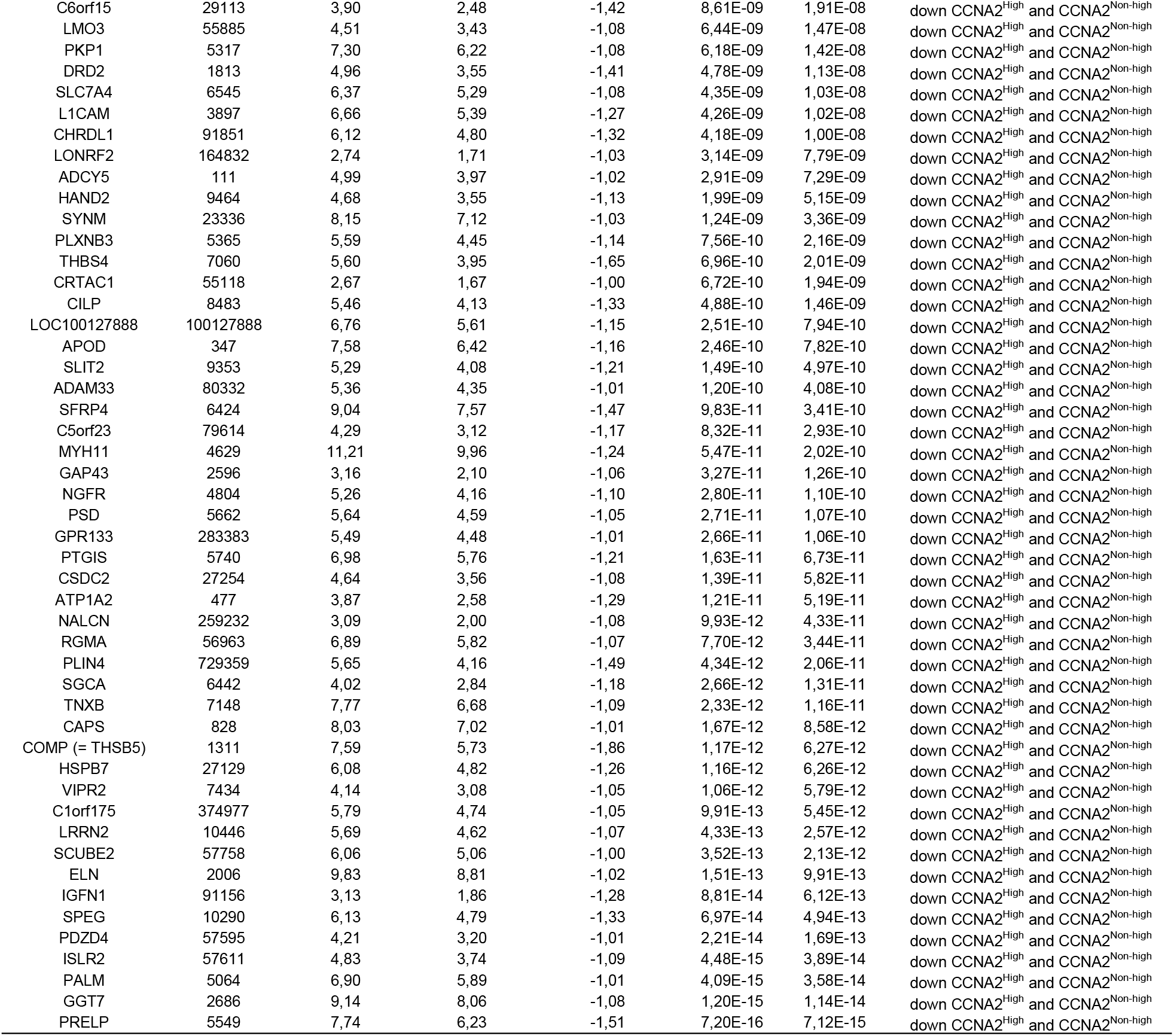
List of 92 genes differentially expressed between the *CCNA2*^High^ and *CCNA2*^Non-high^ classes

## Discussion

Cyclins are well-established drivers of the cell cycle by activating a specific family of kinases, the cyclin-dependent kinases (CDKs). Gene amplification as well as overexpression of cyclins have been frequently detected in cancer tissues, and cyclins have been proposed as biomarkers or therapeutic targets (41). For example, inhibition of cyclin A2 complexes has been shown to impair proliferation of some tumor cell lines (42).

In this report we describe that cyclin A2 deficiency in CECs induces epithelial changes in the mucosa, inflammation, increased cell proliferation and dysplasia in the colon rendering mice more susceptible to chemically-induced colon carcinogenesis. Elevated activation of β-catenin, abnormal mitosis as well as DNA DSBs are already detectable at day 4 after cyclin A2 inactivation and appear therefore to be the initial events. The IHC analysis performed suggests a spatial separation of both events in cyclin A2 deficient colons with nuclear staining of active β-catenin being more frequent in the lower part of colonic crypts, whereas γH2AX staining localized rather in the upper crypt region (Figs 4,5).

The WNT/β-catenin pathway is well established to trigger regeneration in certain tissues by controlling the stem cell compartment and has also been shown to cooperate with the YAP/TAZ pathway during colonic tissue repair and regeneration (33, 43). One study showed that cyclin A2 is involved in stabilizing the Axin/GSK3 destruction complex thus promoting β-catenin degradation (44) that could explain β-catenin nuclear accumulation detected in cyclin A2-deficient colons. In line with this is our previous description that cyclin A2 depletion in murine mammary epithelial cells promotes β-catenin activity (18, 19).

A previous study established a role of cyclin A2 in DNA DSB repair by generating mutant mice bearing cyclin A2 knockout and hypomorphic alleles resulting in lower cyclin A2 expression (45). These mice fail to up-regulate the meiotic recombination 11 (Mre11) nuclease that causes impaired resolution of stalled replication forks, reduced repair of double-stranded DNA breaks, aberrant chromosome segregation and increased spontaneous and chemically-induced tumor development (Kanakkanthara et al., 2016). The same study described that cyclin A2 binds to Mre11 mRNA thus promoting its translation. We detected increased Mre11 foci formation in cyclin A2-deficient CEC suggesting that the increased DNA damage detected in the colons of cyclin A2-deficient mice results rather from a defective regulation of DNA replication and/or chromosome segregation. Indeed, phosphorylation of CDC6 by cyclin A/CDK2 has been shown to prevent re-replication during S phase and G2 (46). Moreover, cyclin A2/CDK activity has been demonstrated to be essential for centrosome duplication and proper sister chromatid segregation (47). Of note, mitoses detected in *VilCreCcna2fl/fl* mice, were mostly located in the middle part of colonic crypts whereas γH2AX positive cells were predominantly identified in the upper part. This is in accordance with the fact that generation of DSB following abnormal mitosis takes place once a cell exits the mitotic process entering the next cell cycle (48) thus during migration toward the top part of the crypts. Notably, high expression levels of *γ*H2AX have been shown to predict a poor prognosis for CRC patients (49) and a more recent report describes a role for cyclin A2 in preventing chromosomal defects (CIN) in colon cancer cells (50).

It is well established that there is a crosstalk between DNA damage and inflammation and that either event can initiate the other one (51). Colons of cyclin A2-deficient mice display different signs of inflammation such as infiltration of macrophages and IL6 production, which correlated with an increased activation of NFκB in the colonic epithelial cells. In addition, rectal prolapse was detectable in *VilCreCcna2fl/fl* mice, a phenomenon which has been previously described in a mouse model, which develops colitis spontaneously (27). Increased expression and activation of NFκB has been observed in IBD and CRC patients, especially in mucosal macrophages and epithelial cells, accompanied by production of pro-inflammatory cytokines including IL6 (26). Finally, YAP/TAZ overexpression has been shown to promote tumor formation (52) and to correlate with poor prognosis in CRC (53). We therefore conclude that DNA damage combined with uncontrolled activation of several regenerative pathways promotes the development of dysplasia in cyclin A2-deficient mice.

Most studies evaluating the prognostic value of cyclin A2 in CRC concluded that cyclin A2 overexpression is an indicator for poor prognosis (54–56) whereas one report described a higher survival rate for patient with tumors expressing high levels of cyclin A2 (57). To clarify the prognostic value of cyclin A2 in CRC we performed a meta-analysis of 11 publicly available transcriptome datasets comprising 2239 primary tumor samples. This analysis revealed elevated *CCNA2* transcript levels in primary tumors compared with normal tissues as well as metastases concurring with our previous analysis of biopsies using IHC and Western blot analysis (17). *CCNA2* transcript expression was higher in tumors of stage I and II patients compared with those of stage III and IV. Importantly, this concurs with cyclin A2 protein expression analysis by IHC on our TMA. For the first time, we described that *CCNA2* expression differs among CMS subtypes, with the highest in CMS1 and lowest in CMS4 consensus subtype. A stratification of the transcriptome samples in high and non-high expressers by taking the *CCNA2* transcript expression ratio of CRC *versus* normal tissue ≥2 as a cut-off displayed a higher RFS in the *CCNA2*^high^ class. Importantly, such favorable prognostic value of high *CCNA2* expression persisted in multivariate analysis incorporating the classical prognostic factors or the CMS subtypes, suggesting independent prognostic value. Whole-transcriptome supervised analysis of the two CCNA2 classes identified that the 20 most significant modulated genes upregulated in the *CCNA2*^high^ versus the *CCNA2*^non-high^ class were all involved in cell proliferation and DNA repair. Though expression data of multicellular tumor tissues cannot be directly correlated with the ones obtained in cell type specific knockout mice, it is remarkable that CCNA2^high^ CRC tumors and cyclin A2 deficiency in murine CEC trigger similar mechanisms such as DNA damage. We conclude that the expression pattern of cyclin A2 in CRC tissues mirrors distinct roles during colon carcinogenesis, such as driving cell proliferation in early stages, when highly expressed, but promoting aggressiveness in later stages, when expression levels are lower.

Taken together, our results establish that cyclin A2 deficiency in murine colonic epithelial cells induces DNA damage, inflammation and activation of different regenerative pathways triggering dysplasia. This renders mice more susceptible to chemically-induced colon carcinogenesis by elevated tumor initiation as well as aggressiveness. The observations made in mice are complemented by a meta-analysis of 2239 primary CRC tumors displaying that high *CCNA2* expression is associated with better prognosis in CRC patients. Moreover, we describe here for the first time that *CCNA2* expression differs between the previously described CMS subtypes. Importantly, dissecting the patient cohort into the consensus molecular subtypes identifies *CCNA2*^non-high^ expression as a bad prognosis factor within CMS1, CMS3 and CMS4, but not CMS2. In conclusion, *CCNA2* expression was strongly correlated with several clinicopathological factors of CRC and may represent a favorable prognostic marker in patients with CRC.

## Methods

### Generation of *cyclin A2* conditional knockout mice in the intestinal epithelium

To generate *cyclin A2* intestinal epithelium-specific knockout mice, *Ccna2 fl/fl* mice (23) were crossed with transgenic mice carrying either a constitutively active Cre recombinase (*vil-Cre*) (24), which is expressed from the embryonic development (day 9) throughout adulthood in intestinal epithelial cells, or a tamoxifen-inducible Cre under the same promoter (*vil-Cre-ER^T2^* ^)^ (24)), which was induced in adult mice by two intraperitoneal injections of tamoxifen (Sigma, 150 mg/kg). Mice were maintained on a C57BL6/129SV background for the *cyclin A2 vil-Cre* strain and C57BL/6 genetic background for the *cyclin A2 vil-Cre-ER^T2^* mice. Experimental groups were caged together according to gender. Genomic DNA from mice tails and colonic epithelial cells was used for genotyping using the primers listed in Supplemental Table 1.

### Immunohistochemical and immunofluorescence analyses on paraffin tissue section

Isolated colons specimens were rolled up lengthwise and embedded in paraffin after fixation in 10% formalin (DiaPATH F0046, MM France) for 24 hours. Immunohistochemistry was performed on 4-μm sections. Tissue slides were deparaffinized and rehydrated by performing the following washes: xylene (2×5 min), 100 % ethanol (5 min), 96% ethanol (3 min), 70% ethanol (3 min) and dH_2_O (5 min). Tissue slides were incubated with 10 mM sodium citrate pH6.0 (T0050, DiaPATH) for 20 min at 100°C or Tris 10mM-EDTA 1mM pH9.0, for antigen retrieval, depending on the antibody. Following two washes in PBS-Tween 0.2%, tissue slides were incubated in PBS-3% hydrogen (#H1009, Sigma) for 5 min at room temperature for peroxidase inactivation. After blocking in 2.5 % blocking serum-5% BSA-5% non-fat milk for 30 min at room temperature, tissue slides were incubated with primary antibody for 1 hour at 4°C. Then corresponding secondary antibody reagents (ImmPRESS^TM^ kit, Vector Laboratories), directed against rabbit, rat or mouse antibodies were used for detection. The following antibodies were used: rabbit anti-Cyclin A2 (Abcam, ab181591, 1 :1000 dilution), rabbit anti-Ki67 (Abcam, ab16667, 1 :200 dilution), rat anti-F4/80 (Hycult biotech, HM1066 1 :800 dilution), rabbit anti-non-phospho (Active) β-catenin (Cell signaling, #8814, 1 :1000 dilution), rabbit anti-γH2AX (Abcam, ab11174, 1 :500), rat anti-IL6 (Novus Biological, IL6/1270, 1:100), rabbit anti-TAZ (Novus Biological, NBP1-85067, 1:100).

For hematoxylin and eosin staining, tissue slides were incubated in Xylene (3×3 min), 100% ethanol (3×3 min), 95% ethanol (3 min), 80% ethanol (3 min) and 5 min in deionized water. Slides were then incubated in hematoxylin (Vector labs) for 3 min, extensively rinsed in tap water and dipped 8 to 12 times in acid ethanol (0.03N HCl-70% ethanol). After rinsing in tap water, tissue slides were transferred in deionized water for 2 minutes then stained in Eosin for 30 second (Vector Labs) followed by three incubations of 5 min in 95% ethanol then 100% ethanol. After three incubations in Xylene, coverslips were mounted using Permount media (Fisher Scientific). For the tissue microarray (TMA) analysis, the rabbit anti-Cyclin A2 antibody (Abcam, ab181591, 1:1000 dilution), validated on *Ccna2*-deficient mouse tissues as well as rabbit anti-Cyclin A2 from Novocastra (NCL-CYCLINA), were used.

To carry out immunofluorescence analysis, 4µm thick tissue slides were deparaffinized and rehydrated as described above and were incubated in 1mM EDTA antigen retrieval buffer (pH9.0) for the alpha and gamma-tubulin, or in 10 mM citrate (pH6) for the anti-Mre11 staining. Blocking was performed for 30 minutes in 2,5 % blocking serum-5% BSA-5% non-fat milk at room temperature, and tissue slides were then incubated with primary antibodies (anti-Me11, Novus Biologicals NB-100-142, 1:500 dilution; anti-alpha-tubulin, Novus Biologicals NB-600-506, 1:1000 dilution, anti-gamma tubulin, Sigma, T3320, 1:1000 dilution) for 1 hour at 4°C. Depending on primary antibodies, slides were incubated with goat anti-mouse, rat or rabbit secondary antibodies conjugated to either Alexa-488 or Alexa-555 (ThermoFisher Scientific) in addition with DAPI (0.1 μg/ml) for 1 hour at room temperature. Acquisition of the Mre11 staining was performed on the Leica confocal SP5-SMD and images were analyzed using Imaris software. Quantification of the Mre11 staining was performed using ImageJ. For the alpha/gamma-tubulin co-staining, images acquisition in the proximal colon was performed using Metamorph software on the Upright Zeiss Axioimager Z2 and images were analyzed using ImageJ software.

### BrdU incorporation assay

For proliferation studies, mice were intraperitoneally injected with 100μg/g of Bromodeoxyuridine (BrdU, B9285, Sigma) diluted in PBS. Colon specimens were dissected 2 hours after injection, flushed with cold PBS, and embedded in paraffin after fixation in 10% formalin for 24hours. Incorporation of BrdU in proliferating intestinal epithelial cells was detected using an anti-BrdU antibody (Biolegend, 1:100 dilution) after deparaffinization of the tissues, antigen retrieval in citrate buffer, as described above, and DNA denaturation using 2N HCl for 1hour at 37°C followed by an incubation in 0.1M borax buffer pH9. Revelation was performed using the Avidin/Biotin Vectastain System kit (Vectorlab, USA) according to the protocol.

### Isolation of colonic epithelial cells (CEC)

Colons were removed from mice and flushed with cold PBS. Fat and adherent connective tissue were taken out, colons were cut longitudinally then placed in 10 ml of cold wash buffer (PBS, 2% FBS). Colons were treated with 10 ml of CEC buffer (PBS, 1% BSA, 1 mM EDTA, 1 mM DTT, 5.6 mM Glucose) at 37°C for 45 minutes under continuous shaking to release CECs. Purity of isolated cells was validated by FACS by using the cell surface markers EpCAM-APC (Biolegend, 118213, 1:50 dilution) and CD45-PE (Biolegend, 103105, 1:100 dilution), CD90.2-FITC (BD, 553013, 1:50 dilution) to identify epithelial and immune cells respectively.

### Isolation of CEC cytoplasmic and nuclear fractions

Cells were treated with ice-cold hypotonic buffer (10 mM Hepes pH 7.9, 10 mM KCl, 0.1 mM EDTA, 0.1 mM EGTA, 0.5 mM PMSF, 1 mM DTT and 1×Complete inhibitor tabs, Roche, for 15 min), then treated with 10% NP40 and vortexed. The nuclei were pelleted by centrifugation (14,000 rpm, 15s) while the supernatant was kept as the cytoplasmic fraction. Nuclei were lysed in 20 mM Hepes pH 7.9, 0.4 mM NaCl, 1 mM EDTA, 1 mM EGTA, 0.5 mM PMSF, 1 mM DTT, 1×complete inhibitor tabs (Sigma). The supernatant was collected as the nucleoplasmic fraction after centrifugation (14,000 rpm, 5min).

### Western blot analysis

Protein samples were prepared in 4xLaemmli buffer, separated in sodium dodecyl sulfate-polyacrylamide gels and electro blotted onto nitrocellulose membranes (Millipore) using a wet/tank transfer system (Bio-Rad, Hercules, CA). The following antibodies were used in this study: anti-Cyclin A2 (Abcam, ab181591, 1:1000 dilution), anti-NF-kB p65 (Cell Signaling, #8242, 1:1000 dilution) and anti-non-phospho (Active) β-catenin (Cell Signaling, #8814, 1:1000 dilution). All the experiments were performed in triplicate and at least three times. Quantification of at least three immunoblots was carried out using the ImageJ software (National Institute of Health). Experimental values are expressed as a ratio of protein levels over control with the TFIIB (Biolegend, 1:2000 dilution) and Histone H3 (Abcam, Ab1791, 1:1000 dilution) signal for nucleoplasmic fraction or β-actin (Sigma, 1:5000 dilution) for total cell lysates.

### RNA sequencing samples preparation and bioinformatic analysis

CEC were isolated as described above from the proximal colon part from 4 weeks-old *Ccna2 fl/fl vil-Cre* (KO) and *Ccna2 fl/fl* (control) mice. RNA was extracted using the Roche High Pure RNA isolation kit (Roche Molecular Systems, Inc.) according to the manufacturer protocol and purity as well as integrity controlled using the Agilent RNA 6000 nano kit (Agilent Technologies). RNA sequencing was performed on an Illumina NextSeq500™ platform. The quantification of the gene transcripts was performed using the Kallisto program (58), which is based on pseudo-alignment principle, with default settings. GRCm38 Refseq was used to annotate the transcripts. We used the STAR program (59) to align the reads on the *Mus musculus* reference genome (GRCm38/mm10). For the differential expression analysis, we first filtered the transcripts with too low counts among the samples by using the *filterByExpr* function of edgeR, with default settings. To perform the data normalization and differential expression analysis (DEA) of *Ccna2* deficient *versus* control samples, we ran standard steps of edgeR and DESeq2 in parallel on the filtered data. For both DEA methods, we applied a cutoff threshold of 0.05 on the adjusted p-values and a cutoff threshold of 2 on fold changes. The two resulting lists of significantly differentially expressed genes were intersected. We separated the up-regulated and the down-regulated genes of this unique list. To perform the functional enrichment analysis, these sub-lists were finally submitted to the dedicated function of the gProfileR R package, available on Bioconductor. The R code used is provided in the supplementary data.

### Colitis-associated carcinogenesis

For induction of colitis-associated carcinogenesis in the *VilCrecyclinA2fl/fl* mice and their corresponding controls, mice were intraperitoneally injected with 6.25 mg/kg azoxymethane (AOM; Sigma). Mice were treated by one cycle of 5 days with 1.75% (w/v) dextran sodium sulphate (DSS; TdB Consultancy AB) administered in the drinking water at day 5 of the protocol. Weight of the mice was checked every two days for two months, except for the week following the DSS treatment where the weight was measured every day. Mice were then sacrificed, and colon specimens prepared for histological analysis after fixation in 10% neutral buffered formalin. Tissues were embedded in paraffin, and 4-μm thick sections were stained with hematoxylin and eosin or incubated with the following primary antibodies (anti-active β-catenin, anti-γH2AX, and anti-IL6) as above reported. Colitis was scored as outlined in Supplemental Table 2.

### Patient samples

Details of the TMA generated at CRB–ICM (agreement number BB-033-00059) in Montpellier (France) are listed in Supplemental Table 3. Some tissue samples were generated as duplicates from the same patients but considered as individual templates, for the statistical analysis, due to tumor heterogeneity.

For the transcriptome meta-analysis, we collected clinicopathological and gene expression data of tissues from normal colon (NC), primary colorectal adenocarcinoma samples (pCRC) and metastases from CRC samples (mCRC) from ten public data sets and one data set generated at the Institut Paoli Calmettes (Marseille, France) comprising at least one probe set representing *CCNA2*. Sets and raw data were collected from the National Center for Biotechnology Information (NCBI)/Genbank GEO, ArrayExpress and TCGA databases (Supplemental Table 5). Samples were profiled using whole-genome DNA microarrays (Affymetrix) or RNA sequencing (Illumina). The analyzed data set contained a total of 2,401 samples, including 95 normal (NC) samples, 2,239 primary tumor (pCRC) samples, and 67 metastases (mCRC) samples included in the present analysis (mCRC, located in the liver (n=47) and lung (n=20)).

### Gene expression data analysis

Data analysis required a step of pre-analytic processing. We first normalized each data set separately, by using Robust Multichip Average (RMA) (60) with the non-parametric quantile algorithm for the raw Affymetrix data. Normalization was done in R using Bioconductor and associated packages. We then mapped hybridization probes across the different microarrays represented as previously reported (61). When multiple probes mapped to the same GeneID, we retained the one with the highest variance in each data set. We log2-transformed the available TCGA RNAseq data that were already normalized. Next, we extracted *CCNA2* mRNA expression and corrected the batch effects through the 11 studies using z-score normalization. Briefly, for each expression value in each study separately, *CCNA2* values were transformed by subtracting the mean of the gene in that data set divided by its standard deviation, mean and standard deviation being measured on primary samples. CCNA2 expression was measured as discrete value after comparison with median expression in the 95 NC samples; upregulation, thereafter designated “CCNA2^high^” was defined by a CRC/NC ratio ≥2 and no upregulation (“CCNA2^non-high^”) by a CRC/NC ratio <2. The Consensus Molecular Subtype (CMS) classification (8) was based on the tool CMScaller made by Eide et al. (62). Finally, to explore the biological pathways associated with our CCNA2-based classification, we applied a supervised analysis using learning and validation sets. The learning set included the 459 samples of the TCGA data set that included 222 CCNA2^non-high^ and 237 CCNA2^high^ samples. We used a moderated t-test with empirical Bayes statistic included in the limma R packages. False discovery rate (FDR) (63) was applied to correct the multiple testing hypothesis and significant genes were defined by the following thresholds: p<5%, q<25% and fold change (FC) superior to |2x|. The resulting gene expression signature (GES) was based on a metagene score defined as the difference between mean expression of genes upregulated and mean expression of genes downregulated in the CCNA2^high^ samples and using a cut-off equal to 0. This score was then applied to both learning and validation sets to test the robustness of the GES (t-test). Ontology analysis of the gene list was based on GO biological processes of the Database for Annotation, Visualization and Integrated Discovery (DAVID; http://david.abcc.ncifcrf.gov/).

### Statistical analysis

Correlations between the CCNA2 expression-based classes (non-high *versus* high) and the clinicopathological factors were calculated with the Fisher’s exact test for the binary variables and the Student’s t-test for the continuous variables. Our primary endpoint, relapse-free survival (RFS), was calculated from the date of diagnosis until the date of metastatic relapse or death from CRC. The follow-up was measured from the date of diagnosis to the date of last news for event-free patients. Survival was calculated using the Kaplan-Meier method and curves were compared with the log-rank test. Univariate and multivariate analyses were done using Cox regression analysis (Wald test). The variables tested in univariate analysis included the CCNA2-based classification (non-high *versus* high), patients’ age and sex, tumor location, pathological stage and grade, MMR status, and the CMS classification. Multivariate analysis incorporated all variables with a p-value inferior to 5% in univariate analysis. All statistical tests were two-sided at the 5% level of significance. Statistical analysis was done using the survival package (version 2.30) in the R software (version 2.15.2). This article is written in accordance with reporting recommendations for tumor marker prognostic studies criteria (REMARK) (64).

Statistical analysis of quantifications for the mouse study was performed in GraphPad using an unpaired t-test except for the Nuclear size distribution where the Kolmogorov-Smirnov test using R software was performed.

### Animal study approval

Mouse experiments were performed in strict accordance with the guidelines of the European Community (86/609/EEC), the French National Committee (87/848) for care and use of laboratory animals and were approved by the Regional Ethics committee under the 01556.02 reference number.

## Supporting information

YGuo et al Supplemental Figures and tables

## Acknowledgments

We are grateful to the excellent platforms in Montpellier, i.e. RIO imaging platform, the histology and animal experimentation platforms RHEM and RAM, as well as the IGMM mouse facility. Many thanks to Thierry Gostan for his help on the statistical analysis and Ula Hibner for carefully reading the manuscript. We are grateful to Maguy Del Rio for her help on cyclin A2 mRNA expression in patients. This work was supported by INCA (project 2013-111), by the SIRIC Montpellier Cancer Grant INCa_Inserm_DGOS_12553 and La Ligue contre Cancer-comité 34 (BL).

## Author’s contribution

YG, MG, CP, VP, RT, HT, JB, CM, QDS, PF, FB-M, CL, SG, RA, EM, BL: conducting experiments, acquiring data and analyzing data.

EM, BL, MH: designing research studies and analyzing data.

YG, FB, JMB, PS, EM, BL, MH: writing manuscript.

---

**Supplemental Figure 1: Cyclin A2-deficiency in colonic epithelial cells induces architectural changes in the mucosa and inflammation. (A).** HE staining illustrating the subdivision of the colon in 3 parts for the histological analysis, i.e. the distal part close to the rectum, transverse and proximal part, close to the caecum. (B). Quantification of the F4/80 staining in control (n=3), constitutive (n=3) and inducible mutant mice (n=4; expressed as mean ±SEM, *p<0.05; two-tailed unpaired t-test) expressed as number of F4/80 positive cells per mm^2^ of colon tissue. (C). Representative HE staining of irregular formed crypts in colons from a constitutive and inducible (day8 following tamoxifen injection) knockout mouse by comparison to colons from control animals. Scale bar: 100μm.

**Supplemental Figure 2: RNA-seq analysis of altered expression of genes involved in cell cycle regulation and DNA double-strand break repair in cyclin A2 deficient colonic epithelial cells (CEC). (A).** Sashimi plot of *Ccna2* reads for each mouse-derived CEC samples analyzed by RNA-seq. Plots show deletion of exons 2 to 7 in transcripts of CEC from *VilCreCcna2fl/fl* mice compared to controls. The coverage for each alignment track is plotted as a bar graph. Arcs representing splice junctions connect exons. Arcs display the number of reads split across the junction (junction depth). Genomic coordinates and the gene annotation track are shown below the junction tracks. (B). Gene Ontology over-representation analysis showing the genes up-regulated in *Ccna2 fl/fl* mutant mice compared to controls. The top 20 of Gene Ontology terms (Biological Process) are shown here. The p-values have been corrected for multiple testing by the Benjamini-Hochberg method. In this dot plot, the color represents the adjusted p-values and the dot size represents the number of genes for each term. The gene ratio is shown on the x-axis. (C). Alterations in the double-strand break KEGG pathway in cyclin A2 deficient CEC. Genes up-regulated in *VilCreCcna2fl/fl* mice samples are in red, whereas downregulated genes are in blue. The scale indicated on the figure represent the log2 Fold change. In grey are the genes absent from our data. For all analyses, only p-values <0.05 were considered as statistically significant. FC: Fold change; DSB: Double-strand break

**Supplemental Figure 3: Increased proliferation of colonic epithelial cells from cyclin A2-deficient mice. (A, B, C).** Quantification of Ki67 expression by IHC of the different parts of crypts from the distal and proximal part of the colon from control (n= 66 crypts analyzed from 3 different mice) and constitutive cyclin A2-deficient mice (**A, B**) and proximal part of the colon from induced knockout mice at day 8 following inactivation (**C**) compared to controls (n=120 from 3 different mice). Mean values ±SEM are provided, **\*** p<0,05, **\*\*** p<0.01 and **\*\*\*** p<0,001; two-tailed unpaired t-test.

**Supplemental Figure 4: Increased nuclear size of colonic epithelial cells in cyclin A2-deficient mice.** Distribution curve representing the nuclear size of *Ccna2fl/fl* colonic epithelial cells (white, n=398 for the left panel and 578 for the right panel from 3 mice), *VilCreCcna2fl/fl* (black, left panel, n=400 from 3 mice, p<1.10e^-16^) and *VilCreERT2Ccna2fl/fl* (black, right panel, n=566, p<1.10e^-16^) colonic epithelial cells; p-values were determined using Kolmogorov-Smirnov test.

**Supplemental Figure 5: Examples of mitoses in colons of cyclin A2-deficient mice. (A).** Immunofluorescence analysis of mitosis (indicated by red circles) using an anti-α-tubulin antibody (green) to stain for the mitotic spindle in combination with γ-tubulin (centrosome in red) and DAPI for DNA in a colon of *VilCreERT2Ccna2fl/fl* mice at day 8 following inactivation (right panel) by comparison to controls (left panel). Scale bar: 100μm. **(B)**. The upper image shows normal mitosis, the lower panel several examples of abnormal mitoses observed in a *Ccna2*-deficient colon. Blow Up: 2.5X

**Supplemental Figure 6: Protocol and weight monitoring of the mice during the colitis-associated carcinogenesis. (A).** Schematic representation of the modified AOM/DSS protocol applied to *VilCreCcna2fl/fl* and control mice (see Figs 6 and 7). **(B).** Monitoring of the relative weight (expressed as percentage relative to the weight at the beginning of the protocol) of *VilCreCcna2fl/fl* and control mice during the AOM/DSS protocol.

**Supplemental Figure 7: Cyclin A2 expression at the mRNA and protein level in CRC patients. (A).** Relapse-Free-Survival (RFS) curve of the overall patients analyzed for cyclin A2 mRNA expression levels**(B).** Metagene-based prediction score of outcome (using Student t-test and expressed as mean ±SD) of the *CCNA2*^high^ samples compared to those of *CCNA2*^non-high^ samples in the learning set (left) and in the independent validation set (right). **(C).** Volcano plot showing the 92 genes differentially expressed in the learning set (TCGA). Genes up-regulated in the *CCNA2*^high^ samples are colored in red and genes down-regulated in green.

**Supplemental Figure 8: Cyclin A2 protein expression in CRC tumor samples from different stages**. **(A).** Cyclin A2 expression analyzed on the same TMA shown in Figure 8, but using a different anti-cyclin A2 antibody from Novocastra. (Mean values ±SEM, p<0.05 for the analysis between stage I and II-MSS, p<0.01 for comparison between stage I and stage III, p<0.001 for stage I to IV, p<0.01 for stage II-MSI to stage IV, two-tailed unpaired t-test). **(B).** Representative immunostaining of stage I, II-MSS, II-MSI, III and IV tumor samples. Scale bar: 100μm

## References

1. http://gco.iarc.fr/today/fact-sheets-cancers

2. Zlobec I, Lugli A. Prognostic and predictive factors in colorectal cancer. J. Clin. Pathol. 2008;61(5):561–569.

3. Cuyle P-J, Prenen H. Current and future biomarkers in the treatment of colorectal cancer. Acta Clin. Belg. 2017;72(2):103–115.

4. Kannarkatt J, Joseph J, Kurniali PC, Al-Janadi A, Hrinczenko B. Adjuvant Chemotherapy for Stage II Colon Cancer: A Clinical Dilemma. J. Oncol. Pract. 2017;13(4):233–241.

5. McCleary NJ, Benson AB, Dienstmann R. Personalizing Adjuvant Therapy for Stage II/III Colorectal Cancer. Am. Soc. Clin. Oncol. Educ. Book Am. Soc. Clin. Oncol. Annu. Meet. 2017;37:232–245.

6. Beaugerie L et al. High risk of anal and rectal cancer in patients with anal and/or perianal Crohn’s disease. Clin. Gastroenterol. Hepatol. Off. Clin. Pract. J. Am. Gastroenterol. Assoc. [published online ahead of print: November 30, 2017]; doi:10.1016/j.cgh.2017.11.041

7. Ma H et al. Pathology and genetics of hereditary colorectal cancer. Pathology (Phila.) 2018;50(1):49–59.

8. Guinney J et al. The consensus molecular subtypes of colorectal cancer. Nat. Med. 2015;21(11):1350–1356.

9. Yasmeen A, Berdel WE, Serve H, Müller-Tidow C. E- and A-type cyclins as markers for cancer diagnosis and prognosis. Expert Rev. Mol. Diagn. 2003;3(5):617–633.

10. Bendris N et al. Cyclin A2: a genuine cell cycle regulator?. Biomol. Concepts 2012;3(6):535–543.

11. Blanchard JM. To be or not to be a proliferation marker?. Oncogene 2014;33(8):954–955.

12. Jirawatnotai S et al. A function for cyclin D1 in DNA repair uncovered by protein interactome analyses in human cancers. Nature 2011;474(7350):230–234.

13. Li Z et al. Cyclin D1 induction of cellular migration requires p27(KIP1). Cancer Res. 2006;66(20):9986–9994.

14. Li Z et al. Cyclin D1 regulates cellular migration through the inhibition of thrombospondin 1 and ROCK signaling. Mol. Cell. Biol. 2006;26(11):4240–4256.

15. Li Z, Wang C, Prendergast GC, Pestell RG. Cyclin D1 functions in cell migration. Cell Cycle Georget. Tex 2006;5(21):2440–2442.

16. McAllister SS, Becker-Hapak M, Pintucci G, Pagano M, Dowdy SF. Novel p27(kip1) C-terminal scatter domain mediates Rac-dependent cell migration independent of cell cycle arrest functions. Mol. Cell. Biol. 2003;23(1):216–228.

17. Arsic N et al. A novel function for Cyclin A2: control of cell invasion via RhoA signalling. J. Cell Biol. 2011;in press.

18. Bendris N et al. Cyclin A2, a novel regulator of EMT [Internet]. Cell Mol Life Sci [published online ahead of print: May 31, 2014];http://www.ncbi.nlm.nih.gov/entrez/query.fcgi?cmd=Retrieve&db=PubMed&dopt=Citation&list_uids=24879294. cited

19. Cheung CT et al. Cyclin A2 modulates EMT via β-catenin and phospholipase C pathways. Carcinogenesis [published online ahead of print: May 19, 2015]; doi:10.1093/carcin/bgv069

20. Wang D et al. Prefoldin 1 promotes EMT and lung cancer progression by suppressing cyclin A expression. Oncogene 2017;36(7):885–898.

21. Kanakkanthara A et al. Cyclin A2 is an RNA binding protein that controls Mre11 mRNA translation. Science 2016;353(6307):1549–1552.

22. Gopinathan L et al. Loss of Cdk2 and cyclin A2 impairs cell proliferation and tumorigenesis. Cancer Res. 2014;74(14):3870–3879.

23. Kalaszczynska I et al. Cyclin A is redundant in fibroblasts but essential in hematopoietic and embryonic stem cells. Cell 2009;138(2):352–365.

24. el Marjou F et al. Tissue-specific and inducible Cre-mediated recombination in the gut epithelium. Genes. N. Y. N 2000 2004;39(3):186–193.

25. Minoo P, Zlobec I, Peterson M, Terracciano L, Lugli A. Characterization of rectal, proximal and distal colon cancers based on clinicopathological, molecular and protein profiles. Int. J. Oncol. 2010;37(3):707–718.

26. Taniguchi K, Karin M. IL-6 and related cytokines as the critical lynchpins between inflammation and cancer. Semin. Immunol. 2014;26(1):54–74.

27. Laubitz D et al. Colonic gene expression profile in NHE3-deficient mice: evidence for spontaneous distal colitis. Am. J. Physiol. Gastrointest. Liver Physiol. 2008;295(1):G63–G77.

28. Sobecki M et al. Cell-Cycle Regulation Accounts for Variability in Ki-67 Expression Levels. Cancer Res. 2017;77(10):2722–2734.

29. Hayashi MT, Karlseder J. DNA damage associated with mitosis and cytokinesis failure. Oncogene 2013;32(39):4593–4601.

30. van den Bosch M, Bree RT, Lowndes NF. The MRN complex: coordinating and mediating the response to broken chromosomes. EMBO Rep. 2003;4(9):844–849.

31. Rein K, Stracker TH. The MRE11 complex: an important source of stress relief. Exp. Cell Res. 2014;329(1):162–169.

32. Karin M, Clevers H. Reparative inflammation takes charge of tissue regeneration. Nature 2016;529(7586):307–315.

33. Yui S et al. YAP/TAZ-Dependent Reprogramming of Colonic Epithelium Links ECM Remodeling to Tissue Regeneration. Cell Stem Cell 2018;22(1):35–49.e7.

34. De Robertis M et al. The AOM/DSS murine model for the study of colon carcinogenesis: From pathways to diagnosis and therapy studies. J. Carcinog. 2011;10:9.

35. Erben U et al. A guide to histomorphological evaluation of intestinal inflammation in mouse models. Int. J. Clin. Exp. Pathol. 2014;7(8):4557–4576.

36. Hydbring P, Malumbres M, Sicinski P. Non-canonical functions of cell cycle cyclins and cyclin-dependent kinases. Nat. Rev. Mol. Cell Biol. 2016;17(5):280–292.

37. Chen W, Lee J, Cho SY, Fine HA. Proteasome-mediated destruction of the cyclin a/cyclin-dependent kinase 2 complex suppresses tumor cell growth in vitro and in vivo. Cancer Res. 2004;64(11):3949–3957.

38. Clevers H, Loh KM, Nusse R. Stem cell signaling. An integral program for tissue renewal and regeneration: Wnt signaling and stem cell control. Science 2014;346(6205):1248012.

39. Kim SI et al. Cyclin-dependent kinase 2 regulates the interaction of Axin with beta-catenin. Biochem. Biophys. Res. Commun. 2004;317(2):478–483.

40. Kanakkanthara A et al. Cyclin A2 is an RNA binding protein that controls Mre11 mRNA translation. Science 2016;353(6307):1549–1552.

41. Petersen BO, Lukas J, Sørensen CS, Bartek J, Helin K. Phosphorylation of mammalian CDC6 by cyclin A/CDK2 regulates its subcellular localization. EMBO J. 1999;18(2):396–410.

42. Kabeche L, Compton DA. Cyclin A regulates kinetochore microtubules to promote faithful chromosome segregation. Nature 2013;502(7469):110–113.

43. Ganem NJ, Pellman D. Linking abnormal mitosis to the acquisition of DNA damage. J. Cell Biol. 2012;199(6):871–881.

44. Lee Y-C et al. High expression of phospho-H2AX predicts a poor prognosis in colorectal cancer. Anticancer Res. 2015;35(4):2447–2453.

45. Li J-A, Liu B-C, Song Y, Chen X. Cyclin A2 regulates symmetrical mitotic spindle formation and centrosome amplification in human colon cancer cells. Am. J. Transl. Res. 2018;10(8):2669–2676.

46. Georgakilas AG, Kotsinas A. Editorial: DNA Damage and Inflammation under Stress. Front. Genet. 2017;8:152.

47. Zanconato F, Cordenonsi M, Piccolo S. YAP/TAZ at the Roots of Cancer. Cancer Cell 2016;29(6):783–803.

48. Yuen H-F et al. TAZ expression as a prognostic indicator in colorectal cancer. PloS One 2013;8(1):e54211.

49. Handa K, Yamakawa M, Takeda H, Kimura S, Takahashi T. Expression of cell cycle markers in colorectal carcinoma: superiority of cyclin A as an indicator of poor prognosis. Int. J. Cancer 1999;84(3):225–233.

50. Bahnassy AA et al. Cyclin A and cyclin D1 as significant prognostic markers in colorectal cancer patients. BMC Gastroenterol. 2004;4:22.

51. Nozoe T, Inutsuka S, Honda M, Ezaki T, Korenaga D. Clinicopathologic significance of cyclin A expression in colorectal carcinoma. J. Exp. Clin. Cancer Res. CR 2004;23(1):127–133.

52. Li J-Q et al. Cyclin A correlates with carcinogenesis and metastasis, and p27(kip1) correlates with lymphatic invasion, in colorectal neoplasms. Hum. Pathol. 2002;33(10):1006–1015.

53. Bray NL, Pimentel H, Melsted P, Pachter L. Near-optimal probabilistic RNA-seq quantification. Nat. Biotechnol. 2016;34(5):525–527.

54. Dobin A et al. STAR: ultrafast universal RNA-seq aligner. Bioinforma. Oxf. Engl. 2013;29(1):15–21.

55. Irizarry RA et al. Exploration, normalization, and summaries of high density oligonucleotide array probe level data. Biostat. Oxf. Engl. 2003;4(2):249–264.

56. Bertucci F, Finetti P, Viens P, Birnbaum D. EndoPredict predicts for the response to neoadjuvant chemotherapy in ER-positive, HER2-negative breast cancer. Cancer Lett. 2014;355(1):70–75.

57. Eide PW, Bruun J, Lothe RA, Sveen A. CMScaller: an R package for consensus molecular subtyping of colorectal cancer pre-clinical models. Sci. Rep. 2017;7(1):16618.

58. Hochberg Y, Benjamini Y. More powerful procedures for multiple significance testing. Stat. Med. 1990;9(7):811–818.

59. McShane LM et al. REporting recommendations for tumor MARKer prognostic studies (REMARK). Nat. Clin. Pract. Urol. 2005;2(8):416–422.

