## Supplementary material for "Loss of cyclin A2 in murine colonic epithelial cells disrupts colon homeostasis by triggering DNA damage and dysplasia and high cyclin A2 expression is a good-prognosis factor in patients with colorectal cancer": YGuo et al Supplemental Figures and tables

### Supplemental Figure 1

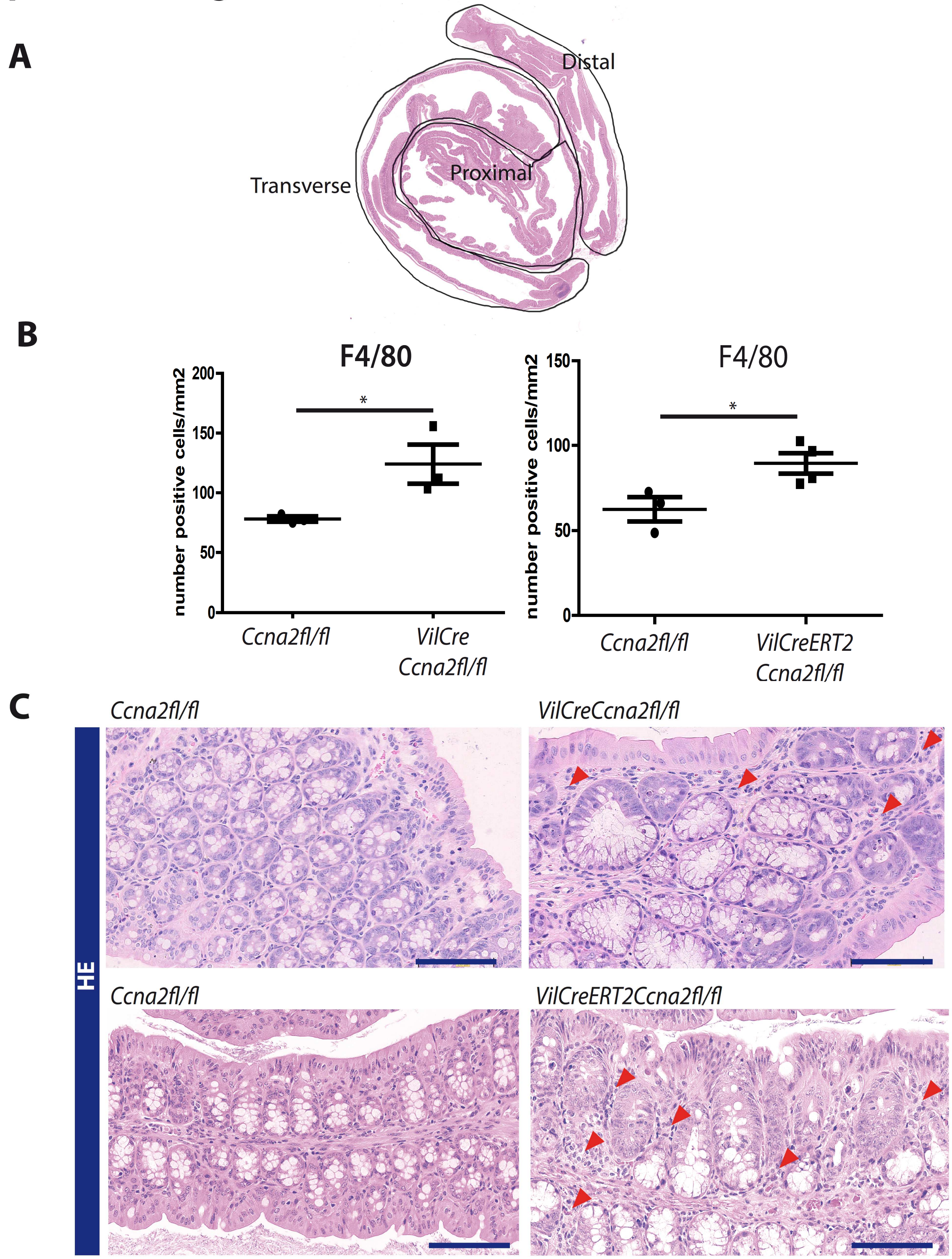


**Supplemental Figure 1**: **Cyclin A2-deficiency in colonic epithelial cells induces architectural changes in the mucosa and inflammation. (A)**. HE staining illustrating the subdivision of the colon in 3 parts for the histological analysis, *i.e.* the distal part close to the rectum, transverse and proximal part, close to the caecum. **(B).** Quantification of the F4/80 staining in control (n=3), constitutive (n=3) and inducible cyclin A2-deficient mice (n=4; expressed as mean±SEM, * $p<0.05$; two-tailed unpaired t-test) expressed as number of F4/80 positive cells per $mm^2$ of colon tissue. **(C)**. Representative HE staining of irregular formed crypts from a constitutive and inducible (day8 following tamoxifen injection) knockout mouse by comparison to a control. Please note the irregular formed crypts and immune cell infiltration (indicated by arrowheads). Scale bar: 100µm.

### Supplemental Figure 2

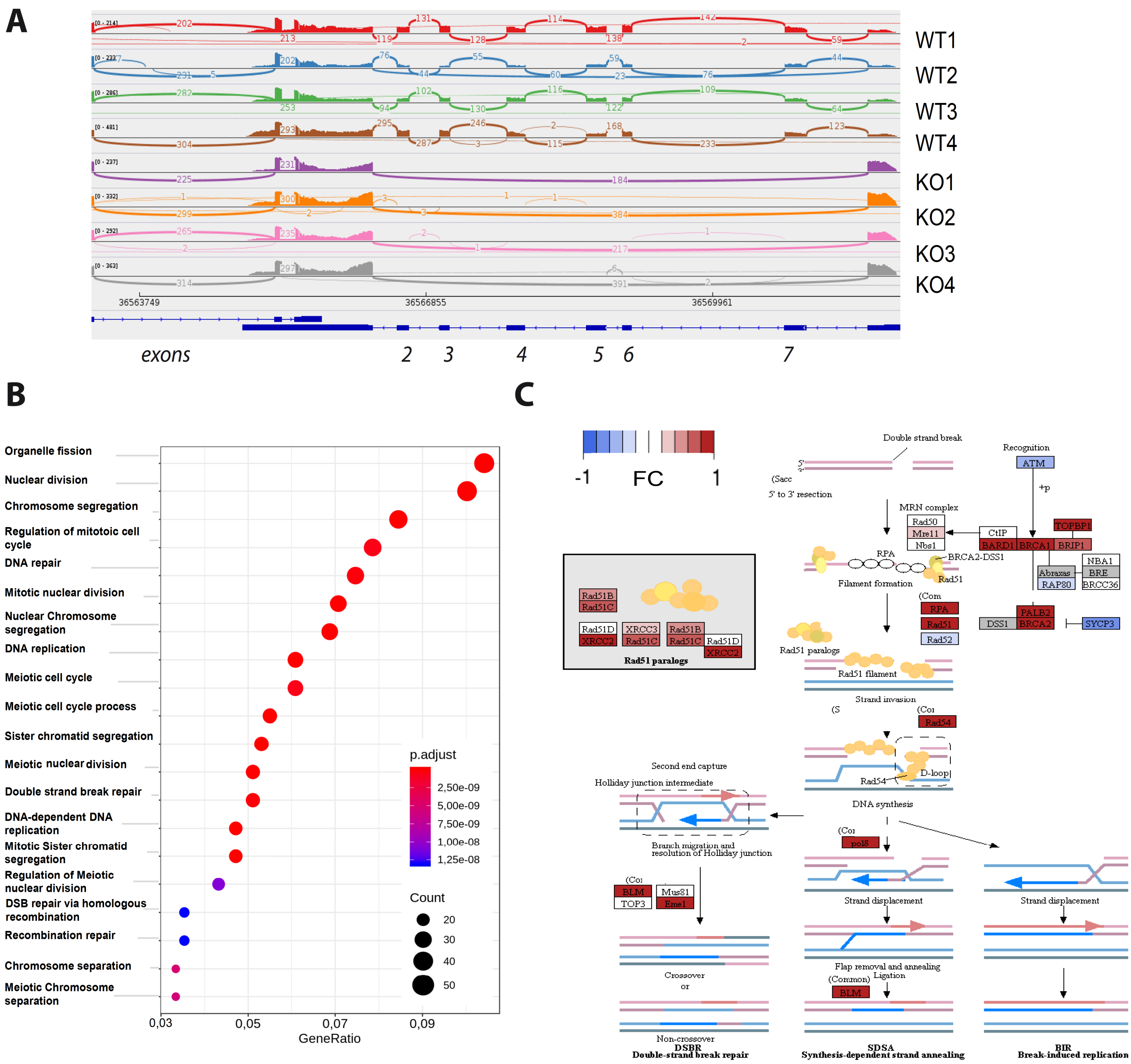


**Supplemental Figure 2: RNA-seq analysis of altered expression of genes involved in cell cycle regulation and DNA double-strand break repair in cyclin A2 deficient colonic epithelial cells (CEC). (A).** Sashimi plot of Ccna2 reads for each mouse-derived CEC samples analyzed by RNA-seq. Plots show deletion of exons 2 to 7 in transcripts of CEC from *VilCreCcna2fl/fl* mice compared to controls. The coverage for each alignment track is plotted as a bar graph. Arcs representing splice junctions connect exons. Arcs display the number of reads split across the junction (junction depth). Genomic coordinates and the gene annotation track are shown below the junction tracks. **(B)**. Gene Ontology over-representation analysis showing the genes up-regulated in *Ccna2 fl/fl* mutant mice compared to controls. The top 20 of Gene Ontology terms (Biological Process) are shown here. The p-values have been corrected for multiple testing by the Benjamini-Hochberg method. In this dot plot, the color represents the adjusted p-values and the dot size represents the number of genes for each term. The gene ratio is shown on the x-axis. **(C).** Alterations in the double-strand break KEGG pathway in cyclin A2 deficient CEC. Genes up-regulated in *VilCreCcna2fl/fl* mice samples are in red, whereas downregulated genes are in blue. The scale indicated on the figure represent the log2 Fold change. In grey are the genes absent from our data. For all analyses, only p-values <0.05 were considered as statistically significant. FC: Fold change; DSB: Double-strand break

### Supplemental Figure 3

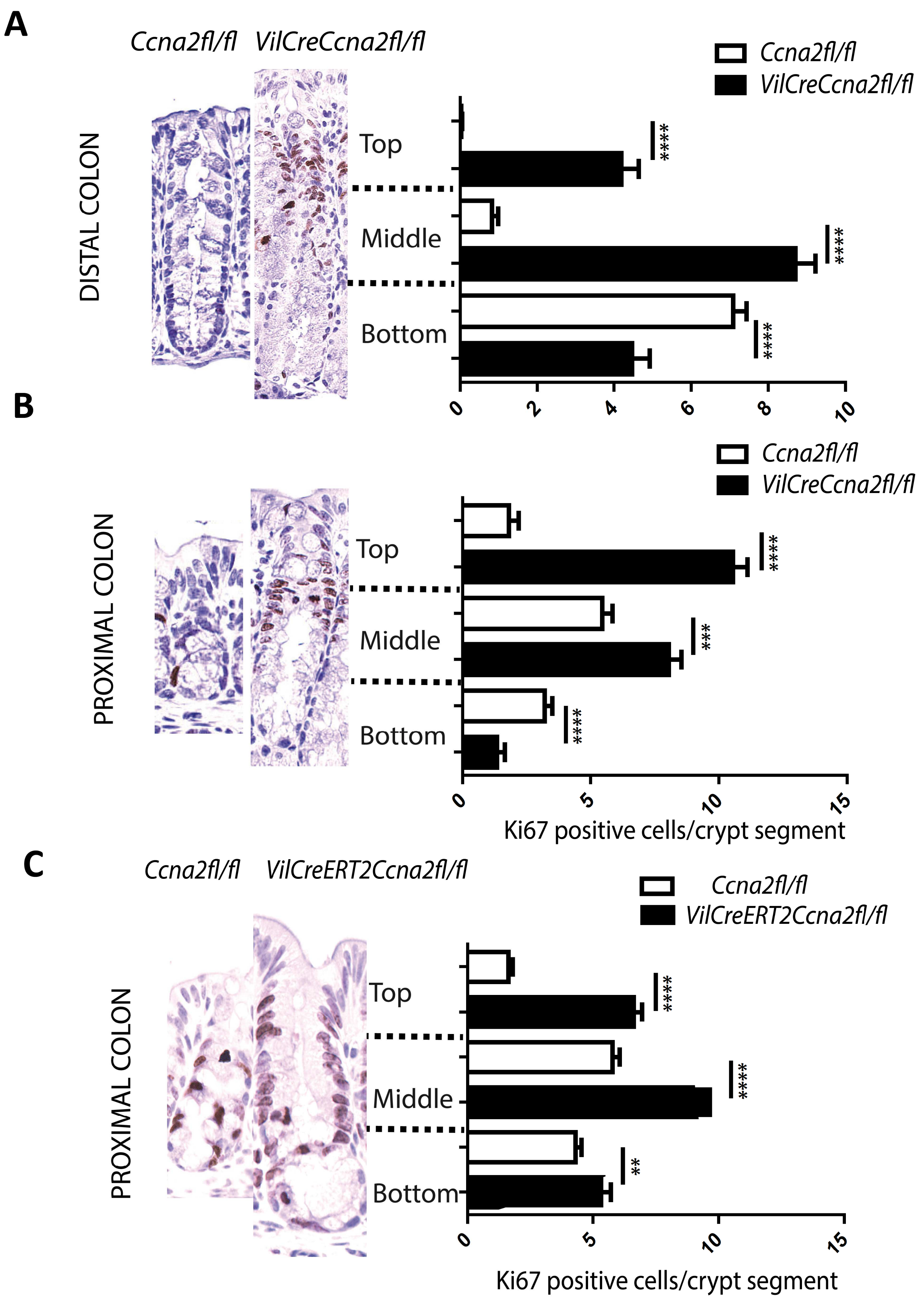


**Supplemental Figure 3**: **Increased proliferation of colonic epithelial cells from cyclin A2-deficient mice**. **(A, B, C)**. Quantification of Ki67 expression by IHC of the different parts of crypts from the distal and proximal part of the colon from control (n= 66 crypts analyzed from 3 different mice) and constitutive cyclin A2-deficient mice (**A, B**) and proximal part of the colon from induced knockout mice at day 8 following inactivation **(C)** compared to controls (n=120 from 3 different mice). Mean values±SEM are provided, * p<0,05, ** p<0.01 and *** p<0,001; two-tailed unpaired t-test.

### Supplemental Figure 4

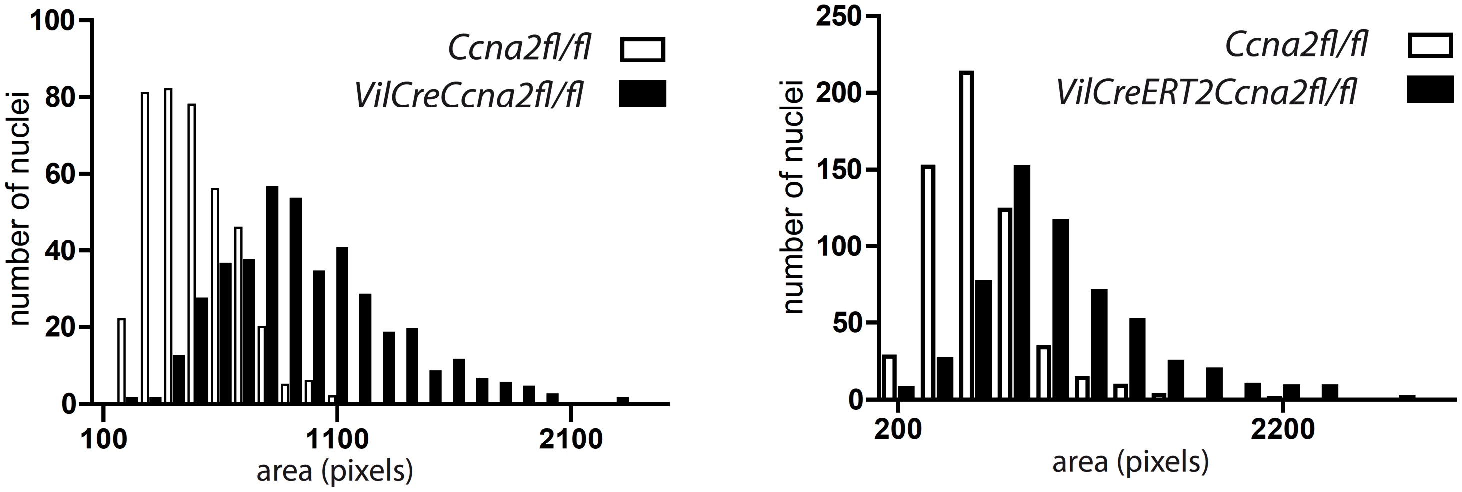


**Supplemental Figure 4**: **Increased nuclear size of colonic epithelial cells in cyclin A2-deficient mice**. Distribution curve representing the nuclear size of *Ccna2fl/fl* colonic epithelial cells (white, n=398 for the left panel and 578 for the right panel from 3 mice), *VilCreCcna2fl/fl* (black, left panel, n=400 from 3 mice, $p<1.10e-16$) and *VilCreERT2Ccna2fl/fl* (black, right panel, n=566, $p<1.10e-16$) colonic epithelial cells; p-values were determined using Kolmogorov-Smirnov test.

### Supplemental Figure 5

**A**

*Ccna2fl/fl* *VilCreERT2Ccna2fl/fl*

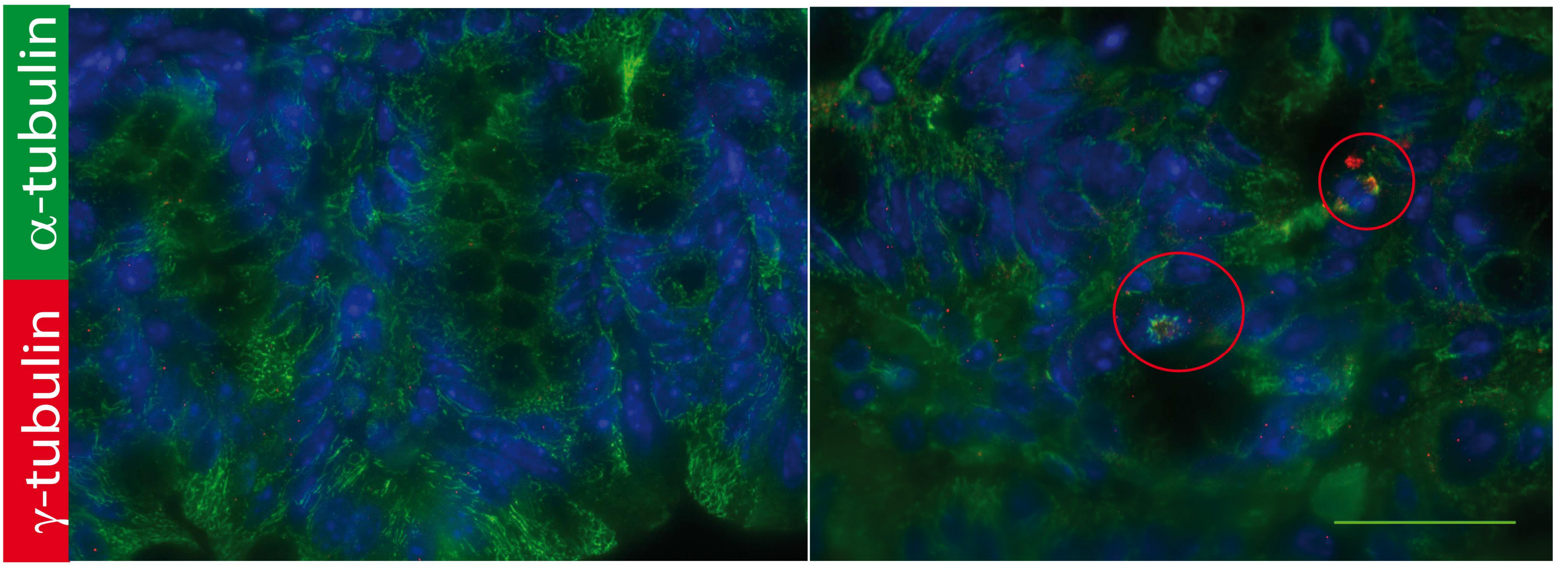


**B**

*VilCreERT2Ccna2fl/fl*

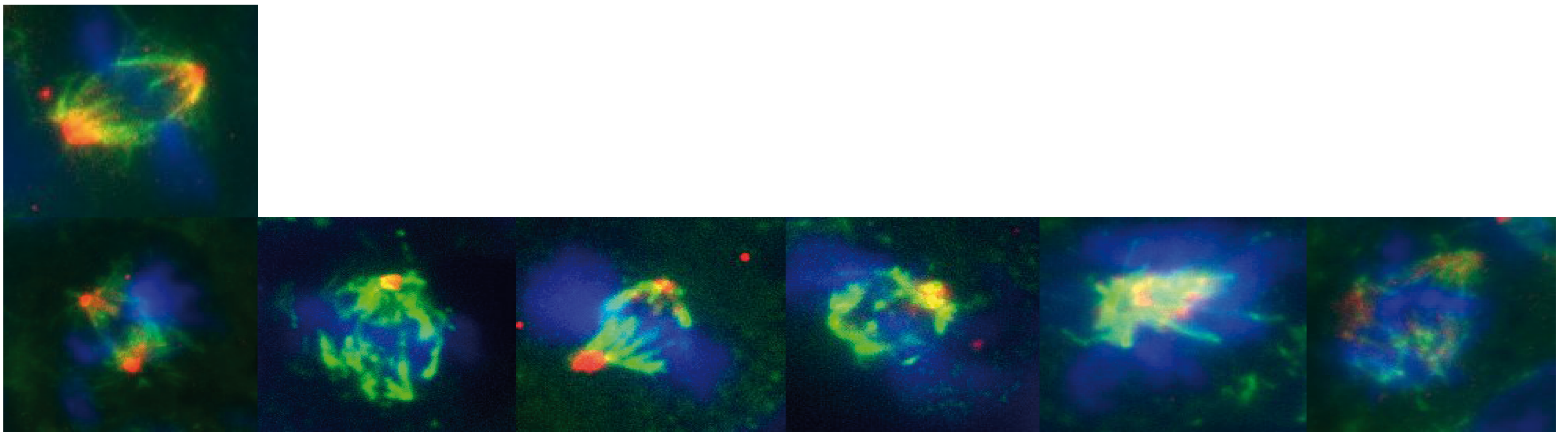

**Supplemental Figure 5**: **Examples of mitoses in colons of cyclin A2-deficient mice**. (**A**). Immunofluorescence analysis of mitosis (indicated by red circles) using an anti γ-tubulin antibody (green) to stain for the mitotic spindle in combination with α-tubulin (centrosome in red) and DAPI for DNA in a colon of *VilCreERT2Ccna2fl/fl* mice at day 8 following inactivation (right panel) by comparison to controls (left panel). Scale bar: 100µm. **(B)**. The upper image shows normal mitosis, the lower panel several examples of abnormal mitoses observed in a *Ccna2*-deficient colon. Blow Up: 2.5X.

#### Supplemental Figure 6

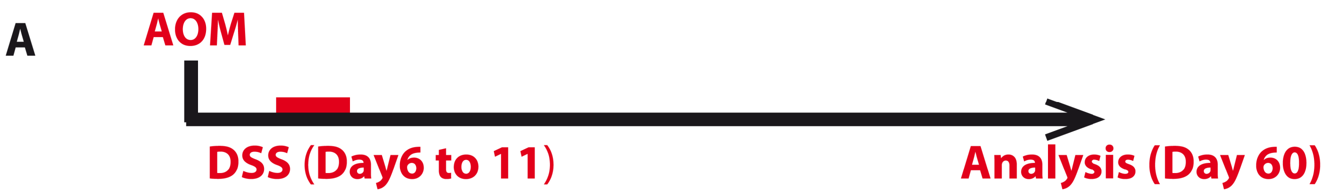


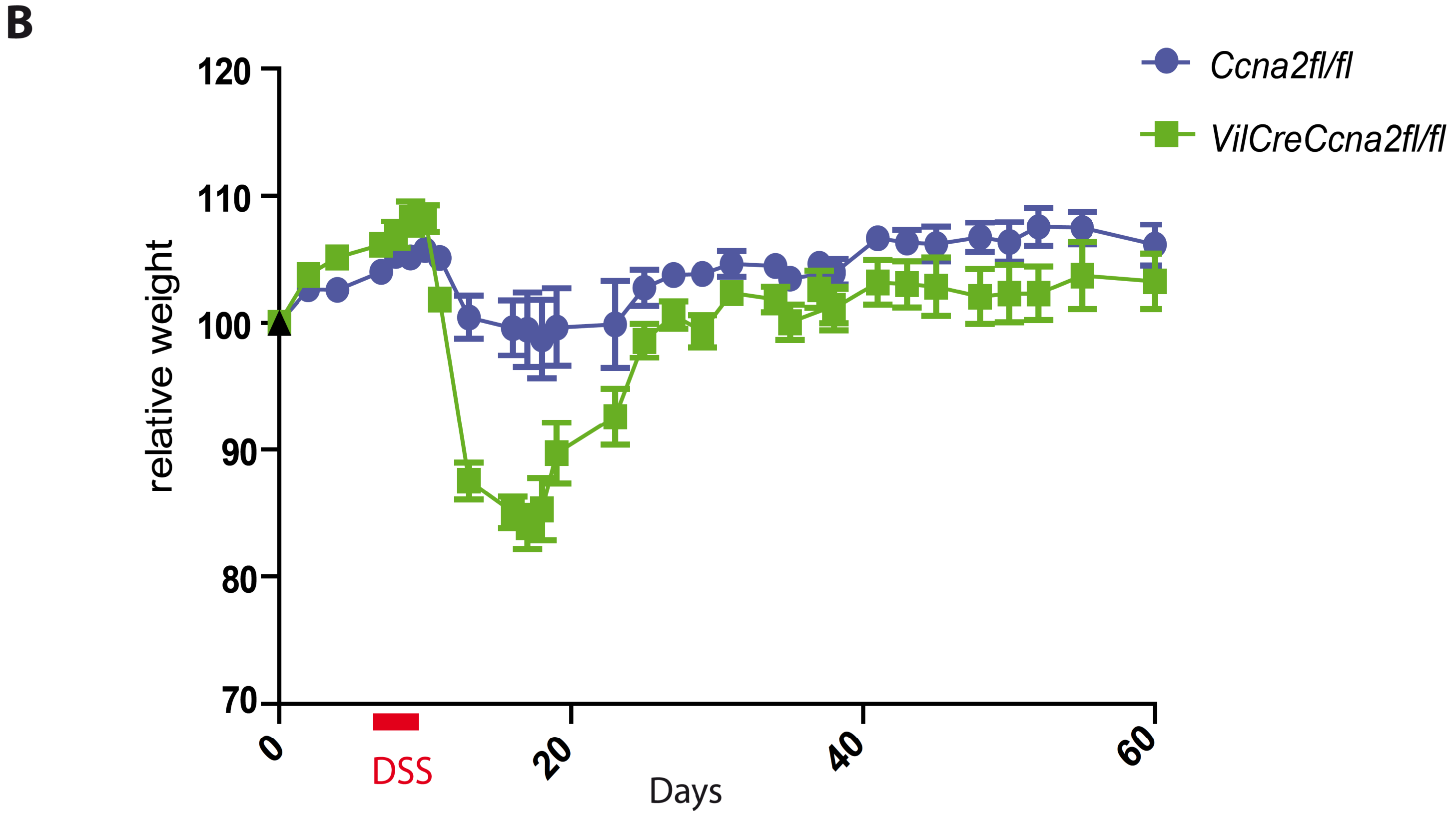


**Supplemental Figure 6**: **Protocol and weight monitoring of the mice during the colitis associated carcinogenesis**. **(A)**. Schematic representation of the modified AOM/DSS protocol applied to *VilCreCcna2fl/fl* (n=7) and control mice (n=6) (see Figs 6 and 7). **(B)**. Monitoring of the relative weight (expressed as percentage relative to the weight at the beginning of the protocol) of *VilCreCcna2fl/fl* and control mice during the AOM/DSS protocol.

### Supplemental Figure 7

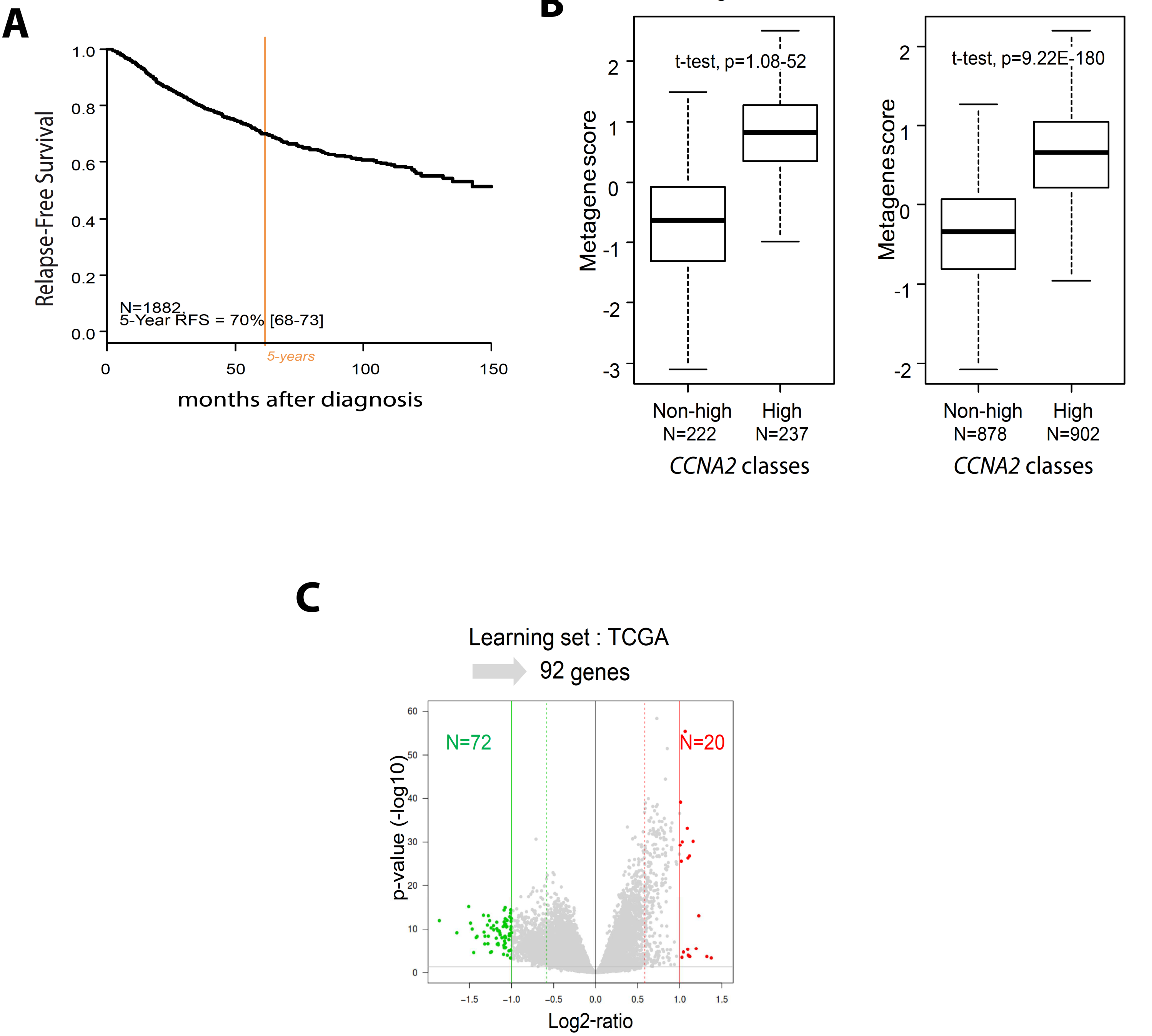


**Supplemental Figure 7: Cyclin A2 expression at the mRNA and protein level in CRC patients**. **(A)**. Relapse free survival (RFS) curve of the overall patients analyzed for cyclin A2 mRNA expression levels. **(B)**. Metagene-based prediction score of outcome (using Student t-test and expressed as mean ±SD) of the *CCNA2*high samples compared to those of *CCNA2*non-high samples in the learning set (left) and in the independent validation set (right). **(C)**. Volcano plot showing the 92 genes differentially expressed in the learning set (TCGA). Genes up-regulated in the *CCNA2*high samples are colored in red and genes down-regulated in green.

### Supplemental Figure 8

A

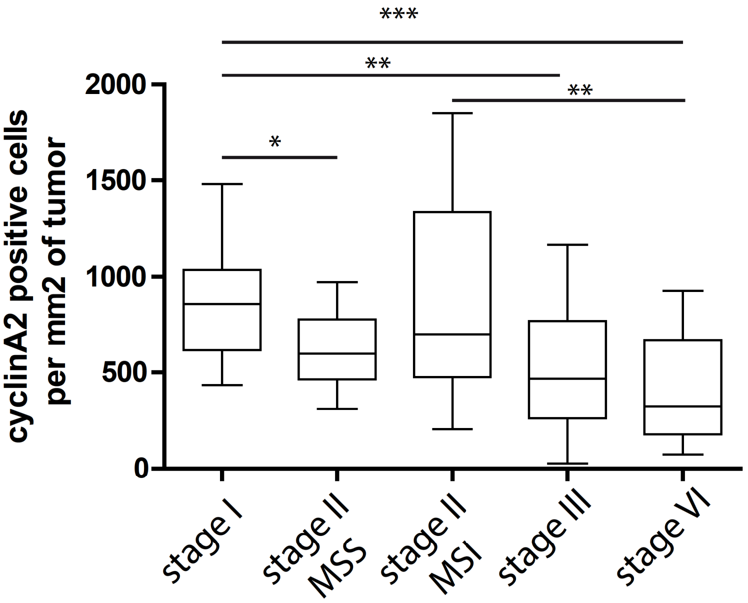


B

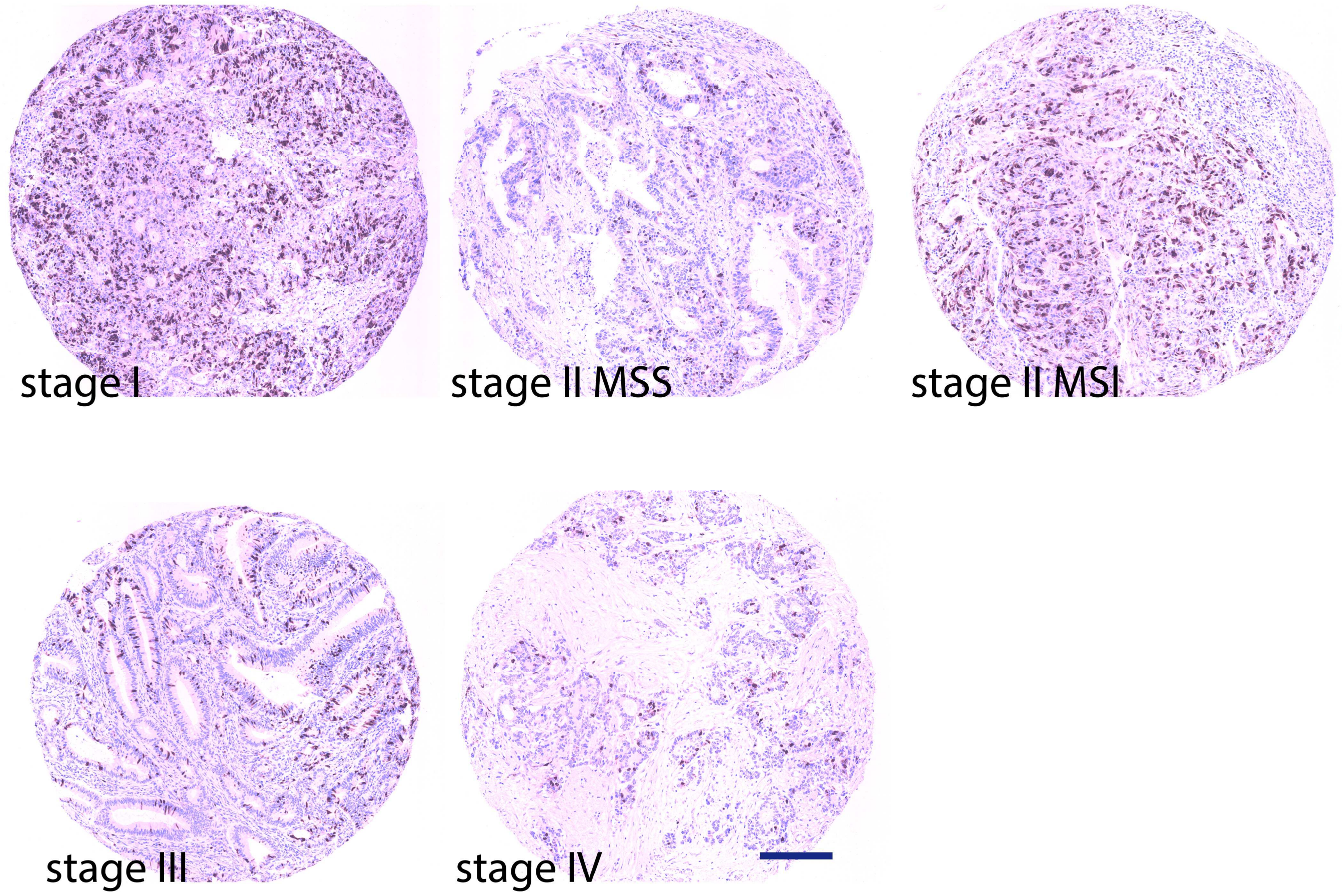


**Supplemental Figure 8**: **Cyclin A2 protein expression in CRC tumor samples from different stages. (A)**. Cyclin A2 expression analyzed on the same TMA shown in Figure 8, but using a different anti-cyclin A2 antibody from Novocastra. (Mean values ±SEM, $p<0.05$ for the analysis between stage I and II-MSS, $p<0.01$ for comparison between stage I and stage III, $p<0.001$ for stage I to IV, $p<0.01$ for stage II-MSI to stage IV, two-tailed unpaired t-test). **(B)**. Representative immunostaining of stage I, II-MSS, II-MSI, III and IV tumor samples. Scale bar: 100µm.

**Supplemental Table 1**

| Target | Primer | Sequence (5'-3') |
|---|---|---|
| ***Ccna**$^{flox\ allele}$* | 5' | CGCAGCAGAAGCTCAAGACTCGAC |
| | 3' | TCTACATCCTAATGCAATGCCTGG |
| ***Ccna*** Δ | 5' | CGCAGCAGAAGCTCAAGACTCGAC |
| | 3' | CACTCACACACTTAGTGTCTCTGG |
| ***Cre*** | 5' | CAAGCCTGGCTCGACGGCC |
| | 3' | CGCGAACATCTTCAGGTTCT |

**Supplemental Table 2.** Histological scoring for colitis (sum of scores 1 and 2) according to (40).

| **Severity** | **Extent** | **Score 1** | **Epithelial changes** | **Mucosal architecture** | **Score 2** |
|---|---|---|---|---|---|
| Mild | Mucosa | 1 | Focal erosions | | 1 |
| Moderate | Mucosa and submucosa | 2 | Erosions | focal ulcerations | 2 |
| Marked | Transmural | 3 | | Extended ulcerations± granulation tissue± pseudopolyps | 3 |

**Supplemental Table 3.** Patients and tumor characteristics of the TMA cohort (n = 65).

| Variable | Value (%) |
|---|---|
| Age at diagnosis (yr) | |
| Median | 66.26 |
| Range | 21-89 |
| Gender | |
| Male | 34 (52.3) |
| Female | 31 (47.7) |
| Tumor Location | |
| Colon | 65 (100) |
| Tumor Stage | |
| I | 13(20)n=23(tissue samples |
| II | 23 (35.4) n=48 |
| III | 15 (23.1)/5 with metachronous liver metastase, n=30 |
| IV | 14 (21.5)/10 with asynchronous liver metastase, n=26 |

**Supplementary Table 4: Ontology analysis of the 1109 genes differentially expressed between the *VilCreCcna2fl/fl* mice (n = 4) and *Ccna2fl/fl* mice samples (n = 4)**

| | | up Vil-Cre-Ccna2 fl/fl mice | | | | | down Vil-Cre-Ccna2 fl/fl mice | | | | |
|---|---|---|---|---|---|---|---|---|---|---|---|
| **ID** | **Description** | **N** | **p-value** | **p-adjusted** | **q-value** | **gene ID** | **N** | **p-value** | **p-adjusted** | **q-value** | **gene ID** |
| GO:0000280 | nuclear division | 51 | 1,53E-23 | 6,01E-20 | 5,31E-20 | Hfm1, Rec8, Stil, Cit, Mybl1, Mki67, Nsl1, Smc2, Brca2, Fanca, Ccne1, Mybl2, Map9, Top2a, Fancd2, Ndc1, Trip13, Fignl1, Rad51, Bmp4, Ccnf, Blm, Espl1, Gen1, Ereg, Dsn1, Kifc5b, Nuf2, Kif11, Ccne2, Ncapg, Incenp, Cdc6, Bub1, Suv39h2, Mastl, Cdca5, Ncapd2, Chtf18, Mad2l1, Kif4, Fbxo5, Ncaph, Eme1, Nek2, Bub1b, Ndc80, Pbp2, Kif18b, Spag5, Pdxp | | | | | |
| GO:0048285 | organelle fission | 53 | 5,04E-23 | 9,88E-20 | 8,73E-20 | Hfm1, Rec8, Stil, Cit, Mybl1, Mki67, Nsl1, Smc2, Brca2, Fanca, Ccne1, Mybl2, Map9, Top2a, Fancd2, Ndc1, Trip13, Fignl1, Bnip3, Rad51, Bmp4, Ccnf, Blm, Espl1, Gen1, Ereg, Dsn1, Kifc5b, Nuf2, Kif11, Ccne2, Ncapg, Incenp, Cdc6, Bub1, Suv39h2, Mastl, Mtfr2, Cdca5, Ncapd2, Chtf18, Mad2l1, Kif4, Fbxo5, Ncaph, Eme1, Nek2, Bub1b, Ndc80, Pbp2, Kif18b, Spag5, Pdxp | | | | | |
| GO:0007059 | chromosome segregation | 43 | 4,29E-22 | 5,61E-19 | 4,95E-19 | Hfm1, Rec8, Stil, Cit, Mki67, Brca1, Nsl1, Smc2, Brca2, Ccne1, Top2a, Fancd2, Ndc1, Rad18, Trip13, Esco2, Blm, Espl1, Gen1, Dsn1, Nuf2, Ccne2, Ncapg, Incenp, Cdc6, Bub1, Ska1, Spc25, Cdca2, Cdca5, Ncapd2, Chtf18, Mad2l1, Kif4, Fbxo5, Oip5, Ncaph, Eme1, Nek2, Bub1b, Ndc80, Kif18b, Spag5 | | | | | |
| GO:0140014 | mitotic nuclear division | 36 | 4,80E-18 | 3,88E-15 | 3,43E-15 | Stil, Cit, Mki67, Nsl1, Smc2, Ccne1, Mybl2, Map9, Trip13, Bmp4, Ccnf, Espl1, Gen1, Ereg, Dsn1, Kifc5b, Nuf2, Kif11, Ccne2, Ncapg, Incenp, Cdc6, Bub1, Cdca5, Ncapd2, Mad2l1, Kif4, Fbxo5, Ncaph, Nek2, Bub1b, Ndc80, Pbp2, Kif18b, Spag5, Pdxp | | | | | |
| GO:0098813 | nuclear chromosome segregation | 35 | 4,95E-18 | 3,88E-15 | 3,43E-15 | Hfm1, Rec8, Stil, Cit, Nsl1, Smc2, Brca2, Ccne1, Top2a, Fancd2, Ndc1, Trip13, Esco2, Blm, Espl1, Gen1, Dsn1, Nuf2, Ccne2, Ncapg, Incenp, Cdc6, Bub1, Cdca5, Ncapd2, Mad2l1, Kif4, Fbxo5, Ncaph, Eme1, Nek2, Bub1b, Ndc80, Kif18b, Spag5 | | | | | |
| GO:0006261 | DNA-dependent DNA replication | 24 | 2,25E-15 | 1,45E-12 | 1,28E-12 | Cdc7, E2f7, Polq, Ticrr, Brca2, Ccne1, E2f8, Rad51, Blm, Wdhd1, Mms22l, Gen1, Mcm10, Dna2, Ccne2, Cdc6, Mcm8, Zfp365, Chtf18, Pole, Fbxo5, Eme1, Cdc45, Pola1 | | | | | |
| GO:1903046 | meiotic cell cycle process | 28 | 2,75E-15 | 1,45E-12 | 1,28E-12 | Hfm1, Rec8, Mybl1, Smc2, Brca2, Fanca, Ccne1, Top2a, Fancd2, Ndc1, Trip13, Fignl1, Rad51, Blm, Espl1, Ereg, Nuf2, Ccne2, Bub1, Suv39h2, Mastl, Ncapd2, Chtf18, Fbxo5, Ncaph, Eme1, Exo1, Bub1b | | | | | |
| GO:0006260 | DNA replication | 31 | 2,95E-15 | 1,45E-12 | 1,28E-12 | Cdc7, Ccdc88a, E2f7, Polq, Rrm2, Ticrr, Brca2, Ccne1, Fam111a, Esco2, E2f8, Rad51, Blm, Wdhd1, Mms22l, Gen1, Ereg, Mcm10, Rrm1, Dna2, Chaf1a, Ccne2, Cdc6, Mcm8, Zfp365, Chtf18, Pole, Fbxo5, Eme1, Cdc45, Pola1 | | | | | |
| GO:0000819 | sister chromatid segregation | 27 | 3,76E-15 | 1,64E-12 | 1,44E-12 | Rec8, Stil, Cit, Nsl1, Smc2, Top2a, Trip13, Esco2, Espl1, Gen1, Dsn1, Nuf2, Ncapg, Incenp, Cdc6, Bub1, Cdca5, Ncapd2, Mad2l1, Kif4, Fbxo5, Ncaph, Nek2, Bub1b, Ndc80, Kif18b, Spag5 | | | | | |
| GO:0140013 | meiotic nuclear division | 26 | 1,17E-14 | 4,60E-12 | 4,06E-12 | Hfm1, Rec8, Mybl1, Smc2, Brca2, Fanca, Ccne1, Top2a, Fancd2, Ndc1, Trip13, Fignl1, Rad51, Blm, Espl1, Ereg, Nuf2, Ccne2, Bub1, Suv39h2, Mastl, Chtf18, Fbxo5, Ncaph, Eme1, Bub1b | | | | | |
| GO:0000070 | mitotic sister chromatid segregation | 24 | 1,92E-14 | 6,84E-12 | 6,04E-12 | Stil, Cit, Nsl1, Smc2, Trip13, Espl1, Gen1, Dsn1, Nuf2, Ncapg, Incenp, Cdc6, Bub1, Cdca5, Ncapd2, Mad2l1, Kif4, Fbxo5, Ncaph, Nek2, Bub1b, Ndc80, Kif18b, Spag5 | | | | | |
| GO:0006302 | double-strand break repair | 26 | 5,41E-13 | 1,77E-10 | 1,56E-10 | Cdc7, Rec8, Polq, Rad54b, Brca1, Brca2, Trip13, Fignl1, Foxm1, Esco2, Rad51, Blm, Mms22l, Rad54l, Gen1, Dna2, Mcm8, Cdca5, Zfp365, Xrcc2, Eme1, Exo1, Rad51ap1, Cdc45, Pola1, Fancb | | | | | |
| GO:0006281 | DNA repair | 38 | 8,11E-13 | 2,44E-10 | 2,16E-10 | Cdc7, Rec8, Polq, Rad54b, Brca1, Smc2, Ticrr, Brca2, Fanca, Fancd2, Rad18, Trip13, Fignl1, Foxm1, Esco2, Rad51, Clspn, Blm, Wdhd1, Mms22l, Rad54l, Bard1, Gen1, Dna2, Chaf1a, Neil3, Mcm8, Cdca5, Zfp365, Xrcc2, Pole, Fanci, Eme1, Exo1, Rad51ap1, Cdc45, Pola1, Fancb | | | | | |
| GO:0007346 | regulation of mitotic cell cycle | 40 | 1,86E-12 | 5,20E-10 | 4,60E-10 | Cdc7, Stil, E2f7, Cit, Ddias, Mki67, Brca1, Ticrr, Zwilch, Brca2, Cdkn2a, Ccne1, Map9, Top2a, Trip13, Clspn, Bmp4, Ccnf, Blm, Trim35, Gen1, Ereg, Dact1, Magi2, Kif11, Ccne2, Incenp, Cdc6, Bub1, Cdca5, Kntc1, Mad2l1, Fbxo5, Eme1, Nek2, Cdc45, Bub1b, Ndc80, Pbp2, Pdxp | | | | | |
| GO:0051321 | meiotic cell cycle | 31 | 2,07E-12 | 5,42E-10 | 4,79E-10 | Hfm1, Rec8, Mybl1, Mki67, Smc2, Brca2, Fanca, Ccne1, Top2a, Fancd2, Ndc1, Trip13, Fignl1, Mns1, Rad51, Blm, Espl1, Ereg, Nuf2, Ccne2, Bub1, Suv39h2, Mastl, Ncapd2, Chtf18, Fbxo5, Ncaph, Eme1, Exo1, Nek2, Bub1b | | | | | |
| GO:0051304 | chromosome separation | 17 | 2,10E-11 | 5,14E-09 | 4,54E-09 | Hfm1, Stil, Cit, Top2a, Trip13, Blm, Espl1, Gen1, Cdc6, Bub1, Ncapd2, Mad2l1, Fbxo5, Ncaph, Eme1, Bub1b, Ndc80 | | | | | |
| GO:0045132 | meiotic chromosome segregation | 17 | 2,50E-11 | 5,77E-09 | 5,10E-09 | Hfm1, Rec8, Smc2, Brca2, Ccne1, Top2a, Fancd2, Ndc1, Trip13, Blm, Espl1, Nuf2, Ccne2, Bub1, Ncaph, Eme1, Bub1b | | | | | |
| GO:0007088 | regulation of mitotic nuclear division | 22 | 4,96E-11 | 1,08E-08 | 9,54E-09 | Stil, Cit, Mki67, Ccne1, Map9, Trip13, Bmp4, Ccnf, Gen1, Ereg, Kif11, Ccne2, Cdc6, Bub1, Cdca5, Mad2l1, Fbxo5, Nek2, Bub1b, Ndc80, Pbp2, Pdxp | | | | | |
| GO:0000724 | double-strand break repair via homologous recombination | 18 | 6,68E-11 | 1,31E-08 | 1,16E-08 | Cdc7, Rec8, Polq, Rad54b, Brca1, Brca2, Fignl1, Rad51, Blm, Mms22l, Rad54l, Gen1, Mcm8, Zfp365, Xrcc2, Rad51ap1, Cdc45, Fancb | | | | | |

| | | | | | | |
|---|---|---|---|---|---|---|
| GO:0000725 | recombinational repair | 18 | 6,68E-11 | 1,31E-08 | 1,16E-08 | Cdc7, Rec8, Polq, Rad54b, Brca1, Brca2, Fignl1, Rad51, Blm, Mms22l, Rad54l, Gen1, Mcm8, Zfp365, Xrcc2, Rad51ap1, Cdc45, Fancb |
| GO:0006310 | DNA recombination | 26 | 2,10E-10 | 3,92E-08 | 3,46E-08 | Cdc7, Hfm1, Rec8, Polq, Rad54b, Brca1, Brca2, Top2a, Rad18, Trip13, Fignl1, Rad51, Blm, Mms22l, Rad54l, Gen1, Mcm8, Zfp365, Ankle1, Xrcc2, Chtf18, Eme1, Exo1, Rad51ap1, Cdc45, Fancb |
| GO:0051983 | regulation of chromosome segregation | 16 | 1,00E-09 | 1,78E-07 | 1,58E-07 | Stil, Cit, Mki67, Rad18, Trip13, Gen1, Cdc6, Bub1, Cdca5, Chtf18, Mad2l1, Fbxo5, Nek2, Bub1b, Ndc80, Spag5 |
| GO:0051783 | regulation of nuclear division | 22 | 1,16E-09 | 1,98E-07 | 1,75E-07 | Stil, Cit, Mki67, Ccne1, Map9, Trip13, Bmp4, Ccnf, Gen1, Ereg, Kif11, Ccne2, Cdc6, Bub1, Cdca5, Mad2l1, Fbxo5, Nek2, Bub1b, Ndc80, Pbp2, Pdxp |
| GO:0007127 | meiosis I | 16 | 1,77E-09 | 2,89E-07 | 2,56E-07 | Hfm1, Rec8, Mybl1, Brca2, Ccne1, Top2a, Fancd2, Ndc1, Trip13, Rad51, Blm, Espl1, Ccne2, Chtf18, Fbxo5, Eme1 |
| GO:0045930 | negative regulation of mitotic cell cycle | 23 | 2,08E-09 | 3,26E-07 | 2,88E-07 | Stil, E2f7, Ddias, Brca1, Ticrr, Zwilch, Top2a, Trip13, Clspn, Bmp4, Blm, Trim35, Gen1, Dact1, Magi2, Cdc6, Bub1, Kntc1, Mad2l1, Fbxo5, Eme1, Bub1b, Ndc80 |
| GO:0044772 | mitotic cell cycle phase transition | 28 | 3,03E-09 | 4,57E-07 | 4,04E-07 | Cdc7, Skp2, Stil, E2f7, Cit, Brca1, Ticrr, Cdkn2a, Ccne1, Trip13, Foxm1, Clspn, Blm, Gen1, Dact1, Ccne2, Cdc6, Bub1, Mastl, Cdca5, Mad2l1, Pole, Fbxo5, Iqgap3, Id4, Cdc45, Bub1b, Ndc80 |
| GO:0061982 | meiosis I cell cycle process | 16 | 3,49E-09 | 5,07E-07 | 4,47E-07 | Hfm1, Rec8, Mybl1, Brca2, Ccne1, Top2a, Fancd2, Ndc1, Trip13, Rad51, Blm, Espl1, Ccne2, Chtf18, Fbxo5, Eme1 |
| GO:0044770 | cell cycle phase transition | 29 | 7,23E-09 | 1,01E-06 | 8,94E-07 | Cdc7, Skp2, Stil, E2f7, Cit, Brca1, Ticrr, Cdkn2a, Ccne1, Dyrk3, Trip13, Foxm1, Clspn, Blm, Gen1, Dact1, Ccne2, Cdc6, Bub1, Mastl, Cdca5, Mad2l1, Pole, Fbxo5, Iqgap3, Id4, Cdc45, Bub1b, Ndc80 |
| GO:0006270 | DNA replication initiation | 9 | 1,42E-08 | 1,92E-06 | 1,69E-06 | Cdc7, Ticrr, Ccne1, Mcm10, Ccne2, Cdc6, Mcm8, Cdc45, Pola1 |
| GO:0045786 | negative regulation of cell cycle | 31 | 1,75E-08 | 2,28E-06 | 2,01E-06 | Stil, E2f7, Ddias, Brca1, Ticrr, Zwilch, Cdkn2a, Top2a, Trip13, Foxm1, E2f8, Clspn, Bmp4, Ccnf, Blm, Trim35, Mdm1, Gen1, Dact1, Magi2, Dna2, Cdc6, Bub1, Kntc1, Mad2l1, Fbxo5, Eme1, Nek2, Bub1b, Ndc80, Mlf1 |
| GO:0007093 | mitotic cell cycle checkpoint | 16 | 3,03E-08 | 3,72E-06 | 3,29E-06 | Stil, Brca1, Ticrr, Zwilch, Top2a, Trip13, Clspn, Blm, Gen1, Cdc6, Bub1, Kntc1, Mad2l1, Eme1, Bub1b, Ndc80 |
| GO:0070192 | chromosome organization involved in meiotic cell cycle | 12 | 3,04E-08 | 3,72E-06 | 3,29E-06 | Rec8, Smc2, Ccne1, Fancd2, Ndc1, Trip13, Rad51, Ccne2, Bub1, Ncapd2, Ncaph, Bub1b |
| GO:0010948 | negative regulation of cell cycle process | 20 | 1,08E-07 | 1,28E-05 | 1,13E-05 | Stil, E2f7, Brca1, Ticrr, Trip13, E2f8, Clspn, Bmp4, Ccnf, Blm, Mdm1, Gen1, Dact1, Cdc6, Bub1, Mad2l1, Fbxo5, Nek2, Bub1b, Ndc80 |
| GO:0045005 | DNA-dependent DNA replication maintenance of fidelity | 9 | 1,28E-07 | 1,47E-05 | 1,30E-05 | Brca2, Rad51, Blm, Mms22l, Gen1, Dna2, Zfp365, Pole, Eme1 |
| GO:0031297 | replication fork processing | 8 | 1,92E-07 | 2,15E-05 | 1,90E-05 | Brca2, Rad51, Blm, Mms22l, Gen1, Dna2, Zfp365, Eme1 |
| GO:0044786 | cell cycle DNA replication | 9 | 3,14E-07 | 3,42E-05 | 3,02E-05 | Cdc7, E2f7, Brca2, E2f8, Rad51, Dna2, Fbxo5, Cdc45, Pola1 |
| GO:0000075 | cell cycle checkpoint | 17 | 3,61E-07 | 3,82E-05 | 3,38E-05 | Stil, Brca1, Ticrr, Zwilch, Top2a, Trip13, Clspn, Blm, Gen1, Dna2, Cdc6, Bub1, Kntc1, Mad2l1, Eme1, Bub1b, Ndc80 |
| GO:0007091 | metaphase/anaphase transition of mitotic cell cycle | 10 | 3,99E-07 | 4,12E-05 | 3,64E-05 | Stil, Cit, Trip13, Gen1, Cdc6, Bub1, Mad2l1, Fbxo5, Bub1b, Ndc80 |
| GO:0010965 | regulation of mitotic sister chromatid separation | 10 | 4,73E-07 | 4,75E-05 | 4,20E-05 | Stil, Cit, Trip13, Gen1, Cdc6, Bub1, Mad2l1, Fbxo5, Bub1b, Ndc80 |
| GO:1901991 | negative regulation of mitotic cell cycle phase transition | 15 | 5,58E-07 | 5,47E-05 | 4,83E-05 | Stil, E2f7, Brca1, Ticrr, Trip13, Clspn, Blm, Gen1, Dact1, Cdc6, Bub1, Mad2l1, Fbxo5, Bub1b, Ndc80 |
| GO:0044784 | metaphase/anaphase transition of cell cycle | 10 | 6,57E-07 | 6,28E-05 | 5,55E-05 | Stil, Cit, Trip13, Gen1, Cdc6, Bub1, Mad2l1, Fbxo5, Bub1b, Ndc80 |
| GO:0006275 | regulation of DNA replication | 13 | 7,10E-07 | 6,62E-05 | 5,85E-05 | Cdc7, Ccdc88a, E2f7, Ticrr, Brca2, Esco2, E2f8, Blm, Ereg, Dna2, Zfp365, Chtf18, Fbxo5 |
| GO:0051306 | mitotic sister chromatid separation | 10 | 9,01E-07 | 8,21E-05 | 7,25E-05 | Stil, Cit, Trip13, Gen1, Cdc6, Bub1, Mad2l1, Fbxo5, Bub1b, Ndc80 |
| GO:0090329 | regulation of DNA-dependent DNA replication | 9 | 1,02E-06 | 9,08E-05 | 8,02E-05 | Cdc7, E2f7, Ticrr, Brca2, E2f8, Blm, Zfp365, Chtf18, Fbxo5 |
| GO:0051985 | negative regulation of chromosome segregation | 9 | 1,22E-06 | 0,00010402 | 9,19E-05 | Stil, Trip13, Gen1, Bub1, Chtf18, Mad2l1, Fbxo5, Bub1b, Ndc80 |
| GO:1905818 | regulation of chromosome separation | 10 | 1,22E-06 | 0,00010402 | 9,19E-05 | Stil, Cit, Trip13, Gen1, Cdc6, Bub1, Mad2l1, Fbxo5, Bub1b, Ndc80 |
| GO:0033045 | regulation of sister chromatid segregation | 11 | 1,95E-06 | 0,00015736 | 0,00013897 | Stil, Cit, Trip13, Gen1, Cdc6, Bub1, Cdca5, Mad2l1, Fbxo5, Bub1b, Ndc80 |
| GO:0006721 | terpenoid metabolic process | 9 | 2,03E-06 | 0,00015736 | 0,00013897 | Akr1b8, Plb1, Rdh9, Rdh1, Bco1, Dhrs9, Akr1c18, Aldh1a3, Rbp1 |
| GO:0030071 | regulation of mitotic metaphase/anaphase transition | 9 | 2,03E-06 | 0,00015736 | 0,00013897 | Stil, Trip13, Gen1, Cdc6, Bub1, Mad2l1, Fbxo5, Bub1b, Ndc80 |
| GO:0002138 | retinoic acid biosynthetic process | 5 | 2,05E-06 | 0,00015736 | 0,00013897 | Rdh9, Dhrs9, Akr1c18, Aldh1a3, Rbp1 |
| GO:0016102 | diterpenoid biosynthetic process | 5 | 2,05E-06 | 0,00015736 | 0,00013897 | Rdh9, Dhrs9, Akr1c18, Aldh1a3, Rbp1 |
| GO:1901988 | negative regulation of cell cycle phase transition | 15 | 2,36E-06 | 0,0001776 | 0,00015685 | Stil, E2f7, Brca1, Ticrr, Trip13, Clspn, Blm, Gen1, Dact1, Cdc6, Bub1, Mad2l1, Fbxo5, Bub1b, Ndc80 |
| GO:0033047 | regulation of mitotic sister chromatid segregation | 10 | 2,49E-06 | 0,00018378 | 0,00016231 | Stil, Cit, Trip13, Gen1, Cdc6, Bub1, Mad2l1, Fbxo5, Bub1b, Ndc80 |
| GO:0045841 | negative regulation of mitotic metaphase/anaphase transition | 8 | 2,54E-06 | 0,00018411 | 0,0001626 | Stil, Trip13, Gen1, Bub1, Mad2l1, Fbxo5, Bub1b, Ndc80 |
| GO:0045839 | negative regulation of mitotic nuclear division | 9 | 2,81E-06 | 0,00020006 | 0,00017668 | Stil, Trip13, Bmp4, Gen1, Bub1, Mad2l1, Fbxo5, Bub1b, Ndc80 |
| GO:2000816 | negative regulation of mitotic sister chromatid separation | 8 | 3,07E-06 | 0,00021513 | 0,00019 | Stil, Trip13, Gen1, Bub1, Mad2l1, Fbxo5, Bub1b, Ndc80 |
| GO:1902099 | regulation of metaphase/anaphase transition of cell cycle | 9 | 3,28E-06 | 0,00022559 | 0,00019923 | Stil, Trip13, Gen1, Cdc6, Bub1, Mad2l1, Fbxo5, Bub1b, Ndc80 |

| | | | | | | |
|---|---|---|---|---|---|---|
| GO:1902100 | negative regulation of metaphase/anaphase transition of cell cycle | 8 | 3,70E-06 | 0,00024674 | 0,00021791 | Stil, Trip13, Gen1, Bub1, Mad2l1, Fbxo5, Bub1b, Ndc80 |
| GO:1901990 | regulation of mitotic cell cycle phase transition | 19 | 3,71E-06 | 0,00024674 | 0,00021791 | Cdc7, Stil, E2f7, Brca1, Ticrr, Cdkn2a, Trip13, Clspn, Blm, Gen1, Dact1, Cdc6, Bub1, Cdca5, Mad2l1, Fbxo5, Cdc45, Bub1b, Ndc80 |
| GO:0045787 | positive regulation of cell cycle | 23 | 4,19E-06 | 0,00027399 | 0,00024197 | Cdc7, Stil, E2f7, Cit, Brca1, Brca2, Cdkn2a, Ccne1, Dyrk3, E2f8, Ccnf, Gen1, Ereg, Ccne2, Cdc6, Gata4, Cdca5, Mad2l1, Fbxo5, Cdc45, Ndc80, Pbp2, Spag5 |
| GO:0001523 | retinoid metabolic process | 8 | 4,44E-06 | 0,00028078 | 0,00024797 | Plb1, Rdh9, Rdh1, Bco1, Dhrs9, Akr1c18, Aldh1a3, Rbp1 |
| GO:1905819 | negative regulation of chromosome separation | 8 | 4,44E-06 | 0,00028078 | 0,00024797 | Stil, Trip13, Gen1, Bub1, Mad2l1, Fbxo5, Bub1b, Ndc80 |
| GO:0044839 | cell cycle G2/M phase transition | 13 | 4,90E-06 | 0,00030506 | 0,00026941 | Cdc7, Skp2, Cit, Brca1, Ticrr, Cdkn2a, Dyrk3, Foxm1, Clspn, Blm, Cdc6, Mastl, Fbxo5 |
| GO:1901987 | regulation of cell cycle phase transition | 20 | 5,17E-06 | 0,00031663 | 0,00027963 | Cdc7, Stil, E2f7, Brca1, Ticrr, Cdkn2a, Dyrk3, Trip13, Clspn, Blm, Gen1, Dact1, Cdc6, Bub1, Cdca5, Mad2l1, Fbxo5, Cdc45, Bub1b, Ndc80 |
| GO:0033048 | negative regulation of mitotic sister chromatid segregation | 8 | 5,30E-06 | 0,00031954 | 0,0002822 | Stil, Trip13, Gen1, Bub1, Mad2l1, Fbxo5, Bub1b, Ndc80 |
| GO:0000910 | cytokinesis | 14 | 6,24E-06 | 0,00036813 | 0,00032511 | E2f7, Cit, Ccp110, Brca2, Map9, E2f8, Anln, Incenp, Cdc6, Unc119, Zfp365, Kif4, Myh10, Pdxp |
| GO:0016101 | diterpenoid metabolic process | 8 | 6,29E-06 | 0,00036813 | 0,00032511 | Plb1, Rdh9, Rdh1, Bco1, Dhrs9, Akr1c18, Aldh1a3, Rbp1 |
| GO:0000086 | G2/M transition of mitotic cell cycle | 12 | 6,50E-06 | 0,00037104 | 0,00032768 | Cdc7, Skp2, Cit, Brca1, Ticrr, Cdkn2a, Foxm1, Clspn, Blm, Cdc6, Mastl, Fbxo5 |
| GO:0010639 | negative regulation of organelle organization | 24 | 6,53E-06 | 0,00037104 | 0,00032768 | Stil, Atad2, Capg, Brca1, Ccp110, Slit2, Top2a, Map2, Trip13, Plekhh2, Bnip3, Bmp4, Ccnf, Nav3, Mdm1, Gen1, Bub1, Mad2l1, Fbxo5, Nek2, Bub1b, Ndc80, Shank3, Pfn2 |
| GO:0033046 | negative regulation of sister chromatid segregation | 8 | 7,44E-06 | 0,00041656 | 0,00036788 | Stil, Trip13, Gen1, Bub1, Mad2l1, Fbxo5, Bub1b, Ndc80 |
| GO:0007076 | mitotic chromosome condensation | 5 | 8,40E-06 | 0,00046382 | 0,00040962 | Smc2, Ncapg, Cdca5, Ncapd2, Ncaph |
| GO:0051052 | regulation of DNA metabolic process | 25 | 9,92E-06 | 0,00053968 | 0,00047662 | Cdc7, Ccdc88a, Gja1, E2f7, Polq, Brca1, Ticrr, Brca2, Cdkn2a, Rad18, Fignl1, Foxm1, Esco2, E2f8, Rad51, Blm, Ereg, Dna2, Zfp365, Ankle1, Chtf18, Fbxo5, Rad51ap1, Nek2, Fancb |
| GO:0007140 | male meiotic nuclear division | 8 | 1,20E-05 | 0,00064383 | 0,0005686 | Rec8, Mybl1, Brca2, Fanca, Trip13, Fignl1, Suv39h2, Chtf18 |
| GO:0007098 | centrosome cycle | 12 | 1,22E-05 | 0,00064444 | 0,00056914 | Stil, Sass6, Brca1, Ccp110, Brca2, Map9, Ccnf, Mdm1, Gen1, Kif11, Xrcc2, Nek2 |
| GO:0016114 | terpenoid biosynthetic process | 5 | 1,24E-05 | 0,00064677 | 0,0005712 | Rdh9, Dhrs9, Akr1c18, Aldh1a3, Rbp1 |
| GO:0051784 | negative regulation of nuclear division | 9 | 1,32E-05 | 0,00068098 | 0,00060141 | Stil, Trip13, Bmp4, Gen1, Bub1, Mad2l1, Fbxo5, Bub1b, Ndc80 |
| GO:0010569 | regulation of double-strand break repair via homologous recombination | 7 | 1,94E-05 | 0,00097689 | 0,00086274 | Polq, Fignl1, Rad51, Blm, Zfp365, Rad51ap1, Fancb |
| GO:0071174 | mitotic spindle checkpoint | 7 | 1,94E-05 | 0,00097689 | 0,00086274 | Stil, Trip13, Gen1, Bub1, Mad2l1, Bub1b, Ndc80 |
| GO:0010212 | response to ionizing radiation | 12 | 2,01E-05 | 0,00099559 | 0,00087925 | Ddias, Rad54b, Brca1, Ticrr, Brca2, Fancd2, Fignl1, Rad51, Blm, Rad54l, Xrcc2, Rad51ap1 |
| GO:1904666 | regulation of ubiquitin protein ligase activity | 5 | 2,46E-05 | 0,00120492 | 0,00106412 | Cdkn2a, Mastl, Mad2l1, Bag2, Fbxo5 |
| GO:0090068 | positive regulation of cell cycle process | 17 | 2,55E-05 | 0,0012359 | 0,00109148 | Cdc7, E2f7, Cit, Brca1, Cdkn2a, Dyrk3, E2f8, Gen1, Ereg, Cdc6, Cdca5, Mad2l1, Fbxo5, Cdc45, Ndc80, Pbp2, Spag5 |
| GO:0046394 | carboxylic acid biosynthetic process | 21 | 2,65E-05 | 0,00126561 | 0,00111772 | Ptges, Myo5a, Apoc2, Gch1, Mtap, Brca1, Fads3, Psat1, Rdh9, Ldhb, Fabp5, Trib3, Hacd4, Ass1, Acot12, Phgdh, Dhrs9, Akr1c18, Ptgis, Aldh1a3, Rbp1 |
| GO:0051307 | meiotic chromosome separation | 6 | 2,68E-05 | 0,00126561 | 0,00111772 | Hfm1, Top2a, Blm, Espl1, Ncaph, Eme1 |
| GO:0031577 | spindle checkpoint | 7 | 2,74E-05 | 0,00127669 | 0,00112751 | Stil, Trip13, Gen1, Bub1, Mad2l1, Bub1b, Ndc80 |
| GO:0016053 | organic acid biosynthetic process | 21 | 2,77E-05 | 0,00127686 | 0,00112765 | Ptges, Myo5a, Apoc2, Gch1, Mtap, Brca1, Fads3, Psat1, Rdh9, Ldhb, Fabp5, Trib3, Hacd4, Ass1, Acot12, Phgdh, Dhrs9, Akr1c18, Ptgis, Aldh1a3, Rbp1 |
| GO:0006720 | isoprenoid metabolic process | 9 | 3,02E-05 | 0,00137849 | 0,00121741 | Akr1b8, Plb1, Rdh9, Rdh1, Bco1, Dhrs9, Akr1c18, Aldh1a3, Rbp1 |
| GO:0051302 | regulation of cell division | 13 | 3,62E-05 | 0,0016294 | 0,001439 | E2f7, Cit, Ccp110, Brca2, Map9, Pdgfc, E2f8, Blm, Ereg, Incenp, Cdc6, Kif18b, Pdxp |
| GO:0031023 | microtubule organizing center organization | 12 | 3,73E-05 | 0,00165001 | 0,0014572 | Stil, Sass6, Brca1, Ccp110, Brca2, Map9, Ccnf, Mdm1, Gen1, Kif11, Xrcc2, Nek2 |
| GO:2001020 | regulation of response to DNA damage stimulus | 15 | 3,75E-05 | 0,00165001 | 0,0014572 | Polq, Ddias, Brca1, Cdkn2a, Dyrk3, Fignl1, Foxm1, Rad51, Blm, Zfp385a, Zfp365, Ankle1, Fbxo5, Rad51ap1, Fancb |
| GO:0045143 | homologous chromosome segregation | 8 | 4,17E-05 | 0,00181538 | 0,00160325 | Rec8, Brca2, Ccne1, Fancd2, Ndc1, Trip13, Espl1, Ccne2 |
| GO:1902969 | mitotic DNA replication | 4 | 4,30E-05 | 0,00183216 | 0,00161807 | Brca2, Rad51, Cdc45, Pola1 |
| GO:1904667 | negative regulation of ubiquitin protein ligase activity | 4 | 4,30E-05 | 0,00183216 | 0,00161807 | Cdkn2a, Mad2l1, Bag2, Fbxo5 |
| GO:0032465 | regulation of cytokinesis | 9 | 4,65E-05 | 0,00196066 | 0,00173155 | E2f7, Cit, Ccp110, Brca2, Map9, E2f8, Incenp, Cdc6, Pdxp |
| GO:0007131 | reciprocal meiotic recombination | 7 | 5,12E-05 | 0,00209075 | 0,00184644 | Hfm1, Top2a, Trip13, Rad51, Blm, Chtf18, Eme1 |
| GO:0030261 | chromosome condensation | 7 | 5,12E-05 | 0,00209075 | 0,00184644 | Smc2, Top2a, Ncapg2, Ncapg, Cdca5, Ncapd2, Ncaph |
| GO:0035825 | homologous recombination | 7 | 5,12E-05 | 0,00209075 | 0,00184644 | Hfm1, Top2a, Trip13, Rad51, Blm, Chtf18, Eme1 |
| GO:0007051 | spindle organization | 13 | 5,75E-05 | 0,00231044 | 0,00204046 | Stil, Mybl2, Map9, Espl1, Kifc5b, Nuf2, Kif11, Spc25, Kif4, Fbxo5, Nek2, Ndc80, Spag5 |
| GO:0001889 | liver development | 11 | 5,80E-05 | 0,00231044 | 0,00204046 | Rarb, E2f7, Mki67, Cadm1, Pdx1, Proc, Cdkn2a, Foxm1, E2f8, Bmp4, Sulf2 |
| GO:0042573 | retinoic acid metabolic process | 5 | 5,84E-05 | 0,00231044 | 0,00204046 | Rdh9, Dhrs9, Akr1c18, Aldh1a3, Rbp1 |
| GO:0033260 | nuclear DNA replication | 6 | 5,99E-05 | 0,00234587 | 0,00207175 | Cdc7, Brca2, Rad51, Dna2, Cdc45, Pola1 |
| GO:0000281 | mitotic cytokinesis | 9 | 6,97E-05 | 0,00270257 | 0,00238677 | Cit, Brca2, Map9, Anln, Incenp, Unc119, Zfp365, Kif4, Myh10 |
| GO:0061008 | hepaticobiliary system development | 11 | 7,30E-05 | 0,00280411 | 0,00247644 | Rarb, E2f7, Mki67, Cadm1, Pdx1, Proc, Cdkn2a, Foxm1, E2f8, Bmp4, Sulf2 |

| | | | | | | |
|---|---|---|---|---|---|---|
| GO:0051298 | centrosome duplication | 8 | 7,56E-05 | 0,00287756 | 0,00254131 | Stil, Sass6, Brca1, Ccp110, Brca2, Ccnf, Mdm1, Gen1 |
| GO:2000779 | regulation of double-strand break repair | 8 | 9,45E-05 | 0,00355952 | 0,00314358 | Polq, Fignl1, Foxm1, Rad51, Blm, Zfp365, Rad51ap1, Fancb |
| GO:0007143 | female meiotic nuclear division | 6 | 0,00010142 | 0,0037853 | 0,00334298 | Top2a, Trip13, Ereg, Mastl, Fbxo5, Ncaph |
| GO:0007052 | mitotic spindle organization | 10 | 0,00011318 | 0,00418434 | 0,00369539 | Stil, Mybl2, Map9, Kifc5b, Nuf2, Kif11, Spc25, Kif4, Nek2, Ndc80 |
| GO:0072330 | monocarboxylic acid biosynthetic process | 16 | 0,00012582 | 0,0046084 | 0,0040699 | Ptges, Myo5a, Apoc2, Brca1, Fads3, Rdh9, Ldhb, Fabp5, Trib3, Hacd4, Acot12, Dhrs9, Akr1c18, Ptgis, Aldh1a3, Rbp1 |
| GO:0042770 | signal transduction in response to DNA damage | 9 | 0,00013327 | 0,00483606 | 0,00427095 | E2f7, Brca1, Brca2, Cdkn2a, Dyrk3, Foxm1, Zfp385a, Gen1, Rps6ka6 |
| GO:0007094 | mitotic spindle assembly checkpoint | 6 | 0,00013997 | 0,00498691 | 0,00440417 | Trip13, Gen1, Bub1, Mad2l1, Bub1b, Ndc80 |
| GO:0071173 | spindle assembly checkpoint | 6 | 0,00013997 | 0,00498691 | 0,00440417 | Trip13, Gen1, Bub1, Mad2l1, Bub1b, Ndc80 |
| GO:0061640 | cytoskeleton-dependent cytokinesis | 9 | 0,00018762 | 0,00662405 | 0,00585001 | Cit, Brca2, Map9, Anln, Incenp, Unc119, Zfp365, Kif4, Myh10 |
| GO:0000076 | DNA replication checkpoint | 4 | 0,0001912 | 0,00669024 | 0,00590846 | Ticrr, Clspn, Dna2, Cdc6 |
| GO:0071103 | DNA conformation change | 14 | 0,00021217 | 0,00735844 | 0,00649858 | Smc2, Cdkn2a, Top2a, Asf1b, Rad51, Blm, Ncapg2, Hist1h3d, Chaf1a, Ncapg, Cdca5, Ncapd2, Oip5, Ncaph |
| GO:0006312 | mitotic recombination | 5 | 0,00022217 | 0,00763774 | 0,00674525 | Brca2, Rad51, Blm, Gen1, Ankle1 |
| GO:1902749 | regulation of cell cycle G2/M phase transition | 9 | 0,00023951 | 0,00816223 | 0,00720845 | Cdc7, Brca1, Ticrr, Cdkn2a, Dyrk3, Clspn, Blm, Cdc6, Fbxo5 |
| GO:0006978 | DNA damage response, signal transduction by p53 class mediator resulting in transcription of p21 class mediator | 4 | 0,00025622 | 0,00850968 | 0,0075153 | Brca2, Foxm1, Zfp385a, Rps6ka6 |
| GO:0042953 | lipoprotein transport | 4 | 0,00025622 | 0,00850968 | 0,0075153 | Apoc2, Apob, Unc119, Mttp |
| GO:0044872 | lipoprotein localization | 4 | 0,00025622 | 0,00850968 | 0,0075153 | Apoc2, Apob, Unc119, Mttp |
| GO:0046605 | regulation of centrosome cycle | 7 | 0,00026562 | 0,00874566 | 0,00772371 | Stil, Map9, Ccnf, Mdm1, Gen1, Kif11, Nek2 |
| GO:0036297 | interstrand cross-link repair | 5 | 0,00026779 | 0,00874566 | 0,00772371 | Fanca, Rad51, Mcm8, Rad51ap1, Fancb |
| GO:0007292 | female gamete generation | 11 | 0,00030226 | 0,00978974 | 0,00864577 | Hfm1, Rec8, Brca2, Top2a, Trip13, Plat, Ereg, Mastl, Mcm8, Fbxo5, Ncaph |
| GO:0000018 | regulation of DNA recombination | 9 | 0,00032673 | 0,01041602 | 0,00919887 | Polq, Rad18, Fignl1, Rad51, Blm, Zfp365, Ankle1, Rad51ap1, Fancb |
| GO:0006323 | DNA packaging | 12 | 0,00032691 | 0,01041602 | 0,00919887 | Smc2, Cdkn2a, Top2a, Asf1b, Ncapg2, Hist1h3d, Chaf1a, Ncapg, Cdca5, Ncapd2, Oip5, Ncaph |
| GO:0042772 | DNA damage response, signal transduction resulting in transcription | 4 | 0,00033574 | 0,01061097 | 0,00937105 | Brca2, Foxm1, Zfp385a, Rps6ka6 |
| GO:2001251 | negative regulation of chromosome organization | 11 | 0,00036183 | 0,01126556 | 0,00994914 | Stil, Atad2, Brca1, Top2a, Trip13, Gen1, Bub1, Mad2l1, Fbxo5, Bub1b, Ndc80 |
| GO:0030330 | DNA damage response, signal transduction by p53 class mediator | 7 | 0,00036429 | 0,01126556 | 0,00994914 | E2f7, Brca2, Cdkn2a, Dyrk3, Foxm1, Zfp385a, Rps6ka6 |
| GO:0010389 | regulation of G2/M transition of mitotic cell cycle | 8 | 0,00036507 | 0,01126556 | 0,00994914 | Cdc7, Brca1, Ticrr, Cdkn2a, Clspn, Blm, Cdc6, Fbxo5 |
| GO:0021756 | striatum development | 4 | 0,00043147 | 0,01321055 | 0,01166686 | Rarb, Slc7a11, Shank3, Aldh1a3 |
| GO:0008608 | attachment of spindle microtubules to kinetochore | 5 | 0,00044783 | 0,01360496 | 0,01201518 | Brca2, Nuf2, Nek2, Ndc80, Spag5 |
| GO:0001556 | oocyte maturation | 5 | 0,00052442 | 0,01561925 | 0,01379409 | Rec8, Brca2, Trip13, Ereg, Fbxo5 |
| GO:0008299 | isoprenoid biosynthetic process | 5 | 0,00052442 | 0,01561925 | 0,01379409 | Rdh9, Dhrs9, Akr1c18, Aldh1a3, Rbp1 |
| GO:1902850 | microtubule cytoskeleton organization involved in mitosis | 10 | 0,00052609 | 0,01561925 | 0,01379409 | Stil, Mybl2, Map9, Kifc5b, Nuf2, Kif11, Spc25, Kif4, Nek2, Ndc80 |
| GO:0051438 | regulation of ubiquitin-protein transferase activity | 6 | 0,00053745 | 0,0157146 | 0,0138783 | Cdkn2a, Trib3, Mastl, Mad2l1, Bag2, Fbxo5 |
| GO:0000712 | resolution of meiotic recombination intermediates | 4 | 0,0005452 | 0,0157146 | 0,0138783 | Hfm1, Top2a, Blm, Eme1 |
| GO:0051444 | negative regulation of ubiquitin-protein transferase activity | 4 | 0,0005452 | 0,0157146 | 0,0138783 | Cdkn2a, Mad2l1, Bag2, Fbxo5 |
| GO:0022412 | cellular process involved in reproduction in multicellular organism | 21 | 0,00054534 | 0,0157146 | 0,0138783 | Hfm1, Rec8, Spag16, Mybl1, Ddias, Bmp8b, Brca2, Fanca, Top2a, Trip13, Fignl1, Bmp4, Zfp37, Ereg, Suv39h2, Mastl, Chtf18, Fbxo5, Ncaph, Pbp2, Jam3 |
| GO:0000079 | regulation of cyclin-dependent protein serine/threonine kinase activity | 7 | 0,00059205 | 0,01687939 | 0,01490698 | Stil, Cdkn2a, Ccne1, Ccnf, Blm, Ccne2, Cdc6 |
| GO:0006633 | fatty acid biosynthetic process | 10 | 0,00059437 | 0,01687939 | 0,01490698 | Ptges, Myo5a, Apoc2, Brca1, Fads3, Fabp5, Trib3, Hacd4, Acot12, Ptgis |
| GO:0007129 | synapsis | 6 | 0,00060301 | 0,01700138 | 0,01501471 | Rec8, Ccne1, Fancd2, Ndc1, Trip13, Ccne2 |
| GO:0044818 | mitotic G2/M transition checkpoint | 5 | 0,00061045 | 0,01708828 | 0,01509146 | Brca1, Ticrr, Clspn, Blm, Cdc6 |
| GO:0001701 | in utero embryonic development | 23 | 0,00062484 | 0,0173671 | 0,0153377 | Eomes, Tgfbr2, Gja1, Stil, E2f7, Myo18b, Jag2, Brca2, Ppp4r4, Slit2, Arnt2, Asf1b, E2f8, Ncapg2, Arhgdig, Mdfi, Apob, Gata4, Xrcc2, Slc34a2, Nek2, B9d1, Myh10 |
| GO:0006282 | regulation of DNA repair | 9 | 0,00066413 | 0,0183291 | 0,01618729 | Polq, Brca1, Fignl1, Foxm1, Rad51, Blm, Zfp365, Rad51ap1, Fancb |
| GO:0033044 | regulation of chromosome organization | 18 | 0,00068246 | 0,01870316 | 0,01651763 | Stil, Atad2, Cit, Mki67, Brca1, Top2a, Trip13, Wdhd1, Gen1, Cdc6, Bub1, Cdca5, Mad2l1, Fbxo5, Nek2, Cdc45, Bub1b, Ndc80 |
| GO:0009798 | axis specification | 8 | 0,00069534 | 0,01892392 | 0,0167126 | Stil, Pcsk6, Tcf7l1, Mns1, Pgap1, Bmp4, Mdfi, Gpc3 |
| GO:0006081 | cellular aldehyde metabolic process | 7 | 0,00070987 | 0,01906943 | 0,01684111 | Aldh3b2, Idh1, Bco1, Akr1c18, Aldh3b3, Pdxp, Aldh1a3 |
| GO:0072331 | signal transduction by p53 class mediator | 10 | 0,00071042 | 0,01906943 | 0,01684111 | Eda2r, E2f7, Phlda3, Brca2, Cdkn2a, Dyrk3, Foxm1, E2f2, Zfp385a, Rps6ka6 |
| GO:0031570 | DNA integrity checkpoint | 9 | 0,00075752 | 0,02019539 | 0,01783549 | Brca1, Ticrr, Top2a, Clspn, Blm, Gen1, Dna2, Cdc6, Eme1 |
| GO:2001021 | negative regulation of response to DNA damage stimulus | 7 | 0,00077536 | 0,02053125 | 0,01813211 | Polq, Ddias, Dyrk3, Blm, Zfp385a, Fbxo5, Fancb |
| GO:1904029 | regulation of cyclin-dependent protein kinase activity | 7 | 0,00084553 | 0,02223902 | 0,01964032 | Stil, Cdkn2a, Ccne1, Ccnf, Blm, Ccne2, Cdc6 |
| GO:0040013 | negative regulation of locomotion | 17 | 0,0009274 | 0,02422996 | 0,02139861 | Fbln1, Il1rn, Slit2, Dusp22, Trim35, Nav3, Sema3d, Adgrb1, Magi2, Il33, Sema3f, Ccl25, Sema7a, Ndrg4, Dpysl3, Pfn2, Adamts9 |

| | | | | | | | | | | | |
|---|---|---|---|---|---|---|---|---|---|---|---|
| GO:0033599 | regulation of mammary gland epithelial cell proliferation | 4 | 0,00101226 | 0,02627192 | 0,02320196 | Brca2, Cdkn2a, Mst1, Iqgap3 | | | | | |
| GO:0072006 | nephron development | 10 | 0,00111387 | 0,02789665 | 0,02463684 | Lama5, Nid1, Npnt, Bmp4, Sulf2, Enpep, Notch3, Irx3, Wwtr1, Gpc3 | | | | | |
| GO:0008300 | isoprenoid catabolic process | 3 | 0,00111757 | 0,02789665 | 0,02463684 | Akr1b8, Bco1, Akr1c18 | | | | | |
| GO:0032875 | regulation of DNA endoreduplication | 3 | 0,00111757 | 0,02789665 | 0,02463684 | E2f7, E2f8, Fbxo5 | | | | | |
| GO:0033314 | mitotic DNA replication checkpoint | 3 | 0,00111757 | 0,02789665 | 0,02463684 | Ticrr, Clspn, Cdc6 | | | | | |
| GO:1903365 | regulation of fear response | 3 | 0,00111757 | 0,02789665 | 0,02463684 | Gja1, Penk, Shank3 | | | | | |
| GO:2000822 | regulation of behavioral fear response | 3 | 0,00111757 | 0,02789665 | 0,02463684 | Gja1, Penk, Shank3 | | | | | |
| GO:2000377 | regulation of reactive oxygen species metabolic process | 12 | 0,00112798 | 0,02797818 | 0,02470885 | Tgfbr2, Gch1, Ulbp1, Brca1, Foxm1, Bnip3, Adgrb1, Ass1, Akr1c18, Ptgis, Plin5, Duoxa1 | | | | | |
| GO:0007141 | male meiosis I | 4 | 0,00121596 | 0,02997083 | 0,02646864 | Rec8, Mybl1, Brca2, Trip13 | | | | | |
| GO:0051271 | negative regulation of cellular component movement | 16 | 0,00139429 | 0,03415142 | 0,03016072 | Fbln1, Il1rn, Slit2, Dusp22, Nav3, Sema3d, Adgrb1, Magi2, Il33, Sema3f, Ccl25, Sema7a, Ndrg4, Dpysl3, Pfn2, Adamts9 | | | | | |
| GO:0021544 | subpallium development | 4 | 0,0014467 | 0,03521494 | 0,03109997 | Rarb, Slc7a11, Shank3, Aldh1a3 | | | | | |
| GO:0048640 | negative regulation of developmental growth | 9 | 0,00148125 | 0,03583353 | 0,03164627 | Tgfbr2, Gja1, Gdf15, Slit2, Map2, Bmp4, Sema3d, Sema3f, Sema7a | | | | | |
| GO:0051225 | spindle assembly | 8 | 0,00150353 | 0,03590655 | 0,03171075 | Mybl2, Map9, Kifc5b, Kif11, Kif4, Fbxo5, Nek2, Spag5 | | | | | |
| GO:0042023 | DNA endoreduplication | 3 | 0,00151176 | 0,03590655 | 0,03171075 | E2f7, E2f8, Fbxo5 | | | | | |
| GO:2000105 | positive regulation of DNA-dependent DNA replication | 3 | 0,00151176 | 0,03590655 | 0,03171075 | Cdc7, E2f7, E2f8 | | | | | |
| GO:0048732 | gland development | 21 | 0,00158312 | 0,0373749 | 0,03300752 | Lama5, Tgfbr2, Rarb, Gja1, E2f7, Mki67, Cadm1, Pdx1, Brca2, Proc, Cdkn2a, Slit2, Foxm1, E2f8, Bmp4, Sulf2, Mst1, Iqgap3, Id4, Hesx1, Aldh1a3 | | | | | |
| GO:0032835 | glomerulus development | 6 | 0,00165485 | 0,03883454 | 0,0342966 | Nid1, Bmp4, Sulf2, Enpep, Notch3, Wwtr1 | | | | | |
| GO:0031345 | negative regulation of cell projection organization | 12 | 0,0016661 | 0,03886583 | 0,03432424 | Cit, Ccp110, Slit2, Map2, Sema3d, Sema3f, Lgals1, Zfp365, Sema7a, Dpysl3, Pagr3, Pfn2 | | | | | |
| GO:0019433 | triglyceride catabolic process | 4 | 0,00170627 | 0,03956726 | 0,0349437 | Apoc2, Apob, Cps1, Plin5 | | | | | |
| GO:0051494 | negative regulation of cytoskeleton organization | 10 | 0,00177378 | 0,04089084 | 0,03611262 | Capg, Slit2, Map2, Plekhh2, Ccnf, Nav3, Mdm1, Nek2, Shank3, Pfn2 | | | | | |
| GO:0090307 | mitotic spindle assembly | 6 | 0,00197317 | 0,04440883 | 0,03921952 | Mybl2, Map9, Kifc5b, Kif11, Kif4, Nek2 | | | | | |
| GO:0007100 | mitotic centrosome separation | 3 | 0,00198304 | 0,04440883 | 0,03921952 | Map9, Kif11, Nek2 | | | | | |
| GO:0007135 | meiosis II | 3 | 0,00198304 | 0,04440883 | 0,03921952 | Bub1, Mastl, Bub1b | | | | | |
| GO:0060433 | bronchus development | 3 | 0,00198304 | 0,04440883 | 0,03921952 | Tgfbr2, Tulp3, Bmp4 | | | | | |
| GO:0061983 | meiosis II cell cycle process | 3 | 0,00198304 | 0,04440883 | 0,03921952 | Bub1, Mastl, Bub1b | | | | | |
| GO:0010972 | negative regulation of G2/M transition of mitotic cell cycle | 5 | 0,00215522 | 0,04771919 | 0,04214305 | Brca1, Ticrr, Clspn, Blm, Cdc6 | | | | | |
| GO:0061098 | positive regulation of protein tyrosine kinase activity | 5 | 0,00215522 | 0,04771919 | 0,04214305 | Ncapg2, Ereg, Gprc5b, Lrp8, Unc119 | | | | | |
| GO:0042110 | T cell activation | | | | | | 51 | 4,70E-25 | 1,71E-21 | 1,35E-21 | Ccl5, Vav1, Cd3g, Apbb1ip, Cd3e, Card11, Cd74, Tbx21, H2-DMa, Ptprc, Zap70, Ptpn22, Xcl1, Rac2, Itk, Gpr18, Dtx1, Rasal3, H2-Ab1, Egr3, Spn, Lax1, Sla2, Il2rg, Dock2, Coro1a, H2-Aa, Thy1, Vsir, Rhoh, Sash3, Lck, Cd48, Il7r, Txk, Cd2, Fyn, Itgb2, Lat, Nlrc3, Was, Cd8a, Nckap1l, Cd3d, Ikzf1, Fcer1g, Gimap1, Ly9, Cdh26, Clec2i, Cd27 |
| GO:0030217 | T cell differentiation | | | | | | 32 | 9,37E-18 | 1,25E-14 | 9,87E-15 | Vav1, Cd3g, Cd3e, Card11, Cd74, Tbx21, H2-DMa, Ptprc, Zap70, Ptpn22, Itk, Gpr18, Dtx1, Egr3, Spn, Il2rg, Dock2, H2-Aa, Vsir, Rhoh, Sash3, Lck, Il7r, Txk, Cd8a, Nckap1l, Cd3d, Ikzf1, Fcer1g, Gimap1, Ly9, Cd27 |
| GO:0030098 | lymphocyte differentiation | | | | | | 38 | 1,03E-17 | 1,25E-14 | 9,87E-15 | Dll1, Vav1, Cd3g, Cd3e, Card11, Cd74, Tbx21, H2-DMa, Inpp5d, Ptprc, Zap70, Ptpn22, Itk, Gpr18, Dtx1, H2-Ab1, Egr3, Spn, Il2rg, Dock2, H2-Aa, Vsir, Rhoh, Sash3, Lck, Dock10, Il7r, Txk, Ikzf3, Cd8a, Nckap1l, Cd3d, Ikzf1, Mfng, Fcer1g, Gimap1, Ly9, Cd27 |
| GO:0007159 | leukocyte cell-cell adhesion | | | | | | 33 | 1,45E-16 | 1,32E-13 | 1,04E-13 | Ccl5, Cd3e, Card11, Cd74, Tbx21, H2-DMa, Ptprc, Zap70, Ptpn22, Xcl1, Rac2, Dtx1, Selplg, Rasal3, H2-Ab1, Egr3, Spn, Lax1, Il2rg, Nr4a3, Coro1a, H2-Aa, Thy1, Skap1, Vsir, Sash3, Lck, Il7r, Itgb7, Itgb2, Nckap1l, Ikzf1, Cd27 |
| GO:1903039 | positive regulation of leukocyte cell-cell adhesion | | | | | | 26 | 1,02E-15 | 7,38E-13 | 5,84E-13 | Ccl5, Cd3e, Card11, Cd74, H2-DMa, Ptprc, Zap70, Ptpn22, Xcl1, Rasal3, H2-Ab1, Egr3, Spn, Il2rg, Nr4a3, Coro1a, H2-Aa, Thy1, Skap1, Vsir, Sash3, Lck, Il7r, Nckap1l, Ikzf1, Cd27 |
| GO:0050863 | regulation of T cell activation | | | | | | 30 | 1,37E-14 | 8,28E-12 | 6,55E-12 | Ccl5, Cd3e, Card11, Cd74, Tbx21, H2-DMa, Ptprc, Zap70, Ptpn22, Xcl1, Rac2, Dtx1, Rasal3, H2-Ab1, Egr3, Spn, Lax1, Il2rg, Coro1a, H2-Aa, Thy1, Vsir, Sash3, Lck, Il7r, Lat, Nckap1l, Ikzf1, Clec2i, Cd27 |
| GO:1903037 | regulation of leukocyte cell-cell adhesion | | | | | | 29 | 1,64E-14 | 8,52E-12 | 6,74E-12 | Ccl5, Cd3e, Card11, Cd74, Tbx21, H2-DMa, Ptprc, Zap70, Ptpn22, Xcl1, Dtx1, Rasal3, H2-Ab1, Egr3, Spn, Lax1, Il2rg, Nr4a3, Coro1a, H2-Aa, Thy1, Skap1, Vsir, Sash3, Lck, Il7r, Nckap1l, Ikzf1, Cd27 |
| GO:0050870 | positive regulation of T cell activation | | | | | | 24 | 2,16E-14 | 9,78E-12 | 7,73E-12 | Ccl5, Cd3e, Card11, Cd74, H2-DMa, Ptprc, Zap70, Ptpn22, Xcl1, Rasal3, H2-Ab1, Egr3, Spn, Il2rg, Coro1a, H2-Aa, Thy1, Vsir, Sash3, Lck, Il7r, Nckap1l, Ikzf1, Cd27 |
| mmu04658 | Th1 and Th2 cell differentiation | | | | | | 19 | 6,86E-14 | 1,62E-11 | 1,25E-11 | 13388, 12502, 12501, 57765, 14998, 14999, 22637, 270118, 12503, 14961, 14969, 16186, 14960, 20849, 16185, 16818, 18037, 16797, 12500 |
| GO:0022409 | positive regulation of cell-cell adhesion | | | | | | 26 | 7,78E-14 | 3,14E-11 | 2,48E-11 | Ccl5, Cd3e, Card11, Cd74, H2-DMa, Ptprc, Zap70, Ptpn22, Xcl1, Rasal3, H2-Ab1, Egr3, Spn, Il2rg, Nr4a3, Coro1a, H2-Aa, Thy1, Skap1, Vsir, Sash3, Lck, Il7r, Nckap1l, Ikzf1, Cd27 |
| mmu04650 | Natural killer cell mediated cytotoxicity | | | | | | 20 | 2,31E-12 | 2,73E-10 | 2,10E-10 | 22324, 22325, 27007, 16643, 14939, 14103, 22637, 19354, 12503, 16818, 18707, 12506, 14360, 16822, 16414, 16797, 22035, 16638, 14127, 23900 |
| mmu04660 | T cell receptor signaling pathway | | | | | | 18 | 1,27E-11 | 9,98E-10 | 7,70E-10 | 22324, 12502, 12501, 108723, 22325, 19264, 22637, 12503, 16428, 16818, 18707, 14360, 16822, 18037, 16797, 17444, 12525, 12500 |

| | | | | | | |
|---|---|---|---|---|---|---|
| GO:0046631 | alpha-beta T cell activation | 20 | 3,49E-12 | 1,27E-09 | 1,00E-09 | Cd3e, Tbx21, Ptprc, Zap70, Ptpn22, Xcl1, Itk, Gpr18, Rasal3, H2-Ab1, Spn, Il2rg, Dock2, Vsir, Sash3, Txk, Nckap1l, Ikzf1, Ly9, Cdh26 |
| GO:0002443 | leukocyte mediated immunity | 35 | 4,41E-12 | 1,46E-09 | 1,15E-09 | Chga, Vav1, H2-T3, Cd74, Tbx21, H2-DMa, Inpp5d, Klrk1, Klrd1, H2-T10, Gzmb, Ptprc, Xcl1, Rac2, H2-Ab1, Cd96, H2-Bl, Spn, Nr4a3, Coro1a, Serpinb9, Vsir, Sash3, Tlr2, Il7r, Itgb2, Myo1f, Lat, Lat2, Was, Cd8a, Klre1, Myo1g, Fcer1g, C9 |
| GO:0045785 | positive regulation of cell adhesion | 31 | 6,06E-12 | 1,83E-09 | 1,45E-09 | Ccl5, Vav1, Apbb1ip, Cd3e, Card11, Cd74, H2-DMa, Thbs1, Ptprc, Zap70, Ptpn22, Xcl1, Smoc2, Rasal3, H2-Ab1, Egr3, Spn, Il2rg, Nr4a3, Coro1a, H2-Aa, Thy1, Skap1, Vsir, Sash3, Lck, Il7r, Nckap1l, Ikzf1, Smoc1, Cd27 |
| GO:0050867 | positive regulation of cell activation | 33 | 8,47E-12 | 2,36E-09 | 1,87E-09 | Ccl5, Cd3e, Card11, Cd74, Tbx21, H2-DMa, Thbs1, Inpp5d, Klrk1, Ptprc, Zap70, Ptpn22, Xcl1, Rasal3, H2-Ab1, Egr3, Spn, Il2rg, Nr4a3, Coro1a, H2-Aa, Thy1, Vsir, Sash3, Lck, Il7r, Itgb2, Nckap1l, Ikzf1, Fcer1g, Capn3, Clec2i, Cd27 |
| GO:0022407 | regulation of cell-cell adhesion | 30 | 1,03E-11 | 2,66E-09 | 2,11E-09 | Ccl5, Adamts18, Cd3e, Card11, Cd74, Tbx21, H2-DMa, Ptprc, Zap70, Ptpn22, Xcl1, Dtx1, Rasal3, H2-Ab1, Egr3, Spn, Lax1, Il2rg, Nr4a3, Coro1a, H2-Aa, Thy1, Skap1, Vsir, Sash3, Lck, Il7r, Nckap1l, Ikzf1, Cd27 |
| GO:0045061 | thymic T cell selection | 10 | 1,69E-11 | 3,98E-09 | 3,15E-09 | Cd3g, Cd3e, Card11, Cd74, H2-DMa, Ptprc, Zap70, Spn, Dock2, Cd3d |
| GO:0002696 | positive regulation of leukocyte activation | 32 | 1,76E-11 | 3,98E-09 | 3,15E-09 | Ccl5, Cd3e, Card11, Cd74, Tbx21, H2-DMa, Thbs1, Inpp5d, Klrk1, Ptprc, Zap70, Ptpn22, Xcl1, Rasal3, H2-Ab1, Egr3, Spn, Il2rg, Nr4a3, Coro1a, H2-Aa, Thy1, Vsir, Sash3, Lck, Il7r, Itgb2, Nckap1l, Ikzf1, Fcer1g, Clec2i, Cd27 |
| GO:0050900 | leukocyte migration | 27 | 3,42E-11 | 7,14E-09 | 5,64E-09 | Chga, Ccl5, Edn2, Vav1, Cd74, Tbx21, Thbs1, Klrk1, Ptpn22, Xcl1, Rac2, Gpr18, Ccl4, Selplg, Spn, Edn1, Pde4b, Coro1a, Thy1, Tlr2, Itgb7, Itgb2, Nckap1l, Myo1g, Fcer1g, Pdgfd, Gpsm3 |
| GO:0043368 | positive T cell selection | 11 | 3,54E-11 | 7,14E-09 | 5,64E-09 | Cd3g, Cd3e, Cd74, Tbx21, H2-DMa, Ptprc, Zap70, Spn, Dock2, Cd3d, Ly9 |
| GO:0032943 | mononuclear cell proliferation | 26 | 3,92E-11 | 7,49E-09 | 5,92E-09 | Ccl5, Cd3e, Card11, Cd74, Inpp5d, Ptprc, Zap70, Ptpn22, Xcl1, Rac2, Rasal3, H2-Ab1, Spn, Dock2, Coro1a, H2-Aa, Vsir, Sash3, Il7r, Fyn, Ikzf3, St6gal1, Itgb2, Nckap1l, Clec2i, Cd27 |
| mmu04659 | Th17 cell differentiation | 17 | 1,46E-10 | 8,64E-09 | 6,66E-09 | 12502, 12501, 57765, 14998, 14999, 22637, 12503, 14961, 60504, 14969, 16186, 14960, 16185, 16818, 18037, 16797, 12500 |
| GO:0002429 | immune response-activating cell surface receptor signaling pathway | 26 | 9,22E-11 | 1,67E-08 | 1,32E-08 | Cd3e, Klrk1, Ptprc, Fyb, Zap70, Ptpn22, Cd247, Itk, Lax1, Nr4a3, Pde4b, Thy1, Skap1, Lck, Tlr2, Txk, Fyn, Lcp2, Lpxn, Trat1, Lat2, Nckap1l, Klre1, Myo1g, Fcer1g, Clec2i |
| GO:0045059 | positive thymic T cell selection | 8 | 9,98E-11 | 1,72E-08 | 1,36E-08 | Cd3g, Cd3e, Cd74, H2-DMa, Ptprc, Zap70, Dock2, Cd3d |
| GO:0070661 | leukocyte proliferation | 26 | 1,14E-10 | 1,87E-08 | 1,48E-08 | Ccl5, Cd3e, Card11, Cd74, Inpp5d, Ptprc, Zap70, Ptpn22, Xcl1, Rac2, Rasal3, H2-Ab1, Spn, Dock2, Coro1a, H2-Aa, Vsir, Sash3, Il7r, Fyn, Ikzf3, St6gal1, Itgb2, Nckap1l, Clec2i, Cd27 |
| GO:0045058 | T cell selection | 12 | 1,72E-10 | 2,71E-08 | 2,14E-08 | Cd3g, Cd3e, Card11, Cd74, Tbx21, H2-DMa, Ptprc, Zap70, Spn, Dock2, Cd3d, Ly9 |
| GO:0046651 | lymphocyte proliferation | 25 | 1,88E-10 | 2,84E-08 | 2,25E-08 | Ccl5, Cd3e, Card11, Cd74, Inpp5d, Ptprc, Zap70, Ptpn22, Xcl1, Rac2, Rasal3, H2-Ab1, Spn, Dock2, Coro1a, H2-Aa, Vsir, Sash3, Il7r, Fyn, Ikzf3, Itgb2, Nckap1l, Clec2i, Cd27 |
| GO:0002768 | immune response-regulating cell surface receptor signaling pathway | 26 | 2,08E-10 | 3,03E-08 | 2,39E-08 | Cd3e, Klrk1, Ptprc, Fyb, Zap70, Ptpn22, Cd247, Itk, Lax1, Nr4a3, Pde4b, Thy1, Skap1, Lck, Tlr2, Txk, Fyn, Lcp2, Lpxn, Trat1, Lat2, Nckap1l, Klre1, Myo1g, Fcer1g, Clec2i |
| GO:0050852 | T cell receptor signaling pathway | 16 | 3,38E-10 | 4,72E-08 | 3,73E-08 | Cd3e, Ptprc, Fyb, Zap70, Ptpn22, Cd247, Itk, Pde4b, Thy1, Skap1, Lck, Txk, Fyn, Lcp2, Trat1, Clec2i |
| GO:1902105 | regulation of leukocyte differentiation | 24 | 3,55E-10 | 4,78E-08 | 3,78E-08 | Ccl5, Card11, Cd74, Tbx21, H2-DMa, Inpp5d, Ptprc, Zap70, Dtx1, Egr3, Il2rg, H2-Aa, Vsir, Sash3, Gpr55, Lck, Fbn1, Il7r, Ikzf3, Hcls1, Nckap1l, Ikzf1, Clec2i, Cd27 |
| GO:0045619 | regulation of lymphocyte differentiation | 19 | 4,28E-10 | 5,54E-08 | 4,39E-08 | Card11, Cd74, Tbx21, H2-DMa, Inpp5d, Ptprc, Zap70, Dtx1, Egr3, Il2rg, H2-Aa, Vsir, Sash3, Lck, Il7r, Ikzf3, Nckap1l, Ikzf1, Cd27 |
| GO:0042098 | T cell proliferation | 20 | 5,68E-10 | 7,11E-08 | 5,62E-08 | Ccl5, Cd3e, Card11, Ptprc, Zap70, Ptpn22, Xcl1, Rac2, Rasal3, H2-Ab1, Spn, Dock2, Coro1a, H2-Aa, Vsir, Sash3, Fyn, Itgb2, Nckap1l, Clec2i |
| GO:0060326 | cell chemotaxis | 24 | 6,21E-10 | 7,32E-08 | 5,79E-08 | Chga, Nr4a1, Ccl5, Edn2, Vav1, Cd74, Thbs1, Klrk1, Xcl1, Rac2, Gpr18, Ccl4, Smoc2, Egr3, Ackr3, Edn1, Pde4b, Coro1a, Cxcr6, Itgb2, Nckap1l, Fcer1g, Pdgfd, Gpsm3 |
| GO:0032944 | regulation of mononuclear cell proliferation | 21 | 6,25E-10 | 7,32E-08 | 5,79E-08 | Ccl5, Cd3e, Card11, Cd74, Inpp5d, Ptprc, Zap70, Ptpn22, Xcl1, Rac2, Rasal3, H2-Ab1, Spn, Coro1a, H2-Aa, Vsir, Sash3, Ikzf3, St6gal1, Nckap1l, Clec2i |
| GO:0045621 | positive regulation of lymphocyte differentiation | 15 | 6,67E-10 | 7,56E-08 | 5,98E-08 | Cd74, H2-DMa, Inpp5d, Ptprc, Zap70, Egr3, Il2rg, H2-Aa, Vsir, Sash3, Lck, Il7r, Nckap1l, Ikzf1, Cd27 |
| GO:0002683 | negative regulation of immune system process | 30 | 6,93E-10 | 7,62E-08 | 6,03E-08 | Dll1, Cd74, Tbx21, Thbs1, Inpp5d, Klrk1, Ptprc, Gpr171, Ptpn22, Xcl1, Gpr18, Tbc1d10c, Dtx1, H2-Ab1, Cd96, Spn, Lax1, H2-Aa, Thy1, Serpinb9, Vsir, Gpr55, Fbn1, Il7r, Lpxn, Nlrc3, Cd300lf, Klre1, Fcer1g, Clec2i |
| GO:0002504 | antigen processing and presentation of peptide or polysaccharide antigen via MHC class II | 8 | 9,18E-10 | 9,80E-08 | 7,75E-08 | Cd74, H2-DMa, Thbs1, H2-DMb1, H2-Ab1, H2-Eb1, H2-Aa, Fcer1g |
| mmu05332 | Graft-versus-host disease | 13 | 2,24E-09 | 1,06E-07 | 8,17E-08 | 15043, 14998, 14999, 16643, 15024, 14939, 14103, 14961, 14963, 14969, 14960, 667803, 16638 |
| GO:0045580 | regulation of T cell differentiation | 17 | 1,03E-09 | 1,07E-07 | 8,49E-08 | Card11, Cd74, Tbx21, H2-DMa, Ptprc, Zap70, Dtx1, Egr3, Il2rg, H2-Aa, Vsir, Sash3, Lck, Il7r, Nckap1l, Ikzf1, Cd27 |

| | | | | | | |
|---|---|---|---|---|---|---|
| GO:0045582 | positive regulation of T cell differentiation | 14 | 1,10E-09 | 1,11E-07 | 8,76E-08 | Cd74, H2-DMa, Ptprc, Zap70, Egr3, Il2rg, H2-Aa, Vsir, Sash3, Lck, Il7r, Nckap1l, Ikzf1, Cd27 |
| GO:0070663 | regulation of leukocyte proliferation | 21 | 1,28E-09 | 1,26E-07 | 9,93E-08 | Ccl5, Cd3e, Card11, Cd74, Inpp5d, Ptprc, Zap70, Ptpn22, Xcl1, Rac2, Rasal3, H2-Ab1, Spn, Coro1a, H2-Aa, Vsir, Sash3, Ikzf3, St6gal1, Nckap1l, Clec2i |
| GO:0019886 | antigen processing and presentation of exogenous peptide antigen via MHC class II | 7 | 1,56E-09 | 1,49E-07 | 1,18E-07 | Cd74, H2-DMa, H2-DMb1, H2-Ab1, H2-Eb1, H2-Aa, Fcer1g |
| GO:0034341 | response to interferon-gamma | 16 | 2,01E-09 | 1,87E-07 | 1,48E-07 | Ccl5, Gbp8, Xcl1, Tgtp1, Ccl4, H2-Ab1, Gbp9, Gbp4, H2-Eb1, H2-Aa, Ciita, Tlr2, Txk, Was, Gbp6, Ubd |
| GO:0051251 | positive regulation of lymphocyte activation | 27 | 2,07E-09 | 1,88E-07 | 1,49E-07 | Ccl5, Cd3e, Card11, Cd74, Tbx21, H2-DMa, Inpp5d, Ptprc, Zap70, Ptpn22, Xcl1, Rasal3, H2-Ab1, Egr3, Spn, Il2rg, Coro1a, H2-Aa, Thy1, Vsir, Sash3, Lck, Il7r, Nckap1l, Ikzf1, Clec2i, Cd27 |
| GO:0002274 | myeloid leukocyte activation | 19 | 2,19E-09 | 1,94E-07 | 1,54E-07 | Chga, Ccl5, Edn2, Thbs1, Klrk1, Rac2, Dock2, Nr4a3, Rhoh, Tlr2, Cd48, Lcp2, Itgb2, Myo1f, Lat, Lat2, Cd300lf, Ubd, Fcer1g |
| GO:0050670 | regulation of lymphocyte proliferation | 20 | 3,22E-09 | 2,78E-07 | 2,20E-07 | Ccl5, Cd3e, Card11, Cd74, Inpp5d, Ptprc, Zap70, Ptpn22, Xcl1, Rac2, Rasal3, H2-Ab1, Spn, Coro1a, H2-Aa, Vsir, Sash3, Ikzf3, Nckap1l, Clec2i |
| GO:0002757 | immune response-activating signal transduction | 28 | 3,79E-09 | 3,20E-07 | 2,53E-07 | Cd3e, Klrk1, Ptprc, Fyb, Zap70, Ptpn22, Cd247, Itk, Lax1, Nr4a3, Pde4b, Thy1, Skap1, Lck, Tlr2, Txk, Fyn, Lcp2, Lpxn, Trat1, Lat2, Cd300lf, Nckap1l, Klre1, Myo1g, Peli3, Fcer1g, Clec2i |
| GO:0033077 | T cell differentiation in thymus | 13 | 4,81E-09 | 3,91E-07 | 3,09E-07 | Cd3g, Cd3e, Card11, Cd74, H2-DMa, Ptprc, Zap70, Egr3, Spn, Il2rg, Dock2, Il7r, Cd3d |
| GO:0019882 | antigen processing and presentation | 14 | 4,85E-09 | 3,91E-07 | 3,09E-07 | H2-T3, Cd74, H2-DMa, Thbs1, H2-DMb1, H2-T10, H2-Ab1, H2-Bl, H2-Eb1, H2-Aa, Was, Psmb8, Fcer1g, Rab3c |
| GO:1902107 | positive regulation of leukocyte differentiation | 17 | 5,11E-09 | 4,03E-07 | 3,19E-07 | Ccl5, Cd74, H2-DMa, Inpp5d, Ptprc, Zap70, Egr3, Il2rg, H2-Aa, Vsir, Sash3, Lck, Il7r, Hcls1, Nckap1l, Ikzf1, Cd27 |
| GO:0002253 | activation of immune response | 30 | 5,45E-09 | 4,21E-07 | 3,33E-07 | Cd3e, Klrk1, Ptprc, Fyb, Zap70, Ptpn22, Cd247, Itk, Lax1, Nr4a3, Pde4b, Thy1, Skap1, Lck, Tlr2, Txk, Fyn, Lcp2, Lpxn, Trat1, Lat2, Cd300lf, Nckap1l, Klre1, Myo1g, Peli3, Fcer1g, Tmem173, Clec2i, C9 |
| GO:0002449 | lymphocyte mediated immunity | 27 | 7,03E-09 | 5,31E-07 | 4,20E-07 | Vav1, H2-T3, Cd74, Tbx21, H2-DMa, Inpp5d, Klrk1, Klrd1, H2-T10, Gzmb, Ptprc, Xcl1, H2-Ab1, Cd96, H2-Bl, Spn, Coro1a, Serpinb9, Vsir, Sash3, Il7r, Was, Cd8a, Klre1, Myo1g, Fcer1g, C9 |
| GO:0002764 | immune response-regulating signaling pathway | 28 | 7,62E-09 | 5,65E-07 | 4,47E-07 | Cd3e, Klrk1, Ptprc, Fyb, Zap70, Ptpn22, Cd247, Itk, Lax1, Nr4a3, Pde4b, Thy1, Skap1, Lck, Tlr2, Txk, Fyn, Lcp2, Lpxn, Trat1, Lat2, Cd300lf, Nckap1l, Klre1, Myo1g, Peli3, Fcer1g, Clec2i |
| GO:0042129 | regulation of T cell proliferation | 17 | 8,09E-09 | 5,87E-07 | 4,65E-07 | Ccl5, Cd3e, Card11, Ptprc, Zap70, Ptpn22, Xcl1, Rac2, Rasal3, H2-Ab1, Spn, Coro1a, H2-Aa, Vsir, Sash3, Nckap1l, Clec2i |
| GO:1903706 | regulation of hemopoiesis | 26 | 1,12E-08 | 7,94E-07 | 6,28E-07 | Dll1, Ccl5, Card11, Cd74, Tbx21, H2-DMa, Inpp5d, Ptprc, Gpr171, Zap70, Dtx1, Egr3, Il2rg, H2-Aa, Vsir, Sash3, Gpr55, Lck, Fbn1, Il7r, Ikzf3, Hcls1, Nckap1l, Ikzf1, Clec2i, Cd27 |
| GO:0002456 | T cell mediated immunity | 15 | 1,28E-08 | 8,90E-07 | 7,04E-07 | H2-T3, Tbx21, H2-T10, Gzmb, Ptprc, Xcl1, H2-Bl, Spn, Serpinb9, Vsir, Sash3, Il7r, Was, Cd8a, Myo1g |
| GO:0002697 | regulation of immune effector process | 26 | 1,76E-08 | 1,21E-06 | 9,55E-07 | Ccl5, Vav1, H2-T3, Cd74, Tbx21, Klrk1, H2-T10, Ptprc, Ptpn22, Xcl1, Rac2, Cd96, H2-Bl, Spn, Gbp4, Nr4a3, Serpinb9, Vsir, Sash3, Tlr2, Il7r, Itgb2, Was, Klre1, Fcer1g, Tmem173 |
| GO:0002703 | regulation of leukocyte mediated immunity | 20 | 1,84E-08 | 1,24E-06 | 9,79E-07 | Vav1, H2-T3, Tbx21, Klrk1, H2-T10, Ptprc, Xcl1, Rac2, Cd96, H2-Bl, Spn, Serpinb9, Vsir, Sash3, Tlr2, Il7r, Itgb2, Was, Klre1, Fcer1g |
| GO:0002495 | antigen processing and presentation of peptide antigen via MHC class II | 7 | 2,12E-08 | 1,40E-06 | 1,11E-06 | Cd74, H2-DMa, H2-DMb1, H2-Ab1, H2-Eb1, H2-Aa, Fcer1g |
| GO:0002819 | regulation of adaptive immune response | 18 | 2,19E-08 | 1,42E-06 | 1,12E-06 | H2-T3, Cd74, Tbx21, H2-DMa, H2-T10, Ptprc, Xcl1, H2-Ab1, H2-Bl, Spn, Skap1, Vsir, Sash3, Cd48, Il7r, Adcy7, Was, Fcer1g |
| GO:0046635 | positive regulation of alpha-beta T cell activation | 11 | 2,39E-08 | 1,52E-06 | 1,20E-06 | Cd3e, Ptprc, Zap70, Ptpn22, Xcl1, Rasal3, H2-Ab1, Il2rg, Sash3, Nckap1l, Ikzf1 |
| GO:0032609 | interferon-gamma production | 14 | 2,78E-08 | 1,74E-06 | 1,38E-06 | Cd3e, Klrk1, Ptpn22, Xcl1, Itk, Cd96, Spn, Pde4b, Vsir, Sash3, Txk, Cd2, Klre1, Cd27 |
| GO:0046634 | regulation of alpha-beta T cell activation | 13 | 3,00E-08 | 1,85E-06 | 1,46E-06 | Cd3e, Tbx21, Ptprc, Zap70, Ptpn22, Xcl1, Rasal3, H2-Ab1, Il2rg, Vsir, Sash3, Nckap1l, Ikzf1 |
| mmu04940 | Type I diabetes mellitus | 12 | 5,88E-08 | 2,31E-06 | 1,79E-06 | 15043, 14998, 14999, 15024, 14939, 14103, 14961, 19275, 14963, 14969, 14960, 667803 |
| GO:0030595 | leukocyte chemotaxis | 18 | 4,00E-08 | 2,42E-06 | 1,91E-06 | Chga, Ccl5, Edn2, Vav1, Cd74, Thbs1, Klrk1, Xcl1, Rac2, Gpr18, Ccl4, Edn1, Pde4b, Coro1a, Itgb2, Nckap1l, Fcer1g, Gpsm3 |
| GO:0002685 | regulation of leukocyte migration | 17 | 4,35E-08 | 2,59E-06 | 2,05E-06 | Ccl5, Edn2, Cd74, Thbs1, Klrk1, Ptpn22, Xcl1, Rac2, Gpr18, Ccl4, Spn, Edn1, Thy1, Tlr2, Nckap1l, Pdgfd, Gpsm3 |
| GO:0001819 | positive regulation of cytokine production | 26 | 4,65E-08 | 2,68E-06 | 2,12E-06 | Ccl5, Cd3e, Card11, Cd74, Thbs1, Klrk1, Ptprc, Ptpn22, Xcl1, Ccl4, Spn, Nr4a3, Pde4b, Sash3, Tlr2, Txk, Cd2, Klre1, Agt, Fcer1g, Ltb, Ly9, Tmem173, Zbtb20, Gpsm3, Cd27 |
| GO:0019932 | second-messenger-mediated signaling | 26 | 4,65E-08 | 2,68E-06 | 2,12E-06 | Chga, Ntrk2, Edn2, Rasd1, Lgr5, Cd3e, Adgrg5, Thbs1, Pclo, Ptprc, Zap70, Neurod1, Tbc1d10c, Ccl4, Ackr3, Edn1, Sla2, Pde4b, Cxcr6, Glp1r, Lat, Trat1, Adcy7, Lat2, Cd8a, Pex5l |
| mmu05340 | Primary immunodeficiency | 9 | 9,10E-08 | 3,07E-06 | 2,37E-06 | 12501, 19264, 22637, 16186, 12265, 16818, 16197, 12525, 12500 |
| GO:0002706 | regulation of lymphocyte mediated immunity | 17 | 5,93E-08 | 3,37E-06 | 2,66E-06 | Vav1, H2-T3, Tbx21, Klrk1, H2-T10, Ptprc, Xcl1, Cd96, H2-Bl, Spn, Serpinb9, Vsir, Sash3, Il7r, Was, Klre1, Fcer1g |
| GO:0002707 | negative regulation of lymphocyte mediated immunity | 9 | 6,23E-08 | 3,48E-06 | 2,75E-06 | Tbx21, Ptprc, Xcl1, Cd96, Spn, Serpinb9, Vsir, Il7r, Klre1 |
| GO:0045577 | regulation of B cell differentiation | 8 | 6,37E-08 | 3,50E-06 | 2,77E-06 | Card11, Inpp5d, Ptprc, Il2rg, Ikzf3, Nckap1l, Ikzf1, Cd27 |
| GO:0045060 | negative thymic T cell selection | 6 | 7,75E-08 | 4,20E-06 | 3,32E-06 | Cd3e, Cd74, Ptprc, Zap70, Spn, Dock2 |

| | | | | | | |
|---|---|---|---|---|---|---|
| GO:0007264 | small GTPase mediated signal transduction | 27 | 8,48E-08 | 4,52E-06 | 3,58E-06 | Rasgrf2, Vav1, Rtn4r, Vav2, Rhobtb3, Farp2, P2ry10, Rac2, Gpr18, Fbp1, Arap3, Rasal3, Dock2, Rab37, Arhgap25, Rhoh, Gpr55, Dock10, Arhgap30, Arhgdib, Cyth4, Lat, Arhgef6, Sipa1, Was, Dock3, Rab3c |
| mmu04612 | Antigen processing and presentation | 13 | 1,54E-07 | 4,55E-06 | 3,51E-06 | 15043, 16149, 14998, 14999, 16643, 15024, 14961, 14963, 14969, 14960, 12265, 667803, 12525 |
| mmu05330 | Allograft rejection | 11 | 2,07E-07 | 5,44E-06 | 4,20E-06 | 15043, 14998, 14999, 15024, 14939, 14103, 14961, 14963, 14969, 14960, 667803 |
| GO:0043383 | negative T cell selection | 6 | 1,27E-07 | 6,69E-06 | 5,29E-06 | Cd3e, Cd74, Ptprc, Zap70, Spn, Dock2 |
| GO:0002478 | antigen processing and presentation of exogenous peptide antigen | 7 | 1,35E-07 | 7,00E-06 | 5,54E-06 | Cd74, H2-DMa, H2-DMb1, H2-Ab1, H2-Eb1, H2-Aa, Fcer1g |
| GO:0042102 | positive regulation of T cell proliferation | 12 | 1,43E-07 | 7,23E-06 | 5,72E-06 | Ccl5, Cd3e, Card11, Ptprc, Zap70, Ptpn22, Xcl1, Rasal3, Spn, Coro1a, Sash3, Nckap1l |
| GO:1903708 | positive regulation of hemopoiesis | 17 | 1,43E-07 | 7,23E-06 | 5,72E-06 | Ccl5, Cd74, H2-DMa, Inpp5d, Ptprc, Zap70, Egr3, Il2rg, H2-Aa, Vsir, Sash3, Lck, Il7r, Hcls1, Nckap1l, Ikzf1, Cd27 |
| GO:0002687 | positive regulation of leukocyte migration | 14 | 1,64E-07 | 8,06E-06 | 6,38E-06 | Ccl5, Edn2, Cd74, Thbs1, Xcl1, Rac2, Ccl4, Spn, Edn1, Thy1, Tlr2, Nckap1l, Pdgfd, Gpsm3 |
| GO:0032946 | positive regulation of mononuclear cell proliferation | 14 | 1,64E-07 | 8,06E-06 | 6,38E-06 | Ccl5, Cd3e, Card11, Cd74, Ptprc, Zap70, Ptpn22, Xcl1, Rasal3, Spn, Coro1a, Sash3, St6gal1, Nckap1l |
| GO:0030593 | neutrophil chemotaxis | 12 | 2,01E-07 | 9,72E-06 | 7,69E-06 | Ccl5, Edn2, Vav1, Cd74, Xcl1, Rac2, Ccl4, Edn1, Pde4b, Itgb2, Nckap1l, Fcer1g |
| GO:0006887 | exocytosis | 23 | 2,12E-07 | 1,01E-05 | 7,99E-06 | Chga, Cacna1a, Ccl5, Baiap3, Grik5, Hck, Pclo, Nrn1, Rac2, Nr4a3, Coro1a, Sept1, Itgb2, Cacna1h, Grm4, Myo1f, Lat, Lat2, Myo1g, Pex5l, Fcer1g, Syp, Rab3c |
| GO:0050851 | antigen receptor-mediated signaling pathway | 20 | 2,18E-07 | 1,03E-05 | 8,12E-06 | Cd3e, Ptprc, Fyb, Zap70, Ptpn22, Cd247, Itk, Lax1, Pde4b, Thy1, Skap1, Lck, Txk, Fyn, Lcp2, Lpxn, Trat1, Lat2, Nckap1l, Clec2i |
| GO:0032623 | interleukin-2 production | 10 | 2,29E-07 | 1,07E-05 | 8,43E-06 | Cd3e, Card11, Tbx21, Ptprc, Xcl1, Cd247, Pde4b, Sash3, Fcer1g, Clec2i |
| GO:0048002 | antigen processing and presentation of peptide antigen | 10 | 2,65E-07 | 1,22E-05 | 9,63E-06 | H2-T3, Cd74, H2-DMa, H2-DMb1, H2-T10, H2-Ab1, H2-Bl, H2-Eb1, H2-Aa, Fcer1g |
| GO:0070665 | positive regulation of leukocyte proliferation | 14 | 2,79E-07 | 1,26E-05 | 1,00E-05 | Ccl5, Cd3e, Card11, Cd74, Ptprc, Zap70, Ptpn22, Xcl1, Rasal3, Spn, Coro1a, Sash3, St6gal1, Nckap1l |
| GO:0071621 | granulocyte chemotaxis | 13 | 2,82E-07 | 1,26E-05 | 1,00E-05 | Ccl5, Edn2, Vav1, Cd74, Thbs1, Xcl1, Rac2, Ccl4, Edn1, Pde4b, Itgb2, Nckap1l, Fcer1g |
| GO:0045576 | mast cell activation | 10 | 3,06E-07 | 1,35E-05 | 1,07E-05 | Chga, Rac2, Nr4a3, Rhoh, Cd48, Lcp2, Lat, Lat2, Cd300lf, Fcer1g |
| GO:0046633 | alpha-beta T cell proliferation | 8 | 3,34E-07 | 1,46E-05 | 1,16E-05 | Cd3e, Ptprc, Zap70, Ptpn22, Xcl1, Rasal3, Dock2, Vsir |
| GO:0002704 | negative regulation of leukocyte mediated immunity | 9 | 4,00E-07 | 1,73E-05 | 1,37E-05 | Tbx21, Ptprc, Xcl1, Cd96, Spn, Serpinb9, Vsir, Il7r, Klre1 |
| mmu05416 | Viral myocarditis | 12 | 7,87E-07 | 1,85E-05 | 1,43E-05 | 15043, 14998, 14999, 15024, 19354, 14961, 14963, 14969, 14960, 667803, 14360, 16414 |
| mmu05321 | Inflammatory bowel disease (IBD) | 10 | 8,63E-07 | 1,85E-05 | 1,43E-05 | 57765, 14998, 14999, 14961, 60504, 14969, 16186, 14960, 20849, 24088 |
| GO:0002710 | negative regulation of T cell mediated immunity | 6 | 4,51E-07 | 1,93E-05 | 1,52E-05 | Tbx21, Ptprc, Xcl1, Spn, Vsir, Il7r |
| GO:0032649 | regulation of interferon-gamma production | 12 | 4,70E-07 | 1,98E-05 | 1,57E-05 | Cd3e, Klrk1, Ptpn22, Xcl1, Cd96, Pde4b, Vsir, Sash3, Txk, Cd2, Klre1, Cd27 |
| GO:0045055 | regulated exocytosis | 18 | 5,48E-07 | 2,28E-05 | 1,81E-05 | Chga, Cacna1a, Baiap3, Grik5, Pclo, Nrn1, Rac2, Nr4a3, Coro1a, Itgb2, Cacna1h, Grm4, Myo1f, Lat, Lat2, Pex5l, Fcer1g, Syp |
| GO:0009914 | hormone transport | 22 | 5,71E-07 | 2,34E-05 | 1,85E-05 | Cacna1e, Chga, Cacna1a, Ccl5, Baiap3, Edn2, Pde1c, Pclo, Pde8b, Neurod1, Hmgn3, Per2, Ptprn, Edn1, Slc16a2, Ptgs1, Scg5, Glp1r, Agt, Myt1, Pex5l, Myrip |
| GO:0002688 | regulation of leukocyte chemotaxis | 12 | 5,74E-07 | 2,34E-05 | 1,85E-05 | Ccl5, Edn2, Cd74, Thbs1, Klrk1, Xcl1, Rac2, Gpr18, Ccl4, Edn1, Nckap1l, Gpsm3 |
| GO:0002709 | regulation of T cell mediated immunity | 11 | 6,02E-07 | 2,43E-05 | 1,92E-05 | H2-T3, Tbx21, H2-T10, Ptprc, Xcl1, H2-Bl, Spn, Vsir, Sash3, Il7r, Was |
| GO:0019884 | antigen processing and presentation of exogenous antigen | 7 | 7,25E-07 | 2,89E-05 | 2,29E-05 | Cd74, H2-DMa, H2-DMb1, H2-Ab1, H2-Eb1, H2-Aa, Fcer1g |
| mmu04514 | Cell adhesion molecules (CAMs) | 16 | 1,80E-06 | 3,13E-05 | 2,42E-05 | 15043, 14998, 14999, 15024, 19264, 20345, 14961, 14963, 20737, 14969, 14960, 667803, 12481, 16421, 16414, 12525 |
| mmu04640 | Hematopoietic cell lineage | 12 | 1,82E-06 | 3,13E-05 | 2,42E-05 | 12502, 12501, 14998, 14999, 12516, 14961, 14969, 14960, 16197, 12481, 12525, 12500 |
| mmu05320 | Autoimmune thyroid disease | 11 | 1,86E-06 | 3,13E-05 | 2,42E-05 | 15043, 14998, 14999, 15024, 14939, 14103, 14961, 14963, 14969, 14960, 667803 |
| GO:0002366 | leukocyte activation involved in immune response | 18 | 8,66E-07 | 3,42E-05 | 2,70E-05 | Chga, Dll1, Apbb1ip, Tbx21, Ptprc, Rac2, Spn, Dock2, Nr4a3, Coro1a, Dock10, Itgb2, Myo1f, Lat, Lat2, Mfng, Fcer1g, Ly9 |
| GO:0050671 | positive regulation of lymphocyte proliferation | 13 | 9,04E-07 | 3,53E-05 | 2,79E-05 | Ccl5, Cd3e, Card11, Cd74, Ptprc, Zap70, Ptpn22, Xcl1, Rasal3, Spn, Coro1a, Sash3, Nckap1l |
| GO:0007204 | positive regulation of cytosolic calcium ion concentration | 20 | 9,98E-07 | 3,85E-05 | 3,05E-05 | Cacna1a, Edn2, Ptprc, Fasl, P2ry10, Xcl1, Gpr18, Ackr3, Edn1, Coro1a, Thy1, Gpr55, Lck, Cxcr6, Fyn, Glp1r, Fzd2, Ric3, Agt, Capn3 |
| GO:0002263 | cell activation involved in immune response | 18 | 1,08E-06 | 4,13E-05 | 3,27E-05 | Chga, Dll1, Apbb1ip, Tbx21, Ptprc, Rac2, Spn, Dock2, Nr4a3, Coro1a, Dock10, Itgb2, Myo1f, Lat, Lat2, Mfng, Fcer1g, Ly9 |
| GO:0050920 | regulation of chemotaxis | 16 | 1,11E-06 | 4,20E-05 | 3,32E-05 | Ccl5, Edn2, Cd74, Thbs1, Klrk1, Xcl1, Rac2, Gpr18, Ccl4, Smoc2, Edn1, St6gal1, Angpt2, Nckap1l, Pdgfd, Gpsm3 |
| GO:1990266 | neutrophil migration | 12 | 1,23E-06 | 4,59E-05 | 3,63E-05 | Ccl5, Edn2, Vav1, Cd74, Xcl1, Rac2, Ccl4, Edn1, Pde4b, Itgb2, Nckap1l, Fcer1g |
| GO:0046641 | positive regulation of alpha-beta T cell proliferation | 6 | 1,26E-06 | 4,67E-05 | 3,69E-05 | Cd3e, Ptprc, Zap70, Ptpn22, Xcl1, Rasal3 |
| mmu04664 | Fc epsilon RI signaling pathway | 10 | 3,33E-06 | 5,24E-05 | 4,04E-05 | 22324, 22325, 16331, 19354, 11689, 18707, 14360, 16822, 16797, 14127 |

| | | | | | | |
|---|---|---|---|---|---|---|
| GO:0046879 | hormone secretion | 21 | 1,45E-06 | 5,32E-05 | 4,21E-05 | Cacna1e, Chga, Cacna1a, Ccl5, Baiap3, Edn2, Pde1c, Pclo, Pde8b, Neurod1, Hmgn3, Per2, Ptprn, Edn1, Ptgs1, Scg5, Glp1r, Agt, Myt1, Pex5l, Myrip |
| GO:0014065 | phosphatidylinositol 3-kinase signaling | 12 | 1,47E-06 | 5,33E-05 | 4,22E-05 | Ccl5, Ntrk2, Pik3r5, Edn1, Pik3cd, Fyn, Hcls1, Agap2, Nlrc3, Agt, Hcst, Pdgfd |
| GO:0001818 | negative regulation of cytokine production | 17 | 1,59E-06 | 5,70E-05 | 4,51E-05 | Dll1, Tbx21, Thbs1, Inpp5d, Ptprc, Ptpn22, Xcl1, Gpr18, Cd96, Gbp4, Srgn, Vsir, Tlr2, Nlrc3, Adcy7, Nckap1l, Peli3 |
| GO:0007266 | Rho protein signal transduction | 15 | 1,65E-06 | 5,86E-05 | 4,63E-05 | Rasgrf2, Vav1, Rtn4r, Vav2, Rhobtb3, Farp2, P2ry10, Rac2, Gpr18, Arap3, Rhoh, Gpr55, Arhgdib, Arhgef6, Was |
| GO:0030183 | B cell differentiation | 13 | 1,73E-06 | 6,10E-05 | 4,82E-05 | Dll1, Card11, Inpp5d, Ptprc, H2-Ab1, Il2rg, Dock10, Ikzf3, Nckap1l, Ikzf1, Mfng, Gimap1, Cd27 |
| GO:0097530 | granulocyte migration | 13 | 1,87E-06 | 6,53E-05 | 5,16E-05 | Ccl5, Edn2, Vav1, Cd74, Thbs1, Xcl1, Rac2, Ccl4, Edn1, Pde4b, Itgb2, Nckap1l, Fcer1g |
| GO:0042492 | gamma-delta T cell differentiation | 5 | 1,91E-06 | 6,60E-05 | 5,22E-05 | Ptprc, Gpr18, Egr3, Lck, Nckap1l |
| GO:0032680 | regulation of tumor necrosis factor production | 13 | 2,18E-06 | 7,47E-05 | 5,91E-05 | Thbs1, Ptprc, Ptpn22, Gpr18, Ccl4, Spn, Vsir, Sash3, Tlr2, Cd2, Nlrc3, Fcer1g, Zbtb20 |
| GO:0051480 | regulation of cytosolic calcium ion concentration | 21 | 2,36E-06 | 8,01E-05 | 6,33E-05 | Chga, Cacna1a, Edn2, Ptprc, Fasl, P2ry10, Xcl1, Gpr18, Ackr3, Edn1, Coro1a, Thy1, Gpr55, Lck, Cxcr6, Fyn, Glp1r, Fzd2, Ric3, Agt, Capn3 |
| GO:0034113 | heterotypic cell-cell adhesion | 8 | 2,46E-06 | 8,23E-05 | 6,51E-05 | Ptprc, Thy1, Skap1, Lck, Cd2, Itgb7, Itgb2, Itgax |
| GO:0051056 | regulation of small GTPase mediated signal transduction | 17 | 2,47E-06 | 8,23E-05 | 6,51E-05 | Rasgrf2, Vav1, Rtn4r, Vav2, Farp2, P2ry10, Gpr18, Fbp1, Arap3, Rasal3, Dock2, Arhgap25, Gpr55, Arhgdib, Cyth4, Arhgef6, Sipa1 |
| GO:0002690 | positive regulation of leukocyte chemotaxis | 10 | 2,52E-06 | 8,23E-05 | 6,51E-05 | Ccl5, Edn2, Cd74, Thbs1, Xcl1, Rac2, Ccl4, Edn1, Nckap1l, Gpsm3 |
| GO:0032640 | tumor necrosis factor production | 13 | 2,54E-06 | 8,23E-05 | 6,51E-05 | Thbs1, Ptprc, Ptpn22, Gpr18, Ccl4, Spn, Vsir, Sash3, Tlr2, Cd2, Nlrc3, Fcer1g, Zbtb20 |
| GO:1903555 | regulation of tumor necrosis factor superfamily cytokine production | 13 | 2,54E-06 | 8,23E-05 | 6,51E-05 | Thbs1, Ptprc, Ptpn22, Gpr18, Ccl4, Spn, Vsir, Sash3, Tlr2, Cd2, Nlrc3, Fcer1g, Zbtb20 |
| GO:0097529 | myeloid leukocyte migration | 15 | 2,58E-06 | 8,28E-05 | 6,55E-05 | Chga, Ccl5, Edn2, Vav1, Cd74, Thbs1, Xcl1, Rac2, Ccl4, Edn1, Pde4b, Itgb2, Nckap1l, Fcer1g, Pdgfd |
| GO:0043299 | leukocyte degranulation | 9 | 2,63E-06 | 8,36E-05 | 6,61E-05 | Chga, Rac2, Nr4a3, Coro1a, Itgb2, Myo1f, Lat, Lat2, Fcer1g |
| GO:0046640 | regulation of alpha-beta T cell proliferation | 7 | 2,71E-06 | 8,56E-05 | 6,77E-05 | Cd3e, Ptprc, Zap70, Ptpn22, Xcl1, Rasal3, Vsir |
| GO:0007265 | Ras protein signal transduction | 22 | 2,96E-06 | 9,26E-05 | 7,33E-05 | Rasgrf2, Vav1, Rtn4r, Vav2, Rhobtb3, Farp2, P2ry10, Rac2, Gpr18, Fbp1, Arap3, Rasal3, Dock2, Rab37, Rhoh, Gpr55, Arhgdib, Cyth4, Lat, Arhgef6, Was, Rab3c |
| GO:0071706 | tumor necrosis factor superfamily cytokine production | 13 | 3,17E-06 | 9,84E-05 | 7,78E-05 | Thbs1, Ptprc, Ptpn22, Gpr18, Ccl4, Spn, Vsir, Sash3, Tlr2, Cd2, Nlrc3, Fcer1g, Zbtb20 |
| GO:0002699 | positive regulation of immune effector process | 17 | 3,41E-06 | 0,00010478 | 8,29E-05 | Vav1, H2-T3, Cd74, Tbx21, Klrk1, H2-T10, Ptprc, Ptpn22, Xcl1, Rac2, H2-Bl, Nr4a3, Sash3, Tlr2, Itgb2, Klre1, Fcer1g |
| GO:2000106 | regulation of leukocyte apoptotic process | 11 | 3,68E-06 | 0,00011218 | 8,87E-05 | Ccl5, Cd3g, Cd74, Nr4a3, Serpinb9, Pik3cd, Il7r, St6gal1, Hcls1, Fcer1g, Cd27 |
| GO:0071887 | leukocyte apoptotic process | 12 | 3,72E-06 | 0,00011237 | 8,89E-05 | Ccl5, Cd3g, Cd74, Fasl, Nr4a3, Serpinb9, Pik3cd, Il7r, St6gal1, Hcls1, Fcer1g, Cd27 |
| GO:0046632 | alpha-beta T cell differentiation | 11 | 4,40E-06 | 0,00013185 | 0,00010429 | Tbx21, Zap70, Itk, Gpr18, Spn, Il2rg, Sash3, Txk, Nckap1l, Ikzf1, Ly9 |
| GO:0071346 | cellular response to interferon-gamma | 11 | 4,80E-06 | 0,00014275 | 0,00011291 | Ccl5, Gbp8, Xcl1, Ccl4, H2-Ab1, Gbp9, Gbp4, Tlr2, Txk, Was, Gbp6 |
| GO:0019722 | calcium-mediated signaling | 15 | 5,01E-06 | 0,0001479 | 0,00011699 | Ntrk2, Edn2, Cd3e, Ptprc, Zap70, Tbc1d10c, Ccl4, Ackr3, Edn1, Sla2, Cxcr6, Lat, Trat1, Lat2, Cd8a |
| GO:0050921 | positive regulation of chemotaxis | 12 | 5,49E-06 | 0,00015979 | 0,0001264 | Ccl5, Edn2, Cd74, Thbs1, Xcl1, Rac2, Ccl4, Smoc2, Edn1, Nckap1l, Pdgfd, Gpsm3 |
| GO:0046883 | regulation of hormone secretion | 18 | 5,50E-06 | 0,00015979 | 0,0001264 | Cacna1e, Chga, Cacna1a, Ccl5, Baiap3, Edn2, Pde1c, Pde8b, Hmgn3, Per2, Edn1, Ptgs1, Scg5, Glp1r, Agt, Myt1, Pex5l, Myrip |
| GO:0046629 | gamma-delta T cell activation | 5 | 6,21E-06 | 0,00017878 | 0,00014141 | Ptprc, Gpr18, Egr3, Lck, Nckap1l |
| GO:0002367 | cytokine production involved in immune response | 10 | 6,33E-06 | 0,00018094 | 0,00014312 | Chga, Cd74, Tbx21, Xcl1, Cd96, Nr4a3, Vsir, Sash3, Tlr2, Fcer1g |
| GO:0002822 | regulation of adaptive immune response based on somatic recombination of immune receptors built from immunoglobulin superfamily domains | 14 | 6,77E-06 | 0,00019171 | 0,00015164 | H2-T3, Tbx21, H2-DMa, H2-T10, Ptprc, Xcl1, H2-Ab1, H2-Bl, Spn, Vsir, Sash3, Il7r, Was, Fcer1g |
| GO:0002718 | regulation of cytokine production involved in immune response | 9 | 6,81E-06 | 0,00019171 | 0,00015164 | Cd74, Tbx21, Xcl1, Cd96, Nr4a3, Vsir, Sash3, Tlr2, Fcer1g |
| GO:0001909 | leukocyte mediated cytotoxicity | 12 | 7,40E-06 | 0,00020669 | 0,00016349 | Vav1, H2-T3, Klrk1, H2-T10, Gzmb, Ptprc, Xcl1, H2-Bl, Coro1a, Serpinb9, Il7r, Klre1 |
| GO:0032729 | positive regulation of interferon-gamma production | 9 | 7,61E-06 | 0,00021084 | 0,00016677 | Cd3e, Klrk1, Ptpn22, Pde4b, Sash3, Txk, Cd2, Klre1, Cd27 |
| GO:0032663 | regulation of interleukin-2 production | 8 | 7,77E-06 | 0,00021368 | 0,00016902 | Cd3e, Card11, Tbx21, Ptprc, Xcl1, Pde4b, Sash3, Clec2i |
| mmu05166 | Human T-cell leukemia virus 1 infection | 18 | 1,58E-05 | 0,00021997 | 0,00016973 | 12502, 12501, 15043, 14998, 14999, 15024, 14961, 14963, 14969, 16186, 14960, 16185, 16818, 18707, 667803, 16414, 11513, 12500 |
| mmu04062 | Chemokine signaling pathway | 16 | 1,58E-05 | 0,00021997 | 0,00016973 | 20304, 22324, 22325, 18796, 15162, 16963, 19354, 320207, 16428, 20303, 14710, 94176, 80901, 18707, 11513, 22376 |
| GO:0006874 | cellular calcium ion homeostasis | 23 | 8,52E-06 | 0,00023237 | 0,0001838 | Chga, Cacna1a, Ccl5, Edn2, Ptprc, Fasl, P2ry10, Xcl1, Gpr18, Ackr3, Edn1, Coro1a, Thy1, Gpr55, Lck, Cxcr6, Fyn, Glp1r, Fzd2, Ric3, Agt, Capn3, Slc24a3 |
| GO:0050866 | negative regulation of cell activation | 14 | 8,62E-06 | 0,00023297 | 0,00018428 | Adamts18, Cd74, Tbx21, Inpp5d, Ptpn22, Xcl1, Tbc1d10c, Dtx1, H2-Ab1, Spn, Lax1, H2-Aa, Vsir, Cd300lf |
| GO:0045579 | positive regulation of B cell differentiation | 5 | 8,67E-06 | 0,00023297 | 0,00018428 | Inpp5d, Il2rg, Nckap1l, Ikzf1, Cd27 |
| GO:0046578 | regulation of Ras protein signal transduction | 15 | 8,82E-06 | 0,00023547 | 0,00018626 | Rasgrf2, Vav1, Rtn4r, Vav2, Farp2, P2ry10, Gpr18, Fbp1, Arap3, Rasal3, Dock2, Gpr55, Arhgdib, Cyth4, Arhgef6 |
| mmu05310 | Asthma | 6 | 1,80E-05 | 0,00023591 | 0,00018203 | 14998, 14999, 14961, 14969, 14960, 14127 |

| | | | | | | |
|---|---|---|---|---|---|---|
| GO:0007162 | negative regulation of cell adhesion | 17 | 9,25E-06 | 0,00024495 | 0,00019375 | Adamts18, Cd74, Tbx21, Thbs1, Ptprc, Ptpn22, Xcl1, Dtx1, H2-Ab1, Spn, Lax1, H2-Aa, Vsir, Lpxn, Myo1f, Angpt2, Spaca7 |
| GO:0042113 | B cell activation | 21 | 9,74E-06 | 0,00025602 | 0,00020251 | Dll1, Card11, Cd74, Tbx21, Inpp5d, Ptprc, Tbc1d10c, H2-Ab1, Lax1, Il2rg, Sash3, Pik3cd, Dock10, Il7r, Ikzf3, Lat2, Nckap1l, Ikzf1, Mfng, Gimap1, Cd27 |
| GO:0048015 | phosphatidylinositol-mediated signaling | 12 | 9,88E-06 | 0,000258 | 0,00020408 | Ccl5, Ntrk2, Pik3r5, Edn1, Pik3cd, Fyn, Hcls1, Agap2, Nlrc3, Agt, Hcst, Pdgfd |
| GO:0002460 | adaptive immune response based on somatic recombination of immune receptors built from immunoglobulin superfamily domains | 22 | 1,00E-05 | 0,00026045 | 0,00020602 | H2-T3, Cd74, Tbx21, H2-DMa, Inpp5d, H2-T10, Gzmb, Ptprc, Xcl1, H2-Ab1, H2-Bl, Spn, Serpinb9, Vsir, Sash3, Il7r, Was, Cd8a, Myo1g, Fcer1g, Ly9, C9 |
| GO:0033627 | cell adhesion mediated by integrin | 8 | 1,14E-05 | 0,00029349 | 0,00023215 | Ccl5, Cd3e, Skap1, Fbn1, Itgb7, Itgb2, Lpxn, Nckap1l |
| GO:0001906 | cell killing | 13 | 1,15E-05 | 0,00029451 | 0,00023296 | Vav1, H2-T3, Klrk1, H2-T10, Gzmb, Ptprc, Xcl1, H2-Bl, Coro1a, Serpinb9, Il7r, Klre1, C9 |
| GO:0042035 | regulation of cytokine biosynthetic process | 10 | 1,21E-05 | 0,00030291 | 0,0002396 | Cd3e, Card11, Thbs1, Inpp5d, Ptprc, Spn, Tlr2, Ltb, Clec2i, Cd27 |
| GO:0048017 | inositol lipid-mediated signaling | 12 | 1,22E-05 | 0,00030291 | 0,0002396 | Ccl5, Ntrk2, Pik3r5, Edn1, Pik3cd, Fyn, Hcls1, Agap2, Nlrc3, Agt, Hcst, Pdgfd |
| GO:0051250 | negative regulation of lymphocyte activation | 12 | 1,22E-05 | 0,00030291 | 0,0002396 | Cd74, Tbx21, Inpp5d, Ptpn22, Xcl1, Tbc1d10c, Dtx1, H2-Ab1, Spn, Lax1, H2-Aa, Vsir |
| GO:0006816 | calcium ion transport | 21 | 1,22E-05 | 0,00030291 | 0,0002396 | Cacna1e, Cacna1a, Ccl5, Ptprc, Fasl, Ptpn22, Xcl1, Ccl4, Pde4b, Coro1a, Thy1, Lck, Fyn, Cacna1h, Glp1r, Oprd1, Agt, Usp2, Capn3, Slc24a3, Hpca |
| GO:0002695 | negative regulation of leukocyte activation | 13 | 1,23E-05 | 0,00030291 | 0,0002396 | Cd74, Tbx21, Inpp5d, Ptpn22, Xcl1, Tbc1d10c, Dtx1, H2-Ab1, Spn, Lax1, H2-Aa, Vsir, Cd300lf |
| GO:0002275 | myeloid cell activation involved in immune response | 9 | 1,29E-05 | 0,00031624 | 0,00025015 | Chga, Rac2, Dock2, Nr4a3, Itgb2, Myo1f, Lat, Lat2, Fcer1g |
| GO:0002821 | positive regulation of adaptive immune response | 12 | 1,31E-05 | 0,00031805 | 0,00025158 | H2-T3, Cd74, Tbx21, H2-DMa, H2-T10, Ptprc, Xcl1, H2-Ab1, H2-Bl, Skap1, Sash3, Fcer1g |
| mmu04015 | Rap1 signaling pathway | 16 | 2,74E-05 | 0,00034023 | 0,00026253 | 54519, 18796, 21825, 227377, 23880, 19354, 106952, 78473, 18707, 16822, 16414, 16797, 11513, 20469, 11601, 71785 |
| mmu04666 | Fc gamma R-mediated phagocytosis | 10 | 3,12E-05 | 0,0003681 | 0,00028404 | 22324, 22325, 15162, 16331, 19264, 19354, 94176, 18707, 16797, 22376 |
| GO:0055074 | calcium ion homeostasis | 23 | 1,52E-05 | 0,00036833 | 0,00029135 | Chga, Cacna1a, Ccl5, Edn2, Ptprc, Fasl, P2ry10, Xcl1, Gpr18, Ackr3, Edn1, Coro1a, Thy1, Gpr55, Lck, Cxcr6, Fyn, Glp1r, Fzd2, Ric3, Agt, Capn3, Slc24a3 |
| GO:0042108 | positive regulation of cytokine biosynthetic process | 8 | 1,84E-05 | 0,00044174 | 0,00034941 | Cd3e, Card11, Thbs1, Ptprc, Spn, Tlr2, Ltb, Cd27 |
| GO:0031589 | cell-substrate adhesion | 18 | 1,89E-05 | 0,00045203 | 0,00035755 | Hoxd3, Cd3e, Limch1, Thbs1, Sned1, Rac2, Smoc2, Cd96, Coro1a, Thy1, Skap1, Itgb7, St6gal1, Itgb2, Angpt2, Agt, Myo1g, Smoc1 |
| GO:0072503 | cellular divalent inorganic cation homeostasis | 23 | 1,92E-05 | 0,00045458 | 0,00035957 | Chga, Cacna1a, Ccl5, Edn2, Ptprc, Fasl, P2ry10, Xcl1, Gpr18, Ackr3, Edn1, Coro1a, Thy1, Gpr55, Lck, Cxcr6, Fyn, Glp1r, Fzd2, Ric3, Agt, Capn3, Slc24a3 |
| GO:0014068 | positive regulation of phosphatidylinositol 3-kinase signaling | 8 | 2,06E-05 | 0,00048537 | 0,00038392 | Ccl5, Ntrk2, Fyn, Hcls1, Agap2, Agt, Hcst, Pdgfd |
| GO:0034765 | regulation of ion transmembrane transport | 22 | 2,09E-05 | 0,00048898 | 0,00038679 | Cacna1e, Rasgrf2, Cacna1a, Kcnh8, Trpm5, Wnk2, Thbs1, Ptpn22, Xcl1, Per2, Hvcn1, Pde4b, Coro1a, Thy1, Fyn, Fxyd5, Cacna1h, Agt, Kcnab2, Capn3, Hpca, Scn3a |
| GO:0007229 | integrin-mediated signaling pathway | 9 | 2,10E-05 | 0,00048941 | 0,00038712 | Vav1, Fyb, Itgae, Thy1, Itgb7, Itgb2, Lat, Itgax, Fcer1g |
| GO:0035023 | regulation of Rho protein signal transduction | 11 | 2,15E-05 | 0,00049748 | 0,0003935 | Rasgrf2, Vav1, Rtn4r, Vav2, Farp2, P2ry10, Gpr18, Arap3, Gpr55, Arhgdib, Arhgef6 |
| GO:0001910 | regulation of leukocyte mediated cytotoxicity | 10 | 2,18E-05 | 0,00050054 | 0,00039593 | Vav1, H2-T3, Klrk1, H2-T10, Ptprc, Xcl1, H2-Bl, Serpinb9, Il7r, Klre1 |
| GO:0071622 | regulation of granulocyte chemotaxis | 7 | 2,29E-05 | 0,00052243 | 0,00041324 | Ccl5, Cd74, Thbs1, Xcl1, Rac2, Edn1, Nckap1l |
| GO:0002228 | natural killer cell mediated immunity | 8 | 2,30E-05 | 0,00052243 | 0,00041324 | Vav1, Klrk1, Klrd1, Gzmb, Cd96, Coro1a, Serpinb9, Klre1 |
| GO:0030072 | peptide hormone secretion | 16 | 2,37E-05 | 0,00053326 | 0,00042181 | Cacna1e, Chga, Ccl5, Baiap3, Pde1c, Pclo, Pde8b, Neurod1, Hmgn3, Per2, Ptprn, Edn1, Glp1r, Myt1, Pex5l, Myrip |
| GO:0070588 | calcium ion transmembrane transport | 16 | 2,48E-05 | 0,00055469 | 0,00043875 | Cacna1e, Cacna1a, Ptprc, Fasl, Ptpn22, Xcl1, Pde4b, Coro1a, Thy1, Lck, Fyn, Cacna1h, Glp1r, Capn3, Slc24a3, Hpca |
| GO:0032760 | positive regulation of tumor necrosis factor production | 9 | 2,53E-05 | 0,0005642 | 0,00044628 | Thbs1, Ptprc, Ccl4, Spn, Sash3, Tlr2, Cd2, Fcer1g, Zbtb20 |
| GO:1903557 | positive regulation of tumor necrosis factor superfamily cytokine production | 9 | 2,78E-05 | 0,00061426 | 0,00048588 | Thbs1, Ptprc, Ccl4, Spn, Sash3, Tlr2, Cd2, Fcer1g, Zbtb20 |
| GO:2000107 | negative regulation of leukocyte apoptotic process | 8 | 2,86E-05 | 0,00062982 | 0,00049819 | Ccl5, Cd74, Serpinb9, Il7r, St6gal1, Hcls1, Fcer1g, Cd27 |
| GO:0014066 | regulation of phosphatidylinositol 3-kinase signaling | 9 | 3,04E-05 | 0,00066393 | 0,00052516 | Ccl5, Ntrk2, Fyn, Hcls1, Agap2, Nlrc3, Agt, Hcst, Pdgfd |
| GO:0042089 | cytokine biosynthetic process | 10 | 3,23E-05 | 0,00069893 | 0,00055285 | Cd3e, Card11, Thbs1, Inpp5d, Ptprc, Spn, Tlr2, Ltb, Clec2i, Cd27 |
| GO:0030073 | insulin secretion | 14 | 3,24E-05 | 0,00069893 | 0,00055285 | Cacna1e, Chga, Ccl5, Baiap3, Pde1c, Pclo, Pde8b, Neurod1, Hmgn3, Per2, Ptprn, Glp1r, Myt1, Myrip |
| GO:0018108 | peptidyl-tyrosine phosphorylation | 17 | 3,93E-05 | 0,00084358 | 0,00066727 | Ccl5, Cd3e, Cd74, Hck, Ptprc, Zap70, Ptpn22, Itk, Thy1, Sla, Lck, Txk, Fyn, Hcls1, Itgb2, Agt, Pdgfd |
| GO:0050868 | negative regulation of T cell activation | 10 | 4,05E-05 | 0,00086502 | 0,00068423 | Cd74, Tbx21, Ptpn22, Xcl1, Dtx1, H2-Ab1, Spn, Lax1, H2-Aa, Vsir |
| GO:0002700 | regulation of production of molecular mediator of immune response | 11 | 4,26E-05 | 0,00090488 | 0,00071576 | Cd74, Tbx21, Ptprc, Ptpn22, Xcl1, Cd96, Nr4a3, Vsir, Sash3, Tlr2, Fcer1g |
| GO:0042107 | cytokine metabolic process | 10 | 4,36E-05 | 0,00092041 | 0,00072804 | Cd3e, Card11, Thbs1, Inpp5d, Ptprc, Spn, Tlr2, Ltb, Clec2i, Cd27 |
| GO:0018212 | peptidyl-tyrosine modification | 17 | 4,43E-05 | 0,00092976 | 0,00073544 | Ccl5, Cd3e, Cd74, Hck, Ptprc, Zap70, Ptpn22, Itk, Thy1, Sla, Lck, Txk, Fyn, Hcls1, Itgb2, Agt, Pdgfd |

| | | | | | | |
|---|---|---|---|---|---|---|
| mmu05169 | Epstein-Barr virus infection | 16 | 8,71E-05 | 0,00097937 | 0,00075572 | 12502, 12501, 15043, 14998, 14999, 15024, 12503, 14961, 14963, 14969, 14960, 18707, 24088, 667803, 18037, 12500 |
| GO:0035710 | CD4-positive, alpha-beta T cell activation | 9 | 5,08E-05 | 0,0010597 | 0,00083822 | Tbx21, Xcl1, Spn, Il2rg, Vsir, Sash3, Nckap1l, Ly9, Cdh26 |
| GO:0001516 | prostaglandin biosynthetic process | 5 | 5,38E-05 | 0,00110234 | 0,00087195 | Edn2, Hpgds, Cd74, Edn1, Ptgs1 |
| GO:0046457 | prostanoid biosynthetic process | 5 | 5,38E-05 | 0,00110234 | 0,00087195 | Edn2, Hpgds, Cd74, Edn1, Ptgs1 |
| GO:0090023 | positive regulation of neutrophil chemotaxis | 5 | 5,38E-05 | 0,00110234 | 0,00087195 | Cd74, Xcl1, Rac2, Edn1, Nckap1l |
| GO:0045088 | regulation of innate immune response | 15 | 5,47E-05 | 0,00111517 | 0,00088209 | Vav1, Cd74, Klrk1, Ptpn22, Xcl1, Cd96, Serpinb9, Tlr2, Txk, Myo1f, Nlrc3, Cd300lf, Klre1, Peli3, Tmem173 |
| GO:0002708 | positive regulation of lymphocyte mediated immunity | 11 | 5,51E-05 | 0,00111699 | 0,00088353 | Vav1, H2-T3, Tbx21, Klrk1, H2-T10, Ptprc, Xcl1, H2-Bl, Sash3, Klre1, Fcer1g |
| GO:0002823 | negative regulation of adaptive immune response based on somatic recombination of immune receptors built from immunoglobulin superfamily domains | 6 | 5,80E-05 | 0,00116836 | 0,00092417 | Tbx21, Ptprc, Xcl1, Spn, Vsir, Il7r |
| GO:0050777 | negative regulation of immune response | 11 | 5,87E-05 | 0,00117607 | 0,00093027 | Tbx21, Inpp5d, Ptprc, Xcl1, Cd96, Spn, Serpinb9, Vsir, Il7r, Nlrc3, Klre1 |
| GO:0002702 | positive regulation of production of molecular mediator of immune response | 9 | 5,98E-05 | 0,0011922 | 0,00094303 | Cd74, Tbx21, Ptprc, Ptpn22, Xcl1, Nr4a3, Sash3, Tlr2, Fcer1g |
| GO:0002431 | Fc receptor mediated stimulatory signaling pathway | 4 | 6,02E-05 | 0,00119329 | 0,00094389 | Ptprc, Nr4a3, Myo1g, Fcer1g |
| GO:0070838 | divalent metal ion transport | 21 | 6,07E-05 | 0,00119748 | 0,0009472 | Cacna1e, Cacna1a, Ccl5, Ptprc, Fasl, Ptpn22, Xcl1, Ccl4, Pde4b, Coro1a, Thy1, Lck, Fyn, Cacna1h, Glp1r, Oprd1, Agt, Usp2, Capn3, Slc24a3, Hpca |
| GO:0045123 | cellular extravasation | 7 | 6,25E-05 | 0,00122606 | 0,00096981 | Ccl5, Selplg, Spn, Thy1, Itgb7, Itgb2, Pdgfd |
| GO:0001913 | T cell mediated cytotoxicity | 8 | 6,37E-05 | 0,00124379 | 0,00098383 | H2-T3, H2-T10, Gzmb, Ptprc, Xcl1, H2-Bl, Serpinb9, Il7r |
| GO:0072511 | divalent inorganic cation transport | 21 | 6,67E-05 | 0,00128976 | 0,0010202 | Cacna1e, Cacna1a, Ccl5, Ptprc, Fasl, Ptpn22, Xcl1, Ccl4, Pde4b, Coro1a, Thy1, Lck, Fyn, Cacna1h, Glp1r, Oprd1, Agt, Usp2, Capn3, Slc24a3, Hpca |
| GO:0031341 | regulation of cell killing | 10 | 6,68E-05 | 0,00128976 | 0,0010202 | Vav1, H2-T3, Klrk1, H2-T10, Ptprc, Xcl1, H2-Bl, Serpinb9, Il7r, Klre1 |
| GO:0046456 | icosanoid biosynthetic process | 6 | 6,72E-05 | 0,00128976 | 0,0010202 | Edn2, Hpgds, Cd74, Edn1, Alox5, Ptgs1 |
| GO:0051924 | regulation of calcium ion transport | 15 | 6,81E-05 | 0,00130018 | 0,00102844 | Cacna1a, Ccl5, Ptpn22, Xcl1, Ccl4, Pde4b, Coro1a, Thy1, Fyn, Glp1r, Oprd1, Agt, Usp2, Capn3, Hpca |
| GO:0002705 | positive regulation of leukocyte mediated immunity | 12 | 7,24E-05 | 0,00136779 | 0,00108192 | Vav1, H2-T3, Tbx21, Klrk1, H2-T10, Ptprc, Xcl1, H2-Bl, Sash3, Itgb2, Klre1, Fcer1g |
| GO:0009749 | response to glucose | 12 | 7,24E-05 | 0,00136779 | 0,00108192 | Pck1, Cacna1e, Baiap3, Pde1c, Thbs1, Pde8b, Neurod1, Hmgn3, Ptprn, Glp1r, Myt1, Zbtb20 |
| mmu05142 | Chagas disease (American trypanosomiasis) | 10 | 0,00013323 | 0,00139263 | 0,0010746 | 20304, 12502, 12501, 18796, 14103, 269643, 12503, 18707, 24088, 12500 |
| mmu05323 | Rheumatoid arthritis | 9 | 0,00013572 | 0,00139263 | 0,0010746 | 20304, 14998, 14999, 14961, 14969, 14960, 24088, 16414, 16994 |
| GO:0038083 | peptidyl-tyrosine autophosphorylation | 6 | 7,75E-05 | 0,00145772 | 0,00115305 | Hck, Itk, Sla, Lck, Txk, Fyn |
| GO:0071624 | positive regulation of granulocyte chemotaxis | 5 | 8,08E-05 | 0,00151197 | 0,00119596 | Cd74, Xcl1, Rac2, Edn1, Nckap1l |
| GO:0030101 | natural killer cell activation | 9 | 8,18E-05 | 0,00152243 | 0,00120423 | Klrk1, Ptprc, Ptpn22, Slamf7, Coro1a, Il2rb, Itgb2, Klre1, Ikzf1 |
| GO:0009746 | response to hexose | 12 | 8,51E-05 | 0,00157576 | 0,00124642 | Pck1, Cacna1e, Baiap3, Pde1c, Thbs1, Pde8b, Neurod1, Hmgn3, Ptprn, Glp1r, Myt1, Zbtb20 |
| GO:0019221 | cytokine-mediated signaling pathway | 17 | 8,84E-05 | 0,00161694 | 0,00127899 | Prlr, Ccl5, Edn2, Cd74, Ptprc, Xcl1, Ccl4, Ptprn, Ackr3, Stat4, Il2rb, Txk, Iigp1, Cd300lf, Peli3, Fcer1g, St18 |
| GO:0002820 | negative regulation of adaptive immune response | 6 | 8,91E-05 | 0,00161694 | 0,00127899 | Tbx21, Ptprc, Xcl1, Spn, Vsir, Il7r |
| GO:0010518 | positive regulation of phospholipase activity | 6 | 8,91E-05 | 0,00161694 | 0,00127899 | Ccl5, Itk, Gpr55, Txk, Agt, Hpca |
| GO:0035458 | cellular response to interferon-beta | 6 | 8,91E-05 | 0,00161694 | 0,00127899 | Tgtp2, Tgtp1, Iigp1, Ifi203, Gbp6, Tmem173 |
| GO:0034284 | response to monosaccharide | 12 | 8,98E-05 | 0,0016206 | 0,00128189 | Pck1, Cacna1e, Baiap3, Pde1c, Thbs1, Pde8b, Neurod1, Hmgn3, Ptprn, Glp1r, Myt1, Zbtb20 |
| GO:1903038 | negative regulation of leukocyte cell-cell adhesion | 10 | 9,35E-05 | 0,00167234 | 0,00132281 | Cd74, Tbx21, Ptpn22, Xcl1, Dtx1, H2-Ab1, Spn, Lax1, H2-Aa, Vsir |
| GO:0002573 | myeloid leukocyte differentiation | 13 | 9,35E-05 | 0,00167234 | 0,00132281 | Ccl5, Cd74, Inpp5d, Farp2, Gpr55, Tlr2, Fbn1, Hcls1, Cd300lf, Ubd, Ikzf1, Fcer1g, Clec2i |
| GO:0002440 | production of molecular mediator of immune response | 17 | 9,51E-05 | 0,00169233 | 0,00133863 | Chga, Card11, Cd74, Tbx21, Ptprc, Ptpn22, Xcl1, H2-Ab1, Cd96, Lax1, Nr4a3, Vsir, Sash3, Tlr2, Il7r, Fcer1g, Cd27 |
| GO:0036037 | CD8-positive, alpha-beta T cell activation | 5 | 9,78E-05 | 0,00173073 | 0,001369 | Ptpn22, Xcl1, Gpr18, Vsir, Nckap1l |
| GO:0097756 | negative regulation of blood vessel diameter | 9 | 0,00010247 | 0,00180515 | 0,00142787 | Add3, Chga, Edn2, Dusp5, Per2, Edn1, Alox5, Ptgs1, Agt |
| GO:0038093 | Fc receptor signaling pathway | 4 | 0,00011171 | 0,00195836 | 0,00154905 | Ptprc, Nr4a3, Myo1g, Fcer1g |
| GO:0061082 | myeloid leukocyte cytokine production | 5 | 0,00011731 | 0,00204669 | 0,00161892 | Chga, Cd74, Nr4a3, Tlr2, Fcer1g |
| GO:0010959 | regulation of metal ion transport | 19 | 0,00012701 | 0,00220529 | 0,00174438 | Cacna1a, Ccl5, Wnk2, Ptpn22, Xcl1, Ccl4, Per1, Pde4b, Coro1a, Thy1, Fyn, Fxyd5, Glp1r, Oprd1, Agt, Usp2, Kcnab2, Capn3, Hpca |
| GO:0001912 | positive regulation of leukocyte mediated cytotoxicity | 8 | 0,000129 | 0,00222924 | 0,00176332 | Vav1, H2-T3, Klrk1, H2-T10, Ptprc, Xcl1, H2-Bl, Klre1 |
| GO:0042100 | B cell proliferation | 9 | 0,00013667 | 0,00235062 | 0,00185933 | Card11, Cd74, Inpp5d, Ptprc, Sash3, Il7r, Ikzf3, Nckap1l, Cd27 |
| mmu04662 | B cell receptor signaling pathway | 8 | 0,00024899 | 0,00244838 | 0,00188925 | 22324, 108723, 22325, 16331, 19354, 18707, 18037, 12517 |
| GO:0071333 | cellular response to glucose stimulus | 10 | 0,00014553 | 0,00249116 | 0,0019705 | Pck1, Cacna1e, Baiap3, Pde1c, Pde8b, Neurod1, Hmgn3, Ptprn, Myt1, Zbtb20 |
| GO:0071331 | cellular response to hexose stimulus | 10 | 0,00016426 | 0,00279859 | 0,00221367 | Pck1, Cacna1e, Baiap3, Pde1c, Pde8b, Neurod1, Hmgn3, Ptprn, Myt1, Zbtb20 |
| mmu04145 | Phagosome | 13 | 0,00029666 | 0,00280048 | 0,00216094 | 15043, 14998, 21825, 14999, 15024, 14961, 14963, 14969, 12721, 14960, 24088, 667803, 16414 |

| | | | | | | |
|---|---|---|---|---|---|---|
| GO:0090022 | regulation of neutrophil chemotaxis | 5 | 0,00016524 | 0,00280207 | 0,00221643 | Cd74, Xcl1, Rac2, Edn1, Nckap1l |
| GO:0043303 | mast cell degranulation | 6 | 0,00016935 | 0,00285852 | 0,00226108 | Chga, Rac2, Nr4a3, Lat, Lat2, Fcer1g |
| GO:0071326 | cellular response to monosaccharide stimulus | 10 | 0,00017436 | 0,00292943 | 0,00231717 | Pck1, Cacna1e, Baiap3, Pde1c, Pde8b, Neurod1, Hmgn3, Ptprn, Myt1, Zbtb20 |
| mmu04670 | Leukocyte transendothelial migration | 10 | 0,00033065 | 0,00300131 | 0,00231591 | 22324, 22325, 19354, 16428, 21838, 74734, 18707, 22165, 16414, 20469 |
| GO:0009743 | response to carbohydrate | 12 | 0,00018188 | 0,00304172 | 0,00240599 | Pck1, Cacna1e, Baiap3, Pde1c, Thbs1, Pde8b, Neurod1, Hmgn3, Ptprn, Glp1r, Myt1, Zbtb20 |
| GO:0002577 | regulation of antigen processing and presentation | 4 | 0,0001894 | 0,00308945 | 0,00244375 | Cd74, Thbs1, H2-Ab1, Was |
| GO:0007202 | activation of phospholipase C activity | 4 | 0,0001894 | 0,00308945 | 0,00244375 | Itk, Gpr55, Txk, Agt |
| GO:0061081 | positive regulation of myeloid leukocyte cytokine production involved in immune response | 4 | 0,0001894 | 0,00308945 | 0,00244375 | Cd74, Nr4a3, Tlr2, Fcer1g |
| GO:2001185 | regulation of CD8-positive, alpha-beta T cell activation | 4 | 0,0001894 | 0,00308945 | 0,00244375 | Ptpn22, Xcl1, Vsir, Nckap1l |
| GO:0002279 | mast cell activation involved in immune response | 6 | 0,0001907 | 0,00308945 | 0,00244375 | Chga, Rac2, Nr4a3, Lat, Lat2, Fcer1g |
| GO:0002448 | mast cell mediated immunity | 6 | 0,0001907 | 0,00308945 | 0,00244375 | Chga, Rac2, Nr4a3, Lat, Lat2, Fcer1g |
| GO:0002720 | positive regulation of cytokine production involved in immune response | 6 | 0,0001907 | 0,00308945 | 0,00244375 | Cd74, Xcl1, Nr4a3, Sash3, Tlr2, Fcer1g |
| GO:0030888 | regulation of B cell proliferation | 7 | 0,0001941 | 0,00313066 | 0,00247634 | Card11, Cd74, Inpp5d, Ptprc, Sash3, Ikzf3, Nckap1l |
| GO:0002824 | positive regulation of adaptive immune response based on somatic recombination of immune receptors built from immunoglobulin superfamily domains | 10 | 0,00020785 | 0,00333763 | 0,00264006 | H2-T3, Tbx21, H2-DMa, H2-T10, Ptprc, Xcl1, H2-Ab1, H2-Bl, Sash3, Fcer1g |
| GO:0032743 | positive regulation of interleukin-2 production | 5 | 0,00022683 | 0,00361039 | 0,00285581 | Cd3e, Card11, Ptprc, Pde4b, Sash3 |
| GO:1902624 | positive regulation of neutrophil migration | 5 | 0,00022683 | 0,00361039 | 0,00285581 | Cd74, Xcl1, Rac2, Edn1, Nckap1l |
| GO:0002532 | production of molecular mediator involved in inflammatory response | 7 | 0,00023289 | 0,00369063 | 0,00291928 | Cd96, Alox5, Per1, Nlrc3, Adcy7, Fcer1g, Gpsm3 |
| GO:0072676 | lymphocyte migration | 8 | 0,00024159 | 0,00381182 | 0,00301513 | Ccl5, Tbx21, Klrk1, Xcl1, Ccl4, Spn, Itgb7, Myo1g |
| GO:0043087 | regulation of GTPase activity | 15 | 0,00024281 | 0,00381448 | 0,00301724 | Ccl5, Ntrk2, Vav1, Rtn4r, Vav2, Rgs1, Klrk1, Xcl1, Tbc1d10c, Ccl4, Arap3, Rasal3, Thy1, Dock10, Tbc1d16 |
| mmu04672 | Intestinal immune network for IgA production | 6 | 0,00043712 | 0,00382076 | 0,00294822 | 14998, 14999, 14961, 14969, 14960, 16421 |
| mmu04713 | Circadian entrainment | 9 | 0,00047282 | 0,00398524 | 0,00307514 | 19416, 18796, 18627, 14710, 18626, 19092, 58226, 11513, 18628 |
| GO:0071322 | cellular response to carbohydrate stimulus | 10 | 0,00026078 | 0,00407913 | 0,00322658 | Pck1, Cacna1e, Baiap3, Pde1c, Pde8b, Neurod1, Hmgn3, Ptprn, Myt1, Zbtb20 |
| GO:0032653 | regulation of interleukin-10 production | 6 | 0,00026769 | 0,00416931 | 0,00329791 | Dll1, Xcl1, Vsir, Sash3, Tlr2, Fcer1g |
| GO:0017157 | regulation of exocytosis | 13 | 0,00029745 | 0,00452465 | 0,00357898 | Cacna1a, Baiap3, Grik5, Pclo, Nrn1, Rac2, Sept1, Itgb2, Cacna1h, Grm4, Fcer1g, Syp, Rab3c |
| GO:0046777 | protein autophosphorylation | 13 | 0,00029745 | 0,00452465 | 0,00357898 | Ntrk2, Wnk2, Hck, Ptprc, Zap70, Itk, Thy1, Sla, Prkg2, Lck, Txk, Fyn, Pdgfd |
| GO:0035456 | response to interferon-beta | 6 | 0,00029816 | 0,00452465 | 0,00357898 | Tgtp2, Tgtp1, Iigp1, Ifi203, Gbp6, Tmem173 |
| GO:0050854 | regulation of antigen receptor-mediated signaling pathway | 6 | 0,00029816 | 0,00452465 | 0,00357898 | Ptprc, Ptpn22, Thy1, Lck, Lpxn, Trat1 |
| GO:1901570 | fatty acid derivative biosynthetic process | 6 | 0,00029816 | 0,00452465 | 0,00357898 | Edn2, Hpgds, Cd74, Edn1, Alox5, Ptgs1 |
| GO:0033630 | positive regulation of cell adhesion mediated by integrin | 4 | 0,00029995 | 0,00452465 | 0,00357898 | Ccl5, Cd3e, Skap1, Nckap1l |
| GO:0045076 | regulation of interleukin-2 biosynthetic process | 4 | 0,00029995 | 0,00452465 | 0,00357898 | Cd3e, Card11, Ptprc, Clec2i |
| GO:0002444 | myeloid leukocyte mediated immunity | 8 | 0,00030048 | 0,00452465 | 0,00357898 | Chga, Rac2, Nr4a3, Itgb2, Myo1f, Lat, Lat2, Fcer1g |
| GO:0097696 | STAT cascade | 11 | 0,00032172 | 0,00481675 | 0,00381003 | Prlr, Ccl5, Ptprc, Neurod1, Ptprt, Lck, Il7r, Fyn, Hcls1, Agap2, Agt |
| GO:0031343 | positive regulation of cell killing | 8 | 0,00032253 | 0,00481675 | 0,00381003 | Vav1, H2-T3, Klrk1, H2-T10, Ptprc, Xcl1, H2-Bl, Klre1 |
| GO:0010517 | regulation of phospholipase activity | 6 | 0,00033128 | 0,00492707 | 0,0038973 | Ccl5, Itk, Gpr55, Txk, Agt, Hpca |
| GO:0001505 | regulation of neurotransmitter levels | 17 | 0,00034189 | 0,00506414 | 0,00400572 | Cacna1a, Baiap3, Ntrk2, Tph1, Grik5, Pclo, Klrk1, Nrn1, Per2, Tlr2, Itgb2, Grm4, Hnmt, Agt, Syn2, Fev, Syp |
| GO:0050796 | regulation of insulin secretion | 11 | 0,00035413 | 0,0052242 | 0,00413232 | Cacna1e, Chga, Ccl5, Baiap3, Pde1c, Pde8b, Hmgn3, Per2, Glp1r, Myt1, Myrip |
| mmu04611 | Platelet activation | 10 | 0,00064356 | 0,00523728 | 0,00404125 | 54519, 18796, 320207, 19092, 18707, 19224, 14360, 16822, 11513, 14127 |
| GO:0019229 | regulation of vasoconstriction | 7 | 0,00035762 | 0,00525428 | 0,00415612 | Add3, Edn2, Dusp5, Per2, Edn1, Alox5, Ptgs1 |
| GO:0001678 | cellular glucose homeostasis | 10 | 0,000361 | 0,00528258 | 0,00417851 | Pck1, Cacna1e, Baiap3, Pde1c, Pde8b, Neurod1, Hmgn3, Ptprn, Myt1, Zbtb20 |
| GO:0001914 | regulation of T cell mediated cytotoxicity | 6 | 0,00036721 | 0,00530919 | 0,00419955 | H2-T3, H2-T10, Ptprc, Xcl1, H2-Bl, Il7r |
| GO:0032613 | interleukin-10 production | 6 | 0,00036721 | 0,00530919 | 0,00419955 | Dll1, Xcl1, Vsir, Sash3, Tlr2, Fcer1g |
| GO:0060193 | positive regulation of lipase activity | 6 | 0,00036721 | 0,00530919 | 0,00419955 | Ccl5, Itk, Gpr55, Txk, Agt, Hpca |
| GO:0070269 | pyroptosis | 4 | 0,00036974 | 0,00531586 | 0,00420483 | Gsdmc3, Gsdmc2, Gsdmc4, Gsdma |
| GO:0042310 | vasoconstriction | 8 | 0,0003706 | 0,00531586 | 0,00420483 | Add3, Edn2, Dusp5, Per2, Edn1, Alox5, Ptgs1, Agt |
| GO:0022408 | negative regulation of cell-cell adhesion | 11 | 0,00038925 | 0,00556132 | 0,00439899 | Adamts18, Cd74, Tbx21, Ptpn22, Xcl1, Dtx1, H2-Ab1, Spn, Lax1, H2-Aa, Vsir |
| GO:0002698 | negative regulation of immune effector process | 9 | 0,00040619 | 0,0057807 | 0,00457251 | Tbx21, Ptprc, Xcl1, Cd96, Spn, Serpinb9, Vsir, Il7r, Klre1 |
| GO:0097553 | calcium ion transmembrane import into cytosol | 9 | 0,0004305 | 0,00610266 | 0,00482719 | Ptprc, Fasl, Xcl1, Coro1a, Thy1, Lck, Fyn, Glp1r, Capn3 |
| GO:0042094 | interleukin-2 biosynthetic process | 4 | 0,00045041 | 0,0063265 | 0,00500424 | Cd3e, Card11, Ptprc, Clec2i |
| GO:0032479 | regulation of type I interferon production | 7 | 0,000455 | 0,0063265 | 0,00500424 | Ptpn22, Gbp4, Tlr2, Nlrc3, Peli3, Tmem173, Zbtb20 |

| | | | | | | |
|---|---|---|---|---|---|---|
| GO:0006636 | unsaturated fatty acid biosynthetic process | 5 | 0,00045675 | 0,0063265 | 0,00500424 | Edn2, Hpgds, Cd74, Edn1, Ptgs1 |
| GO:0006692 | prostanoid metabolic process | 5 | 0,00045675 | 0,0063265 | 0,00500424 | Edn2, Hpgds, Cd74, Edn1, Ptgs1 |
| GO:0006693 | prostaglandin metabolic process | 5 | 0,00045675 | 0,0063265 | 0,00500424 | Edn2, Hpgds, Cd74, Edn1, Ptgs1 |
| GO:0032633 | interleukin-4 production | 5 | 0,00045675 | 0,0063265 | 0,00500424 | Cd3e, Itk, Sash3, Txk, Fcer1g |
| mmu05140 | Leishmaniasis | 7 | 0,00088535 | 0,0065843 | 0,00508066 | 14998, 14999, 14961, 14969, 14960, 24088, 16414 |
| mmu05152 | Tuberculosis | 12 | 0,00089018 | 0,0065843 | 0,00508066 | 16149, 14998, 14999, 14961, 14969, 12721, 14960, 12265, 24088, 16414, 16411, 14127 |
| mmu05145 | Toxoplasmosis | 9 | 0,00089279 | 0,0065843 | 0,00508066 | 14998, 14999, 320207, 14961, 11689, 14969, 14960, 12265, 24088 |
| GO:0048872 | homeostasis of number of cells | 15 | 0,00048739 | 0,00672527 | 0,00531966 | Card11, Cd74, Inpp5d, Cd7, Pde4b, Coro1a, Sash3, Pik3cd, Dock10, Il7r, Hcls1, Lat, Nckap1l, Ikzf1, Fcer1g |
| GO:0072330 | monocarboxylic acid biosynthetic process | 13 | 0,00049691 | 0,00683057 | 0,00540296 | Slc27a2, Edn2, Hpgds, Cd74, Chst12, Per2, Fbp1, Edn1, Ptgs1, Malrd1, Prox1, Agt, Zbtb20 |
| GO:0070371 | ERK1 and ERK2 cascade | 16 | 0,000503 | 0,00688822 | 0,00544855 | Ccl5, Ntrk2, Wnk2, Cd74, Ptprc, Ptpn22, Xcl1, Tbc1d10c, Ccl4, Ackr3, Glipr2, Gpr55, Tlr2, Agt, Ror1, Pdgfd |
| GO:0070374 | positive regulation of ERK1 and ERK2 cascade | 13 | 0,00051609 | 0,00701985 | 0,00555268 | Ccl5, Ntrk2, Cd74, Ptprc, Ptpn22, Xcl1, Ccl4, Ackr3, Glipr2, Gpr55, Tlr2, Ror1, Pdgfd |
| GO:0030316 | osteoclast differentiation | 8 | 0,00051668 | 0,00701985 | 0,00555268 | Ccl5, Inpp5d, Farp2, Gpr55, Fbn1, Cd300lf, Fcer1g, Clec2i |
| GO:0033628 | regulation of cell adhesion mediated by integrin | 5 | 0,00051841 | 0,00701985 | 0,00555268 | Ccl5, Cd3e, Skap1, Lpxn, Nckap1l |
| GO:0001911 | negative regulation of leukocyte mediated cytotoxicity | 4 | 0,00054288 | 0,00726976 | 0,00575036 | Ptprc, Serpinb9, Il7r, Klre1 |
| GO:0043153 | entrainment of circadian clock by photoperiod | 4 | 0,00054288 | 0,00726976 | 0,00575036 | Per2, Per1, Per3, Usp2 |
| GO:1901623 | regulation of lymphocyte chemotaxis | 4 | 0,00054288 | 0,00726976 | 0,00575036 | Ccl5, Klrk1, Xcl1, Ccl4 |
| GO:0032606 | type I interferon production | 7 | 0,00057245 | 0,00763753 | 0,00604126 | Ptpn22, Gbp4, Tlr2, Nlrc3, Peli3, Tmem173, Zbtb20 |
| GO:0090276 | regulation of peptide hormone secretion | 12 | 0,00059375 | 0,00787243 | 0,00622707 | Cacna1e, Chga, Ccl5, Baiap3, Pde1c, Pde8b, Hmgn3, Per2, Glp1r, Myt1, Pex5l, Myrip |
| GO:0002315 | marginal zone B cell differentiation | 3 | 0,00059656 | 0,00787243 | 0,00622707 | Dll1, Dock10, Mfng |
| GO:0038094 | Fc-gamma receptor signaling pathway | 3 | 0,00059656 | 0,00787243 | 0,00622707 | Ptprc, Myo1g, Fcer1g |
| GO:0051209 | release of sequestered calcium ion into cytosol | 8 | 0,00062465 | 0,00821321 | 0,00649662 | Ptprc, Fasl, Xcl1, Coro1a, Thy1, Lck, Glp1r, Capn3 |
| GO:0002719 | negative regulation of cytokine production involved in immune response | 4 | 0,00064808 | 0,00845996 | 0,0066918 | Tbx21, Xcl1, Cd96, Vsir |
| GO:0009648 | photoperiodism | 4 | 0,00064808 | 0,00845996 | 0,0066918 | Per2, Per1, Per3, Usp2 |
| GO:0032922 | circadian regulation of gene expression | 6 | 0,00065138 | 0,00847258 | 0,00670178 | Per2, Per1, Prkg2, Ciart, Per3, Usp2 |
| GO:0030099 | myeloid cell differentiation | 17 | 0,00065662 | 0,0085102 | 0,00673154 | Dll1, Ccl5, Cd74, Inpp5d, Gpr171, Farp2, Fli1, Gpr55, Tlr2, Fbn1, Hcls1, Cd300lf, Nckap1l, Ubd, Ikzf1, Fcer1g, Clec2i |
| GO:1902622 | regulation of neutrophil migration | 5 | 0,00066026 | 0,00852705 | 0,00674486 | Cd74, Xcl1, Rac2, Edn1, Nckap1l |
| GO:0001776 | leukocyte homeostasis | 8 | 0,00066443 | 0,00855045 | 0,00676337 | Cd74, Pde4b, Coro1a, Pik3cd, Dock10, Lat, Nckap1l, Fcer1g |
| mmu05162 | Measles | 10 | 0,00123691 | 0,00884578 | 0,00682569 | 12502, 12501, 14103, 16186, 16185, 18707, 24088, 14360, 12500, 22035 |
| GO:1904892 | regulation of STAT cascade | 10 | 0,00069083 | 0,00885875 | 0,00700724 | Ccl5, Ptprc, Neurod1, Ptprt, Lck, Il7r, Fyn, Hcls1, Agap2, Agt |
| GO:0051283 | negative regulation of sequestering of calcium ion | 8 | 0,00070624 | 0,0090244 | 0,00713827 | Ptprc, Fasl, Xcl1, Coro1a, Thy1, Lck, Glp1r, Capn3 |
| mmu05168 | Herpes simplex virus 1 infection | 21 | 0,00130218 | 0,00903869 | 0,00697455 | 20304, 210503, 15043, 16149, 14998, 14999, 15024, 14103, 78921, 14961, 14963, 14969, 14960, 18707, 24088, 210104, 667803, 237758, 230162, 72512, 76373 |
| mmu05170 | Human immunodeficiency virus 1 infection | 14 | 0,00134464 | 0,0090667 | 0,00699616 | 12502, 12501, 15043, 15024, 14103, 19354, 12503, 14710, 14963, 18707, 24088, 667803, 12500, 72512 |
| GO:0035773 | insulin secretion involved in cellular response to glucose stimulus | 7 | 0,0007128 | 0,00907627 | 0,0071793 | Cacna1e, Baiap3, Pde1c, Pde8b, Hmgn3, Ptprn, Myt1 |
| GO:0007015 | actin filament organization | 17 | 0,00073481 | 0,00932385 | 0,00737513 | Add3, Acta1, Limch1, Rhobtb3, Rac2, Coro1a, Arhgap25, Rhoh, Tlr2, Hcls1, Gdpd2, Myom3, Prox1, Was, Nckap1l, Clec2i, Pstpip1 |
| GO:0032103 | positive regulation of response to external stimulus | 14 | 0,00074071 | 0,00936601 | 0,00740848 | Ccl5, Edn2, Cd74, Thbs1, Xcl1, Rac2, Ccl4, Smoc2, Edn1, Tlr2, Nckap1l, Fcer1g, Pdgfd, Gpsm3 |
| GO:0043113 | receptor clustering | 6 | 0,00077657 | 0,0097514 | 0,00771332 | Cacna1a, Grik5, Thy1, Itgb7, Itgb2, Clec2i |
| GO:2000514 | regulation of CD4-positive, alpha-beta T cell activation | 6 | 0,00077657 | 0,0097514 | 0,00771332 | Tbx21, Xcl1, Il2rg, Vsir, Sash3, Nckap1l |
| GO:0051282 | regulation of sequestering of calcium ion | 8 | 0,00079619 | 0,00996341 | 0,00788102 | Ptprc, Fasl, Xcl1, Coro1a, Thy1, Lck, Glp1r, Capn3 |
| GO:0001865 | NK T cell differentiation | 3 | 0,00080956 | 0,01002694 | 0,00793128 | Itk, Txk, Ikzf1 |
| GO:0014824 | artery smooth muscle contraction | 3 | 0,00080956 | 0,01002694 | 0,00793128 | Edn2, Edn1, Agt |
| GO:0033632 | regulation of cell-cell adhesion mediated by integrin | 3 | 0,00080956 | 0,01002694 | 0,00793128 | Ccl5, Cd3e, Skap1 |
| GO:0052548 | regulation of endopeptidase activity | 15 | 0,00081257 | 0,01003004 | 0,00793373 | Nr4a1, Pcsk1n, Casp12, Thbs1, Fasl, Serpinb9, Vsir, Lck, Fyn, Cyfip2, Nek5, Psmb8, Tnfsf10, Agt, St18 |
| GO:1903524 | positive regulation of blood circulation | 7 | 0,00082058 | 0,0100945 | 0,00798471 | Add3, Chga, Edn2, Edn1, Alox5, Ptgs1, Glp1r |
| GO:0007160 | cell-matrix adhesion | 11 | 0,00082365 | 0,01009807 | 0,00798754 | Hoxd3, Cd3e, Limch1, Thbs1, Sned1, Cd96, Thy1, Skap1, Itgb7, Itgb2, Agt |
| GO:0002534 | cytokine production involved in inflammatory response | 5 | 0,00082932 | 0,01013335 | 0,00801544 | Cd96, Per1, Nlrc3, Adcy7, Gpsm3 |
| GO:0070372 | regulation of ERK1 and ERK2 cascade | 15 | 0,0008379 | 0,01020377 | 0,00807115 | Ccl5, Ntrk2, Wnk2, Cd74, Ptprc, Ptpn22, Xcl1, Tbc1d10c, Ccl4, Ackr3, Glipr2, Gpr55, Tlr2, Ror1, Pdgfd |
| GO:0051208 | sequestering of calcium ion | 8 | 0,0008445 | 0,01024985 | 0,0081076 | Ptprc, Fasl, Xcl1, Coro1a, Thy1, Lck, Glp1r, Capn3 |
| mmu04810 | Regulation of actin cytoskeleton | 13 | 0,00162659 | 0,01066321 | 0,00822808 | 22324, 22325, 19354, 16407, 18707, 76884, 16421, 16414, 73341, 22376, 105855, 16411, 71785 |
| GO:0052547 | regulation of peptidase activity | 17 | 0,00089089 | 0,01071335 | 0,00847422 | Nr4a1, Pcsk1n, Casp12, Tfpi, Thbs1, Fasl, Serpinb9, Vsir, Lck, Fyn, Cyfip2, Cst7, Nek5, Psmb8, Tnfsf10, Agt, St18 |

| | | | | | | |
|---|---|---|---|---|---|---|
| GO:0006925 | inflammatory cell apoptotic process | 4 | 0,00090041 | 0,01071335 | 0,00847422 | Ccl5, Fasl, Pik3cd, St6gal1 |
| GO:0009649 | entrainment of circadian clock | 4 | 0,00090041 | 0,01071335 | 0,00847422 | Per2, Per1, Per3, Usp2 |
| GO:0014821 | phasic smooth muscle contraction | 4 | 0,00090041 | 0,01071335 | 0,00847422 | Edn2, Edn1, Sstr2, Agt |
| GO:0031342 | negative regulation of cell killing | 4 | 0,00090041 | 0,01071335 | 0,00847422 | Ptprc, Serpinb9, Il7r, Klre1 |
| GO:0071402 | cellular response to lipoprotein particle stimulus | 4 | 0,00090041 | 0,01071335 | 0,00847422 | Mylip, Abcg1, Itgb2, Fcer1g |
| GO:1903522 | regulation of blood circulation | 12 | 0,00090846 | 0,01077383 | 0,00852206 | Cacna1e, Add3, Chga, Edn2, Dusp5, Per2, Edn1, Alox5, Pde4b, Ptgs1, Glp1r, Agt |
| GO:0002831 | regulation of response to biotic stimulus | 9 | 0,00091633 | 0,01079658 | 0,00854006 | Ccl5, Klrk1, Ptpn22, Xcl1, Spn, Gbp4, Tlr2, Klre1, Tmem173 |
| GO:0019233 | sensory perception of pain | 9 | 0,00091633 | 0,01079658 | 0,00854006 | Cacna1e, Cacna1a, Edn1, Alox5, Ptgs1, Fyn, Cacna1h, Oprd1, Scn3a |
| GO:0006836 | neurotransmitter transport | 14 | 0,00096134 | 0,01129034 | 0,00893062 | Cacna1a, Baiap3, Ntrk2, Grik5, Slc6a20a, Pclo, Nrn1, Per2, Edn1, Grm4, Fcer1g, Syn2, Fev, Syp |
| GO:0042267 | natural killer cell mediated cytotoxicity | 6 | 0,00099824 | 0,01168584 | 0,00924345 | Vav1, Klrk1, Gzmb, Coro1a, Serpinb9, Klre1 |
| GO:0050764 | regulation of phagocytosis | 7 | 0,00100595 | 0,01173819 | 0,00928487 | Hck, Ptprc, Dock2, Tlr2, Cd300lf, Nckap1l, Fcer1g |
| GO:0002347 | response to tumor cell | 4 | 0,00104942 | 0,01212848 | 0,00959359 | Klrk1, Xcl1, Rbms3, Klre1 |
| GO:0010863 | positive regulation of phospholipase C activity | 4 | 0,00104942 | 0,01212848 | 0,00959359 | Itk, Gpr55, Txk, Agt |
| GO:0070233 | negative regulation of T cell apoptotic process | 4 | 0,00104942 | 0,01212848 | 0,00959359 | Ccl5, Serpinb9, Il7r, Cd27 |
| GO:0045086 | positive regulation of interleukin-2 biosynthetic process | 3 | 0,00106533 | 0,01223445 | 0,00967741 | Cd3e, Card11, Ptprc |
| GO:0061469 | regulation of type B pancreatic cell proliferation | 3 | 0,00106533 | 0,01223445 | 0,00967741 | Nr4a1, Ptprn, Nr4a3 |
| GO:0043583 | ear development | 12 | 0,00109182 | 0,01249909 | 0,00988674 | Sobp, Dll1, Lgr5, H2-DMa, Fzd3, Neurod1, Edn1, Nr4a3, Fzd2, Prox1, Ror1, Hpca |
| GO:0048839 | inner ear development | 11 | 0,00113696 | 0,01297494 | 0,01026313 | Sobp, Dll1, Lgr5, H2-DMa, Fzd3, Neurod1, Nr4a3, Fzd2, Prox1, Ror1, Hpca |
| GO:0030890 | positive regulation of B cell proliferation | 5 | 0,00114073 | 0,01297712 | 0,01026486 | Card11, Cd74, Ptprc, Sash3, Nckap1l |
| GO:0002724 | regulation of T cell cytokine production | 4 | 0,00121492 | 0,01373508 | 0,0108644 | Tbx21, Xcl1, Vsir, Sash3 |
| GO:0042119 | neutrophil activation | 4 | 0,00121492 | 0,01373508 | 0,0108644 | Ccl5, Itgb2, Myo1f, Fcer1g |
| GO:1903305 | regulation of regulated secretory pathway | 10 | 0,00123719 | 0,01394339 | 0,01102918 | Cacna1a, Baiap3, Grik5, Nrn1, Rac2, Itgb2, Cacna1h, Grm4, Fcer1g, Syp |
| GO:0032418 | lysosome localization | 6 | 0,00126557 | 0,01417519 | 0,01121252 | Chga, Rac2, Nr4a3, Lat, Lat2, Fcer1g |
| GO:0046637 | regulation of alpha-beta T cell differentiation | 6 | 0,00126557 | 0,01417519 | 0,01121252 | Tbx21, Zap70, Il2rg, Sash3, Nckap1l, Ikzf1 |
| GO:0010950 | positive regulation of endopeptidase activity | 9 | 0,00128905 | 0,01434962 | 0,0113505 | Casp12, Fasl, Vsir, Lck, Fyn, Cyfip2, Nek5, Tnfsf10, St18 |
| GO:0060402 | calcium ion transport into cytosol | 9 | 0,00128905 | 0,01434962 | 0,0113505 | Ptprc, Fasl, Xcl1, Coro1a, Thy1, Lck, Fyn, Glp1r, Capn3 |
| GO:1903169 | regulation of calcium ion transmembrane transport | 9 | 0,00135106 | 0,01494097 | 0,01181825 | Cacna1a, Ptpn22, Xcl1, Pde4b, Coro1a, Thy1, Fyn, Capn3, Hpca |
| GO:0002551 | mast cell chemotaxis | 3 | 0,00136688 | 0,01494097 | 0,01181825 | Chga, Ccl5, Rac2 |
| GO:0014820 | tonic smooth muscle contraction | 3 | 0,00136688 | 0,01494097 | 0,01181825 | Edn2, Edn1, Agt |
| GO:0030240 | skeletal muscle thin filament assembly | 3 | 0,00136688 | 0,01494097 | 0,01181825 | Acta1, Myom3, Prox1 |
| GO:0034116 | positive regulation of heterotypic cell-cell adhesion | 3 | 0,00136688 | 0,01494097 | 0,01181825 | Thy1, Skap1, Lck |
| GO:0042535 | positive regulation of tumor necrosis factor biosynthetic process | 3 | 0,00136688 | 0,01494097 | 0,01181825 | Thbs1, Spn, Tlr2 |
| GO:1901617 | organic hydroxy compound biosynthetic process | 11 | 0,00137845 | 0,01502221 | 0,01188252 | Pck1, Slc27a2, Ces1d, Tph1, Per2, Abcg1, Nr4a2, Cacna1h, Malrd1, Prox1, Fev |
| GO:0007200 | phospholipase C-activating G protein-coupled receptor signaling pathway | 7 | 0,0013865 | 0,01506466 | 0,0119161 | Chga, Plcb2, P2ry10, Gpr18, Edn1, Gpr55, Oprd1 |
| GO:0045671 | negative regulation of osteoclast differentiation | 4 | 0,00139786 | 0,01514279 | 0,01197789 | Inpp5d, Gpr55, Fbn1, Clec2i |
| GO:0051668 | localization within membrane | 10 | 0,0014035 | 0,01515866 | 0,01199045 | Cacna1a, Grik5, Ptprc, Grasp, Dock2, Thy1, Itgb7, Itgb2, Hpca, Clec2i |
| GO:0045637 | regulation of myeloid cell differentiation | 11 | 0,00143146 | 0,0154148 | 0,01219305 | Dll1, Ccl5, Cd74, Inpp5d, Gpr171, Gpr55, Fbn1, Hcls1, Nckap1l, Ikzf1, Clec2i |
| mmu04072 | Phospholipase D signaling pathway | 10 | 0,00244844 | 0,01561709 | 0,01205066 | 18796, 320207, 18707, 14360, 72318, 268934, 11513, 11606, 14127, 71785 |
| GO:0002761 | regulation of myeloid leukocyte differentiation | 8 | 0,00146926 | 0,01575833 | 0,01246479 | Ccl5, Cd74, Inpp5d, Gpr55, Fbn1, Hcls1, Ikzf1, Clec2i |
| GO:0002711 | positive regulation of T cell mediated immunity | 6 | 0,00147205 | 0,01575833 | 0,01246479 | H2-T3, H2-T10, Ptprc, Xcl1, H2-Bl, Sash3 |
| GO:0002791 | regulation of peptide secretion | 19 | 0,00148311 | 0,01581584 | 0,01251028 | Cacna1e, Chga, Ccl5, Baiap3, Pde1c, Cd74, Pde8b, Ptpn22, Hmgn3, Per2, Srgn, Tlr2, Cd2, Malrd1, Glp1r, Agt, Myt1, Pex5l, Myrip |
| GO:0045089 | positive regulation of innate immune response | 11 | 0,00148614 | 0,01581584 | 0,01251028 | Vav1, Cd74, Klrk1, Ptpn22, Xcl1, Tlr2, Txk, Cd300lf, Klre1, Peli3, Tmem173 |
| GO:0007249 | I-kappaB kinase/NF-kappaB signaling | 10 | 0,00158756 | 0,01684573 | 0,01332491 | Card11, Cd74, Per1, Rhoh, Tlr2, Fyn, Nlrc3, Ubd, Ror1, Capn3 |
| GO:0051085 | chaperone cofactor-dependent protein refolding | 4 | 0,00159916 | 0,01688569 | 0,01335652 | Cd74, H2-DMa, H2-DMb1, Dnajb4 |
| GO:0043547 | positive regulation of GTPase activity | 11 | 0,00160063 | 0,01688569 | 0,01335652 | Ccl5, Vav1, Rtn4r, Rgs1, Xcl1, Tbc1d10c, Ccl4, Arap3, Thy1, Dock10, Tbc1d16 |
| GO:0070227 | lymphocyte apoptotic process | 7 | 0,00166366 | 0,01747835 | 0,01382531 | Ccl5, Cd3g, Cd74, Fasl, Serpinb9, Il7r, Cd27 |
| GO:0001916 | positive regulation of T cell mediated cytotoxicity | 5 | 0,00168089 | 0,01747835 | 0,01382531 | H2-T3, H2-T10, Ptprc, Xcl1, H2-Bl |
| GO:0046638 | positive regulation of alpha-beta T cell differentiation | 5 | 0,00168089 | 0,01747835 | 0,01382531 | Zap70, Il2rg, Sash3, Nckap1l, Ikzf1 |
| GO:0050850 | positive regulation of calcium-mediated signaling | 5 | 0,00168089 | 0,01747835 | 0,01382531 | Cd3e, Zap70, Ccl4, Trat1, Cd8a |

| | | | | | | |
|---|---|---|---|---|---|---|
| GO:0070229 | negative regulation of lymphocyte apoptotic process | 5 | 0,00168089 | 0,01747835 | 0,01382531 | Ccl5, Cd74, Serpinb9, Il7r, Cd27 |
| GO:1903707 | negative regulation of hemopoiesis | 9 | 0,00169812 | 0,01755217 | 0,0138837 | Dll1, Cd74, Tbx21, Inpp5d, Gpr171, Dtx1, Gpr55, Fbn1, Clec2i |
| GO:0060191 | regulation of lipase activity | 6 | 0,00170324 | 0,01755217 | 0,0138837 | Ccl5, Itk, Gpr55, Txk, Agt, Hpca |
| GO:0014866 | skeletal myofibril assembly | 3 | 0,00171701 | 0,01755217 | 0,0138837 | Acta1, Myom3, Prox1 |
| GO:0033631 | cell-cell adhesion mediated by integrin | 3 | 0,00171701 | 0,01755217 | 0,0138837 | Ccl5, Cd3e, Skap1 |
| GO:0070486 | leukocyte aggregation | 3 | 0,00171701 | 0,01755217 | 0,0138837 | Rac2, Spn, Nr4a3 |
| GO:0097531 | mast cell migration | 3 | 0,00171701 | 0,01755217 | 0,0138837 | Chga, Ccl5, Rac2 |
| GO:0043122 | regulation of I-kappaB kinase/NF-kappaB signaling | 9 | 0,00177541 | 0,01809823 | 0,01431564 | Card11, Cd74, Per1, Rhoh, Fyn, Nlrc3, Ubd, Ror1, Capn3 |
| GO:0002285 | lymphocyte activation involved in immune response | 10 | 0,00179072 | 0,01820311 | 0,01439859 | Dll1, Apbb1ip, Tbx21, Ptprc, Spn, Coro1a, Dock10, Mfng, Fcer1g, Ly9 |
| GO:0008277 | regulation of G protein-coupled receptor signaling pathway | 8 | 0,00180363 | 0,0182832 | 0,01446195 | Chga, Ccl5, Rgs1, Edn1, Pde4b, Fzd2, Rgs13, Syp |
| GO:0032660 | regulation of interleukin-17 production | 4 | 0,00181977 | 0,01829347 | 0,01447007 | Vsir, Tlr2, Nckap1l, Ly9 |
| GO:0035767 | endothelial cell chemotaxis | 4 | 0,00181977 | 0,01829347 | 0,01447007 | Nr4a1, Thbs1, Smoc2, Egr3 |
| GO:1900274 | regulation of phospholipase C activity | 4 | 0,00181977 | 0,01829347 | 0,01447007 | Itk, Gpr55, Txk, Agt |
| GO:0010952 | positive regulation of peptidase activity | 9 | 0,0018555 | 0,0184989 | 0,01463257 | Casp12, Fasl, Vsir, Lck, Fyn, Cyfip2, Nek5, Tnfsf10, St18 |
| GO:0035296 | regulation of tube diameter | 9 | 0,0018555 | 0,0184989 | 0,01463257 | Add3, Chga, Edn2, Dusp5, Per2, Edn1, Alox5, Ptgs1, Agt |
| GO:0097746 | regulation of blood vessel diameter | 9 | 0,0018555 | 0,0184989 | 0,01463257 | Add3, Chga, Edn2, Dusp5, Per2, Edn1, Alox5, Ptgs1, Agt |
| GO:0050848 | regulation of calcium-mediated signaling | 7 | 0,00187113 | 0,01855281 | 0,01467521 | Cd3e, Zap70, Tbc1d10c, Ccl4, Sla2, Trat1, Cd8a |
| GO:1902106 | negative regulation of leukocyte differentiation | 7 | 0,00187113 | 0,01855281 | 0,01467521 | Cd74, Tbx21, Inpp5d, Dtx1, Gpr55, Fbn1, Clec2i |
| GO:0050730 | regulation of peptidyl-tyrosine phosphorylation | 12 | 0,00189645 | 0,01875261 | 0,01483325 | Ccl5, Cd3e, Cd74, Ptprc, Ptpn22, Thy1, Lck, Fyn, Hcls1, Itgb2, Agt, Pdgfd |
| GO:1904062 | regulation of cation transmembrane transport | 14 | 0,00197977 | 0,0195233 | 0,01544286 | Rasgrf2, Cacna1a, Wnk2, Ptpn22, Xcl1, Pde4b, Coro1a, Thy1, Fyn, Fxyd5, Agt, Kcnab2, Capn3, Hpca |
| GO:0045933 | positive regulation of muscle contraction | 5 | 0,00201203 | 0,0197877 | 0,01565201 | Chga, Edn2, Edn1, Lck, Ptgs1 |
| GO:0035025 | positive regulation of Rho protein signal transduction | 4 | 0,0020606 | 0,02021056 | 0,01598648 | Rtn4r, P2ry10, Gpr18, Gpr55 |
| GO:0015711 | organic anion transport | 16 | 0,00208995 | 0,02040113 | 0,01613722 | Slco2b1, Slc27a2, Cacna1a, Ntrk2, Slc6a20a, Thbs1, Per2, Slc37a2, Abcg1, Edn1, Lrrc8c, Emb, Nfkbie, Atp8b2, Agt, Slc15a1 |
| GO:0042593 | glucose homeostasis | 12 | 0,00209127 | 0,02040113 | 0,01613722 | Pck1, Cacna1e, Cacna1a, Baiap3, Pde1c, Pde8b, Neurod1, Hmgn3, Ptprn, Pygl, Myt1, Zbtb20 |
| GO:0033089 | positive regulation of T cell differentiation in thymus | 3 | 0,00211833 | 0,02049981 | 0,01621528 | Egr3, Il2rg, Il7r |
| GO:0051923 | sulfation | 3 | 0,00211833 | 0,02049981 | 0,01621528 | Sult1c2, Gal3st2, Sult1a1 |
| GO:0071404 | cellular response to low-density lipoprotein particle stimulus | 3 | 0,00211833 | 0,02049981 | 0,01621528 | Mylip, Itgb2, Fcer1g |
| GO:0033500 | carbohydrate homeostasis | 12 | 0,00215974 | 0,02084493 | 0,01648827 | Pck1, Cacna1e, Cacna1a, Baiap3, Pde1c, Pde8b, Neurod1, Hmgn3, Ptprn, Pygl, Myt1, Zbtb20 |
| GO:0060401 | cytosolic calcium ion transport | 9 | 0,0022052 | 0,0212272 | 0,01679064 | Ptprc, Fasl, Xcl1, Coro1a, Thy1, Lck, Fyn, Glp1r, Capn3 |
| GO:1904894 | positive regulation of STAT cascade | 7 | 0,00221929 | 0,02130637 | 0,01685326 | Ccl5, Lck, Il7r, Fyn, Hcls1, Agap2, Agt |
| GO:0061178 | regulation of insulin secretion involved in cellular response to glucose stimulus | 6 | 0,00224718 | 0,02151719 | 0,01702003 | Cacna1e, Baiap3, Pde1c, Pde8b, Hmgn3, Myt1 |
| GO:0003018 | vascular process in circulatory system | 10 | 0,00226011 | 0,02158402 | 0,01707288 | Add3, Chga, Edn2, Dusp5, Per2, Ccl4, Edn1, Alox5, Ptgs1, Agt |
| GO:0032733 | positive regulation of interleukin-10 production | 4 | 0,00232256 | 0,02212227 | 0,01749864 | Xcl1, Sash3, Tlr2, Fcer1g |
| GO:0046942 | carboxylic acid transport | 13 | 0,00238742 | 0,02257191 | 0,0178543 | Slco2b1, Slc27a2, Cacna1a, Ntrk2, Slc6a20a, Thbs1, Per2, Edn1, Lrrc8c, Emb, Nfkbie, Agt, Slc15a1 |
| GO:0032608 | interferon-beta production | 5 | 0,00238843 | 0,02257191 | 0,0178543 | Ptpn22, Tlr2, Nlrc3, Tmem173, Zbtb20 |
| GO:0072678 | T cell migration | 5 | 0,00238843 | 0,02257191 | 0,0178543 | Ccl5, Xcl1, Spn, Itgb7, Myo1g |
| GO:0002260 | lymphocyte homeostasis | 6 | 0,00240155 | 0,02257832 | 0,01785937 | Cd74, Coro1a, Pik3cd, Dock10, Lat, Nckap1l |
| GO:0070228 | regulation of lymphocyte apoptotic process | 6 | 0,00240155 | 0,02257832 | 0,01785937 | Ccl5, Cd3g, Cd74, Serpinb9, Il7r, Cd27 |
| GO:2001056 | positive regulation of cysteine-type endopeptidase activity | 8 | 0,00241484 | 0,02264455 | 0,01791176 | Casp12, Fasl, Lck, Fyn, Cyfip2, Nek5, Tnfsf10, St18 |
| GO:0015849 | organic acid transport | 13 | 0,00245827 | 0,02299244 | 0,01818694 | Slco2b1, Slc27a2, Cacna1a, Ntrk2, Slc6a20a, Thbs1, Per2, Edn1, Lrrc8c, Emb, Nfkbie, Agt, Slc15a1 |
| mmu04060 | Cytokine-cytokine receptor interaction | 15 | 0,00375408 | 0,02331481 | 0,01799046 | 110075, 19116, 20304, 14103, 16963, 20303, 12778, 60504, 16186, 16185, 80901, 16197, 22035, 16994, 21940 |
| GO:0002283 | neutrophil activation involved in immune response | 3 | 0,00257329 | 0,02388351 | 0,01889177 | Itgb2, Myo1f, Fcer1g |
| GO:0002295 | T-helper cell lineage commitment | 3 | 0,00257329 | 0,02388351 | 0,01889177 | Tbx21, Spn, Ly9 |
| GO:0070571 | negative regulation of neuron projection regeneration | 3 | 0,00257329 | 0,02388351 | 0,01889177 | Rtn4rl1, Rtn4r, Thy1 |
| GO:0002715 | regulation of natural killer cell mediated immunity | 5 | 0,00259468 | 0,02402062 | 0,01900022 | Vav1, Klrk1, Cd96, Serpinb9, Klre1 |
| GO:0036230 | granulocyte activation | 4 | 0,00260657 | 0,0240408 | 0,01901618 | Ccl5, Itgb2, Myo1f, Fcer1g |
| GO:0050864 | regulation of B cell activation | 12 | 0,00261011 | 0,0240408 | 0,01901618 | Card11, Cd74, Tbx21, Inpp5d, Ptprc, Tbc1d10c, Il2rg, Sash3, Ikzf3, Nckap1l, Ikzf1, Cd27 |
| mmu04151 | PI3K-Akt signaling pathway | 17 | 0,00404576 | 0,02448206 | 0,01889116 | 18534, 19116, 15370, 18212, 21825, 14103, 269643, 320207, 14710, 16186, 16185, 18707, 24088, 16197, 16421, 11601, 71785 |
| GO:0031349 | positive regulation of defense response | 14 | 0,00275572 | 0,02531775 | 0,02002625 | Ccl5, Vav1, Cd74, Klrk1, Ptpn22, Xcl1, Tlr2, Txk, Cd300lf, Klre1, Peli3, Fcer1g, Tmem173, Gpsm3 |
| GO:0050922 | negative regulation of chemotaxis | 5 | 0,00281353 | 0,02571866 | 0,02034337 | Thbs1, Klrk1, Gpr18, St6gal1, Angpt2 |
| GO:2000401 | regulation of lymphocyte migration | 5 | 0,00281353 | 0,02571866 | 0,02034337 | Ccl5, Klrk1, Xcl1, Ccl4, Spn |

| | | | | | | |
|---|---|---|---|---|---|---|
| GO:0007187 | G protein-coupled receptor signaling pathway, coupled to cyclic nucleotide second messenger | 11 | 0,00289527 | 0,02623612 | 0,02075268 | Chga, Lgr5, Adgrg5, Edn1, Pde4b, Grm4, Glp1r, Fzd2, Adcy7, Sstr2, Oprd1 |
| GO:0043367 | CD4-positive, alpha-beta T cell differentiation | 6 | 0,0029127 | 0,02623612 | 0,02075268 | Tbx21, Spn, Il2rg, Sash3, Nckap1l, Ly9 |
| GO:0002701 | negative regulation of production of molecular mediator of immune response | 4 | 0,00291352 | 0,02623612 | 0,02075268 | Tbx21, Xcl1, Cd96, Vsir |
| GO:0033032 | regulation of myeloid cell apoptotic process | 4 | 0,00291352 | 0,02623612 | 0,02075268 | Ccl5, Pik3cd, St6gal1, Fcer1g |
| GO:0051084 | 'de novo' posttranslational protein folding | 4 | 0,00291352 | 0,02623612 | 0,02075268 | Cd74, H2-DMa, H2-DMb1, Dnajb4 |
| GO:2000516 | positive regulation of CD4-positive, alpha-beta T cell activation | 4 | 0,00291352 | 0,02623612 | 0,02075268 | Xcl1, Il2rg, Sash3, Nckap1l |
| GO:1901215 | negative regulation of neuron death | 11 | 0,00299201 | 0,02687621 | 0,02125899 | Chga, Cacna1a, Ccl5, Ntrk2, Nr4a3, Coro1a, Fyn, Nr4a2, Agap2, Glp1r, Agt |
| GO:0050880 | regulation of blood vessel size | 9 | 0,00306122 | 0,02736496 | 0,02164559 | Add3, Chga, Edn2, Dusp5, Per2, Edn1, Alox5, Ptgs1, Agt |
| GO:0032700 | negative regulation of interleukin-17 production | 3 | 0,00308412 | 0,02736496 | 0,02164559 | Vsir, Tlr2, Nckap1l |
| GO:0043373 | CD4-positive, alpha-beta T cell lineage commitment | 3 | 0,00308412 | 0,02736496 | 0,02164559 | Tbx21, Spn, Ly9 |
| GO:0097091 | synaptic vesicle clustering | 3 | 0,00308412 | 0,02736496 | 0,02164559 | Pclo, Pcdh17, Syn2 |
| GO:0140131 | positive regulation of lymphocyte chemotaxis | 3 | 0,00308412 | 0,02736496 | 0,02164559 | Ccl5, Xcl1, Ccl4 |
| GO:1903321 | negative regulation of protein modification by small protein conjugation or removal | 6 | 0,00309992 | 0,02743808 | 0,02170342 | Per2, Gbp4, Sh3rf2, Fyn, Peli3, Capn3 |
| GO:0035150 | regulation of tube size | 9 | 0,00318444 | 0,02811756 | 0,02224089 | Add3, Chga, Edn2, Dusp5, Per2, Edn1, Alox5, Ptgs1, Agt |
| GO:0006690 | icosanoid metabolic process | 7 | 0,00322559 | 0,02837475 | 0,02244432 | Edn2, Hpgds, Cd74, Edn1, Alox5, Tlr2, Ptgs1 |
| GO:0048511 | rhythmic process | 12 | 0,0032292 | 0,02837475 | 0,02244432 | Ntrk2, Edn2, Dpyd, Per2, Ptprn, Per1, Prkg2, Ciart, Prox1, Per3, Agt, Usp2 |
| GO:0006458 | 'de novo' protein folding | 4 | 0,00324428 | 0,02843836 | 0,02249464 | Cd74, H2-DMa, H2-DMb1, Dnajb4 |
| GO:0051091 | positive regulation of DNA-binding transcription factor activity | 11 | 0,00329793 | 0,028839 | 0,02281155 | Card11, Neurod1, Hmgn3, Edn1, Tlr2, Hcls1, Itgb2, Fzd2, Ror1, Capn3, Tmem173 |
| GO:0007188 | adenylate cyclase-modulating G protein-coupled receptor signaling pathway | 10 | 0,00337462 | 0,02943866 | 0,02328588 | Chga, Lgr5, Adgrg5, Edn1, Pde4b, Grm4, Glp1r, Adcy7, Sstr2, Oprd1 |
| GO:0046394 | carboxylic acid biosynthetic process | 14 | 0,00340031 | 0,02959166 | 0,0234069 | Slc27a2, Edn2, Hpgds, Cd74, Chst12, Per2, Fbp1, Edn1, Alox5, Ptgs1, Malrd1, Prox1, Agt, Zbtb20 |
| GO:0016053 | organic acid biosynthetic process | 14 | 0,0034889 | 0,03025136 | 0,02392872 | Slc27a2, Edn2, Hpgds, Cd74, Chst12, Per2, Fbp1, Edn1, Alox5, Ptgs1, Malrd1, Prox1, Agt, Zbtb20 |
| GO:0032945 | negative regulation of mononuclear cell proliferation | 6 | 0,00350112 | 0,03025136 | 0,02392872 | Inpp5d, Xcl1, H2-Ab1, Spn, H2-Aa, Vsir |
| GO:0050672 | negative regulation of lymphocyte proliferation | 6 | 0,00350112 | 0,03025136 | 0,02392872 | Inpp5d, Xcl1, H2-Ab1, Spn, H2-Aa, Vsir |
| GO:0070231 | T cell apoptotic process | 5 | 0,00355008 | 0,03059336 | 0,02419924 | Ccl5, Fasl, Serpinb9, Il7r, Cd27 |
| GO:0050708 | regulation of protein secretion | 17 | 0,00356473 | 0,03059336 | 0,02419924 | Cacna1e, Chga, Ccl5, Baiap3, Pde1c, Pde8b, Ptpn22, Hmgn3, Per2, Srgn, Tlr2, Cd2, Malrd1, Glp1r, Agt, Myt1, Myrip |
| GO:0043524 | negative regulation of neuron apoptotic process | 9 | 0,00357767 | 0,03059336 | 0,02419924 | Cacna1a, Ntrk2, Nr4a3, Coro1a, Fyn, Nr4a2, Agap2, Glp1r, Agt |
| GO:0001562 | response to protozoan | 4 | 0,00359971 | 0,03059336 | 0,02419924 | Gbp9, Spn, Iigp1, Gbp6 |
| GO:0016601 | Rac protein signal transduction | 4 | 0,00359971 | 0,03059336 | 0,02419924 | Farp2, Rac2, Arap3, Dock2 |
| GO:0045987 | positive regulation of smooth muscle contraction | 4 | 0,00359971 | 0,03059336 | 0,02419924 | Edn2, Edn1, Lck, Ptgs1 |
| GO:0051482 | positive regulation of cytosolic calcium ion concentration involved in phospholipase C-activating G protein-coupled signaling pathway | 4 | 0,00359971 | 0,03059336 | 0,02419924 | P2ry10, Gpr18, Edn1, Gpr55 |
| GO:0010810 | regulation of cell-substrate adhesion | 10 | 0,00361755 | 0,0306731 | 0,02426232 | Cd3e, Limch1, Thbs1, Rac2, Smoc2, Thy1, Skap1, St6gal1, Angpt2, Smoc1 |
| GO:0006909 | phagocytosis | 13 | 0,00364491 | 0,03075739 | 0,02432899 | Vav1, Hck, Thbs1, Ptprc, Dock2, Coro1a, Arhgap25, Tlr2, Itgb2, Cd300lf, Nckap1l, Myo1g, Fcer1g |
| GO:0051235 | maintenance of location | 13 | 0,00364491 | 0,03075739 | 0,02432899 | Grik5, Ptprc, Fasl, Xcl1, Abcg1, Coro1a, Thy1, Srgn, Lck, Fbn1, Glp1r, Pex5l, Capn3 |
| GO:0032695 | negative regulation of interleukin-12 production | 3 | 0,00365292 | 0,03075739 | 0,02432899 | Thbs1, Tlr2, Nlrc3 |
| mmu04927 | Cortisol synthesis and secretion | 6 | 0,00521792 | 0,03078575 | 0,02375529 | 15370, 18796, 218461, 58226, 11513, 11606 |
| GO:0007259 | JAK-STAT cascade | 9 | 0,00371689 | 0,03116676 | 0,0246528 | Prlr, Ccl5, Ptprc, Neurod1, Lck, Fyn, Hcls1, Agap2, Agt |
| GO:0051090 | regulation of DNA-binding transcription factor activity | 15 | 0,00371871 | 0,03116676 | 0,0246528 | Card11, Hck, Xcl1, Neurod1, Hmgn3, Edn1, Tlr2, Hcls1, Itgb2, Fzd2, Nlrc3, Prox1, Ror1, Capn3, Tmem173 |
| GO:2000146 | negative regulation of cell motility | 12 | 0,00374025 | 0,03127507 | 0,02473847 | Limch1, Ldlrad4, Thbs1, Klrk1, Gpr18, Arap3, Thy1, Ptprt, Arhgdib, Was, Angpt2, Olfm1 |
| GO:0034394 | protein localization to cell surface | 5 | 0,00382371 | 0,03189939 | 0,0252323 | Slc28a2, Cd247, Ric3, Kcnab2, Fcer1g |
| GO:0001782 | B cell homeostasis | 4 | 0,00398068 | 0,03290635 | 0,02602881 | Cd74, Pik3cd, Dock10, Nckap1l |
| GO:0002369 | T cell cytokine production | 4 | 0,00398068 | 0,03290635 | 0,02602881 | Tbx21, Xcl1, Vsir, Sash3 |
| GO:0032620 | interleukin-17 production | 4 | 0,00398068 | 0,03290635 | 0,02602881 | Vsir, Tlr2, Nckap1l, Ly9 |
| GO:0033028 | myeloid cell apoptotic process | 4 | 0,00398068 | 0,03290635 | 0,02602881 | Ccl5, Pik3cd, St6gal1, Fcer1g |
| mmu05164 | Influenza A | 10 | 0,00576519 | 0,03318498 | 0,02560661 | 20304, 14998, 14999, 14103, 14961, 14969, 14960, 12265, 18707, 22035 |
| GO:0046579 | positive regulation of Ras protein signal transduction | 5 | 0,0041121 | 0,03386721 | 0,02678885 | Rtn4r, P2ry10, Gpr18, Dock2, Gpr55 |
| GO:0006865 | amino acid transport | 8 | 0,00411558 | 0,03386721 | 0,02678885 | Cacna1a, Ntrk2, Slc6a20a, Per2, Lrrc8c, Nfkbie, Agt, Slc15a1 |

| | | | | | | |
|---|---|---|---|---|---|---|
| GO:0051656 | establishment of organelle localization | 17 | 0,00423181 | 0,03474489 | 0,02748309 | Chga, Cacna1a, Grik5, Pclo, Tacc1, Nrn1, Rac2, Nr4a3, Sept1, Grm4, Lat, Lat2, Map1b, Fcer1g, Myrip, Kif1a, Syp |
| GO:0002363 | alpha-beta T cell lineage commitment | 3 | 0,0042816 | 0,03476045 | 0,02749539 | Tbx21, Spn, Ly9 |
| GO:0002716 | negative regulation of natural killer cell mediated immunity | 3 | 0,0042816 | 0,03476045 | 0,02749539 | Cd96, Serpinb9, Klre1 |
| GO:0006699 | bile acid biosynthetic process | 3 | 0,0042816 | 0,03476045 | 0,02749539 | Slc27a2, Malrd1, Prox1 |
| GO:0030852 | regulation of granulocyte differentiation | 3 | 0,0042816 | 0,03476045 | 0,02749539 | Inpp5d, Hcls1, Ikzf1 |
| GO:0031579 | membrane raft organization | 3 | 0,0042816 | 0,03476045 | 0,02749539 | Ptprc, Dock2, Clec2i |
| GO:0051588 | regulation of neurotransmitter transport | 9 | 0,00431675 | 0,03496761 | 0,02765926 | Cacna1a, Baiap3, Ntrk2, Grik5, Nrn1, Per2, Grm4, Fev, Syp |
| GO:2000403 | positive regulation of lymphocyte migration | 4 | 0,00438801 | 0,03546564 | 0,0280532 | Ccl5, Xcl1, Ccl4, Spn |
| GO:0070664 | negative regulation of leukocyte proliferation | 6 | 0,00441766 | 0,03562601 | 0,02818005 | Inpp5d, Xcl1, H2-Ab1, Spn, H2-Aa, Vsir |
| mmu05163 | Human cytomegalovirus infection | 13 | 0,00670185 | 0,03765799 | 0,02905813 | 20304, 15043, 18796, 15024, 14103, 19354, 20303, 14710, 14963, 18707, 667803, 11513, 72512 |
| mmu05150 | Staphylococcus aureus infection | 7 | 0,00690167 | 0,03787892 | 0,02922861 | 14998, 14999, 20345, 14961, 14969, 14960, 16414 |
| GO:0050856 | regulation of T cell receptor signaling pathway | 4 | 0,00482251 | 0,03871875 | 0,03062639 | Ptpn22, Thy1, Lck, Trat1 |
| GO:1900015 | regulation of cytokine production involved in inflammatory response | 4 | 0,00482251 | 0,03871875 | 0,03062639 | Per1, Nlrc3, Adcy7, Gpsm3 |
| GO:0003013 | circulatory system process | 17 | 0,00489673 | 0,0392279 | 0,03102913 | Cacna1e, Add3, Chga, Dll1, Edn2, Dusp5, Per2, Ccl4, Edn1, Alox5, Pde4b, Fli1, Ptgs1, Fyn, Glp1r, Agt, Adamts16 |
| GO:0019226 | transmission of nerve impulse | 6 | 0,00493709 | 0,03946407 | 0,03121594 | Cacna1e, Cacna1a, Ntrk2, Cacna1h, Agt, Scn3a |
| GO:0002313 | mature B cell differentiation involved in immune response | 3 | 0,00497193 | 0,03948167 | 0,03122986 | Dll1, Dock10, Mfng |
| GO:0033033 | negative regulation of myeloid cell apoptotic process | 3 | 0,00497193 | 0,03948167 | 0,03122986 | Ccl5, St6gal1, Fcer1g |
| GO:0072148 | epithelial cell fate commitment | 3 | 0,00497193 | 0,03948167 | 0,03122986 | Dll1, Neurod1, Prox1 |
| mmu05020 | Prion diseases | 4 | 0,00763141 | 0,04093213 | 0,03158456 | 20304, 12364, 14360, 12279 |
| GO:0045823 | positive regulation of heart contraction | 4 | 0,00528498 | 0,04187594 | 0,03312373 | Chga, Edn2, Edn1, Glp1r |
| GO:0043254 | regulation of protein complex assembly | 16 | 0,00538026 | 0,04253801 | 0,03364742 | Add3, Ldlrad4, Hck, Farp2, Coro1a, Skap1, Tlr2, Hcls1, Nlrc3, Was, Nckap1l, Oprd1, Ikzf1, Bmf, Map1b, Clec2i |
| GO:0050766 | positive regulation of phagocytosis | 5 | 0,00542194 | 0,04277439 | 0,03383439 | Ptprc, Dock2, Cd300lf, Nckap1l, Fcer1g |
| GO:0040013 | negative regulation of locomotion | 13 | 0,0055304 | 0,04353537 | 0,03443633 | Limch1, Ldlrad4, Thbs1, Klrk1, Gpr18, Arap3, Thy1, Ptprt, Arhgdib, St6gal1, Was, Angpt2, Olfm1 |
| GO:0050871 | positive regulation of B cell activation | 10 | 0,00555388 | 0,04362559 | 0,03450769 | Card11, Cd74, Tbx21, Inpp5d, Ptprc, Il2rg, Sash3, Nckap1l, Ikzf1, Cd27 |
| mmu04924 | Renin secretion | 6 | 0,00832721 | 0,04367157 | 0,0336984 | 13615, 18575, 18796, 13614, 19092, 11606 |
| GO:0002834 | regulation of response to tumor cell | 3 | 0,00572551 | 0,04439719 | 0,03511803 | Klrk1, Xcl1, Klre1 |
| GO:0002837 | regulation of immune response to tumor cell | 3 | 0,00572551 | 0,04439719 | 0,03511803 | Klrk1, Xcl1, Klre1 |
| GO:0042533 | tumor necrosis factor biosynthetic process | 3 | 0,00572551 | 0,04439719 | 0,03511803 | Thbs1, Spn, Tlr2 |
| GO:0042534 | regulation of tumor necrosis factor biosynthetic process | 3 | 0,00572551 | 0,04439719 | 0,03511803 | Thbs1, Spn, Tlr2 |
| GO:0043369 | CD4-positive or CD8-positive, alpha-beta T cell lineage commitment | 3 | 0,00572551 | 0,04439719 | 0,03511803 | Tbx21, Spn, Ly9 |
| GO:2000482 | regulation of interleukin-8 secretion | 3 | 0,00572551 | 0,04439719 | 0,03511803 | Ptpn22, Tlr2, Cd2 |
| GO:0030239 | myofibril assembly | 5 | 0,00579058 | 0,04480601 | 0,0354414 | Acta1, Edn1, Myom3, Prox1, Capn3 |
| GO:2000116 | regulation of cysteine-type endopeptidase activity | 10 | 0,00591258 | 0,04565264 | 0,03611108 | Nr4a1, Casp12, Thbs1, Fasl, Lck, Fyn, Cyfip2, Nek5, Tnfsf10, St18 |
| GO:0007269 | neurotransmitter secretion | 9 | 0,00594202 | 0,04568561 | 0,03613716 | Cacna1a, Baiap3, Ntrk2, Grik5, Pclo, Nrn1, Grm4, Syn2, Syp |
| GO:0099643 | signal release from synapse | 9 | 0,00594202 | 0,04568561 | 0,03613716 | Cacna1a, Baiap3, Ntrk2, Grik5, Pclo, Nrn1, Grm4, Syn2, Syp |
| GO:0043523 | regulation of neuron apoptotic process | 11 | 0,00603518 | 0,04630374 | 0,0366261 | Cacna1a, Ntrk2, Grik5, Fasl, Nr4a3, Coro1a, Fyn, Nr4a2, Agap2, Glp1r, Agt |
| GO:0046888 | negative regulation of hormone secretion | 6 | 0,00610848 | 0,04676723 | 0,03699272 | Chga, Edn2, Pde1c, Pde8b, Edn1, Ptgs1 |
| GO:0032956 | regulation of actin cytoskeleton organization | 13 | 0,006412 | 0,04898771 | 0,03874911 | Add3, Limch1, Hck, Edn1, Coro1a, Tlr2, Arhgdib, Hcls1, Myo1f, Prox1, Was, Nckap1l, Clec2i |
| GO:0006631 | fatty acid metabolic process | 14 | 0,00654041 | 0,04957732 | 0,03921549 | Pck1, Slc27a2, Ces1d, Edn2, Hpgds, Cd74, Eci3, Per2, Edn1, Alox5, Nr4a3, Ptgs1, Agt, Acsbg1 |
| GO:0014850 | response to muscle activity | 3 | 0,00654382 | 0,04957732 | 0,03921549 | Fndc5, Agt, Capn3 |
| GO:0060749 | mammary gland alveolus development | 3 | 0,00654382 | 0,04957732 | 0,03921549 | Prlr, Tph1, Agap2 |
| GO:0061377 | mammary gland lobule development | 3 | 0,00654382 | 0,04957732 | 0,03921549 | Prlr, Tph1, Agap2 |
| GO:0051057 | positive regulation of small GTPase mediated signal transduction | 5 | 0,0065802 | 0,04974906 | 0,03935133 | Rtn4r, P2ry10, Gpr18, Dock2, Gpr55 |

**Supplemental Table 5 :** Description of the 11 colon data sets included in the analysis of *CCNA2* expression in CRC patients

| Reference | Source of data | Technological platform | N° of probe sets/genes | N° of samples | Primary (N) | Normal (N) | Metastasis (N) |
|---|---|---|---|---|---|---|---|
| Jorissen et al.,<br>Clin Cancer Res 2009 | GEO database,<br>GSE14333 | Affymetrix,<br>array U133 Plus 2.0 | 54K | 251 | 251 | | |
| Sheffer et al.,<br>Proc Natl Acad Sci 2009 | GEO database,<br>GSE41258 | Affymetrix,<br>array U133 A | 22K | 289 | 168 | 54 | 67 |
| Staub et al.,<br>J Mol Med (Berl) 2009 | GEO database,<br>GSE12945 | Affymetrix,<br>array U133 Plus 2.0 | 54K | 29 | 29 | | |
| Smith et al.,<br>Gastroenterology 2010 | GEO database,<br>GSE17538 | Affymetrix,<br>array U133 Plus 2.0 | 54K | 232 | 232 | | |
| de Sousa et al.,<br>Cell Stem Cell 2011 | GEO database,<br>GSE33113 | Affymetrix,<br>array U133 Plus 2.0 | 54K | 90 | 90 | | |
| Kenned et al.,<br>J Clin Oncol 2011 | Array-Express database,<br>E-MTAB-863 | Affymetrix custom array,<br>ADXCRCG2a520319 | 62K | 215 | 215 | | |
| Sveen et al.,<br>Genome Med 2011 | GEO database,<br>GSE24551 | Affymetrix,<br>Human Exon 1.0 ST array | 22K | 160 | 160 | | |
| Laibe et al.,<br>OMICS 2012 | GEO database,<br>GSE37892 | Affymetrix,<br>array U133 Plus 2.0 | 54K | 130 | 130 | | |
| Marisa et al.,<br>PLoS Med 2013 | GEO database,<br>GSE39582 | Affymetrix,<br>array U133 Plus 2.0 | 54K | 455 | 455 | | |
| TCGA, COAD | TCGA portal,<br>https://tcga-data.nci.nih.gov | Illumina,<br>RNA sequencing V2 | 25K | 500 | 459 | 41 | |
| IPC | ---- submission to ArrayExpress ongoing ---- | Affymetrix,<br>array U133 Plus 2.0 | 54K | 50 | 50 |  |  |
| | Total | | | 2401 | 2239 | 95 | 67 |
